# Consensus Through Diversity: A Comprehensive Benchmark of Multi-Omic Approaches for Precision Breast Oncology

**DOI:** 10.64898/2026.04.17.719159

**Authors:** Aristeidis Sionakidis, Karen Pinilla Alba, Jean Abraham, Nikola Simidjievski

**Affiliations:** Department of Oncology, University of Cambridge, Hills Road, Cambridge CB2 0QQ, United Kingdom; Télécom Paris, Institut Polytechnique de Paris, 19 Pl. Marguerite Perey, Paris 91120, France

**Keywords:** multi-omics, subtyping, consensus, breast cancer, integration, benchmark

## Abstract

Emerging multi-omic profiling has made it feasible to subtype disease using multiple molecular layers. However, inconsistent preprocessing, heterogeneous implementations, variable evaluation, and limited reproducibility often constrain method selection. Here, we systematically benchmark 22 publicly available unsupervised approaches for bulk data on the TCGA-BRCA cohort across five modalities (RNA-seq, miRNA, DNA methylation, copy numbers, single nucleotide polymorphisms) and validate findings in two independent datasets, enabling a multi-layered comparison of performance, heterogeneous data support and interpretability. Most approaches fuse multi-omic data to produce a two-cluster solution largely aligned with ER status, with higher-resolution approaches further refining these into four coherent subclasses (angiogenic luminal, oxidative-phosphorylation/HER2-low luminal, immune-inflamed basal-like, and hyper-proliferative basal-like). Our benchmarking results indicate that methods based on similarity networks can efficiently produce stable, reliable partitions. Matrix factorisation and Bayesian factorisation algorithms produce rich latent representations, allowing quantification of feature and modality contributions, albeit at higher computational cost. Consensus clustering can be used on a case-by-case basis and refine partitions into more robust and generalisable findings. We aggregate our insights into a decision workflow that aligns with study goals, data characteristics, and computational resources, enabling optimal analytic strategies. This comprehensive assessment provides a practical roadmap for investigators seeking to extract reproducible, biologically meaningful subtypes from complex multi-omic datasets. We higlight the different technical and practical benefits and trade-offs that shape the selection and development of multi-omic approaches applied in precision oncology.

## 1 Background

Recent multi-omic technologies have revolutionised biomedical research, producing high-quality heterogeneous data at different scales [1]. These capabilities provide an unprecedented opportunity to unravel the complex molecular mechanisms underlying diseases such as cancer [1]. By integrating data modalities across different molecular layers, such as genomics, transcriptomics, proteomics, and epigenomics, researchers can gain a more holistic view of biological processes [1, 2]. These approaches enable a deeper understanding of disease biology, as integrated multi-omic data analysis can often reveal interactions and regulatory pathways that otherwise would have remained obscure if each -omic modality had been analysed individually [1–5]. Recent approaches to integrative multi-omic analyses are capable of addressing many computational challenges inherent to the scale and complexity of multi-omic datasets [1–3], thus allowing the discovery of biologically meaningful patterns that could inform diagnosis, prognosis and treatment [1, 2, 6].

Multi-omic analyses have been successfully applied to a wide range of tasks, including the identification of disease biomarkers, elucidation of intricate molecular pathways, prediction of outcome and disease subtyping [1, 2, 4, 7, 8]. More specifically, disease subtyping has gained significant attention in the past two decades, becoming a cornerstone of computational biology, bioinformatics and molecular genetics [9, 10]. In cancer research, subtyping based on multi-omic profiles has proven especially valuable for uncovering distinct molecular subgroups correlated with clinical outcomes, which could guide personalised treatment strategies [6, 10]. The ability to stratify patients into molecularly defined subgroups facilitates the development of targeted therapies and improves the understanding of tumour heterogeneity [6].

Despite these advances, several critical challenges remain that may often limit the translational significance of recent multi-omic approaches. Current studies often employ inconsistent data processing pipelines and evaluation strategies, with methods tested on disparate datasets, limiting objective performance assessment [11]. Recent reviews and comparative studies lack standardised empirical benchmarking frameworks with consistent evaluation criteria, data preprocessing protocols and parameter tuning strategies. In particular, existing reviews of multi-omic subtyping approaches primarily focus on the theoretical foundations and the classification of the discussed methods into distinct groups [1–5, 10]. Some other reviews have constrained their analysis by applying only a limited selection of clustering algorithms to the same collection of datasets and evaluating them [12, 13]. These limitations impede fair comparisons among multi-omic approaches and undermine reproducibility across studies. Moreover, existing work provides limited practical guidance on method selection, parameter tuning, appropriate data-type combinations, and the real-world implications of computational complexity.

In this work, we aim at addressing the aforementioned shortcomings and present a comprehensive review of tools and methodologies focusing on unsupervised subtyping of multiomic data. Specifically, we evaluate the tools’ methodological foundations, computational demands, robustness and stability, capacity to integrate diverse data types, interpretability of outputs, and alignment with existing biological knowledge. We establish explicit selection criteria for the algorithms incorporated in our analysis and provide a detailed justification for the parameters we assess during algorithm tuning, as well as those we keep fixed. Furthermore, we evaluate each method’s capability to handle binary data, such as Single Nucleotide Polymorphisms (SNPs), independently and/or with reasonable/minor modifications. Our work delivers a broad spectrum of downstream results, enabling more detailed comparisons across different algorithms. We investigate when and whether consensus strategies offer superior performance compared to a meticulously optimised single method, thereby providing deeper insights into the effectiveness of multi-omic subtyping approaches. We highlight the strengths and limitations of various methods, offering insights into best practices for multi-omic data integration. Finally, we discuss current challenges and future directions in the field, with a focus on improving the robustness and biological relevance of subtyping results.

Our contributions can be summarised as follows:

- We benchmark 22 popular multi-omic integration methods on the TCGA-BRCA dataset using five omic layers.
- We evaluate the selected methods in terms of scalability, robustness, stability, interpretability, plausibility, capability to handle heterogenous data and user-friendliness.
- We examine consensus approaches and when they can be used to refine analysis results.
- We provide a straightforward workflow for deciding the optimal multi-omic analysis strategy.

## Results

### Baseline analysis

For our baseline multi-omic clustering analysis, we used 1) an analysis of the two groups produced by estrogen receptor (ER) status classification (ER+ vs. ER-tumours) and 2) a consensus multi-omic analysis with ten multi-omic integration methods with fixed parameters, as these are implemented in the *MOVICS* package in R [14]: Cancer Integration via Multikernel LeaRning (CIMLR) [15], iClusterBayes [16], moCluster [17], Cluster Of Cluster Assignments (COCA) [18], Consensus Clustering [19], Integrative Non-negative Matrix Factorization (IntNMF) [20], Low Rank Approximation clustering (LRAcluster) [21], NEighborhood-based Multi-Omics clustering (NEMO) [22], Perturbation clustering for data INtegration and disease Subtyping (PINSPlus) [23] and Similarity Network Fusion (SNF) [24]. A subset of these methods (CIMLR [15], iClusterBayes [16], COCA [18], LRAcluster [21], NEMO [22], and SNF [24]) were then additionally evaluated along with methods retrieved from the literature: association-signal-annotation boosted Similarity Network Fusion (ab-SNF) [25], Affinity Network Fusion (ANF) [26], Kernel Learning Integrative Clustering (KLIC) [27], Multi-omics Data Integration for Clustering to identify Cancer subtypes (MDICC) [28], Multiple Factor Analysis (MFA) [29], Multi-Omics Factor Analysis (MOFA) [30], Multi Omic clustering by Non-Exhaustive Types (MONET) [31], Multiple Similarity Network Embedding (MSNE) [32], Random Walk with Restart for multi-dimensional data Fusion (RWR-F) [33], Random Walk with Restart and Neighbor information-based multi-dimensional data Fusion (RWR-NF) [33], Spectrum [34] and weight-boosted Multi-Kernel Learning (wMKL) [35].

Table 1 summarises the methods examined in this work and distinguishes between those included in the baseline analysis, individual runs or both scenarios. For more details on data preprocessing, method selection, tuning and post-hoc analysis see Sections Data characteristics and preprocessing, Exclusion of methods from individual runs, Individual method multi-omic clustering and General pipeline. For details on benchmarking methods with respect to robustness, stability and scalability, see Section Benchmarks. The general pipeline followed in the baseline analysis, individual runs and the consensus analysis is described in Section General pipeline and summarised in Figure 1. Supplementary material (methodology, results, tables and figures) can be found in Supplementary File 1. We also encourage readers to interactively explore results from this survey using the user-friendly R Shiny app we designed (https://github.com/sionaris/MO_survey_Shiny).

**Table 1.** Summary of methods.

| Category | Method | Software | Usage | Reference |
| --- | --- | --- | --- | --- |
| **Similarity Network** | ab-SNF | R | Trials* | Ruan et al.[25] |
|  | ANF | R | Trials | Ma et al.[26] |
|  | MDICC | R & Python | Trials | Yang et al.[28] |
|  | MSNE | Python | Trials | Xu et al.[32] |
|  | NEMO | R | Both | Rappoport et al.[22] |
|  | RWR-(N)F | R | Trials | Wen et al.[33] |
|  | SNF | R | Both | Wang et al.[24] |
|  | Spectrum | R | Trials | John et al.[34] |
| **Multiple Kernels** | CIMLR | R | Both | Ramazzotti et al.[15] |
|  | wMKL | R | Trials | Cao et al.[35] |
|  | KLIC | R | Trials | Cabassi et al.[27] |
| **Matrix Factorisation** | IntNMF | R | Baseline | Chalise et al.[20] |
|  | LRAcluster | R | Both | Wu et al. [21] |
|  | MFA | R | Trials | Lê et al.[36] |
|  | moCluster | R | Baseline | Meng et al.[17] |
|  | MOFA | R & Python | Trials | Argelaguet et al.[30] |
| **Graph-based learning** | MONET | Python | Trials | Rappoport et al.[31] |
| **Bayesian learning** | iClusterBayes | R | Both | Mo et al.[16] |
| **Ensemble Clustering** | PINSPlus | R | Baseline | Nguyen et al.[23, 37, 38] |
|  | COCA | R | Both | Various[18, 39] |
|  | Consensus Clustering (original) | R | Baseline | Monti et al.[19] |

\***Trials:** refers to methods not used in the baseline analysis.

**Figure 1.**
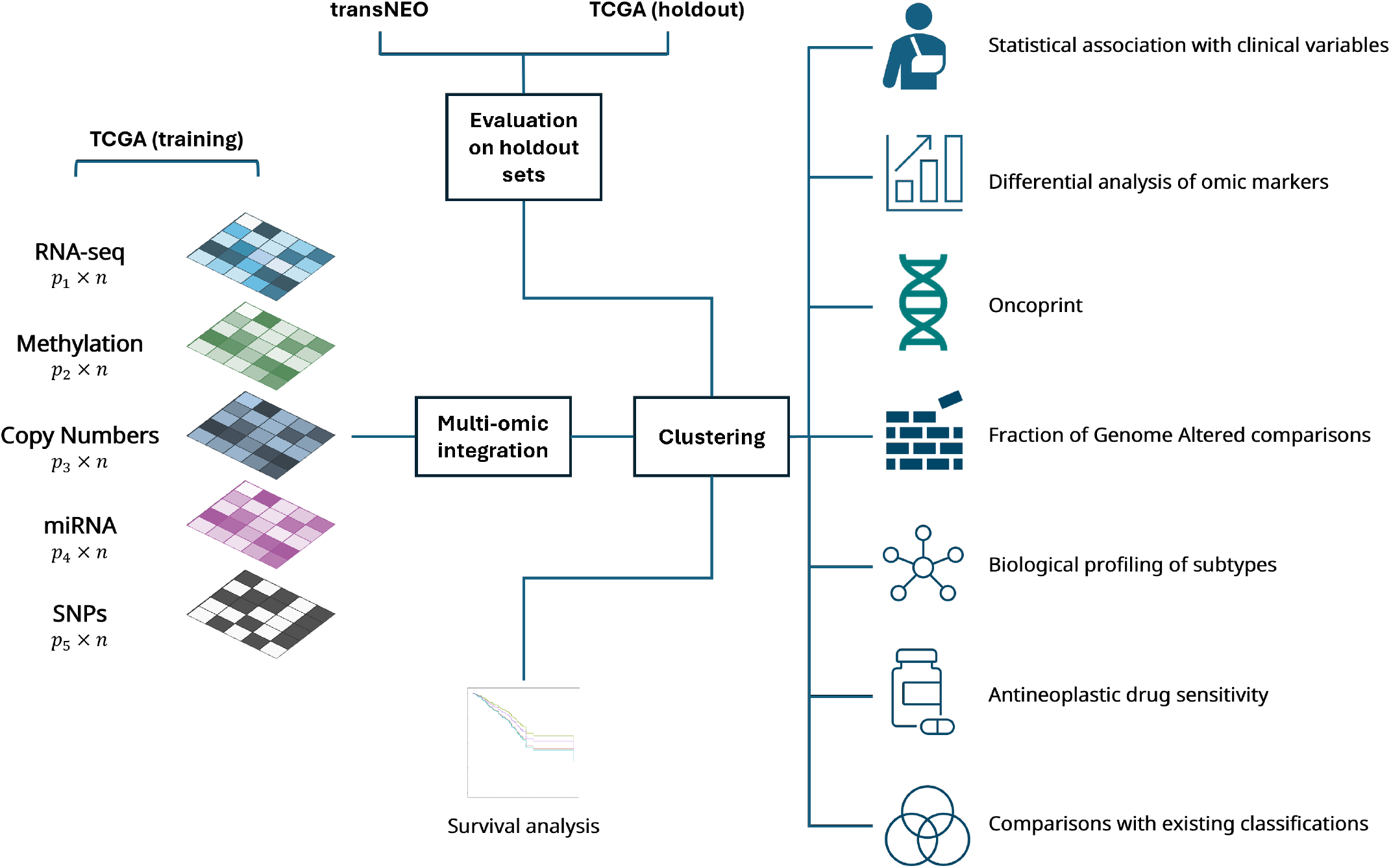
General multi-omic integration pipeline. The five modalities (distinct dimensionalities: *p*_*i*_; same sample set: *n*) are integrated and clustered using one of the selected methods (or an ensemble of methods in consensus approaches). The obtained clusters are examined (in the training cohort) for predicted drug sensitivity, distinct mutational and pathway patterns and agreement with established tumour classification systems. Differences in Kaplan-Meier curves between clusters, and associations with clinical variables are examined in the training cohort and both holdout sets.

### ER baseline analysis

ER status baseline analysis returned the results expected to be seen when a clustering produces two pure clusters: i) ER+ samples, ii) ER-samples. Regarding mutations (Figure 2e), the ER+ subset was characterised by higher proportions of *PIK3CA* (38.3% vs. 13.1% in ER-, *p*_*adj*_ = 6.8 ·10^−8^) and *CDH1* mutations (10.3% vs. 1.5% in ER-, *p*_*adj*_ = 1.7 ·10^−3^), while the ER-subset demonstrated higher proportions of *TP53* mutations (60.8% vs. 18.4% in ER+, *p*_*adj*_ = 1.3 ·10^−18^). In terms of Fraction of Genome Altered (FGA) (Figure 2c), a significantly larger fraction of the genome appeared to be altered in the ERsubset, which also demonstrated a significantly higher proportion of Fraction of Genome Lost (FGL), using the Catalogue Of Somatic Mutations In Cancer (COSMIC) criteria (https://cancer.sanger.ac.uk/cosmic/help/cnv/overview). ER+ tumours were enriched for cilium organisation and cytoskeleton-dependent intracellular and vesicle-transport programmes, whereas ER-tumours showed strong enrichment of DNA replication, mitotic cell-cycle, ribonucleoprotein and messenger RNA (mRNA) processing pathways, together with adaptive and innate immune response and cytokine-production signatures (Supplementary Figure 2). In other words, the baseline ER+ versus ER-contrast is dominated by a ciliated/secretory phenotype on the ER+ side and a highly proliferative, immune-active phenotype on the ER-side.

**Figure 2.**
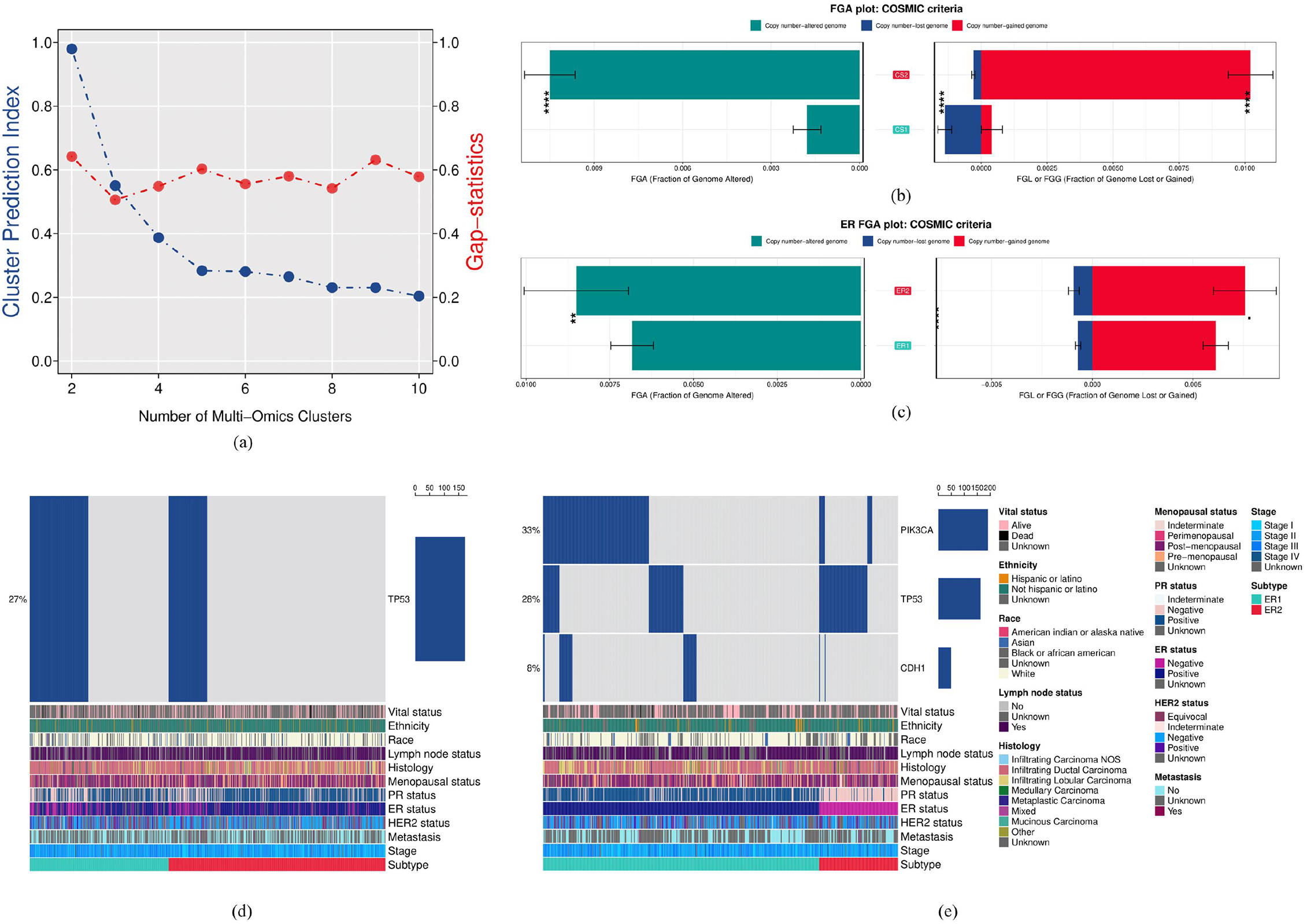
Baseline MOVICS analysis. **(a)** Optimal number of clusters determined by *MOVICS*. **(b)** Fraction of Genome Altered plot: COSMIC criteria - *MOVICS* CS subtypes **(c)** Fraction of Genome Altered plot: COSMIC criteria - ER subtypes. **(d)** Oncoprint plot: *MOVICS* CS subtypes. **(e)** Oncoprint plot: ER subtypes. Sample annotations at the bottom right.

### MOVICS baseline analysis

*MOVICS* provides a multi-omic integration suite of ten popular tools (marked as “Baseline” or “Both” in Table 1). *MOVICS* identified 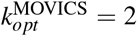 as the optimal number of multi-omic clusters in the training cohort (Figure 2a). All ten methods were then run with default settings (regardless of the distributional properties of the multi-omic data input) for a predetermined number of clusters 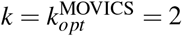. Consensus clustering on the clusters identified by each method produced the final two consensus subtypes (CS1 and CS2) that were used as the *MOVICS* baseline.

The clearest separation with respect to the original omics matrices is observed in Copy Number Variant (CNV) data (Supplementary Figure 3j). Samples in the two consensus subtypes are well-separated according to pair-wise Euclidean distances (calculated using the consensus matrix) and the corresponding silhouette metrics (Supplementary Figure 3b,c). When compared to other tumour classifications, there is significant association between the consensus subtypes and tumour stage (*p* = 0.039), ER (*p* < 0.001), progesterone receptor (PR) (*p* = 0.001) and Human Epidermal growth factor Receptor 2 (HER2) (*p* < 0.001) subtypes (Supplementary Figure 3d), without, however, approaching pure separations. Consensus subtypes are also significantly associated with histological type (*p <* 0.001).

There are significantly different FGA patterns between the two consensus subtypes (Figure 2b), as defined by COSMIC criteria. CS1 is associated with up-regulation of cell cycle pathways compared to CS2, while CS2 is associated with increased chemotaxis, extracellular matrix and angiogenesis activity (Supplementary Figure 3i). CS1 has a significantly higher proportion of *TP53* mutations compared to CS2 (42.2.% vs. 17.8%, *p*_*adj*_ = 4.31 · 10^−10^; Figure 2d).

### Individual runs

The output clusterings of all methods, for all cohorts can be found in Supplementary File 2.

#### Multi-omic profiles

Similarity network-fusion algorithms (ab-SNF, ANF, NEMO, RWR-NF, SNF and Spectrum) converged on highly concordant two-cluster solutions. The partitions separate ER+ from ER-tumours and are associated with significant differences in: i) *PIK3CA, TP53* and occasionally *CDH1* mutation frequencies, ii) FGA - predominantly driven by FGL - in the ER clusters, iii) expression profiles and iv) methylation patterns (Supplementary Figures 5-12). RWR-F also yields two clusters: one predominantly ER+ and one containing a mixture of ER+/-samples (*p <* 10^−5^; Supplementary Figure 15a). No significant associations are detected for HER2 status, tumour stage, metastatic status, menopausal status or lymph-node presence; the latent structure is driven chiefly by CNV data (Supplementary Figure 15b,c). MDICC produces two markedly unbalanced clusters (607/18 samples) without noteworthy molecular distinctions. By contrast, MSNE returns five well-resolved clusters that differ across expression, CNVs, methylation and microRNAs (miRNAs) (Supplementary Figure 13a-d). MSNE1/4 are mainly ER-, whereas MSNE2/3/5 are predominantly ER+. Within the ER+ group, MSNE3 shows elevated *TP53* mutations (31.4%) relative to MSNE2 (9.8%) and MSNE5 (7.6%), while the *PIK3CA* mutation rates are higher in MSNE3/5 (MSNE2 25.9%, MSNE3 41.3%, MSNE5 51.7%; see Supplementary Figures 13e,f). MSNE3 is also characterised by the highest FGL across MSNE clusters and no Fraction of Genome Gained (FGG) (Supplementary Figure 13g).

With respect to multiple-kernel learners, CIMLR identifies two subtypes that separate ER+ and ER-samples with statistical significance (*p <* 10^−5^), but none of the two clusters approaches purity (Supplementary Figure 16a). The cluster which is richer in ER-tumours, (CIMLR2), exhibits significantly higher mutation frequencies in *TTN, MUC16, RYR2, HMCN1, SYNE1, FLG* and *TP53* (Supplementary Figure 16d) and a statistically significantly higher FGL (Supplementary Figure 16c). wMKL distinguishes eight subtypes with distinct combinations of receptor status and mutation profiles (*TP53, PIK3CA* and *RYR2*) (Supplementary Figure 20a,b,d). wMKL4 is mainly ER- and harbours frequent *TP53* (62.0%), as well as characteristic methylation, miRNA and RNA sequencing (RNA-seq) profiles (Supplementary Figure 19). Most remaining ER-tumours fall into wMKL8, and also have a high rate of *TP53* mutations (Supplementary Figure 20d). The remaining six clusters are strongly ER+. wMKL1 and wMKL6 are highly mutated in *PIK3CA* (62.5%, 66.7% respectively), wMKL3 carries the highest proportion of *RYR2* mutations (13.3%). wMKL6 and wMKL7 are predominantly ER+, but are additionally mildly enriched for HER2+ tumours (26.9% and 25.0%, respectively). A low proportion (∼11%) of wMKL6 HER2+ samples carry *TP53* mutations, whereas in wMKL7 HER2+ samples show a 75/25% *TP53*-mutant/wild-type split. In terms of FGA, wMKL3/6 are characterised by relatively high FGL, while wMKL1/4/7/8 are characterised by relatively high FGG (Supplementary Figure 20c). In the kernel-Principal Component Analysis (PCA) projection of the wMKL similarity matrix, wMKL2 and wMKL6 appear as outlier clusters, whereas wMKL2/3/4/6 separate from the other subtypes in low-dimensional CNV space owing to higher copy-numbers (Supplementary Figure 20e,f). Finally, in KLIC, two largely imbalanced clusters are uncovered, with the main differences observed in RNA-seq and methylation profiles, and a significant (*p <* 10^−5^), yet not pure separation of ER status across clusters (Supplementary Figures 17,18).

Bayesian and matrix-factorisation methods generated modality-balanced latent spaces. MOFA yields three clusters that capture the most variance in CNV, RNA-seq and methylation data (Supplementary Figure 31). MOFA3 is characterised by significantly higher FGL compared to MOFA1/2 (*p <* 10^−3^, Supplementary Figure 32c). MFA also produces three clusters, distinguished by *PIK3CA* and *CDH1* mutation patterns (Supplementary Figure 24) and clearly separated across all five modalities in dimensionality-reduction plots (Supplementary Figure 25). LRAcluster finds four subtypes, two largely ER+ and two largely ER-, with distinct *TP53* mutation profiles (Supplementary Figure 22a,e). LRAcluster4 contains an elevated proportion of HER2+ samples (26.5%, *p <* 0.001) and is characterised by significantly high FGL (Supplementary Figure 22b,c). iClusterBayes assigns tumours to five clusters (two predominantly ER- and three predominantly ER+) that are well separated in low-dimensional CNV space and exhibit distinct *TP53* mutation landscapes (Supplementary Figure 37).

From the remaining methods, MONET uncovers three molecular subtypes significantly associated with ER status (*p <* 10^−5^), HER2 status (*p* = 0.0089), lymph-node presence (*p* = 0.0042) and tumour stage (*p* = 0.0019); see Supplementary Figure 39. MONET1 is mainly ER-, whereas MONET2 and MONET3 are ER+. The two ER+ clusters share similar methylation and RNA-seq profiles, but differ with respect to CNV and, to a lesser extent, miRNA and SNPs. MONET1 diverges from MONET2 across all omics layers; MONET1 and MONET3 differ mainly in RNA-seq and methylation, while remaining comparable in CNV and SNP patterns (Supplementary Figures 40-42). Mutation frequencies for *TP53* (61.0%, 7.2%, 23.8%), *PIK3CA* (16.4%, 51.0%, 22.7%) and *CDH1* (4.5%, 14.4%, 2.2%) are characteristic of different clusters (*p*_*adj*_ *<* 10^−15^, 4.04 · 10^−15^ and 6.37 · 10^−6^, respectively; see Supplementary Figure 40b). Finally, COCA clusters are driven largely by CNV variation and are only significantly associated with HER2 status (*p* = 0.01). No significant association with mutations or other clinical variables is observed.

#### Biological profiles of multi-omic clusters

Most algorithms (17/18; exception: COCA) converge on a common biological signature that includes enrichment for DNA double-strand-break repair, nucleotide metabolism, cell-cycle progression and coordinated regulation of leukocyte activity and apoptosis. When the partitions align well with ER status, ER+ clusters are characterised by up-regulated hormone-mediated signalling, vesicle-mediated intracellular and extracellular transport, endothelial proliferation and angiogenesis, whereas ER-clusters are dominated by mitotic programmes: chromosome segregation, DNA replication, checkpoint signalling, and by homologous recombination repair pathways (Supplementary Figures 12, 15d, 16b).

Three-cluster solutions (MOFA, MFA, MONET) show a recurrent pattern: one ER-subset (MOFA2, MFA3, MONET1) marked by increased cell-cycle activity and immune activation, and two ER+ subsets: one angiogenic/extracellular-matrix-driven (MOFA1, MFA1, MONET2) subset and one enriched for mitochondrial bioenergetics and mitophagy (MOFA3, MFA2, MONET3). These results are summarised in Supplementary Figures 26, 33, 43, respectively. Four-cluster output from LRAcluster resolves two ER-groups: an immune-dominant subset (LRAcluster1) and a proliferative subset (LRAcluster3), together with two ER+ groups: an angiogenic/hormone-responsive subset (LRAcluster2) and a mitochondrial subset (LRAcluster4); see Supplementary Figure 23.

Five-cluster solutions (MSNE, iClusterBayes) yield a similar 3 ER+/2 ER-split. The ER-subsets are either immune-enriched (MSNE4, iClusterBayes1) or cell-cycle–driven (MSNE1, iClusterBayes4). Among the ER+ subtypes, extracellular-matrix and angiogenesis pathways predominate in MSNE5 and iClusterBayes2, organelle-remodelling and proteostasis in MSNE2, and mitochondrial/mitophagy programmes in MSNE3 and iClusterBayes5 (see also Supplementary Figures 14, 38). The eight-cluster wMKL solution preserves this theme as well: the principal ER-subset (wMKL4) demostrates up-regulation of imune processes, whereas ER+ subsets partition into angiogenic (wMKL1), immune (wMKL2) and mitochondrial/mitophagy (wMKL3) profiles (Supplementary Figure 21).

#### Modality contribution & feature importance

This section discusses insights from built-in functionality for feature-level and/or omic-level interpretation, excluding inspection of dimensionality reduction plots. For Similarity network methods, the lists of highly ranked features consisted mainly of methylation, RNA-seq and miRNA features for ab-SNF, ANF, NEMO, RWR-NF and SNF (Supplementary Figure 9). Notably, RWR-F’s respective list was enriched in CNV features, accompanied by methylation, RNA-seq and SNP features (Supplementary Figure 15e). MDICC learns kernel weights at the omic-level, with RNA-seq heavily influencing the results (Kernel weight: 1.0 vs. 4.096 · 10^−9^ for all other modalities). The sum of each modality’s set of 55 kernel weights, indicates that the omic-level importance ranking for wMKL is: SNPs: 0.38 *>* RNA-seq: 0.25 *>* Methylation: 0.20 *>* miRNA: 0.17 *>* CNV: 2.4 · 10^−3^. Similarly, the corresponding order for CIMLR is: RNA-seq: 0.462 *>* miRNA: 0.400 *>* CNV: 0.117 *>* Methylation: 0.023 *>* SNPs: 4.096· 10^−9^. By taking the average weight per modality in KLIC, we get an order of importance: RNA-seq: 0.278 *>* miRNA: 0.214 *>* Methylation: 0.182 *>* CNV: 0.173 *>* SNPs: 0.153. The three modules (not clusters) produced by MONET demonstrate separation with respect to ER status and HER2 status (Supplementary Figure 44). Upon inspecting the excess within-module connectivity, none of the five data types provides above-background cohesion, indicating that these patient modules are not dominated by any single molecular layer.

Plots of explained variance for each modality in each latent dimension produced by MOFA reveal that the elected 7 latent space dimensions capture variability in the CNV data, RNA-seq, methylation and to a lesser extent miRNA (Supplementary Figure 31). In factor 1, CNV features dominate with loadings near 1, followed by the *TP53* SNP only with a weight around 0.3 (Supplementary Figure 34). From factor 2 and on, RNA-seq features have consistently high weights and they dominate factor 4. MOFA factor 2 captures the differences in ER status, whereas factors 6 and 7 capture stage IV variability (Supplementary Figures 35, 36). In MFA, feature-level contribution plots indicate that the top ranking features are miRNA features, followed by RNA-seq and methylation (Supplementary Figure 27). RNA-seq, miRNA and methylation contributions dominate the first two dimensions, whereas the former two dominate the third. SNPs and CNVs are the main contributors in dimensions 4 and 5 respectively (Supplementary Figure 28). The first dimension captures ER status and to a lower extent tumour stage variability, while the second dimension shows marginal HER2+/-separation (Supplementary Figures 29, 30). In iClusterBayes, the top 1000 contributing features consist of 613 RNA-seq, 259 SNP and 128 methylation features. Overall, the average contribution score for modalities ranks as follows: methylation (1.91) *>* RNA-seq (1.71) *>* SNPs (1.55) *>* CNVs (0.90) *>* miRNAs (0.49).

### Method comparisons

Table 2 summarises how methods that have been used in the individual method runs compare against each other with respect to silhouette metrics, aggregated runtimes, heterogenous data handling, documentation and explainability/interpretability. For more details on these criteria see Section Method comparisons. Three methods (iClusterBayes [16], LRAcluster and MOFA [30]) have pre-built binary data handling functionality, while the rest of the methods are either amenable to use binary data, e.g., by incorporating appropriate distance metrics, or can handle binary data upon small modifications in their source code. Seven methods (CIMLR [15], iClusterBayes [16], MOFA [30], MSNE [32], RWR-F [33], RWR-NF [33] and wMKL [35]) have implemented some form of task parallelism in each job to speed up the underlying process. In terms of handling missing values, MSNE [32], NEMO [22], MONET [31], COCA [18, 39] and MOFA [30] have functionality to handle missing values, although missing data scenarios were outside the scope of this work and were therefore, not examined. Most of the methods have adequate or extended documentation (installation instructions, tutorials/examples). A few methodologies offer interpretability options at the omic or feature level, which extend beyond the inspection of low-dimensional projection plots. MFA [36] and MOFA [30] stand out with extensive functionality for examining feature and modality contribution to the final results.

**Table 2.**
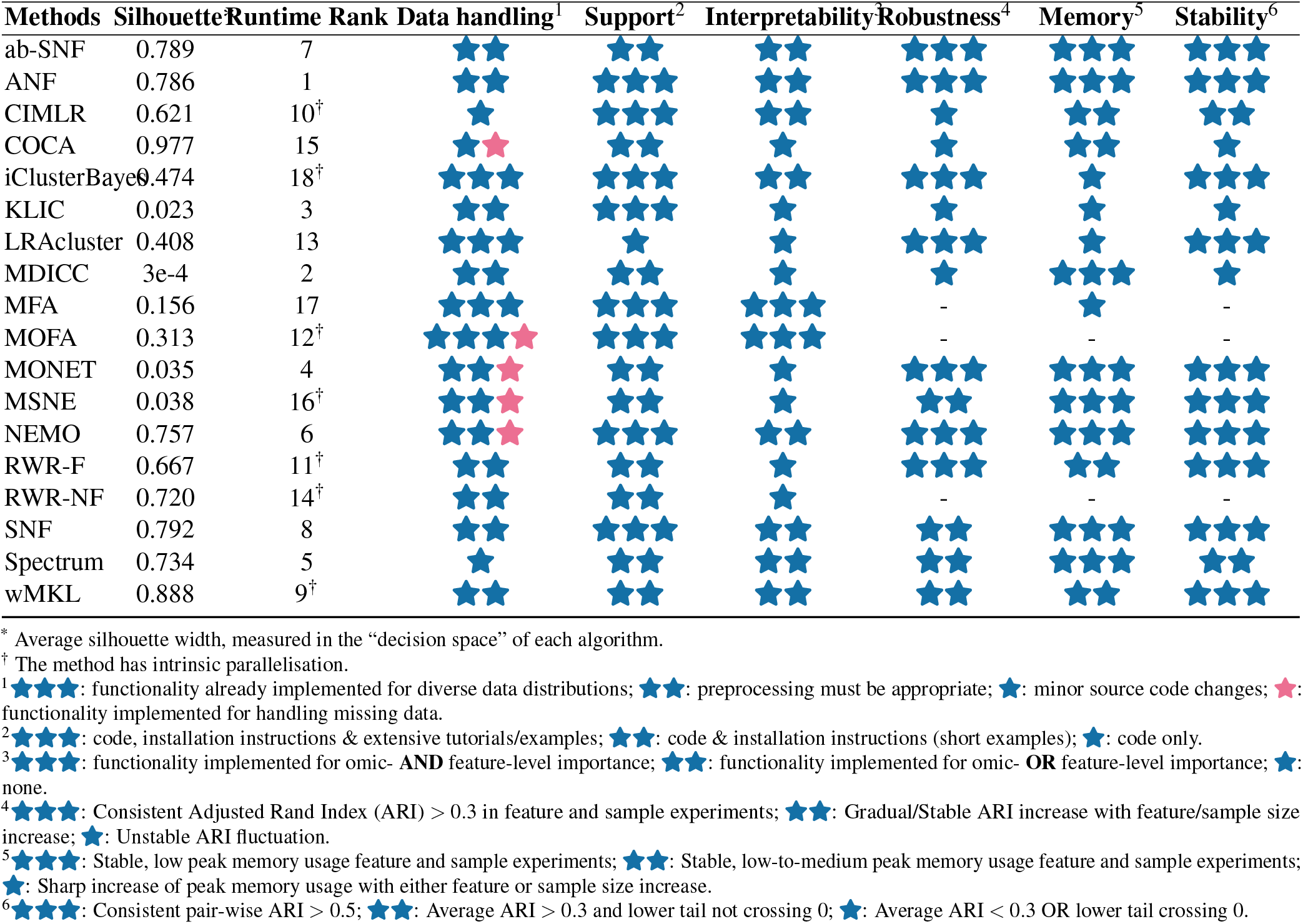
Comparison of multi-omics clustering methods.

**Table 3.**
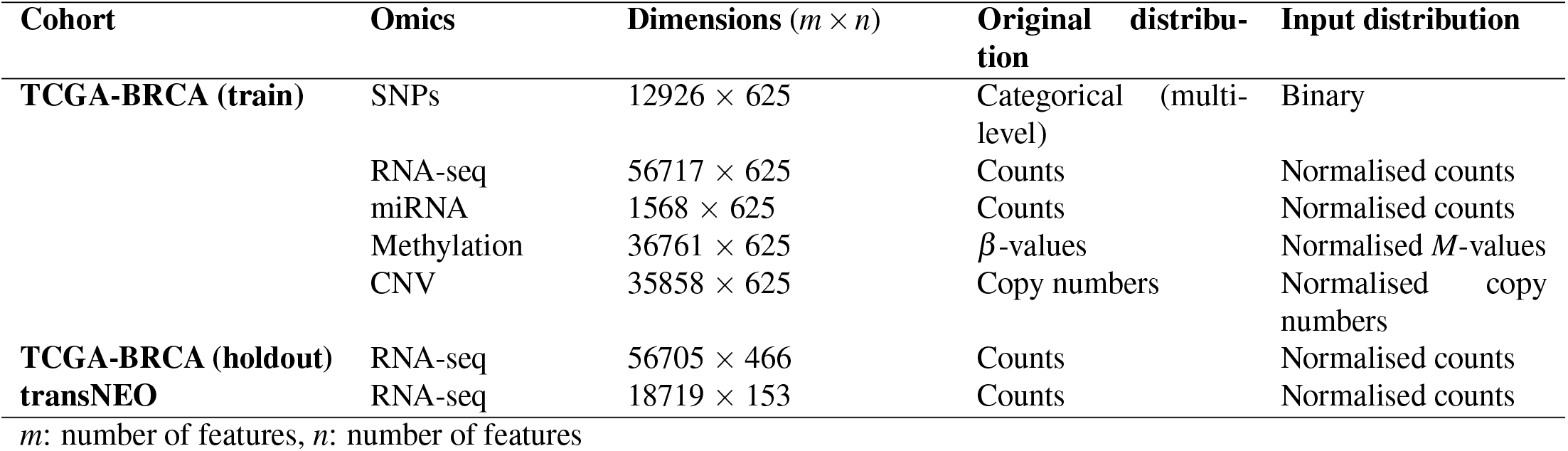
Overview of omic modalities.

**Table 4.**
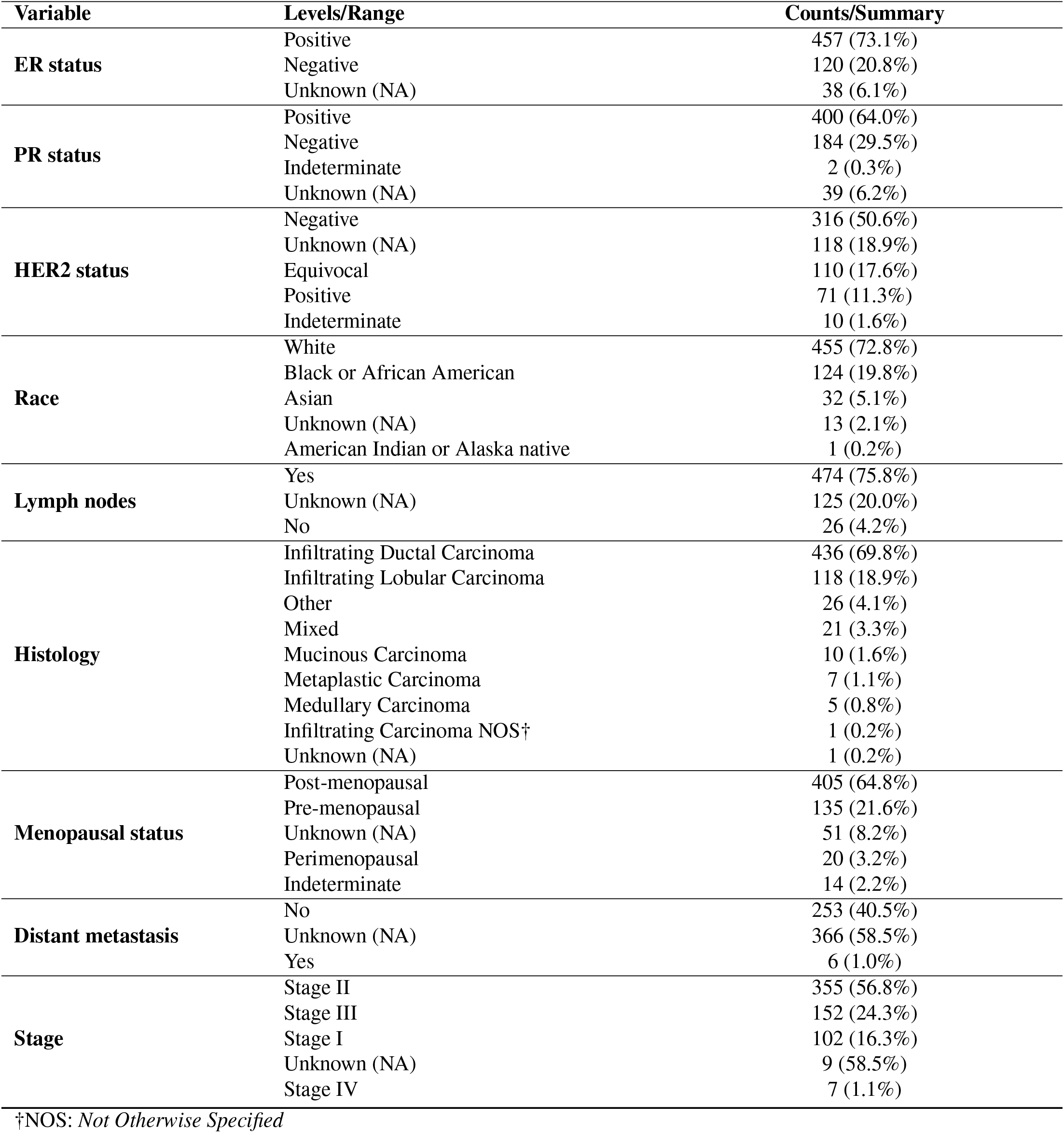
Sample characteristics in the training set (*N* = 625)

**Table 5.**
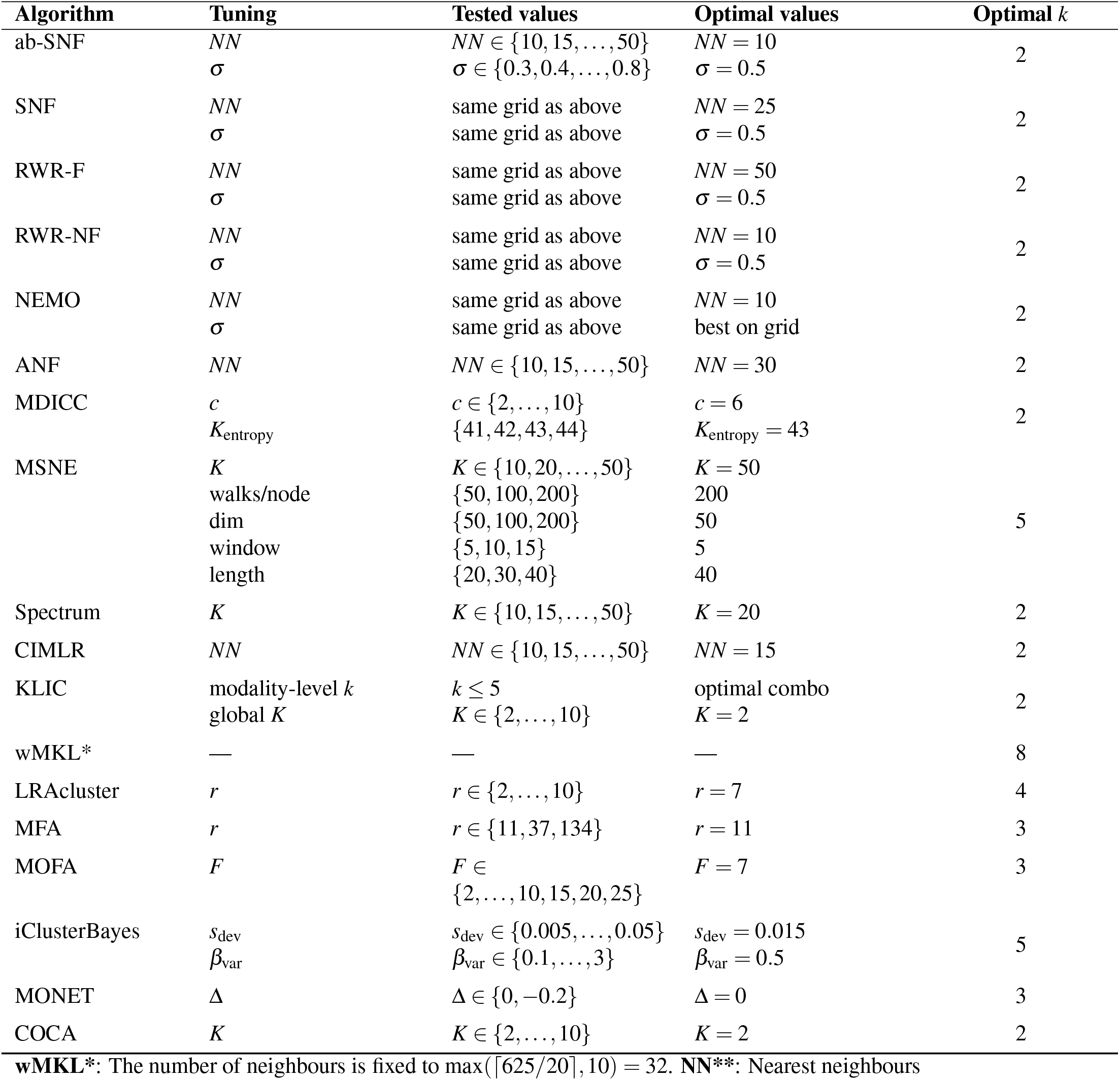
Method optimisation.

| Algorithm | Tuning | Tested values | Optimal values | Optimal $k$ |
| --- | --- | --- | --- | --- |
| ab-SNF | $NN$<br>$\sigma$ | $NN \in \{10, 15, \dots, 50\}$<br>$\sigma \in \{0.3, 0.4, \dots, 0.8\}$ | $NN = 10$<br>$\sigma = 0.5$ | 2 |
| SNF | $NN$<br>$\sigma$ | same grid as above<br>same grid as above | $NN = 25$<br>$\sigma = 0.5$ | 2 |
| RWR-F | $NN$<br>$\sigma$ | same grid as above<br>same grid as above | $NN = 50$<br>$\sigma = 0.5$ | 2 |
| RWR-NF | $NN$<br>$\sigma$ | same grid as above<br>same grid as above | $NN = 10$<br>$\sigma = 0.5$ | 2 |
| NEMO | $NN$<br>$\sigma$ | same grid as above<br>same grid as above | $NN = 10$<br>best on grid | 2 |
| ANF | $NN$ | $NN \in \{10, 15, \dots, 50\}$ | $NN = 30$ | 2 |
| MDICC | $c$<br>$K_{\text{entropy}}$ | $c \in \{2, \dots, 10\}$<br>$\{41, 42, 43, 44\}$ | $c = 6$<br>$K_{\text{entropy}} = 43$ | 2 |
| MSNE | $K$<br>walks/node<br>dim<br>window<br>length | $K \in \{10, 20, \dots, 50\}$<br>$\{50, 100, 200\}$<br>$\{50, 100, 200\}$<br>$\{5, 10, 15\}$<br>$\{20, 30, 40\}$ | $K = 50$<br>200<br>50<br>5<br>40 | 5 |
| Spectrum | $K$ | $K \in \{10, 15, \dots, 50\}$ | $K = 20$ | 2 |
| CIMLR | $NN$ | $NN \in \{10, 15, \dots, 50\}$ | $NN = 15$ | 2 |
| KLIC | modality-level $k$<br>global $K$ | $k \leq 5$<br>$K \in \{2, \dots, 10\}$ | optimal combo<br>$K = 2$ | 2 |
| wMKL* | — | — | — | 8 |
| LRacluster | $r$ | $r \in \{2, \dots, 10\}$ | $r = 7$ | 4 |
| MFA | $r$ | $r \in \{11, 37, 134\}$ | $r = 11$ | 3 |
| MOFA | $F$ | $F \in \{2, \dots, 10, 15, 20, 25\}$ | $F = 7$ | 3 |
| iClusterBayes | $s_{\text{dev}}$<br>$\beta_{\text{var}}$ | $s_{\text{dev}} \in \{0.005, \dots, 0.05\}$<br>$\beta_{\text{var}} \in \{0.1, \dots, 3\}$ | $s_{\text{dev}} = 0.015$<br>$\beta_{\text{var}} = 0.5$ | 5 |
| MONET | $\Delta$ | $\Delta \in \{0, -0.2\}$ | $\Delta = 0$ | 3 |
| COCA | $K$ | $K \in \{2, \dots, 10\}$ | $K = 2$ | 2 |

**wMKL\***: The number of neighbours is fixed to $\max(\lceil 625/20 \rceil, 10) = 32$ . **NN\*\***: Nearest neighbours

#### Method pair-wise agreement

Figure 3a illustrates the clustering agreement between different algorithms, as quantified by ARI. ARI is preferrable to Normalised Mutual Information (NMI) for measuring clustering agreement, due to NMI’s biased behavior, particularly when the number of clusters is large [40]. Among methods from the same broad category, similarity network methods exhibit the highest levels of agreement. Other notably concordant pairs include MOFA-LRAcluster, iClusterBayes-RWR-F. Overall, MFA and iClusterBayes exhibit moderate levels of agreement with each other, a set of highly concordant similarity network methods (ab-SNF, ANF, NEMO, RWR-NF, SNF and Spectrum) and wMKL. The majority of clustering ARI agreements are reflected at the pathway level as well, apart from a few notable cases: MFA-MONET, MFA-MSNE, CIMLR-MDICC and MOFA-similarity network methods.

**Figure 3.**
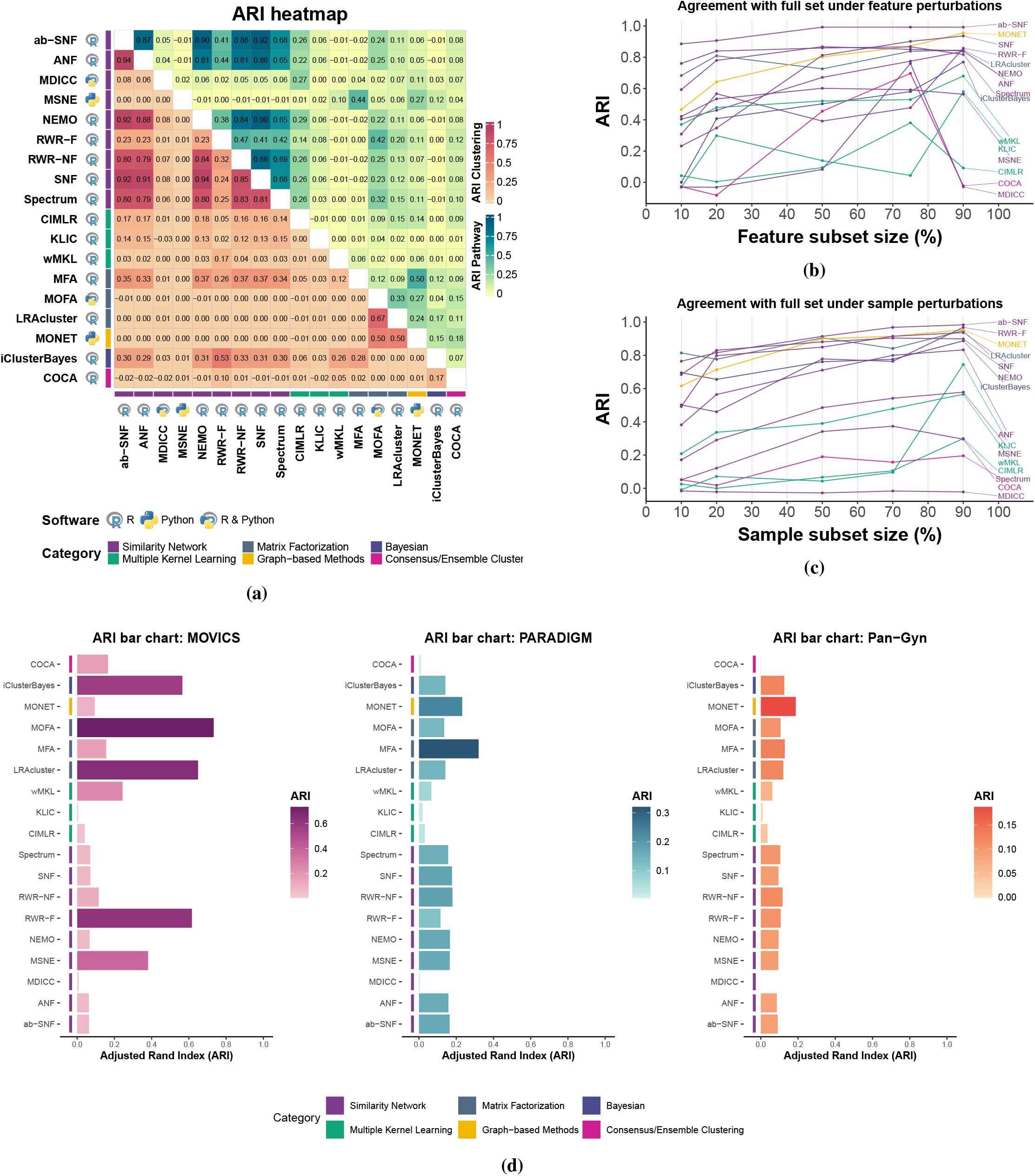
Comparison of clustering agreement and robustness across methods. **(a)** Adjusted Rand Index (ARI) heatmap. Pair-wise ARI values between method clusterings (lower triangle) and discovered enriched pathways (upper triangle). **(b)** ARI agreement under feature perturbations. Comparisons between feature subsets and full feature set for each method (MFA, RWR-NF and MOFA excluded). **(c)** ARI agreement under sample perturbations. Comparisons between sample subsets and full sample set for each method (MFA, RWR-NF and MOFA excluded). **(d)** Method ARI agreement with the baseline MOVICS analysis, PARADIGM and Pan-Gyn clusters on TCGA.

#### Robustness, scalability and stability

We defined robustness as the ARI-measured agreement between a method’s clustering under feature/sample perturbations (i.e., feature/sample subsets of the full dataset) and the clustering obtained from the full dataset under the same method parametrisation (Figures 3b,3c). Peak memory and core runtime (measuring the main integration step of each method) were recorded for both feature and sample perturbation scenarios to characterise scaling with increasing numbers of features and/or samples (Figures 4a,4b,4c,4d). We assessed method stability using the 10 replicates of the 90% sample subset (independent subsamplings of the full set without replacement, stratified for ER status) by computing pairwise ARIs on the overlap of samples and summarising them by the median, comparing methods (Figure 4f). MFA, MOFA and RWR-NF were excluded from these analyses (see Section Robustness, scalability and stability benchmarks).

**Figure 4.**
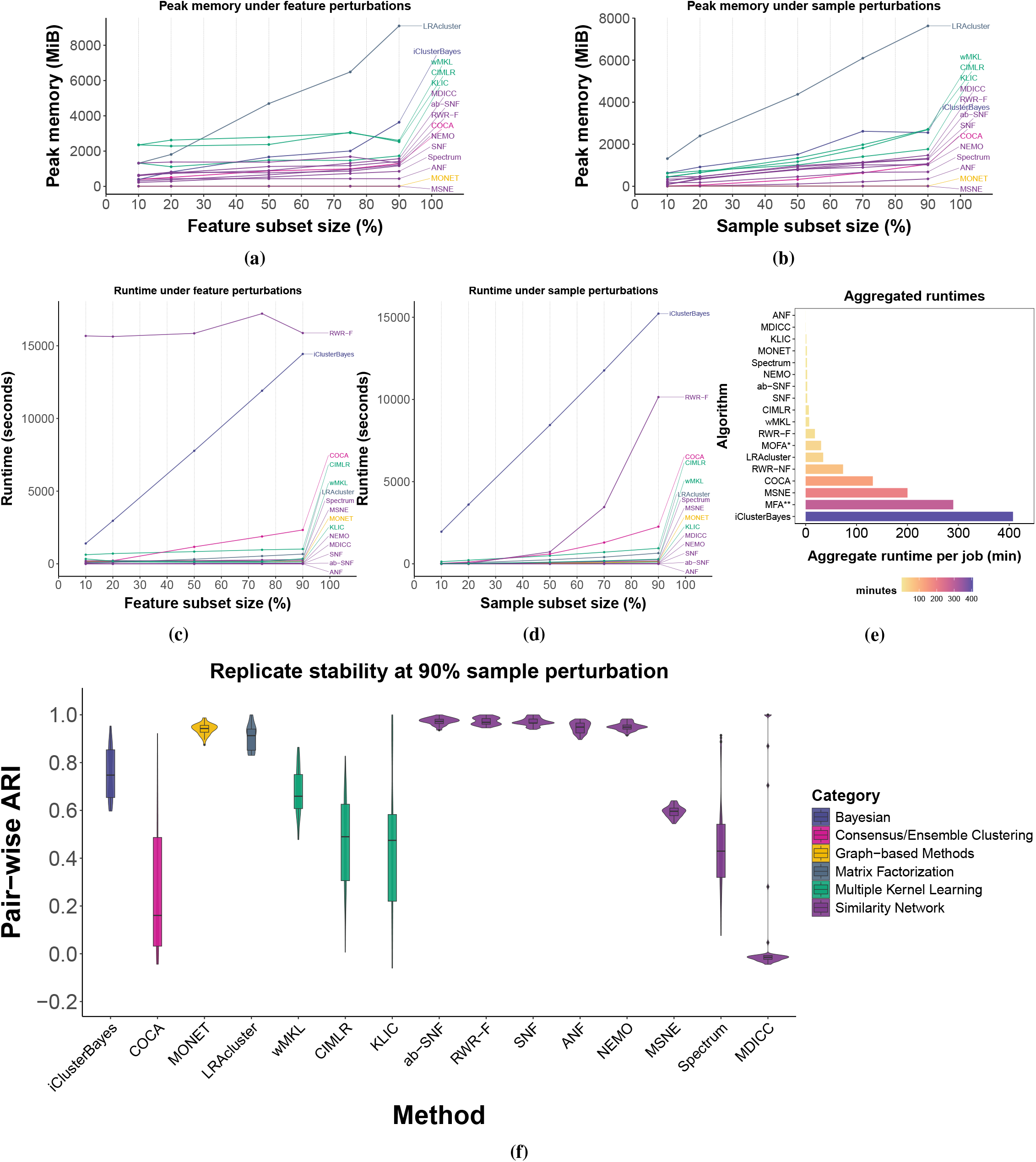
Peak memory, runtime and resampling stability comparisons. **(a)** Peak memory under feature perturbations. **(b)** Peak memory under sample perturbations. **(c)** Runtime under feature perturbations. **(d)** Median runtime under sample perturbations. **(e)** Aggregated runtime per job in non-benchmark runs. **(f)** Replicate stability at 90% sample subset. Violin plots of ARI values between different resamplings (90% of the full set) for each method. MFA, RWR-NF and MOFA excluded from subfigures (a), (b), (c), (d), (f).

**Figure 5.**
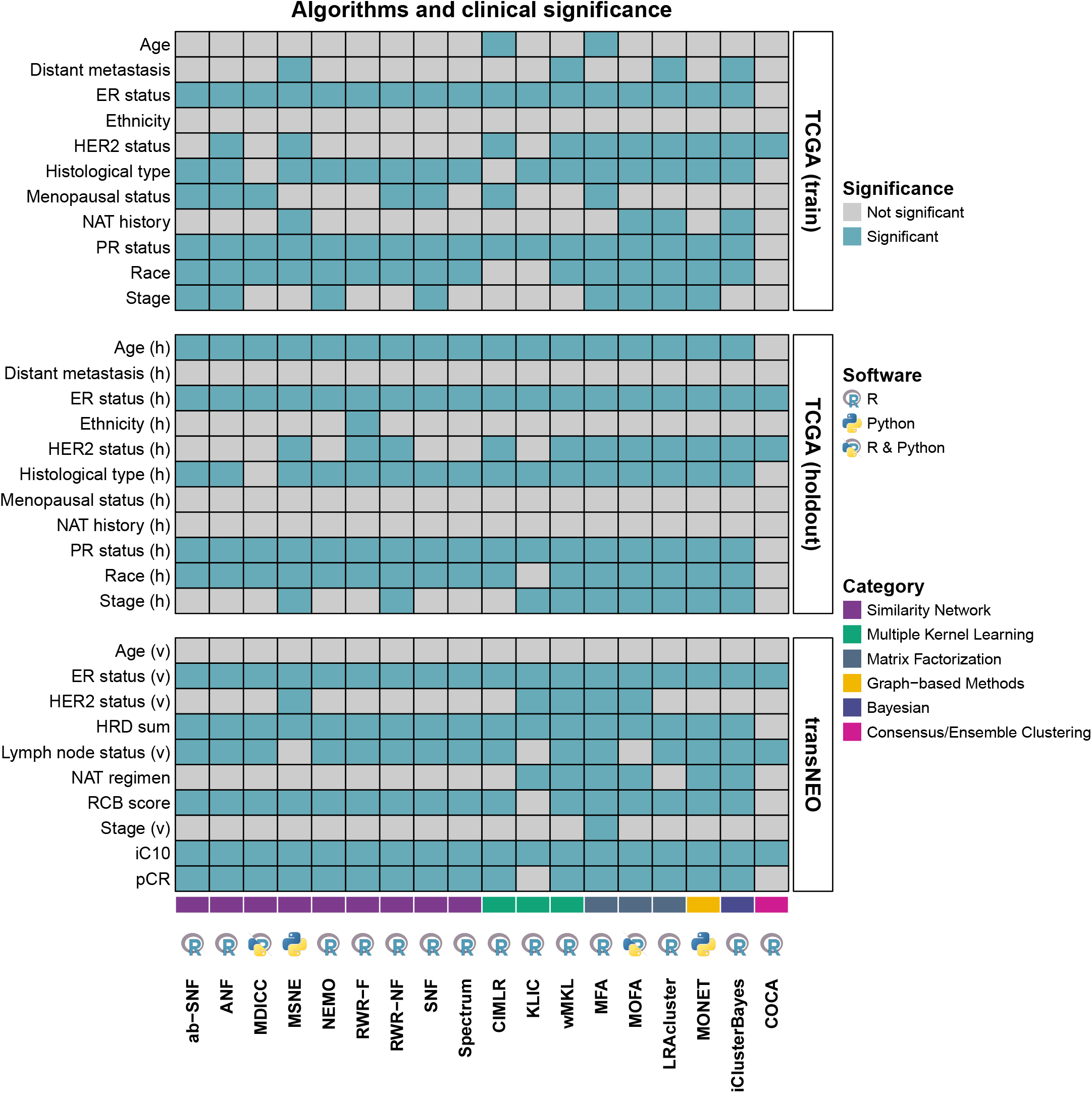
Heatmap of clinical associations. Blue tiles indicate statistically significant relationships and grey tiles indicate non-significant relationships. Statistical significance was determined through non-parametric tests (𝒳^2^-test and Kruskal-Wallis test for categorical and continuous variables, respectively).

Most similarity network-based methods (ab-SNF, ANF, SNF, RWR-F, NEMO), together with MONET and LRAcluster, were robust to both feature and sample perturbations, showing good agreement with the full-data clustering (ARI *>* 0.5). Spectrum was robust under feature perturbations but performed consistently worse under sample perturbations (0.1 *<* ARI *<* 0.4). MSNE showed stably moderate robustness in the feature experiments (0.4 *<* ARI *<* 0.5) and improved as the sample fraction increased. Multiple kernel-learning methods were generally poorer (KLIC, CIMLR) or moderate (wMKL) relative to similarity network methods, only occasionally exceeding ARI = 0.5, most notably at the 90% feature and sample subsets; KLIC, in particular, increased sharply from approximately 0.1 to *>* 0.5 at 90% in both perturbation settings. iClusterBayes maintained consistently high robustness across sample subsets (ARI *>* 0.6) and increased gradually under feature subsampling, whereas COCA and MDICC showed poor robustness in both scenarios.

Regarding stability analyses, ab-SNF, ANF, NEMO, RWR-F, SNF, LRAcluster, and MONET were the most stable (median ARI *>* 0.9). iClusterBayes followed (median ARI = 0.75), then wMKL (median ARI = 0.66) and MSNE (median ARI = 0.60). CIMLR, KLIC, and Spectrum showed lower median stability (0.49, 0.47, and 0.43, respectively) and substantially greater variability across replicate pairs. MDICC and COCA exhibited the lowest stability overall.

LRAcluster was the most memory-demanding method as feature and sample sizes increased, followed by iClusterBayes and the multiple kernel-learning approaches; all other methods required consistently less memory. For runtime under feature perturbations, RWR-F was the slowest, remaining constant at *>* 4 h across feature subsets, while iClusterBayes increased approximately linearly with the number of features (𝒪 (*p*)). COCA showed only a modest runtime increase with larger feature proportions and remained substantially faster than RWR-F and iClusterBayes; all other methods exhibited largely stable and low runtimes across feature subsets. Under sample perturbations, iClusterBayes again took the longest, scaling linearly with sample size and increasing from ∼0.5 h at the smallest subsets to *>* 4 h at 90%. RWR-F followed, with a steep, cubic-like dependence on sample size (estimated 𝒪 (*n*^3.69^)), yet still completing within *<* 3 h at 90%, consistent with a predominant dependence on *n* rather than *p*. COCA showed a mild quadratic increase (𝒪 (*n*^2^)) but remained faster than both. The remaining methods exhibited linear-to-quadratic scaling with significantly lower absolute runtimes, and were therefore comparatively quick in both perturbation settings.

#### Aggregated runtimes

Aggregated runtimes provide a more practical estimate of end-to-end computational cost than the single-core, no-multithreading benchmarks reported in Section Robustness, scalability and stability. Specifically, these aggregated estimates are derived from our non-benchmark runs and average runtime across tuning and fitting while accounting for (i) the number of cores allocated, (ii) the number of jobs executed during hyperparameter tuning, and (iii) whether parallelisation was enabled where supported. Similarity network methods rank higher on the list, while Matrix Factorisation methods and the Bayesian iClusterBayes tend to be the slowest (Figure 4e; Table 2: Runtime Rank). MFA jobs with subsets (or the full set) of features in proportions higher than 33% (50%, 75%, 100%) failed due to exceeding the 12-hour time limit, which essentially ranks MFA as the slowest method overall.

#### TCGA Ground Truth

Figure 3d summarises the concordance between the partitions generated by each individual algorithm and three references: the baseline *MOVICS* analysis and the previously published PARADIGM and PanGyn multi-omic clusters for the The Cancer Genome Atlas (TCGA) training cohort, as these are stored in the *TCGAbiolinks* package in R [41–43]. The highest ARI scores are obtained against the *MOVICS* baseline (highest, MOFA: *ARI* = 0.73), reflecting the fact that both analyses were performed on identical input. In contrast, PARADIGM and PanGyn were derived from distinct sample subsets and modality combinations in their original TCGA studies, leading to systematically lower concordance. Among the methods evaluated, MFA shows the strongest agreement with PARADIGM (*ARI* = 0.32), whereas agreement with PanGyn is uniformly weaker; here, MONET attains the top score (*ARI* = 0.19).

#### Clinical variables

The identified clusters from each algorithm are associated with clinical variables of interest in the training and evaluation cohorts (Figure 5). All algorithms apart from COCA identify subtypes significantly associated with ER status. In most algorithms which identify an optimal number of *k* = 2 clusters (ab-SNF, ANF, NEMO, RWR-NF, SNF, Spectrum), the produced partitioning separates the ER+ and ER-samples. Other variables which we found regularly to be associated with clusterings are histological type, HER2 status, age (in the TCGA holdout set only) and stage. In the transNEO cohort, significant associations were found between clusters and Homologous Recombination Deficiency (HRD) score, lymph node presence, pathological Complete Response (pCR) and integrative clusters (IC10) subtypes.

### Consensus analysis

Consensus analysis on the full list of results (all 18 algorithms), revealed two clusters (*k* = 2; silhouette = 0.613, Figure 6a,c,e). The highest agreement is observed between the consensus clustering results and results from similarity network methods which exhibited high ARIs with each other (Figure 6e). The two Consensus Clusters (CCs), CC1 and CC2, reflect results from the individual clusterings produced earlier. CC1 is enriched in ER+, *PIK3CA*-mutated samples and shows up-regulation of ciliary, vesicle transport and extra-cellular transport pathways, alongside cell–matrix adhesion and angiogenesis, while CC2 is rich in ER-, *TP53*-mutated samples characterised by deregulated cell cycle pathways (Supplementary Figures 46, 47).

**Figure 6.**
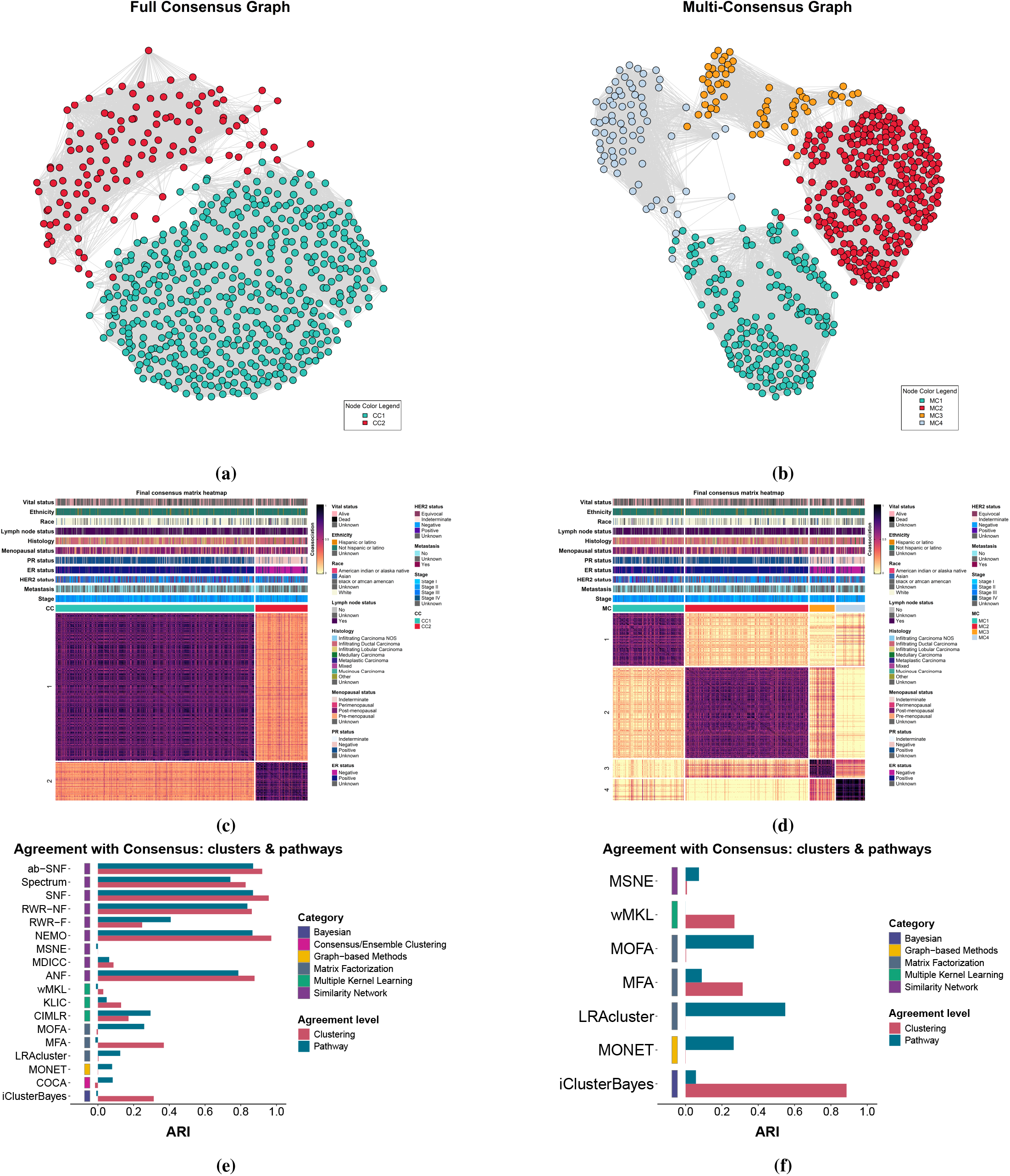
Consensus analysis. **(a)** Consensus graph using results from all methods (full consensus). **(b)** Consensus graph using results with more than 2 clusters (multi-consensus). **(c)** Full consensus co-association heatmap. **(d)** Multi-consensus co-association heatmap. **(e)** ARI bar chart: method agreement with full consensus. **(f)** ARI bar chart: method agreement with multi-consensus.

Repeating the procedure with only those algorithms that originally produced more than two clusters yielded a solution with four Multi-Consensus (MC) clusters (*k* = 4; silhouette = 0.610, Figure 6b,d,f), effectively splitting both the ER+ and ER-groups into two subsets: MC1 and MC2 are ER+, while MC3 and MC4 are ER-. The two ER+ subsets have high percentages of *PIK3CA* mutations, while the two ER-subsets have high *TP53* percentages (Supplementary Figures 48a,d). In terms of FGA, MC1 has significantly high FGL compared to the rest of the clusters (Supplementary Figure 48c). MC1 also has a higher proportion of HER2+ samples (21.8%, Supplementary Figure 48b) and is further characterised by up-regulated intracellular transport and mitochondrial pathways (Supplementary Figure 49). The other ER+ cluster, MC2 demonstrates up-regulation in extracellular pathways, angiogenesis and hormonal signalling processes. With respect to the ER-clusters, MC3 has higher immune activity than MC4, while MC4 exhibits higher cell cycle deregulation (Supplementary Figure 48). Overall, ER+/ER-separation is preserved in low-dimensional RNA-seq, CNV and methylation spaces (Supplementary Figure 50). MC1/2 replicate the functional split seen in MOFA/MFA/MONET, whereas MC3/4 recapitulate the immune-versus-proliferative dichotomy observed in MSNE and iClusterBayes.

### Selecting a strategy

The selection of a method (or a series of methods which will may be combined in a consensus approach) can be based on one or more of the following criteria: resource availability, scalability, robustness, stability, interpretability, data heterogeneity and expected/desirable granularity.

For researchers who do not have access to/do not want to rely on a High-Performance Computing (HPC), focus can be set on lighter methods which do not require extensive memory and/or running time, such as similarity network methods (SNF, ab-SNF, ANF, NEMO) or the graph-based MONET. These algorithms required an aggregated *<* 15 min per run and converged on highly concordant partitions (both between them and in stability experiments). These methods are quick, stable, robust and produce biologically plausible partitions, albeit at a high level and with limited post-hoc importance quantification. SNF, ab-SNF, ANF and NEMO offer post-hoc feature-level contribution analysis options, whereas MONET offers omic-level contribution quantification with respect to the discovered modules (not clusters) which is more biologically interpretable and is useful for clarifying the effects of single omic layers and combinations thereof. The properties of these methods make them a reasonable choice when the sample size is small, resources are limited and deep contextual feature- or omic-level importance is not required.

Multiple kernel learners (CIMLR, KLIC, wMKL) can also be run locally with the added benefit of quantifying the contribution of each omic to the final output, although they scored lower with respect to robustness and stability, and should therefore be run with higher sample sizes for more reliable results.

The similarity network methods Spectrum and MSNE, matrix factorisation methods and Bayesian methods generally have extensive parameter tuning, and are better run in a HPC. MOFA runs can be significantly enhanced by using Graphics Processing Units (GPUs). MOFA and MFA result in rich latent representations for which omic-level and feature-level contribution can be quantified in a method-contextual way (as opposed to post-hoc distinct methodologies) and add interpretative power. Finally, if a researcher prefers to retain original data distributions (e.g. count data for RNA-seq) without coercing them into normal distributions (potentially with loss of information), this can be accommodated with methods such as MOFA, iClusterBayes and LRAcluster.

If 3 or more models are used, consensus approaches can also be explored. If the methods produce relatively concordant solutions with a similar number of clusters, the consensus will refine the partitions, whereas if the number of clusters varies significantly across methods, the consensus is not guaranteed to refine the partitions and may deviate from the results of every individual method. In these cases, we suggest considering the expected number of clusters and potential number of nested partitions of these, if applicable, and limit the consensus to the methods which produce a number of clusters *k*_*exp*_ ± 1, where *k*_*exp*_ is the expected number of clusters.

## Discussion

The rapid expansion of multi-omic technologies presents both an opportunity and a challenge for biomedical research: while diverse molecular assays offer complementary insights, the accompanying proliferation of computational tools makes method selection increasingly difficult. Here, we deliver a comprehensive survey of unsupervised machine learning approaches for multi-omic subtyping. By benchmarking 22 algorithms across five molecular layers and two external validation cohorts, and by comparing them with a consensus pipeline, we show how method choice, data transformations and parametrisation influence subgroup discovery and agreement with established clinical knowledge.

Our baseline experiment consisted of a fully automated run of the ten algorithms included in *MOVICS* and deliberately treated heterogeneous omics as if they were continuous and identically distributed. The resulting two baseline consensus subtypes (CS1 and CS2) were clinically informative, yet closer inspection revealed that copy-number variation dominated the cluster separation over the remaining modalities. Baseline subtypes were significantly associated with ER status, however, the separation of ER+ and ER-samples between the two subtypes was not pure. These observations suggest that default settings which overlook data-type idiosyncrasies may inflate performance estimates and obscure biologically relevant structure. A more carefully designed, distributionally appropriate tuning strategy is, therefore, required. In parallel, our baseline ER+ versus ER-analysis showed that a large fraction of the observed molecular and pathway differences can already be anticipated from a simple ER-based partition of the data. This underlines the importance of distinguishing structure that genuinely arises from multi-omic integration from patterns that algorithms capture because they have effectively rediscovered ER status.

Despite methodological differences, most tools recover a remarkably coherent biological axis in breast cancer. Across 17 of 18 methods, we observed consistent enrichment of DNA double-strand-break repair, nucleotide metabolism, and core cell-cycle programmes, along with modulation of leukocyte activity and apoptosis. The axis manifests as two broad subtypes: (1) an ER-positive, hormone-responsive phenotype marked by angiogenesis and vesicle-mediated export, and (2) an ER-negative, highly proliferative phenotype dominated by mitotic checkpoint and homologous-recombination pathways. Our ER-stratified baseline analysis indicates that many of these differences are intrinsic to the ER+/ER-split itself, so we interpret methods that primarily recover this axis as providing a robust, multi-omic realisation of established ER-driven biology rather than as discovering an entirely new taxonomy. Methods resulting in more than two subtypes further refine this backbone by adding more nested layers. Three-cluster solutions generated by factorisation models (MOFA, MFA) and graph methods (MONET) consistently split the *ER*+ arm into an angiogenic–extracellular-matrix subset and a mitochondrial–mitophagy subset, while retaining a single proliferative ER– subset. Four- and five-cluster outputs from LRAcluster, MSNE, and iClusterBayes refine the ER– branch into immune-enriched versus cell-cycle–enriched groups and resolve the ER+ branch into discrete angiogenic, organelle-remodelling, and bioenergetic subclasses.

Concordance between methods was strongest among similarity network methods (SNF, ab-SNF, ANF, NEMO, RWR-NF, Spectrum), which converged on near-identical bipartitions (*ARI >* 0.85). Across robustness, stability, and computational scaling analyses, similarity network-based methods (ab-SNF, ANF, SNF, RWR-F, NEMO) along with MONET and LRAcluster emerged as the most reliable overall, combining strong agreement with full-data clusterings under both feature and sample perturbations (ARI *>* 0.5), with high resampling stability (median ARI *>* 0.9). In contrast, Spectrum and the multiple kernel-learning methods were less consistent, with reduced robustness under sample subsampling and greater variability across resamples, while COCA and MDICC performed poorly across settings. These performance differences were accompanied by clear computational trade-offs: LRAcluster incurred the largest memory growth, whereas iClusterBayes and RWR-F dominated runtimes, with iClusterBayes scaling roughly linearly with respect to both number of features and samples, and RWR-F showing steep sample-size dependence.

Finally, aggregated runtimes from end-to-end runs (including tuning and parallel execution) reinforced these trends in practical usage, with similarity network methods remaining comparatively efficient, and Bayesian and matrix factorisation approaches among the slowest, with MFA frequently exceeding the 12-hour limit at higher feature proportions.

Building a consensus from all 18 tuned algorithms distilled the data into two robust clusters (silhouette = 0.617). This bipartition aligned almost perfectly with the similarity-network outputs and recapitulated a familiar axis between an ER+, *PIK3CA*-mutant lineage and an ER-, *TP53*-mutant lineage, as described in large sequencing cohorts where *PIK3CA* mutations are concentrated in ER+/luminal disease whereas *TP53* mutations are enriched in basal-like and HER2-enriched tumours [18]. The ER+ cluster (CC1) combined prominent ciliary and vesicle-/secretory-transport programmes with extracellular transport and cell matrix adhesion and angiogenesis signals, whereas the ER– cluster (CC2) was strongly enriched for mitotic cell-cycle, DNA-replication and chromosome-segregation pathways. This contrast aligns with the canonical luminal-basal divide seen in intrinsic subtype and integrative classifications: luminal/ER+ tumours with frequent *PIK3CA* mutation versus highly proliferative, *TP53*-mutant basal-like groups [44–46].

Consensus of the higher resolution methods yielded a four-cluster consensus solution (silhouette = 0.617). Each lineage of the previous consensus bipartition is further partitioned here. Within ER+ tumours, one subset (MC1) combined moderate HER2 enrichment, a higher burden of large-scale copy-number loss and up-regulation of mitochondrial respiratory-chain, Adenosine Tri-Phosphate (ATP)-synthesis and protein-import/folding pathways, consistent with a metabolically aberrated luminal state with heightened mitochondrial activity. The sister ER+ cluster (MC2) was instead characterised by extracellular-transport and secretory programmes, including angiogenesis, extracellular matrix assembly and multiple hormone-transport and hormone-response pathways, together with prominent cilium/axoneme-related signatures, suggestive of a more stromal–secretory, hormone-responsive phenotype. Within the ER-subgroups, one cluster (MC3) was clearly immune-enriched, with strong presence of antigen-presentation, lymphocyte-mediated immunity and interferon-response pathways, whereas the companion cluster (MC4) had low immune enrichment, but was dominated by chromosome condensation, mitotic cell cycle and DNA-replication/repair programmes. This immune-versus-proliferative contrast mirrors patterns previously described in triple-negative breast cancer [47, 48], while the metabolic–stromal split within the ER+ partitions represents an additional, previously described layer of ER+ tumour heterogeneity [45, 49, 50]. These within-lineage refinements are not captured by a simple ER+/ER-baseline comparison, indicating that the higher-resolution multi-omic methods are capturing structure beyond ER status alone.

Taken together, the two-cluster consensus recapitulates the primary dimension of breast tumour variation, while the four-cluster refinement reproduces the second-order structure reported for mitochondrial activity-high/HER2-low luminal, angiogenic lobular-rich luminal, immune-rich basal-like and hyper-proliferative basal-like tumours [45, 47, 48, 51]. The fact that a single, fully data-driven, label-agnostic ensemble can recover all four archetypes in one framework argues that the approach provides a principled bridge from broad classification to fine-grained, treatment-relevant stratification.

To translate these insights into practice, we propose a decision framework (Figure 7) that guides researchers from data characteristics and research goals to an actionable subset of algorithms. Researchers without access to (or preferring not to rely on) a HPC can focus on faster, lighter methods such as similarity network approaches or the graph-based MONET, all of which complete in under 15 minutes per run and yield highly concordant partitions. Multiple kernel learners can also be run locally, preferrably with larger sample sizes for more robust results, while quantifying each omic’s contribution. Higher-memory or heavily parameterised methods (MSNE, matrix-factorisation tools, Bayesian models) are best run on a HPC, may benefit from GPU acceleration (MOFA), and produce interpretable latent representations with richer context. Finally, when using three or more methods, consensus clustering can enhance robustness: if base methods produce similar *k*, consensus refines partitions, however, if *k* varies widely, it may drift away from each solution. We recommend limiting the ensemble to partitions whose *k* lies within *k*_*exp*_ ± 1.

**Figure 7.**
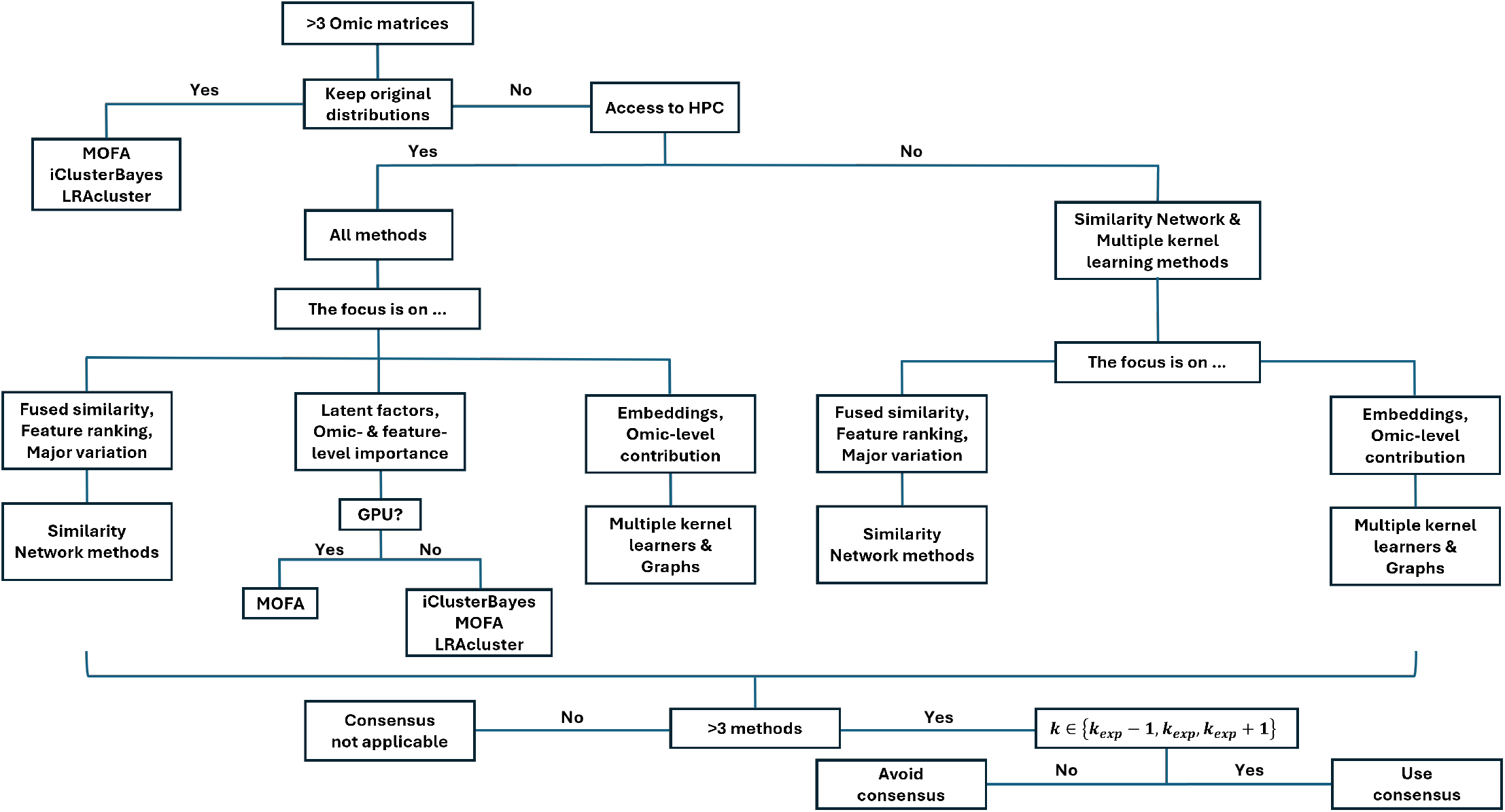
Selecting a workflow. Determine a sensible strategy based on available data, resources and desired research outcomes.

There are a few limitations to our stduy worth noting. First, our survey focused on algorithms with publicly available implementations in R or Python. We therefore excluded approaches released only as stand-alone executables or as proprietary software. Our external validation label-mapping approach used only transcriptomic data, precluding a full assessment of cross-modal generalizability, which is a task beyond the scope of this work. Although all methods were run on identically pre-processed data, we did not exhaustively explore every possible hyperparameter choice; performance may vary with alternative settings or additional omic layers. No feature pre-filtering (based on quality or other data-driven criteria) is carried out in any of the individual runs, because our goal was to also examine the algorithms’ ability to handle large matrices with noisy features. Feature filtering is a step described as a reasonable first step in the integration process usually, and it would be expected to affect results and downstream analyses for a handful of methods, as we showed in our feature perturbation experiments. Furthermore, the algorithms examined in this work can handle binary data either by default or after appropriate input preprocessing and/or minor source code modifications. Several methods which are built for continuous data only are not considered in this work. Runtime and memory benchmarks should be interpreted cautiously, as all methods were evaluated under a deliberately constrained single-core, no-multithreading setup that does not reflect the parallel execution available in typical HPC use. Finally, deep learning approaches were excluded, and therefore, the conclusions of this work are not applicable to this category of methods.

In summary, our results demonstrate that modern multiomic clustering algorithms, when appropriately tuned, converge on a shared biological landscape, yet offer complementary strengths in resolution, interpretability and computational efficiency. We provide a decision flowchart to help investigators match study constraints to optimal tool choice, and we encourage the usage of an ensemble consensus as a robust default for large-scale integrative studies.

## Conclusions

Modern unsupervised multi-omic algorithms, when tuned on harmonised data, converge on a shared multi-layered luminal–basal axis in breast cancer and can be ensembled to expose clinically relevant sub-strata. Method selection heavily depends on the distributional properties of data, the available resources and the end goals of each research endeavour, with respect to resolution, robustness and interpretability. Similarity network-based methods offer a fast, reliable baseline. Matrix factorisation and Bayesian factor models add resolution and mechanistic interpretability at higher computational cost. Our simple decision workflow provides a practical route from omics to actionable subtypes.

## Methods

A more detailed description of the methodology along with extended details on the different methods that were examined in this work can be found in the Supplementary Material (Supplementary File 1; available in Zenodo.

### Data characteristics and preprocessing

Throughout this work, we used the same training and evaluation cohorts for each method. The clinical annotation of these cohorts can be found in Supplementary File 3. The processed omic data for all cohorts can be found in Supplementary File 4 (available in Zenodo).

#### Training cohort

The training cohort is a subset of the TCGA-BRCA dataset. More specifically, primary tumour open-access data were downloaded from TCGA and preprocessed using the *TCGAbiolinks* package [41–43] in R [52]. Five different types of omics datasets were downloaded from TCGA, plus clinical data: RNA-seq (STAR counts), miRNA (miRNA-Seq), CNVs (gene level copy numbers; ABSOLUTE LiftOver pipeline [53]), methylation (Illumina Human Methylation 450; *beta*-values were converted to *M*-values) and SNPs (masked somatic mutations; aliquot ensemble somatic variant merging and masking).

Regardless of omics dataset, the first step of the preprocessing was to identify the male samples that were downloaded and remove them from the analysis. For RNA-seq and miRNA data, unstranded counts and read counts, respectively, were filtered to retain only rows with non-zero values. The median of ratios normalisation method was applied using the *DESeq2* package [54], followed by log 2-transformation with a data-driven selection of pseudocount of 1 [55], to mitigate the effect of outliers. Rows annotated with the same Human Genome Organization (HUGO) symbol were reduced to the one with the highest variance, and any remaining rows with zero values were removed. The resulting matrices were then standardised.

For CNV data, raw copy number values were filtered similarly, and rows with identical HUGO annotations were combined by retaining the one with the highest variance. Methylation data underwent filtering to remove rows with more than 25% missing values, followed by *k*-NN imputation [56]. Methylation *β* -values were converted to *M*-values for statistical suitability [57], with rows standardised post-conversion. SNP data were transformed into a binary matrix indicating the presence or absence of SNPs (of any category) per gene, and rows with all-zero values were discarded.

For extensive details on data preprocessing, see Section “Preprocessing” in Supplementary File 1.

#### Evaluation cohorts

The evaluation cohorts used throughout this work are the transNEO cohort [58] and the remaining samples from TCGA-BRCA with RNA-seq data available. The TCGA evaluation cohort’s expression data (only modality used for label propagation) have been preprocessed identically to the expression data from the training cohort, but separately. Regarding transNEO, the preprocessing scripts follow the original transNEO preprocessing steps with minor or no modifications and are designed for using within the official transNEO repository (https://github.com/cclab-brca/neoadjuvant-therapy-response-predictor). After obtaining the log 2-normalised versions in both cases, the matrices were then standardised.

### Compiling a list of methods

We compiled a list of methods described and/or mentioned by name in a series of reviews on unsupervised multi-omic analysis [1–8, 10, 59–61]. Articles which cited the reviews or were deemed similar by the PubMed database (https://pubmed.ncbi.nlm.nih.gov/) and ConnectedPapers (https://www.connectedpapers.com/) were also considered. After compiling an initial list of methods, we then filtered this list, for methods which:

1. are publicly available in either **Python or R**
2. are unsupervised and can be used for subtyping samples/patients
3. are flexible/reasonably amenable to work with both numerical and categorical multi-omic data

The full list, along with justifications for excluded methods, can be found in Section “Compiling a list of methods” in Supplementary File 1 (Supplementary Table 1).

### General pipeline

We implemented a general pipeline *after* each clustering output was produced, which borrows and adapts functionalities predominantly from the *MOVICS* package [14] and several other R packages (*pathfindR* [62], *fastcluster* [63], *pheatmap* [64], *ComplexHeatmap* [65], *M3C* [66, 67], *rcompanion* [68], *igraph* [69, 70]).

*Firstly, we ran a baseline consensus clustering analysis (more details in section MOVICSbaselineanalysis)with default settings on 10 popular multi-omic clustering algorithms included in MOVICS*: CIMLR [15], iClusterBayes [16], moCluster [17], COCA [18], Consensus Clustering [19], IntNMF [20], LRAcluster [21], NEMO [22], PINSPlus [23] and SNF [24]. Subsequently, we tested individual algorithms for the same multi-omic clustering task (including more methods not available within *MOVICS*, see section Individual method multi-omic clustering). Finally, we examined consensus strategies (see section Consensus analyses). The post-processing pipeline was adapted for each algorithm while maintaining a consistent structure for meaningful comparisons. Deviations from the default settings are noted where applicable (see Supplementary File 1 for more details).

Upon completion of the clustering task, results were visualised using heatmaps annotated with clinical data, and silhouette plots to examine clustering performance. Associations between subtypes and clinical variables were evaluated using non-parametric statistical tests. For mutation data, genes mutated in fewer than 5% of samples were filtered out, and Fisher’s exact test was used to assess subtype associations, with results visualised in oncoprint plots. Differential analysis was conducted using the *limma* package for the standardised RNA-seq, miRNA and methylation data, focusing on features with adjusted *p*-values lower than 0.05 (*p*_*adj*_ *<* 0.05). CNV analysis included calculating the FGA using original copy number counts, with gains and losses annotated using one of two sets of criteria:

- **Simple criteria**: Gains are defined as copy number *>* 2, losses as copy number *<* 2, and normal as copy number equal to 2.
- **COSMIC criteria**: Gains and losses are determined based on ploidy-adjusted thresholds, with gain defined as copy number ≥5 for ploidy ≤2.7 or ≥9 for ploidy *>* 2.7, and loss defined as copy number = 0 for ploidy ≤ 2.7 or below a specific threshold for ploidy *>* 2.7.

In our results, we report CNV patterns based on the more robust COSMIC criteria. Non-parametric tests, such as the Wilcoxon rank-sum test (2 subtypes) or Kruskal-Wallis test (*>* 2 subtypes), were used to compare FGA across subtypes.

Drug sensitivity analysis on the identified clusters and anti-neoplastic drugs was carried out using the *pRRophetic* package [71]. The agremeent between the identified clusters and known subtypes, e.g. ER+/-classification, was measured using four indices, the Rand Index (RI), the Adjusted Mutual Information (AMI), the Jaccard index (JI) and the Fowlkes-Mallows (FM) index. Pathway analysis employed Gene Set Enrichment Analysis (GSEA) [72, 73] to identify subtype-specific pathways, followed by single-sample enrichment scoring via the *GSVA* package [74]. Dimensionality reduction plots, using either PCA or Multi-dimensional Scaling (MDS), and graphs produced by the *igraph* package [69, 70] (where applicable) were generated to explore subtype distributions and visualise similarity matrices. External validation was performed using Nearest Template Prediction (NTP) and Partioning Around Medoids (PAM) [75, 76]. When NTP was successful, these results are used for the analysis on the evaluation cohorts.

Survival analysis for both the TCGA training cohort and the independent TCGA hold-out set was performed in R with an extended wrapper around *TCGAbiolinks*’ `TCGAanalyze survival()` function [41–43]. Clinical end-points (overall survival time and vital status) were imported from *cBioPortal* [77–79], whose annotations are more complete compared to *TCGAbiolinks* data with respect to survival. For every algorithm’s clusters we produced Kaplan–Meier curves and fitted an age-adjusted Cox model; the corresponding figures display the Likelihood Ratio Test (LRT) *p*-value for the cluster term. Alongside the primary comparison we computed four diagnostics: proportional hazards test, variance inflation factor for collinearity, martingale-residual check for linearity of numeric covariates, and detection of infinite coefficients caused by separation. With the exception of wMKL which failes the collinearity criterion, all other algorithms in the hold-out cohort satisfied these criteria, confirming that model assumptions did not influence the main LRT result. However, the proportional hazards tests failed in the training cohort. Results from survival analysis are, therefore, limited.

### Estrogen receptor-based reference analysis

We performed a direct comparison between ER+ and ER-samples in the training set to characterise ER-associated profiles and to provide context for interpreting the multi-omic clustering results. This baseline ER-driven analysis defines the patterns (expression, copy numbers, methylation, mutations, pathways, clinical variable associations) expected from a purely ER-based partition of the data, and the main results are summarised in Figure 2. By establishing this reference, we avoid “over-attributing” these patterns to the clustering algorithms themselves when they may simply reflect accurate partitioning based on ER status. We follow applicable steps from the previously described general pipeline (Section General pipeline). Enriched pathways were identified and visualised using the WEB-based GEne SeT AnaLysis Toolkit [80] to perform gene ontology set enrichment analysis for the differentially expressed genes identified through Differential Gene Expression Analysis (DGEA) between ER+ and ER-samples.

Based on the results presented in Section ER baseline analysis, when a multi-omic algorithm reproduces this ER+/ER-separation together with its expected combination of pathway, mutation (*PIK3CA, TP53, CDH1*) and copy-number (FGA, FGL) differences, this indicates that it is successfully aligning information across RNA-seq, single-nucleotide variant and CNV data. However, this alone does not reveal which omic layers actually drive the partition, since an algorithm clustering primarily on expression could still inherit the correct mutation and copy-number patterns through their correlation with ER status. We therefore complement our multi-omic analysis pipeline with low-dimensional projections, sample relationship heatmaps and other interpretability techniques where applicable, to assess the relative contribution of each data type and/or individual features and to distinguish genuine multi-omic integration from clustering that is predominantly driven by one omic modality.

### MOVICS baseline analysis

*MOVICS* is used to perform multi-omics integrative clustering and visualisation for cancer subtyping research [14]. It provides a unified interface for 10 state-of-the-art multi-omics clustering algorithms: CIMLR [15], iClusterBayes [16], moCluster [17], COCA [18], Consensus Clustering [19], IntNMF [20], LRAcluster [21], NEMO [22], PINSPlus [23] and SNF [24]. *MOVICS* suggests an estimate of the optimal number of multi-omic clusters *k*, then proceeds by using each clustering algorithm to cluster the multi-omic data in *k* subtypes and continues with performing consensus clustering on the outputs.

The first step in the pipeline is to identify the optimal number of clusters for multi-omic data integration. To find the best number of clusters, *MOVICS* employs a Non-negative Matrix Factorization (NMF) approach, which helps simplify the data and detect patterns across different omic layers. The process involves testing various numbers of clusters (ranging from 2 to 10) and assessing how stable the clusters are using two metrics: Cluster Prediction Index (CPI) and the GAP statistic. CPI measures the stability of cluster assignments across repeated runs, and the GAP statistic compares the tightness of clusters to what would be expected in random data. The cluster number *k* for which the sum of the normalised values from both metrics is maximised, is used as the optimal number of clusters.

After fixing the number of clusters at the optimal *k*, each of the ten methods is run with default settings and the resulting cluster labels are assembled into a consensus procedure described by Monti et al. [19]. For every method a binary *N*× *N* co-association matrix is generated (entry = 1 if the two samples fall in the same cluster, 0 otherwise);. These matrices are averaged, yielding a consensus similarity matrix *S* whose values range from 0 (never clustered together) to 1 (always clustered together). The algorithm then converts *S* to a dissimilarity matrix *D* = 1 −*S* and performs Ward’s agglomerative hierarchical clustering using *D* as a distance matrix. The tree is cut at the predefined *k* to obtain the final consensus labels.

### Individual method multi-omic clustering

After the baseline analysis was carried out, a series of different Machine Learning (ML) methods were run using the same training and evaluation cohorts. The output for each method was further analysed as described in section General pipeline and, in more detail, in Supplementary File 1. When the authors of a method mention a way for selecting the optimal number of clusters *k* in the corresponding paper and/or offer such a way in the tool itself, we use that approach - as long as it does not fix values of tunable hyperparmeters - to determine the optimal number of clusters *k*_*opt*_ and cluster the data into *k*_*opt*_ clusters. Otherwise, the determination of the optimal *k* is carried out on a case-by-case basis, mainly depending on the method output (e.g. similarity matrix, latent embeddings etc.).

Multi-omic clustering methods cannot always be allocated into exclusive methodological categories. It is possible that a method’s approach could be classified into multiple of the groups mentioned below, which do not constitute an exhaustive list of method classifications. We provide a broader categorisation here and refer the reader to the Supplementary File 1 to explore each tool’s methodology in more detail.

#### Similarity network methods

In this category, we classify methods whose underlying methodology predominantly depends on the construction of a similarity network between patients/samples (excluding multiple kernel learning methods which are mentioned below). These include: SNF [24], ab-SNF [25], ANF [26], MDICC [28], MSNE [32], NEMO [22], RWR-F [33], RWR-NF [33] and Spectrum [34].

SNF constructs networks of samples (patients) for each available data type and then efficiently fuses these into one network that represents the full spectrum of the underlying data [24]. ab-SNF is following the same approach, but also incorporates feature weighting based on biological knowledge or data-driven criteria [25]. ANF derives a “smoothed” network from each view via *k*-NN-based pruning and weighting and then directly combines those single-pass networks into a final fused affinity, in one iteration (one pass) [26]. MDICC applies network fusion by incorporating weighted local kernels, low-rank and sparsity constraints, and entropy-based integration [28]. MSNE integrates sample similarities by performing random walks on multiple similarity networks and projects them into a low-dimensional space [32]. It employs a random walk strategy designed to prevent comparing edge weights across different similarity networks [32]. NEMO uses the same approach as SNF to create similarity matrices and then proceeds by creating a relative similarity matrix for each omic to adjust similarities based on local neighbourhood data, making the comparison between different omics more consistent [22]. The relative similarity matrices are then averaged to form an Average Relative Similarity matrix [22]. RWR-F constructs a *k*-NN similarity graph for each omics layer, links identical samples across layers to form a heterogeneous multiplex network, and fuses them by running a random walk with restart and the steady-state probabilities give the integrated sample-similarity matrix [33]. RWR-NF follows the same scheme but applies a degree-normalisation to every layer before the walk, ensuring that no single, denser omics network overwhelms the fusion [33]. Finally, Spectrum employs a self-tuning, density-aware kernel to strengthen similarities among shared nearest neighbours, integrates data using tensor product graph diffusion to reduce noise and reveal structures, and introduces an eigenvector-based approach to automatically determine the optimal number of clusters for both Gaussian and non-Gaussian datasets [34].

#### Matrix factorisation methods

Matrix factorisation methods work by decomposing the original data matrix into the product of two lower dimensionality matrices. A widely used approach for matrix factorisation is NMF, which requires the input matrix and the low-dimensional matrices to have strictly non-negative values. Methods included in our work which are classified (mainly) as matrix factorisation methods are: IntNMF [20], MFA [36], MOFA [30] and LRAcluster [21].

IntNMF is a NMF approach to multi-omics where one common basis is used for all modalities, but separate sets of coefficients for each modality’s features [20]. In NMF terminology, in the multi-omic case, the modalities share a basis vector matrix *W*, while each individual modality *i* has its corresponding coefficients matrix *H*_*i*_ [20]. The algorithm converges to a set of basis vectors through multiple iterations [20]. MFA is a multivariate technique that summarises and visualises complex datasets by grouping quantitative or qualitative variables, accounting for each group’s contribution to individual distances while allowing varying numbers and types (continuous, categorical or frequencies) of variables within each group [29, 36]. It effectively is a combination of PCA and Multiple Correspondence Analysis (MCA) [29, 36]. MOFA is a factor analysis method which models the multi-omic matrices with a common factor matrix and individual weight matrices (similarly to IntNMF), but also implements, regularisation, noise modelling and a variational Bayesian framework which allows for the use of specific distributions for the different omic datasets [30]. Finally, LRAcluster iteratively fits a shared low-rank matrix to multiple data matrices (flexible with binary, Gaussian or Poisson distributions), updating it with exponential-family likelihoods so that the matrix’s singular vectors give a common low-dimensional embedding [21].

#### Multiple kernel learning methods

Multiple kernel learning methods use an often predefined set of kernels and learn an optimal linear or non-linear combination of kernels. Methods in this category include: CIMLR [15], wMKL [35] and KLIC [27].

CIMLR learns a measure of similarity between each pair of samples in a multi-omic dataset by combining multiple Gaussian kernels per data type, corresponding to different, complementary representations of the data [15]. It enforces a block structure in the resulting similarity matrix, which is then used for dimension reduction and subsequently, clustering [15]. wMKL is based on the CIMLR idea and further builds on it by integrating prior weights into each kernel, enforcing block structures with rank constraints, and applying graph diffusion to strengthen weak similarities [35]. It effectively handles data heterogeneity by assigning different weights to each omics type, incorporates prognostic information to prioritise clinically relevant features, and improves robustness against kernel parameter sensitivity [35]. KLIC takes as input a list of datasets and builds one kernel per dataset, using consensus clustering [27]. The kernels are then combined through localised kernel *k*-means to obtain the final clustering [27].

#### Graph-based methods

Graph-based methods create graphs based on a mathematical representation of pair-wise sample relationships. Technically, network fusion methods involve some kind of graph-related functionality within their approach, however, here, we focus on methods which are predominantly graph-based: MONET [31]. MONET converts every omics layer into a patient-similarity network and then iteratively grows, splits, or merges groups of patients so as to maximise the total edge weight only in those omics where a given group is unusually cohesive [31]. Evaluated independently, each module may rely on a different subset of omics, so an omics layer ignored for one patient group can still drive the discovery of others [31].

#### Bayesian methods

Bayesian methods for cancer subtyping utilise probabilistic models to integrate various data types and incorporate prior biological knowledge, enabling the identification of distinct cancer subgroups with posterior probabilities. iClusterBayes fits a low-dimensional latent-factor model that captures the shared structure of continuous, count and binary omics datasets, enabling tumour samples to be clustered in the resulting factor space [16]. Its Bayesian formulation simultaneously delivers posterior probabilities for the association of every feature with the latent factors, thereby pinpointing the molecular features most likely to drive the observed clustering [16].

#### Consensus/Ensemble methods

In this section, we allocate methods which employ a form of consensus/ensemble clustering in their approach. These methods include: PINSPlus [23, 37, 38], COCA [18, 39] and the original Consensus Clustering [19]. PINSPlus estimates how often each pair of patients is grouped together in the following scenarios: (i) when data are perturbed, (ii) when using different data types and (iii) when using different clustering techniques; it then partitions patients into subgroups that are strongly connected in all scenarios [23, 37, 38]. COCA is clustering each modality separately either by PAM, hierarchical clustering or *k*-means, determines the best *k* for each modality based on the silhouette index and then aggregates the results in an indicator matrix for the clusters of all modalities [18, 39]. Finally, the original Consensus Clustering combines multiple clustering results from resampled data subsets to identify stable and reliable subgroups [19]. It is essentially unimodal, but is applied in *MOVICS* in a multi-omic setting, by using the concatenated omics as input.

#### Exclusion of methods from individual runs

The following methods were only used in the baseline analysis: moCluster [17], Consensus Clustering [19], IntNMF [20] and PINSPlus [23, 37, 38]. While in the baseline analysis we used all methods available in *MOVICS* with default settings, regardless of whether they consider input distributional properties, individual method runs were thoroughly designed and tuned, and care was taken to ensure the methods used are compatible with binary data. moCluster is built on consensus PCA, i.e. it finds joint latent variables by maximising covariances across blocks that have first been centred, scaled and weighted [17]. Those operations assume continuous (approximately Gaussian) measurements and would not be directly applicable to our case because the SNPs dataset is binary. Similarly, PINSPlus depends on projecting data assumed to be continuous into lower dimensions and cannot currently incorporate appropriate binary data handling without major source code modifications [23, 37, 38]. As described earlier, IntNMF uses a separate coefficients matrix *H*_*i*_ for each modality *i* and a shared basis vector matrix *W* for all modalities. However, because the optimisation minimises a Euclidean (Gaussian) reconstruction error on non-negative continuous values, feeding a 0/1 matrix breaks this assumption: the binary block offers virtually no variance once shifted to non-negativity, so the NMF updates cannot extract meaningful factors and its contribution to the shared *W* becomes numerically unreliable. Although binary NMF methods do exist [81, 82], merging the objective function of the binary case and the continuous (Gaussian) case into one objective for simultaneous integration, is not straightforward. Finally, *MOVICS* uses the Consensus Clustering approach [19] for multi-omic integration by providing the concatenated omics as input. The method essentially is “uni-modal” and treats the input as one matrix, not a series for different modalities. This does not fall within the scope of this work which is mainly focused on intermediate and late integration methods, and is therefore, not used in the individual method runs.

#### Method comparisons

We compare the methods in terms of:

- **Performance**. Performance of unsupervised methods is hard to measure owing to the lack of ground truth labels. Here, we measure contextual silhouette indices (in the “decision space” of each method) and also comment on the biological plausibility of results: agreement with established clinical knowledge (ER status, known mutational patterns and enriched pathways). The space into which silhouette indices are calculated differs for each method in terms of dimensionality and process of generation. Therefore, comparisons across methods are treated with caution. Additionally, high silhouette values do not necessarily indicate clinically relevant clusters (e.g. COCA).
- **Running times & memory**. The aggregated runtime scoring and ranking for every method was determined by calculating the runtime (min) per job (see also Section Aggregated running times), as opposed to the total runtime consumed in order to examine exhaustive hyperparameter scenarios and tune the models. User-implemented parallelisation is taken into account here to allow for fair comparisons between methods. We also report when authors included parallelisation intrinsically to each method, to make it faster. Additionally, we compare runtimes and memory usage across feature and sample perturbation scenarios for a fixed parametrisation (see also Section Robustness, scalability and stability benchmarks).
- **Robustness and stability**. Robustness and stability benchmark jobs were run as described in detail in Section Robustness, scalability and stability benchmarks. Robustness was defined as agreement between the results of each method under a series of perturbations, while stability was defined as agreement between the results of each method under different 90% sub-samplings of the full dataset.
- **Data handling**. All the methods used in the individual runs are capable of handling binary data. It is possible that a method can handle binary input i) by default, ii) upon appropriate input preprocessing (e.g. providing a distance matrix calculated with a binary distance metric for binary data) or iii) after a minor modification of source code which does not otherwise compromise the underlying methodology (e.g. by allowing for the estimation of diverse distance metrics depending on input data type). Regarding missing values, this scenario was not examined as an ablation because the majority of the methods cannot deal with partial omics/missing values. However, methods with pre-built functionality to deal with types of missingness are reported on Table 2.
- **Intepretability**. Can the final clustering results be attributed (preferrably in a quantitative way) to specific omics and/or features? A few methods offer functionality to quantify modality contribution (omic-level importance) and/or rank features (feature-level importance) for improved interpretability of results. When none of these contextual functionalities is included in a method, omic-level contribution is restricted to visual inspection of the dimensionality reduction plots and feature-level contribution is restricted to the results of differential analysis (expression, methylation, miRNA).
- **User-engagement/Documentation**. Here, we assess the methods by ranking the available support in each case: Packages with instructions *>* Installable GitHub repos (R) / Repos with appropriate `.yml` or `.toml` files (Python) *>* Repos which require a good deal of manual work *>* Repos which contain outdated code which needs major modifications to run.

An overview of these comparisons is shown in Table 2.

### Case-by-case method tuning

In each case, parameter tuning was carried out on parameters which do not have clearly established defaults, e.g. fixed values selected by original authors after extensive trials. At the same time, the ranges of tested parameters and values were selected while considering the total expected running time for a method’s individual runs/jobs. The possible number of multi-omic clusters for each method was consistently set to 2, 3, 4, 5, 6, 7, 8, 9, 10 . Wherever the input consists of the distance matrices of the individual omics, and unless specified otherwise, Euclidean distances were calculated for all modalities other than SNPs, for which we used the binary distance implemented in `dist(…, method = “binary”)` in R, i.e. the proportion of loci where the two samples disagree (one carries a variant and the other does not) among all loci where at least one of them carries a variant, with shared absences ignored.

Similarity network methods including SNF, ab-SNF and NEMO were tuned on two key hyperparameters: the number of nearest neighbours ∈ {10, 15,…, 50} and the regularisation hyperparameter *σ* ∈ {0.3, 0.4,…, 0.8}, while the maximum number of iterations was fixed on *t* = 50. ab-SNF requires a weighting scheme to boost input signals. We weight each continuous feature in proportion to its median absolute deviation (weights normalised to sum to one) and give binary SNPs a weight of 0.8 if the locus maps to a COSMIC breast-cancer driver gene and 0.2 otherwise, using the resulting weights to compute weighted squared Euclidean (continuous) or weighted binary distance matrices. ANF used the same range of number of neighbours (*σ* not applicable here) and fixed values for the ANF affinity parameters *α* = *β* = 1*/*6 and the separate fusion random walk *alpha*_*fusion*_ = (1, 1, 0, 0, 0, 0, 0, 0) vector (values suggested by the authors) [26]. In the case of RWR-F and RWR-NF we use the aforementioned grid of number of neighbours and *σ*, and fix some additional parameters: number of maximum iterations (1000, source code default), *γ* = 0.7, random walk neighbours (10, source code default; RWR-NF only), random walk *α*_*RW*_ (0.9, source code default; RWR-NF only), random walk *β*_*RW*_ (0.9, source code default; RWR-NF only). For MDICC we tuned the number of subspace dimensions (top *c* eigenvalues: {2, 3, …, 10}) and the number of nearest neighbours used when computing the adaptive entropy-sparsity regularisation parameters ({41, 42, 43, 44} ; range used in the publication) [28]. We fix the number of neighbours used for the local affinity matrices to 18 (value used in the publication) [28]. MSNE was exhaustively tuned over the consistent neighbourhood size {10, 20,…, 50}, random-walks per node {50, 100, 200}, embedding dimension {50, 100, 200}, Skip-gram window {5, 10, 15} and walk length {20, 30, 40 }, ranges that match the authors’ recommended search space [32], while all other arguments remained at default settings. The number of clusters was fixed at 5 which is expected in breast cancer (5 intrinsic subtypes) in order to reduce the already very high number of jobs, while achieveing high resolution (this *k*-fixing was only applied in MSNE). Finally, for Spectrum, we use the adaptive density-aware kernel, the multimodality gap method to determine the best *k* for *k*-means clustering and implement kernel tuning. We set the number of nearest neighbours to use *σ* parameters to 3 (default), the number of nearest neighbours to use for the common nearest neighbours to 7 (multimodality gap method parameter; default), fraction to find the last substantial drop to 2 (multimodality gap method parameter; default), the threshold of points ahead to keep searching to 7 (multimodality gap method parameter; default) and the number of diffusion iterations to 5 (default: 4). We tune the number of *K*-nearest neighbours used when making the intermediate *K*-NN graphs over the same range we used throughout the similarity network methods.

Regarding multiple kernel learners (CIMLR, KLIC and wMKL) we used squared Euclidean distances for RNA-seq, methylation, CNV and miRNA, as described in the original methodologies [15, 27, 35]. For CIMLR we use the same range of neighbours as before. In wMKL, the number of neighbours is hard-coded to be max(⌈*n/*20, 10⌉), where *n* is the number of samples (in our case: max(⌈625*/*20, 10⌉) = 32 neighbours are used). In KLIC, consensus matrices were built using the COCA approach (see Supplementary File 1), by setting the maximum number of clusters in each modality to 5, *B* = 250 bootstrap iterations and the subsampling proportion to 80%. Subsequently, KLIC’s local kernel *k*-means approach was run for all possible combinations of numbers of clusters per modality, and the number of global clusters ranging from 2 to 10 as before, for a maximum of 100 iterations (1024 combinations). In wMKL, we used the same feature-weighting system as in ab-SNF.

With respect to matrix factorisation methods: We restricted the choice of latent dimensions to the range {2, 3, …, 10} for LRAcluster, and selected the optimal number of dimensions (*r*_*opt*_ = 7) by inspecting the corresponding “explained variance” plot and selecting the number of dimensions where an elbow is observed and slow incremental increase follows in explained variance from then on (Supplementary Figure 22d). Due to long running times, MFA was run at the highest proportion of features per modality (33%) that allowed job completion within 12 hours. The subsets were selected based on highest mean absolute deviation (continuous modalities: RNA-seq, CNV, methylation and miRNA), while for SNPs we kept the COSMIC cancer drivers plus the top 33% of other SNPs, ranked by proportion of samples that carry a mutation. The maximum number of components was set to 200. Similarly to LRAcluster, by visually inspecting the “explained variance” plot we selected 11 as the optimal number of dimensions for MFA. We also used the Unit Invariant Knee (UIK) method from the *inflection* package [83, 84] to determine elbow points in a more objective way. UIK identifies 37 as the elbow point in terms of eigenvalues and variance explained, as well as 134 as the elbow point in cumulative variance explained. All three cases of latent dimensions were tested before selecting the optimal number of clusters as described in Section Optimal number of clusters. Finally, we fit MOFA models with possible number of factors *F* ∈ {2, …, 10, 15, 20, 25} . The models were run on a GPU, with a fixed number of maximum iterations (20,000). The critical point in the cumulative explained variance (Supplementary Figure 31) and consistently low inter-factor correlations up to *F* = 7 pointed to *F*_opt_ = 7. A 25-factor run with Automatic Relevance Determination (ARD) -a Bayesian shrinkage prior that automatically suppresses non-informative factors-reproduced the same *R*^2^ profile, confirming this choice for *F*_opt_.

For iClusterBayes, we performed a grid search over *s*_dev_ ∈ {0.005, 0.01, 0.015, 0.02, 0.025, 0.03, 0.05} and *β*_var_ ∈ {0.1, 0.2, 0.3, 0.4, 0.5, 0.8, 1, 1.25, 1.5, 2, 2.5, 3}, fixing *n*_burnin_ = 1200, *n*_draw_ = 1800, `thin`=3, pp_cutoff_ = 0.5 and *γ* = (0.5, 0.5, 0.5, 0.5, 0.5) as in the original specification (which also allows for running times below 12 hours). Each of the 7 ×12 = 84 pairs was run for *K* = 2:10 (`K=1:9` in code) on a HPC. For every fit we recorded the median acceptance rates 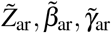 and defined

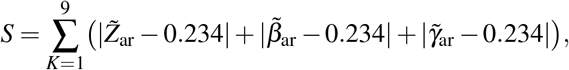

using 0.234 as the optimal random-walk acceptance target [85, 86]. A penalty of +0.05 was added whenever a median laid outside [0.1, 0.8]. The lowest penalised score was obtained for *s*_dev_ = 0.015, *β*_var_ = 0.5.

In MONET, we computed a 625 × 625 Pearson-correlation matrix for each omic (equivalent to *φ* -correlation for binary data), and, for a sensitivity check, an “offset” version (−0.2 shift) as in the original MONET paper [31]. We then ran the main MONET loop twice: once with the original versions and once with the “offset” versions of the correlation matrices. We fixed the maximum number of iterations to 10,000, the number of different seeds to 100, the number of samples in a seed to 10, the minimum size of modules to 10, the maximum number of patients per action to 10 (upper bound on patients moved per MONET step), the percentile of removed edges to 80% (drops the lowest 20% of edge weights), and applied no global re-centering of weights. Weight histograms revealed that the 0.2 offset drove *>* 95% of edges negative in every omic, whereas the raw matrices were well-balanced. We, therefore, kept the “no-offset” solution.

Finally in COCA, we first built a 625 ×625 Multi-Omic Consensus (MOC) matrix with `buildMOC()` using all five omics (*M* = 5) and evaluating *K*∈ {2, …, 10} ; single-omic partitions were produced with Ward hierarchical clustering on Jaccard distance for SNPs (“binary”) and Euclidean distance for the four continuous omics. The resulting MOC was converted to a Jaccard dissimilarity matrix and reclustered with Ward’s method, yielding a consensus dendrogram.

### Optimal number of clusters

When a method has a pre-built way to determine the optimal number of clusters *k*_*opt*_, we use that to select the optimal partition. When the output is a similarity/dissimilarity matrix, this is used as input for spectral clustering for a range of *k* ∈ {2, …, 10} and the *k*_*opt*_ is determined based on the silhouette index (see below). When the output of a method is instead embeddings of the samples in a latent space, we use the *M3C* package [66, 67] to carry out Monte-Carlo reference-based *k*-means clustering on the embeddings’ space, for a range of *k* ∈ {2,…, 10}.

We used a contextual version of the silhouette index to assess the clustering output of each algorithm. In the case of algorithms that produce some sort of embeddings, Euclidean distances were calculated in the embeddings’ space and fed into the silhouette formula. When clustering was carried out on a latent space projection/embeddings of the original data using the *M3C* package, the output of main `M3C()` function was used to infer statistical significance and stability of the clustering. For similarity network methods which suggest spectral clustering as the final step, the distances were measured in the eigenspace that is intrinsically computed during spectral clustering and on which the final partition is based. Otherwise, if a final multi-omic matrix with symmetric relationships is provided, it is converted to a distance matrix on a case-by-case basis and used for silhouette index calculation.

In similarity network methods (ab-SNF, RWR-F, RWR-NF, SNF), the final affinity matrices were significantly influenced by the chosen number of nearest neighbours. The affinity matrices for a fixed value of nearest neighbours and varying values of the regularisation parameter *σ* were practically equivalent. Therefore, we set *σ* to 0.5 (halfway between 0 and 1) and kept the nine corresponding fused affinity matrices, for *σ* = 0.5 and a varying number of nearest neighbours. Each of these matrices was used as input for the built-in optimal *k* estimation method provided by SNF [87], and the optimal partition was selected based on the average silhouette width after runnning spectral clustering on the “fused affinity matrices - estimated optimal *k*” pairs (optimal nearest neighbour values: ab-SNF/10, RWR-F/50, RWR-NF/10, SNF/25, optimal *k* = 2 for all of them). In NEMO, both the number of neighbours and *σ* significantly influence the resulting fused affinity matrix. All fused affinity matrices were used as input for the optimal *k* estimation method [87] and the final partition was selected based on average silhouette width (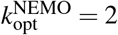 for 10 nearest neighbours). In ANF, only the number of nearest neighbours is tuned, and different values yield sufficiently different matrices. The same process (excluding the *σ* filtering) is followed to determine the optimal 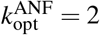for 30 nearest neighbours. For MDICC, the optimal 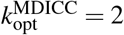 was determined by selecting the clustering whose final similarity matrix yielded the highest average silhouette width (top *c*_opt_ = 6 eigenvalues and number of neighbours: 43). The same exhaustive approach is followed in the case of MSNE, only this time the silhouette is calculated on the provided embeddings (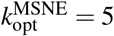, neighbourhood size: 50, random-walks per node: 200, embedding dimensions: 50, Skip-gram window: 5 and walk length: 40). In Spectrum, the clusterings obtained by the different *K*-NN values are assessed in terms of silhouette widths in the corresponding eigenspaces to determine the optimal *K*-NN/*k* pair (20/2).

In CIMLR, the final similarity matrix and clustering was chosen after the *t*-SNE embeddings produced for different values of nearest neighbours were used as input for reference-based Monte-Carlo consensus *k*-means clustering [66], using the *M3C* package [67]. We set the number of Monte Carlo iterations to 100, the number of resampling iterations for both reference and real data to 250, the objective function to “entropy” (default), the maximum number of clusters to 10, the reference data generation method to “reverse PCA” (default) and the fraction of points resampled in each iteration to 0.8 (default). This parameter configuration was used whenever the *M3C* package is used in this work. The total of 81 clusterings was examined in terms of stability, as measured by the Relative Cluster Stability Index (RCSI) and statistical significance (clustering Monte-Carlo *p*-value). The best performance is attained for 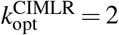 and 15 nearest neighbours.

The parameter combination that achieves the highest average silhouette width in KLIC is and yields two clusters. In wMKL, the omic-specific kernels are merged into one similarity graph, then every possible value of number of clusters is scored with CIMLR’s cluster-quality metric, and 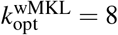 is the *k* beyond which the score stops improving—i.e. the point that best balances having enough clusters without over-splitting the data (see Supplementary File 1 for more details).

For all three matrix factorisation methods, the optimal number of clusters was determined using the previously described *M3C* approach [66, 67]. The optimal number of clusters 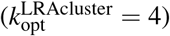 for LRAcluster results was determined using the low-dimensional coordinates of the samples as input (*r*_opt_ = 7). The same approach was followed for MFA, but using three distinct versions of the low-dimensional projections (with 11, 37 and 134 dimensions; see Section Case-by-case method tuning) and determining the final low-dimensional representation (11 dimensions) and number of clusters 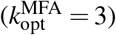, by considering stability and statistical significance metrics across all runs. Similarly, in MOFA, the *F*_opt_ = 7-factor solution was used as input for *M3C* clustering 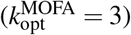).

Within the selected hyperparameter setting in iCluster-Bayes, the minimum Bayesian Information Criterion (BIC), a deviance-ratio elbow and a well-structured heatmap all agreed at *K* = 5; we therefore selected *K*_opt_ = 5 and the corresponding cluster labels for down-stream analyses. In MONET, the final module set ℳ contained three non-overlapping modules, which we mapped to {1, 2, 3} to obtain 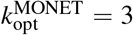 sample clusters. For each candidate cut *K* ∈ {2, …, 10} of the COCA consensus dendrogram, we stored the resulting cluster labels and computed the mean silhouette width 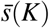 using the Jaccard-based dissimilarity matrix. We also computed Tibshirani’s gap statistic Gap(*K*) with *B* = 100 bootstrap reference draws, using the same hierarchical clustering cut as the clustering function. Across the tested range, 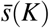 increased up to *K* = 6 (with near-maximal values at *K* = 5 and *K* = 6), while the gap statistic became ill-defined for *K*≥ 6 due to degenerate within-cluster dispersion estimates. Although *K* = 5 would typically be favoured by 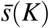, the corresponding partition was highly imbalanced (cluster sizes: 549, 68, 3, 1, 4), indicating that the additional clusters were effectively singletons/outliers rather than stable modules. We therefore selected a parsimonious cut of 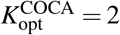 for all downstream analyses.

### Feature/modality importance

This section discusses the usage of built-in funcionality for feature-level and/or omic-level interpretation, excluding inspection of dimensionality reduction plots. For Similarity network methods, we extracted feature rankings (top 1000 features) and weighted fused graphs using the approaches described in the original SNF publication [24]. For MDICC, the source code of the main function needed to be slightly modified to allow for the return of omic-level kernel weights which quantify the contribution of each modality to the final results.

A similar kernel summing approach was followed to estimate omic-level contributions in the CIMLR case. KLIC generates a *n*× *p* weight matrix, where *n* is the number of samples and *p* the number of modalities. We rank modalities in terms of importance by taking the average weight per modality. For MONET, we inspect the excess within-module connectivity (the degree to which samples in a module are more tightly linked than the cohort-wide average for each omic), to investigate whether modules are dominated by subsets of modalities. In MOFA, we examine feature weights in the loading vectors of each factor to quantify feature importance per factor and, at the omic level, examine plots of explained variance per factor. Additionally, we look at the distribution of three key clinical variables in the different factors: ER status, HER2 status and tumour stage. We follow similar steps in MFA to estimate feature- and omic-level contribution in the obtained factors, as well as assess the factors’ relationships with clinical variables.

### Consensus analyses

The clustering results from all individual methods were used as input to perform consensus clustering similar to the baseline *MOVICS* pipeline. We collated the class labels produced by all 18 individual methods and, over 100 bootstrap iterations, in each of which 80% of the tumours were subsampled, calculated a pair-wise co-association value, i.e. the fraction of methods that placed each sample pair in the same class. The mean co-association matrix was converted to a “1 – consensus” dissimilarity, and used as input for average-linkage hierarchical clustering. The final partition was chosen as the value of *k* (evaluated for *k* = {2 … 10 }, *k* ∈ ℕ) that maximised the average silhouette width. The output was then examined as described in Section General pipeline. A separate consensus analysis was then run for methods yielding more than 2 clusters in their corresponding output, to examine how the consensus would be affected when considering only more granular clusterings. The preprocessing, consensus clustering and downstream analyses remained the same as in the case of the consensus analysis on all the clustering results.

### Benchmarks

#### Aggregated running times

A first running time comparison was carried out by aggrgeating results from the individual runs of each method. The methods were ranked according to the mean wall-clock time required to execute a single analytical job, defined as one call to the core multi-omic integration routine provided by a given package or library. Several algorithms require hyperparameter optimisation, which is implemented either by looping over an exhaustive grid or by submitting multiple jobs to a HPC cluster. In both situations, we normalised for the number of hyperparameter configurations: the total elapsed time for the full grid search (or the sum of wall times for all HPC array jobs) was divided by the number of parameter combinations explored. The resulting average per-job runtime forms the basis of the cross-method ranking (Figure 4e). We note in the case of MFA that a subset of features (33%) was used for each modality, for the method to run within the 12-hour limit (the full set of features would extend the runtime beyond the limit) and, for MOFA, that a GPU was used to accelerate the jobs significantly.

#### Robustness, scalability and stability benchmarks

We excluded MFA, RWR-NF, and MOFA from comparative runs: MFA was originally evaluated on a non-matching subset (as described above, and extending it to larger/smaller subsets would exceed memory and the 12 h wall-time limit), RWR-NF did not complete within 12 h on all subsets, and MOFA’s GPU-based execution is not directly comparable to Central Processing Unit (CPU)-only methods. All benchmark jobs were run single-threaded on one CPU per task to reduce variability (e.g., BLAS/OpenMP threads fixed to 1 and `srun --hint=nomultithread --cpu-bind=cores`). Runtime and peak memory were measured consistently by profiling only each method’s *core* optimisation step (e.g. network fusion, factorisation, kernel learning etc.) using system-level tooling (e.g., `/usr/bin/time -v` for elapsed time and Max RSS), while excluding downstream post-processing not intrinsic to the method.

Robustness was assessed with hyperparameters fixed to previously tuned optima, using ARI to quantify agreement with full-data results under two perturbation families: (i) *feature perturbations*, retaining modality-wise top-ranked features from a method-agnostic ranking at fixed centiles (10%, 20%, 50%, 75%, 90%); and (ii) *sample perturbations*, evaluating predefined sample subsets shared across methods at fractions (10%, 20%, 50%, 70%, 90%) with 10 replicates per fraction (stratified to preserve ER+/ER-proportions), summarising robustness by the median ARI across replicates. Resampling stability was quantified at the 90% sample level via pair-wise ARI between replicate clusterings (computed on overlapping samples), summarised by the median and interquartile range. Scalability was characterised by fitting method-specific log– log power-law regressions of runtime versus effective feature count *p* (feature perturbations) or sample size *n* (sample perturbations), reporting scaling exponents with confidence intervals, *R*^2^, and corresponding *O*(·) notation (see also Supplementary Tables 4,5).

#### Benchmarking limitations

We took extensive steps to ensure fair and reproducible comparisons across methods. Nevertheless, as with any large-scale benchmarking on shared HPC infrastructure, there remains a possibility of unobserved system-level effects (e.g., scheduler variability, background contention or sporadic node-specific issues) influencing individual runs. We therefore interpret absolute runtime and memory values with appropriate caution, while emphasising that all results reflect our best effort to standardise conditions and minimise confounding variability. The power-law scalability fits are empirical approximations based on a small number of subset levels (five effective sizes per method and perturbation type). Consequently, the estimated scaling exponents should be interpreted as coarse summaries over the tested range rather than definitive asymptotic complexities. We report the coefficient of determination (*R*^2^) alongside each exponent, enabling readers to gauge how well the log–log linear model captures the observed runtime trend (Supplementary Tables 4,5).

We also emphasise that the benchmarking results are derived from one-core/no multithreading scenarios. A few methods offer intrinsic parallelisation or are amenable to parallelised runs. Total runtime decreases a lot if the user’s system (either local or HPC) allows for such setup. Total runtime also depends on hyperparameter choices and the tuning strategy. In particular, methods that use multiple random initialisations, expose optimisation settings (e.g., learning rate), or require an explicit number of iterations can vary substantially in computational cost depending on the selected values. Therefore, although the benchmarks do offer a straightforward comparison between methods with respect to runtimes (given the originally derived optimal parametrisation), the aggregated runtimes and corresponding ranking in Table 2 offer a more “real-world” comparison between runtimes.

## Declarations

### Data availability

All data and code to reproduce the analysis of this survey are openly available on GitHub at https://github.com/sionaris/MultiOmicsSurvey and Zenodo). An interactive R Shiny app to explore the results of our study is available in https://github.com/sionaris/MO_survey_Shiny.

### Authors’ contributions

A.S. and N.S. conceived and designed the analysis. A.S. implemented and ran the analysis (preprocessing, method runs, post-hoc analysis, consensus analysis, cross-method comparisons). A.S. and N.S. analysed the findings. All authors wrote and edited the manuscript. J.A. and N.S. supervised the work.

## Acknowledgements

We thank Dr. Steven Bell, Katrina J.Q. Xian and Dr. Charlotte King for their insights on the design of molecular data preprocessing. We thank Dr. Md Mostafa Kamal Sarker, Dr. Gurpreet Ghattaoraya and Dr. Nikolaos Demiris for their constructive feedback on the write-up of this manuscript. The results shown here are in part based upon data generated by the TCGA Research Network: https://www.cancer.gov/tcga.

## Abbreviations

**Acronyms**

ab-SNF: association-signal-annotation boosted Similarity Network Fusion
AMI: Adjusted Mutual Information
ANF: Affinity Network Fusion
ARD: Automatic Relevance Determination
ARI: Adjusted Rand Index
ATP: Adenosine Tri-Phosphate
BIC: Bayesian Information Criterion
CC: Consensus Cluster
CIMLR: Cancer Integration via Multikernel LeaRning
CNV: Copy Number Variant
COCA: Cluster Of Cluster Assignments
COSMIC: Catalogue Of Somatic Mutations In Cancer
CPI: Cluster Prediction Index
CPU: Central Processing Unit
DGEA: Differential Gene Expression Analysis
ER: estrogen receptor
FGA: Fraction of Genome Altered
FGG: Fraction of Genome Gained
FGL: Fraction of Genome Lost
FM: Fowlkes-Mallows
GPU: Graphics Processing Unit
GSEA: Gene Set Enrichment Analysis
HER2: Human Epidermal growth factor Receptor 2
HPC: High-Performance Computing
HRD: Homologous Recombination Deficiency
HUGO: Human Genome Organization
IC10: integrative clusters
IntNMF: Integrative Non-negative Matrix Factorization
KLIC: Kernel Learning Integrative Clustering
LRAcluster: Low Rank Approximation clustering
LRT: Likelihood Ratio Test
MC: Multi-Consensus
MCA: Multiple Correspondence Analysis
MDICC: Multi-omics Data Integration for Clustering to identify Cancer subtypes
MDS: Multi-dimensional Scaling
MFA: Multiple Factor Analysis
miRNA: microRNA
ML: Machine Learning
MOC: Multi-Omic Consensus
MOFA: Multi-Omics Factor Analysis
MONET: Multi Omic clustering by Non-Exhaustive Types
mRNA: messenger RNA
MSNE: Multiple Similarity Network Embedding
NEMO: NEighborhood-based Multi-Omics clustering
NMF: Non-negative Matrix Factorization
NMI: Normalised Mutual Information
NTP: Nearest Template Prediction
PAM: Partioning Around Medoids
PCA: Principal Component Analysis
pCR: pathological Complete Response
PINSPlus: Perturbation clustering for data INtegration and disease Subtyping
PR: progesterone receptor
RCSI: Relative Cluster Stability Index
RI: Rand Index
RNA-seq: RNA sequencing
RWR-F: Random Walk with Restart for multi-dimensional data Fusion
RWR-NF: Random Walk with Restart and Neighbor information-based multi-dimensional data Fusion
SNF: Similarity Network Fusion
SNP: Single Nucleotide Polymorphism
TCGA: The Cancer Genome Atlas
UIK: Unit Invariant Knee
wMKL: weight-boosted Multi-Kernel Learning

