## Supplementary File 1 for "Consensus Through Diversity: A Comprehensive Benchmark of Multi-Omic Approaches for Precision Breast Oncology"

### Contents

**Acronyms** **5**

**1 Introduction** **8**

**2 Compiling a list of methods** **9**

**3 Preprocessing** **11**

### Acronyms

*ab-SNF* association-signal-annotation boosted Similarity Network Fusion
*AMI* Adjusted Mutual Information
*ANF* Affinity Network Fusion
*ANOVA* ANalysis Of VAriance
*ARD* Automatic Relevance Determination
*ARI* Adjusted Rand Index

*BCC* Bayesian Consensus Clustering
*BIC* Bayesian Information Criterion

*CC* Consensus Cluster
*CIMLR* Cancer Integration via Multikernel LeaRning
*CNV* Copy Number Variant
*COCA* Cluster Of Cluster Assignments
*COSMIC* Catalogue Of Somatic Mutations In Cancer
*CPCA* Consensus Principal Component Analysis
*CPI* Cluster Prediction Index
*CPU* Central Processing Unit

*DEG* Differentially Expressed Gene
*DGEA* Differential Gene Expression Analysis

*ER* estrogen receptor

*FDR* False Diacovery Rate
*FGA* Fraction of Genome Altered
*FGG* Fraction of Genome Gained
*FGL* Fraction of Genome Lost
*FM* Fowlkes-Mallows
*FPKM* Fragments Per Kilobase Million

*GPU* Graphics Processing Unit
*GSEA* Gene Set Enrichment Analysis
*GSVA* Gene Set Variation Analysis

*HER2* Human Epidermal growth factor Receptor 2
*HPC* High-Performance Computing
*HUGO* Human Genome Organization

*IDC* Invasive Ductal Carcinoma
*IGP* In-Group Proportion
*ILC* Invasive Lobular Carcinoma
*IntNMF* Integrative Non-negative Matrix Factorization
*iPF* integrative Phenotypic Framework
*IS-k-means* Integrative Sparse k-means

*JIVE* Joint and Individual Variation Explained
*JLV* Joint Latent Variable

*KLIC* Kernel Learning Integrative Clustering

*LRAcluster* Low Rank Approximation clustering

*MCA* Multiple Correspondence Analysis
*MCIA* Multiple Co-Inertia Analysis
*MCMC* Markov Chain Monte Carlo
*MDICC* Multi-omics Data Integration for Clustering to identify Cancer subtypes
*MDS* Multi-dimensional Scaling
*MFA* Multiple Factor Analysis
*miRNA* microRNA
*MOC* Multi-Omic Consensus
*MOCA* Multiple Omics data integrative Cluster Analysis
*MOFA* Multi-Omics Factor Analysis
*MONET* Multi Omic clustering by Non-Exhaustive Types
*MSNE* Multiple Similarity Network Embedding

*NEMO* NEighborhood-based Multi-Omics clustering
*NMF* Non-negative Matrix Factorization
*NMI* Normalised Mutual Information
*NNDSVD* Non-Negative Double Singular Value Decomposition
*NTP* Nearest Template Prediction

*PAM* Partioning Around Medoids
*PAM50* Prediction Analysis of Microarray 50
*PAMOGK* Pathway-based MultiOmic Graph Kernel
*PARADIGM* Pathway Recognition Algorithm using Data Integration on Genomic Models
*PCA* Principal Component Analysis
*pCR* pathological Complete Response
*PINSPlus* Perturbation clustering for data INtegration and disease Subtyping
*PR* progesterone receptor

*RCB* residual cancer burden
*RCSI* Relative Cluster Stability Index
*RGCCA* Regularised Generalised Canonical Correlation Analysis
*RI* Rand Index
*RNA-seq* RNA sequencing
*RPKM* Reads Per Kilobase Million
*RWR-F* Random Walk with Restart for multi-dimensional data Fusion
*RWR-NF* Random Walk with Restart and Neighbor information-based multi-dimensional data Fusion

*SGCCA* Sparse Generalised Canonical Correlation Analysis

*SIMLR* Single-cell Interpretation via Multi-kernel LeaRning
*SNF* Similarity Network Fusion
*SNP* Single Nucleotide Polymorphism

*TCGA* The Cancer Genome Atlas
*TPM* Transcripts Per Million

*UIK* Unit Invariant Knee

*wMKL* weight-boosted Multi-Kernel Learning
*WSNF* Weighted Similarity Network Fusion

### 1 Introduction

In this supplementary file, we describe our methodology in more detail. All work has been performed in R [1] version 4.4.0 (2024-04-24), platform: x86_64-w64-mingw32/x64, running under: Windows 11 x64 (build 22631). The Python versions and environment setups are explicitly mentioned for each Python method. For each script in our repository, the corresponding `sessionInfo()` outputs in R and `.yml`/`.toml`/`requirements.txt` and/or poetry files in Python, are provided.

**Custom functions:** In some parts of the code, custom functions are used that are either defined in-place or loaded into the RStudio environment by sourcing scripts specifically created for this purpose. Two such scripts exist in the repository:

1. `Scripts/automated_scripts/custom_functions.R`, which contains a series of useful functions that are defined in this file and help us keep our code more concise for the main analysis scripts.
2. `Scripts/automated_scripts/modified_MOVICS_functions.R`, which contains updated functions from the *MOVICS* package [2]. The updates we introduce only ensure functionality in the presence of more recent versions of accompanying packages or include some additional automations for report generation.

Custom Python scripts are included in the corresponding method's folder in the repository. For all processes that incorporate randomness, we use `RNGversion("4.2.2")` and `set.seed(123)` in R and the same random seed (`123`) in Python, where possible. Whenever a process has been run on a High-Performance Computing (HPC) cluster, this is explicitly mentioned. When not specified, the analysis has been run on a Windows machine. Some parts of the analysis are highly demanding in terms of processing power and/or RAM. The upper limit of RAM provided by the Windows machine for this analysis was 25GB and up to 10 cores at a time were used in parallel jobs within the Windows machine.

For HPC cluster runs, the maximum time limit for a process (accounting for all implemented parallelisation) was 12 hours. Any job which would require more than 12 hours of running time was excluded from the analysis. The 12-hour limit is applicable to the majority of free tiers for university HPC clusters.

The descriptions of the different methodologies across algorithms are adapted from the original publications. The notation is not necessarily consistent across algorithms; each description has its own scope.

#### 2 Compiling a list of methods

We compiled a list of methods described and/or mentioned by name in a series of reviews on unsupervised multi-omic analysis [3–14]. Articles that cited the reviews or were deemed similar by the PubMed database (https://pubmed.ncbi.nlm.nih.gov/) and ConnectedPapers (https://www.connectedpapers.com/) were also considered. After compiling an initial list of methods, we then filtered this list for methods which:

1. are available in either **Python or R**
2. are unsupervised and can be used for subtyping
3. are flexible to work with both numerical and categorical multi-omic data

Regarding the final criterion, by "flexible" we mean that the methods are either adaptable to the data type or their source code can be slightly modified to incorporate the analysis of binary data, **without compromising the logic of the original method**.

**For each method we record and report:**

- The underlying approach
- Running times for the main part of the method
- Types of data used as input and required transformations if applicable
- Software required
- Generalizability and mapping to external cohort capabilities
- Interpretability

Table 1 presents all unsupervised methods available in either R or Python, which can be used for multi-omic subtyping. The "**Verdict**" column states whether the method was selected for this work or not. Justifications for the excluded methods, follow below.

With respect to the excluded methods:

- **BCC & Clusternomics:** There were consistent errors for both binary data and continuous data, due to some hard-coded checks in the source code (e.g. not capable to handle low variance etc). In the GitHub repository of MONET, the authors compared their method against Bayesian Consensus Clustering (BCC) and Clusternomics, and provide their modifications so that errors are avoided. These modifications did not work in our case, possibly due to newer versions of R.
- **iClusterPlus:** We ran iClusterBayes which is building on iClusterPlus and is more flexible with heterogenous data distributions.
- **iPF:** integrative Phenotypic Framework (iPF) is designed for 2 omics datasets. For 3 or more datasets, all pairwise comparisons must be considered first before proceeding. The method is outdated and not well-maintained.
- **IS-k-means:** Integrative Sparse k-means (IS-k-means) is based on concatenation (early fusion). The implementation of functionality for the handling of binary data is not straightforward, particularly due to the sparsity bits of the algorithm.
- **JIVE:** Joint and Individual Variation Explained (JIVE) is relatively old and not well-maintained. It cannot incorporate binary modalities straightforwardly.
- **MCIA:** Multiple Co-Inertia Analysis (MCIA) cannot incorporate binary modalities straightforwardly. We use Multiple Factor Analysis (MFA) instead.

**Table 1.** List of unsupervised multi-omic subtyping methods.

| Algorithm | Link | Software* | Verdict |
|---|---|---|---|
| ab-SNF [15] | GitHub | R | ✓ |
| ANF [16] | GitHub | R | ✓ |
| BCC [17] | GitHub | R | ✗ |
| CIMLR [18] | GitHub | R | ✓ |
| Clusternomics [19] | GitHub | R | ✗ |
| COCA [20–22] | CRAN*** | R | ✓ |
| Consensus Clustering** [23, 24] | Bioconductor | R | ✓ |
| iClusterBayes [25, 26] | Bioconductor | R | ✓ |
| iClusterPlus [26, 27] | Bioconductor | R | ✗ |
| IntNMF (iNMF)** [28, 29] | CRAN | R | ✓ |
| iPF [30] | Link to download | R | ✗ |
| IS-k-means [31] | GitHub | R | ✗ |
| JIVE [32] | CRAN | R | ✗ |
| KLIC [22] | CRAN | R | ✓ |
| LRAcluster [33] | Source code | R | ✓ |
| MCIA [34, 35] | Bioconductor | R | ✗ |
| MDICC [36] | GitHub | R + Python+ | ✓ |
| MFA [37] | CRAN | R | ✓ |
| MixKernel [38] | CRAN | R | ✗ |
| MoCluster** [39] | Bioconductor | R | ✓ |
| MOFA [40] | Website | Python | ✓ |
| MONET [41] | GitHub | Python | ✓ |
| MSNE [42] | GitHub | Python | ✓ |
| NEMO [43] | GitHub | R | ✓ |
| PAMOGK [44] | GitHub | Python | ✗ |
| PARADIGM [45] | GitHub | Python | ✗ |
| PINSPlus** [46–48] | CRAN | R | ✓ |
| rGCCA [49, 50] | Bioconductor | R | ✗ |
| sGCCA [51] | Bioconductor | R | ✗ |
| RWR-(N)F [52] | GitHub | R | ✓ |
| SIMLR [53] | GitHub | R | ✗ |
| SNF [54] | CRAN | R | ✓ |
| Spectrum [55] | CRAN | R | ✓ |
| wMKL [56] | GitHub | R | ✓ |
| WSNF [57] | GitHub | R | ✗ |

*Some methods may be available in multiple programming languages.
We report the one used in this work.
**Baseline analysis only. As included in *MOVICS*.
***Package developed years after the original papers.
+"R+Python" means the method uses both Python and R functionality.

- **MixKernel:** Consistent errors about kernel positivity and semi-definite matrices. Required modifications, but still did not work.
- **PAMOGK:** Pathway-based MultiOmic Graph Kernel (PAMOGK) is not applicable to miRNA data, because they are not translated into proteins. Could still work with the remaining four modalities, but would require modifications for handling binary data.

- **PARADIGM:** Pathway Recognition Algorithm using Data Integration on Genomic Models (PARADIGM) is no longer directly available, but can be provided for researchers if requested. All code has been removed from GitHub.
- **RGCCA/SGCCA:** Regularised Generalised Canonical Correlation Analysis (RGCCA) and Sparse Generalised Canonical Correlation Analysis (SGCCA) cannot work with binary data (see forum too).
- **SIMLR:** Single-cell Interpretation via Multi-kernel LeaRning (SIMLR) was originally developed for single-cell data. We are using Cancer Integration via Multikernel LeaRning (CIMLR) and weight-boosted Multi-Kernel Learning (wMKL) here which introduced improvements to the original algorithm.
- **WSNF**: Weighted Similarity Network Fusion (WSNF) requires transcription factor data to run and is not applicable in our case.

### 3 Preprocessing

#### 3.1 TCGA training cohort

The information provided in these sections refers to the analysis carried out by running the `Scripts/Download_TCGA_data.R` script for the preprocessing of our training cohort, which is a subset of the The Cancer Genome Atlas (TCGA) breast cancer project.

##### 3.1.1 Querying TCGA for omics datasets

We downloaded data from TCGA using the *TCGAbiolinks* package [58–60] in R [1]. We queried the database for open access primary tumour data in the TCGA-BRCA project. Five different types of omics datasets were downloaded from TCGA, plus clinical data and each omic dataset was then preprocessed using the built-in function `GDCprepare()` with default options:

1. **Gene expression data.** RNA sequencing (RNA-seq) data were downloaded using `GDCquery()` and setting the following options:
   - `data.category = "Transcriptome profiling"`
   - `data.type = "Gene Expression Quantification"`
   - `workflow.type = "STAR - Counts"`
2. **microRNA expression data.** microRNA (miRNA) data were downloaded using `GDCquery()` and setting the following options:
   - `data.category = "Transcriptome profiling"`
   - `data.type = "miRNA Expression Quantification"`
   - `experimental.strategy = "miRNA-Seq"`
3. **Copy Number Variation.** Copy Number Variant (CNV) data were downloaded using `GDCquery()` and setting the following options:
   - `data.category = "Copy Number Variation"`
   - `data.type = "Gene Level Copy Number"`

When this analysis was performed, primary tumour gene level CNV data were profiled using the ABSOLUTE LiftOver pipeline [61].

4. **Methylation.** Methylation data were downloaded using `GDCquery()` and setting the following options:
   - `data.category = "DNA methylation"`
   - `data.type = "Methylation Beta Value"`
   - `platform = "Illumina Human Methylation 450"`

   Although $\beta$-values were downloaded from TCGA, $M$-values were actually used in the analysis; see section 3.1.2.4.
5. **Single Nucleotide Polymorphisms.** Single Nucleotide Polymorphism (SNP) data were downloaded using `GDCquery()` and setting the following options:
   - `data.category = "Simple Nucleotide Variation"`
   - `data.type = "Masked Somatic Mutation"`
   - `workflow.type = "Aliquot Ensemble Somatic Variant Merging and Masking"`

   The generation of gene level SNP data is described in section 3.1.2.5 in detail.
6. **Clinical information.** Clinical data were downloaded using `GDCquery()` and setting the following options:
   - `data.category = "Clinical"`
   - `data.type = "Clinical supplement"`

##### 3.1.2 Preprocessing of omics datasets

Each of the following subsections describes the step-by-step approach we took to preprocess each omics dataset to make it suitable for multi-omics analysis using the selected algorithms. After the final list of datasets was derived, this was consistently used for multi-omics workflows and downstream analysis. All algorithms and methodologies that are compared in this paper, used this list of datasets as input.

Regardless of omics dataset, the first step of the preprocessing was to identify the male samples that were downloaded and remove them from the analysis. Additionally, the columns of each data matrix (samples) were annotated with sample identifiers of the format `TCGA-##-####-###`.

###### 3.1.2.1 RNA-seq

From the RNA-seq data object which was downloaded, we kept the unstranded counts. The row identifiers (feature names) were set to Human Genome Organization (HUGO) symbols. Rows that only contained zero values were removed.

Several ways of normalizing RNA-seq counts have been proposed over the years, including Reads Per Kilobase Million (RPKM) (originally used for single-end RNA-seq), Fragments Per Kilobase Million (FPKM) (used for paired-end RNA-seq), Transcripts Per Million (TPM) and more. RPKM and FPKM are produced for each sample after considering the total number of counts in each sample and each gene's length in kilobases. TPM are produced similarly, but the series of steps is reversed: the counts are first divided by the length of each transcript in kilobases and then scaling for each sample based on its total number of counts follows. This ensures that the sum of all TPM in each sample are the same. Therefore, counts are transformed into a proportion of reads within a sample.

RPKM and FPKM cannot be used by definition for comparisons across samples. TPM should be used with caution for such purposes, because the scaling factor differs for each sample and therefore, equal proportions of counts do not guarantee equal counts across samples. We, thus, used a different form of count normalisation, implemented in the *DESeq2* package [62] in R, namely the "median of rations" method [63], which has been shown to provide more robust normalisation and yield output that is more suited for downstream tasks like clustering [64–66].

Eventually we want to have all continuous modalities on similar scales. Therefore, the final step in the preprocessing of each omics dataset is standardisation of features across samples. However, there exist extreme outliers in the normalised counts as shown in Figures 1a and 1b (x-axis truncated at 30, on purpose). To avoid the strong effect of these outliers in the standardisation step, we further transform the data by logarithmic transformation and the addition of a pseudocount. The choice pseudocount was performed according to the procedure described by Lun et al. [67]. Through this process, we choose a pseudocount of 1.

After transforming each count entry $K_{ij}$ by $\log_2(K_{ij}+1)$ transformation, we proceed by identifying the rows that are annotated with the same HUGO symbol. Whenever two or more rows were annotated with the same HUGO symbol, the row with the highest variance was kept. In the case of ties, the tied rows were averaged into one row.

As a final filtering step before standardisation, rows with missing values or rows containing only zero values are removed. The resulting expression matrix of logarithmically transformed normalised counts was subjected to standardisation (each row, across all samples). The final matrix of dimensions $p \times 625$ was used as input for downstream analysis.

###### 3.1.2.2 miRNA

From the miRNA-seq data object which was downloaded, we kept the read counts as our data matrix. The row identifiers (feature names) were set to miRNA `hsa-miR-###`. Rows that only contained zero values were removed. Normalised miRNA-seq counts were obtained similarly to the RNA-seq case, using the median of ratios method and the choice of pseudocount for the logarithmic transformation was made after following the procedure described by Lun et al. [67]. The resulting matrix was then subjected to standardisation of rows across samples.

###### 3.1.2.3 CNV

From the CNV data object which was downloaded, we kept the copy number counts as our data matrix. The row identifiers (feature names) were set to HUGO symbols. Rows with missing values were then removed. We then proceed by identifying the rows that are annotated with the same HUGO symbol. Whenever two or more rows

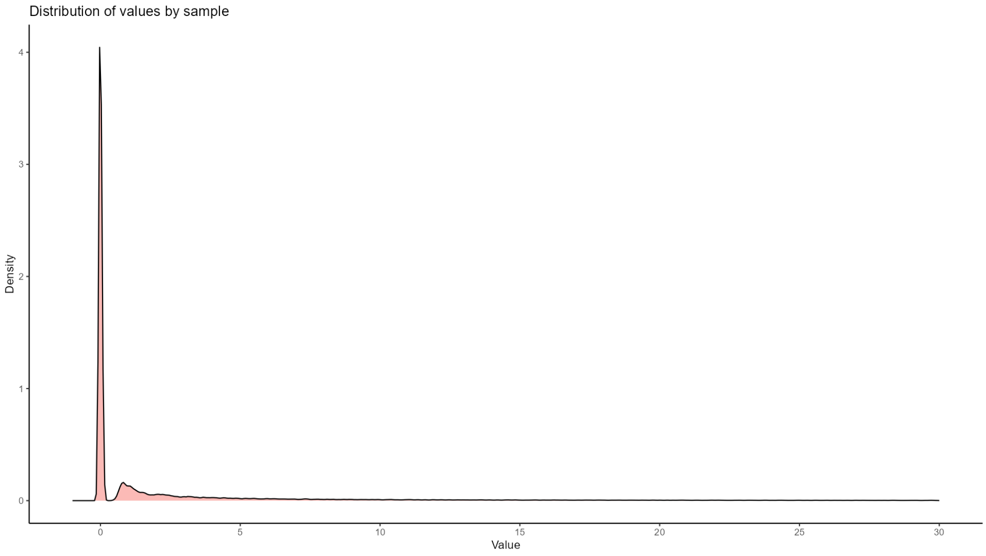

**(a)**

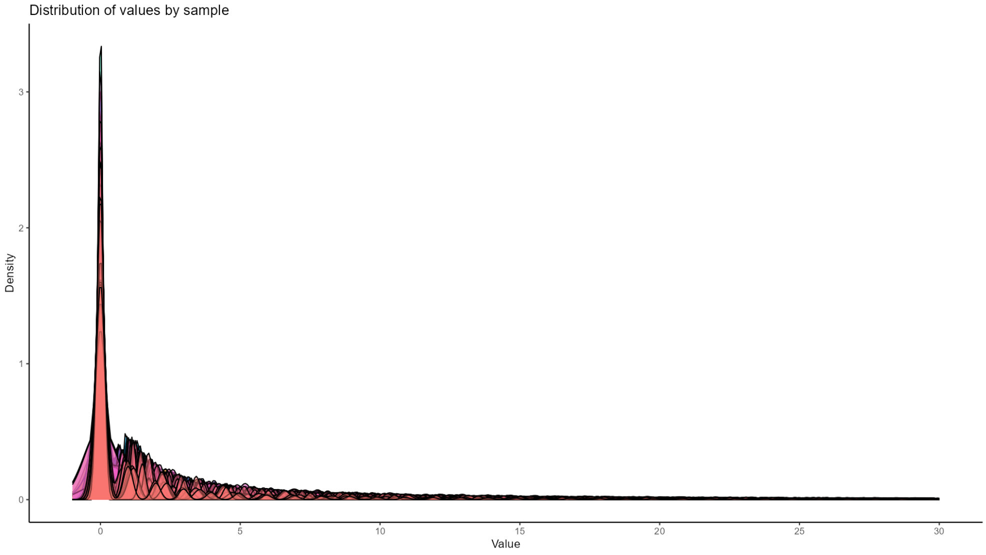

**(b)**

**Figure 1. Normalised counts distribution in RNA-seq. (a) Distribution of normalised counts in RNA-seq prior to transformation.** The majority of normalised counts lie in the rough area between 0 and 5. The x-axis is truncated at 30 for visualisation purposes. **(b) Density curves of normalised counts in RNA-seq prior to transformation, per sample.** The x-axis is truncated at 30 for visualisation purposes.

were annotated with the same HUGO symbol, the row with the highest variance was kept. In the case of ties, the rows are summarised in one row using the **median** of each column. After removing any remaining rows with missing values and/or only zero values, the rows were standardised across samples.

###### 3.1.2.4 Methylation

From the methylation data object which was downloaded, we kept the $\beta$-values as our data matrix. The row identifiers (feature names) were set to HUGO symbols. Rows with missing values in more than 25% of the samples were then removed. The rest of the values were imputed using the *impute* package in R, which employs $k$-Nearest-neighbour ($k$-NN) imputation [68, 69]. We set $k = 10$.

The package offers a few additional options. We set `rowmax = 0.25` to keep rows with less than 25% missing values (already ensured in a previous step), `colmax = 0.8` to keep samples which have methylation levels recorded for at least 80% of the rows and `maxp = 1500`. `maxp` is the largest block of genes imputed using the $k$-NN algorithm inside `impute.knn()` (default 1500); larger blocks are divided by two-means clustering (recursively) prior to imputation. If `maxp` is set equal to the number of features, only $k$-NN imputation is done. Due to the considerable size of the dataset, we left `maxp` equal to the default value, 1500.

TCGA is using the SeSAMe software package [70] for its methylation pipeline. We used functionality from the corresponding R package to convert the $\beta$-values to $M$-values, which have more appropriate statistical properties (especially with respect to heteroscedasticity) and are more suitable as input for downstream tasks [71]. An $M$-value around 0 means that the methylated and unmethylated signals are close to equal. A positive $M$-value means that the methylated signal is higher, while a negative M-value means that the unmethylated signal is higher.

We then proceed by identifying the rows that are annotated with the same HUGO symbol. Whenever two or more rows were annotated with the same HUGO symbol, the row with the highest variance was kept. In the case of ties, the rows are summarised in one row using the average of each column. After removing any remaining rows with missing values and/or only zero values, the rows were standardised across samples.

###### 3.1.2.5 SNPs

For SNPs, the output of the `GDCprepare()` function was used as input in the `read.maf()` function from package *maftools* [72], which creates a summary of the mutation data. We filter the summarised dar for for `Variant_Type == "SNP"` which results in 62,325 variants, distributed as: 56,025 missense, 4901 nonsense, 63 translation start site, 1262 splice and 74 nonstop mutations. By filtering for `Variant_Type == "SNP"`, we explicitly exclude frameshift and in-frame insertions/deletions (which are classified as `Variant_Type == "INDEL"`). We then use HUGO symbols for row annotation. Finally, we convert the data frame into a binary matrix which has gene symbols in rows and sample identifiers in columns. Each entry $(i, j)$ in the matrix is set equal to 1 if there is at least one SNP on gene $i$ in sample $j$, or 0 otherwise. All rows which contained only zero values were removed.

###### 3.1.2.6 List of omic inputs

The five resulting matrices, with dimensions as shown in the table below, were put in a `list` object, with names as shown in the first column. The object was exported and used as input for multi-omics analyses and downstream analyses.

**Table 2.** Overview of multi-omic input

| Cohort | Omics | Dimensions | Type |
|---|---|---|---|
| **TCGA-BRCA** | SNPs | 12926 × 625 | Binary |
| | RNA-seq | 56717 × 625 | Continuous |
| | miRNA | 1568 × 625 | Continuous |
| | Methylation | 36761 × 625 | Continuous |
| | CNV | 35858 × 625 | Continuous |
| **transNEO** | RNA-seq | 18719 × 153 | Continuous |

##### 3.1.3 Clinical information

Clinical data were downloaded and prepared as described in section 3.1.1. We used the patient barcode column to filter for the samples which have measurements in all five modalities. For a small number of patient identifiers, there existed multiple entries with clinical information. In each case, we kept the entry with the fewest missing values across all clinical variables. The final object is available at `Resources/TCGA/clinical_data.xlsx` in the GitHub repository. All clinical data can be also found in Supplementary File 3.

#### 3.2 Evaluation cohorts

The evaluation cohorts used throughout this work are the transNEO cohort [73] and the remaining samples from TCGA-BRCA. The preprocessing outputs for this study are available in Supplementary File 4 (available in Zenodo). The TCGA evaluation cohort's expression data (only modality used for label propagation) have been preprocessed identically to the expression data from the training cohort, but separately. The corresponding code for the holdout TCGA cohort can be found in the `Scripts/Survival analysis/Survival_TCGA_holdout.R`.

Regarding transNEO, the preprocessing scripts follow the original transNEO preprocessing steps with minor or no modifications and are designed for use within the official transNEO repository (https://github.com/cclab-brca/neoadjuvant-therapy-response-predictor).For transNEO, the order of the scripts is as follows:

1. `File_prep.R` for initial preparation of data.
2. `Preprocessing.R` for cleaning and extracting objects of interest.

3. `transNEO_pseudocount_determination.R` for determining the optimal pseudocount for transNEO normalised counts. This script can be used after delivering the resources of interest produced through the first two scripts, in the `Resources/transNEO` directory.

We produced the files we needed for our analysis using input from the transNEO repository and then included them in our `Resources` folder.

For both TCGA holdout and transNEO evaluations, we used the gene expression matrix and clinical data as input for the Nearest Template Prediction (NTP) and Partioning Around Medoids (PAM) methods described below [74, 75]. The expression data were $\log 2$-transformed with the addition of a data-driven pseudocount as in the training data. The grouping variable that was used to determine the optimal pseudocount (1) here was the binary pathological Complete Response (pCR) variable in transNEO, and the estrogen receptor (ER) status in the TCGA holdout. The matrices were then centered and scaled.

### 4 General pipeline

#### 4.1 Overview

In this work, we ran a baseline consensus multi-omic clustering approach (Section 7) using the *MOVICS* package [2] with default settings and subsequently, we examined a series of different algorithms separately for the task of multi-omic clustering. Finally, we use an updated consensus multi-omic clustering approach with selected algorithms.

Throughout this work, the overall pipeline is adapted for each algorithm, however, its higher order structure remains the same to ensure robust and meaningful comparisons across methods. The following sections describe the default workflow used for each method, including the baseline and final analysis. Whenever parameters of this workflow are modified, this is explicitly mentioned within the corresponding subsections in Sections 7 - 9. For most of the following steps we use functionality from the *MOVICS* package.

#### 4.2 Clustering output inspection

After clustering is completed, we provide multi-omic heatmap plots and heatmaps of final similarities (if available), annotated with clinical information in the form of colorful rags over the heatmap. Additionally, we provide a silhouette plot that illustrates the robustness of the produced clustering, by estimating average within-cluster distances and between-cluster distances. Silhouette values range from $-1$ to 1. If a sample has a value close to 1, then that the sample is "well-clustered", and its cluster is very different from neighbouring clusters. Conversely, if a sample has a value close to $-1$, the sample is probably "misclassified", as it is closer to a neighbouring cluster than to its own cluster. Values close to 0 indicate that a sample lies close to the boundary between two clusters. The silhouette values of all samples are then averaged to give a metric for the full clustering output.

#### 4.3 Subtypes and clinical variables

*MOVICS* provides wrapper functions that assess a set of variables selected by the user for association with the identified subtypes. By default, chi-square tests are carried out to test associations of discrete variables with the identified subtypes and either parametric $t$-tests and ANalysis Of VAriance (ANOVA) tests for continuous variables or non-parametric Wilcoxon rank sum tests and Kruskal-Wallis tests, for the continuous variables the users specifies to be non-normal. In all cases, we used non-parametric tests for continuous variables.

Bar charts for the discrete variables which are found to be significantly associated with subtypes are generated and annotated with corrected Chi-square statistics, the corresponding $p$-value and Cramér's $V$; the latter is used to determine effect size of associations, using the *rcompanion* package [76].

For selected variables which exhibit significant associations with the subtypes, a sunburst plot is also produced in HTML format and is available within the `Supplement` subdirectory of each method's `Results` directory.

#### 4.4 Mutations

*MOVICS* provides a wrapper function (`compMut()`) which compares mutation frequencies across subtypes.

1. **Frequency Filtering**: - Genes that are mutated in fewer than 5% of samples are filtered out. This threshold is defined by the user through the `freq.cutoff` parameter. We leave this to the default `0.05`.
2. **Statistical Testing**: - The association between each gene's mutation status and the subtype is tested using either Fisher's exact test (default) or the chi-square test. We use Fisher's exact test, unless the number of subtypes is large and therefore leads to increased computational intensity. - $p$-values are adjusted for multiple testing using methods such as the Benjamini-Hochberg procedure (we keep it as such), which controls the False Diacovery Rate (FDR).
3. **Oncoprint Plot**: - A visual representation (oncoprint) of mutation events across clusters is generated for genes with significant associations with the produced clusters ($p_{adj} < 0.05$).

#### 4.5 Differential analysis

*MOVICS* allows the user to carry out Differential Gene Expression Analysis (DGEA) using one of three methods, coming from three different packages: *DESeq2* [77] and *edgeR* [78] for RNA-seq count data and *limma* [79] for microarray data. Due to the preprocessing (normalisation and standardisation) of the RNA-seq data, we selected the method from *limma*, which works with linear models of expression [79].

After the Differentially Expressed Genes (DEGs) are identified, we choose the top 100 non-overlapping up-regulated or down-regulated DEGs (arranged by magnitude of differential expression value, after filtering for the $p$-value and $p_{adj}$ thresholds) for each subtype and produce the corresponding heatmaps.

We use the same approach to create heatmaps of the top 100 non-overlapping markers in miRNA and methylation data.

The inputs that we use when performing differential analysis in all of these modalities is the standardised data for each modality, with large values ($< -3$ and $> 3$) truncated to $-3$ and 3. The truncation helps in limiting the effect of outliers and producing heatmaps with smoother color gradients.

#### 4.6 Fraction of genome altered

In the case of CNVs, we generate a plot of the fraction of genome that is altered (Fraction of Genome Altered (FGA)) in the samples of each identified subtype. This plot uses **original counts of copy number segments** as input and a custom definition of gain and loss. In our work, we used two approaches in annotating segments in a sample with one of three possible labels: "gain", "loss" and "normal".

In the first, simpler approach, we define gain as a copy number $> 2$, loss as a copy number $< 2$ and normal as copy number equal to 2. In the second approach, we use Catalogue Of Somatic Mutations In Cancer (COSMIC) criteria to define gain and loss, which also take into account the average ploidy of each sample. More specifically, we define five cases:

$$\text{Status} = \begin{cases} \text{gain} & \text{if ploidy} \leq 2.7 \text{ and copy number} \geq 5, \\ \text{gain} & \text{if ploidy} > 2.7 \text{ and copy number} \geq 9, \\ \text{loss} & \text{if ploidy} \leq 2.7 \text{ and copy number} = 0, \\ \text{loss} & \text{if ploidy} > 2.7 \text{ and copy number} < \text{ploidy} - 2.7, \\ \text{normal} & \text{otherwise.} \end{cases}$$

We modified the *MOVICS* `compFGA()` function which compares subtypes based on the FGA, Fraction of Genome Gained (FGG), and Fraction of Genome Lost (FGL). The function performs statistical tests to compare the subtypes using either parametric or non-parametric methods, depending on the number of subtypes and the chosen test method. We modified the function to allow for predefined gain/loss/normal annotations. The code is included in the script `Scripts/automated_scripts/modified_MOVICS_functions.R`.

Mathematically, FGA, FGG, and FGL are defined as described in Equations 1, 2 and 3.

$$\text{FGA} = \frac{\sum(\text{bases where classification is "gain" or "loss"})}{\sum(\text{total bases})} \tag{1}$$

$$\text{FGG} = \frac{\sum(\text{bases where classification is "gain"})}{\sum(\text{total bases})} \tag{2}$$

$$\text{FGL} = \frac{\sum(\text{bases where classification is "loss"})}{\sum(\text{total bases})} \tag{3}$$

For every method (including baseline, individual trials and the final consensus pipeline), we used original copy number counts and non-parametric tests:

- **Wilcoxon Rank-Sum Test (Non-parametric, 2 subtypes)**: If there are two subtypes, the Wilcoxon rank-sum test is used to compare the values of FGA, FGG, and FGL between the two subtypes.
- **Kruskal-Wallis Test (Non-parametric, more than 2 subtypes)**: When there are more than two subtypes, the Kruskal-Wallis test (`kruskal.test`) is applied. This is a non-parametric version of ANOVA used to detect differences between multiple subtypes.
  Additionally, pairwise comparisons between subtypes are performed using the Wilcoxon test with the Benjamini-Hochberg $p$-value adjustment.

In all cases, the function summarizes the FGA, FGG, and FGL values for each subtype and calculates the respective $p$-values to assess statistical significance.

#### 4.7 Pathway analysis

With the results from DGEA, the *MOVICS* pipeline proceeds with identifying up- and down-regulated pathways in the subtypes, using Gene Set Enrichment Analysis (GSEA) [80, 81]. When running GSEA, we specify the minimum size of considered gene sets to `5` and maximum size to `500`. Additionally, we define significant pathways by requiring that: $p < 0.05$ and $p_{adj} < 0.05$ and set the number of permutations to `nPerm = 10000`. The background gene sets used for enrichment analysis were downloaded from https://www.gsea-msigdb.org/gsea/msigdb/human/collections.jsp (GO-BP C5, Immunogenic C7 and Oncogenic C6).

After getting the subtype-specific pathways, genes that are members of these pathways are retrieved to calculate a single sample enrichment score by using the *GSVA* R package [82]. Subsequently, subtype-specific enrichment scores are represented by the mean value within the subtype and further visualised by heatmap. Typically, we choose 20 pathways per subtype, ranked by enrichment score, but for a larger number of clusters we use 10 pathways per subtype.

We use functionality from the *pathfindR* package in R [83] to perform hierarchical clustering on the identified pathways based on the common contributing genes to get a clear higher level overview of the deregulated biological processes in each subtype. We modify the original hierarchical clustering implementation of pathfindR to use the *fastcluster* package's implementation of `hclust` [84] to perform hierarchical clustering more efficiently.

*MOVICS* also offers a Gene Set Variation Analysis (GSVA) approach through which specific pathways of interest can be examined across individual samples and subtypes, visually. For this function, we use a set of 25 manually selected immune processes to inspect the data for potential differential immune activation across clusters. The input that we use when performing GSVA is the standardised RNA-seq data, with large values ($< -3$ and $> 3$) truncated to $-3$ and 3.

#### 4.8 Evaluation on external cohort

*MOVICS* offers two methods to evaluate results from the training data (expression data only; although adaptable to use another modality instead) on an external cohort: a) NTP [74] and b) PAM [75, 85].

##### 4.8.1 Nearest Template Prediction (NTP)

Cosine similarity is used to compare external samples to class templates. Class templates are generated using the up-regulated (or down-regulated) genes in the form of a binary matrix with columns: "cluster #", where an entry is 1 if the gene is up-regulated in the cluster # or -1 otherwise. The nearest class for each sample is determined by the smallest distance (converted from cosine similarity here). Each sample's result includes the nearest class prediction, the distance to all templates, and the $p$-value of the prediction. The number of markers used for template creation is consistently set to `1000` for all methods unless specified otherwise.

The function randomly permutes the sample labels multiple times (here `nPerm = 10000`) to generate a null distribution of similarity scores between samples and class templates. For each observed sample-to-template distance, the function compares it against the null distribution to calculate a $p$-value. This $p$-value indicates the likelihood that the observed match could have occurred by chance. The $p$-values are adjusted using the FDR method to control for multiple hypothesis testing.

##### 4.8.2 Partition Around Medoids (PAM)

In this approach, *MOVICS* first trains a PAM classifier in the discovery (training) cohort to predict the subtype for patients in the external validation (testing) cohort, and each sample in the validation cohort was assigned to the subtype label whose centroid had the highest Pearson correlation with the sample. Finally, the In-Group Proportion (IGP) statistic (the proportion of samples within a cluster that were classified into the same cluster upon random permutation of the data) is estimated to evaluate the similarity and reproducibility of the acquired subtypes between discovery and validation cohorts (i.e. are the clusters identified in the discovery cohort reproducible in the validation cohort after label permutation?).

#### 4.9 Dimensionality reduction plots

After generating the subtypes of interest using a particular method, we create dimensionality reduction plots to investigate - in an exploratory way - how the identified subtypes are distributed in each modality's lower dimensional space. When the method in question is using similarity matrices to perform clustering (e.g. Similarity Network Fusion (SNF) [54, 86], NEighborhood-based Multi-Omics clustering (NEMO) [43] etc.), we use a modified function from the *M3C* package [87], `kernelPCA()` to produce low-dimensional projections of the original similarity matrices and the final similarity matrix (if applicable). Otherwise, we produce traditional Principal Component Analysis (PCA) plots using the original modality matrices.

In the case of binary mutations (our SNPs modality; see more in Section 3.1.2.5), we use Multi-dimensional Scaling (MDS) with binary distance. This approach is more robust and appropriate for binary data, since PCA assumes normally distributed features in the high-dimensional space, which is not an appropriate assumption for binary data.

#### 4.10 Drug sensitivity tests

The *MOVICS* function `compDrugsen()` performs a drug sensitivity analysis by estimating IC50 (half-maximal inhibitory concentration) values based on normalised gene expression data. The predicted IC50 values are compared across subtypes identified by the examined method, and different statistical tests are applied depending on the number of subtypes and the chosen method (non-parametric or parametric).

For each drug, ridge regression using the *pRRophetic* package [88] is used to predict the IC50 values based on the transformed gene expression data. Ridge (L2-regularised) regression models are built using the drug sensitivity data from cell lines in the Cancer Genome Project dataset [89] for the tissue type specified by the user (`tissueType = "breast"` in our case).

We choose non-parametric tests for our data. In the case of 2 subtypes, the Wilcoxon rank-sum test is applied to compare the IC50 values between the two subtypes. In the case of more than 2 subtypes, the Kruskal-Wallis test is applied to test for differences in IC50 distributions across subtypes. Additionally, a pairwise Wilcoxon test is conducted with Benjamini-Hochberg adjustment for multiple comparisons.

#### 4.11 Subtype agreement

*MOVICS* provides functionality that measures the agreement between the produced subtypes and known/established subtypes. More specifically, four metrics of agreement are calculated: the Rand Index (RI), the Adjusted Mutual Information (AMI), the Jaccard index ($JI$) and the Fowlkes-Mallows (FM) index. A bar chart with the corresponding metrics for each known/established subtype is produced along with an accompanying alluvial plot.

#### 4.12 Graphs from similarity matrices

For methods which construct similarity matrices for individual modalities and/or a final similarity matrix, we use the *igraph* package in R [90, 91] to construct graphs of these relationships.

If during the similarity matrix generation a specific number of neighbours for each node is used for thresholding and filtering for stronger relationships between nodes, we use that number to limit the number of edges that are drawn on the graph. We weigh edges by the magnitude of the recorded similarity value for each pair of nodes (weight). More specifically, the width of the edges is set based on the square root of their weights, scaled by a factor of 10. This transformation is applied to improve visibility, making stronger connections more prominent in the plot. Edges with weight equal to 0 are not drawn, regardless of whether a specific number of neighbours is used or not. From the rest of the edges only those on the 4th weight quartile are drawn (apart from wMKL, where we draw edges from the 3rd and 4th quartiles).

In our approach, we use the Fruchterman-Reingold layout algorithm [92] to position the nodes. This layout spreads nodes in a way that reduces edge overlap and maximizes visual clarity.

In the case of SNF and other methods which construct graphs and are amenable to this, we also draw a version of the final graph with coloured edges, depending on the modality (or combination of modalities) that mainly affect(s) the respective sample-to-sample relationship. For that, we use the approach described in the supplement of the SNF original publication [54], which is as follows: The edge is considered supported by a single data type if the weight (patient similarity) in that data type's network is more than 10% higher than the similarity of the same edge in the other data types' networks. If the difference between two highest edge weights from the corresponding data types is less than 10%, the edge is considered to be supported by those 2 data types and if the difference between all of the edge weights between all data types is less than 10%, it is considered to be supported by all data types.

#### 4.13 Interpretability

Some methods contain built-in ways to rank features and/or omics, other than the ones we can use post-hoc for all results (e.g. DGEA to select subtype-specific markers). When feature ranking results are available, these are presented in the corresponding section.

For Similarity network methods, we extracted feature rankings (top 1000 features) and weighted fused graphs using the approaches described in the original SNF publication [54] (see also Section 4.12). The feature ranking in SNF uses a Normalised Mutual Information (NMI)-based approach: for each feature an affinity graph is build only on that feature and spectrally clustered. Then, that single-feature clustering is compared back to the "full" (fused) clustering in terms of NMI. The features with the highest NMI metrics are ranked the highest.

For Multi-omics Data Integration for Clustering to identify Cancer subtypes (MDICC), the source code of the main function needed to be slightly modified to allow for the return of omic-level kernel weights which quantify the contribution of each modality to the final results. A similar kernel summing approach was followed to estimate omic-level contributions in the CIMLR case. Kernel Learning Integrative Clustering (KLIC) generates a $n \times p$ weight matrix, where $n$ is the number of samples and $p$ the number of modalities. We rank modalities in terms of importance by taking the average weight per modality. For Multi Omic clustering by Non-Exhaustive Types (MONET), we inspect the excess within-module connectivity (the degree to which samples in a module are more tightly linked than the cohort-wide average for each omic), to investigate whether modules are dominated by subsets of modalities.

In Multi-Omics Factor Analysis (MOFA), we examine feature weights in the loading vectors of each factor to quantify feature importance per factor and, at the omic level, examine plots of explained variance per factor. Additionally, we look at the distribution of three key clinical variables in the different factors: ER status, Human Epidermal growth factor Receptor 2 (HER2) status and tumour stage. We follow similar steps in MFA to estimate feature- and omic-level contribution in the obtained factors, as well as assess the factors' relationships with clinical variables.

### 5 Closing remarks

The previous section describes the generic steps we take for the analysis of each method that was examined in this work, including the baseline analysis, the individual algorithm trials and the final consensus pipeline. Deviations from the specifics of the processes we discussed can occur, and these are explicitly mentioned in the corresponding sections. Most of the time that a process is modified, it is due to computational intensity problems.

The baseline analysis is conducted using the *MOVICS* package and the 10 included algorithms (see Section 7) using default settings. For each method included in *MOVICS* and the individual methods we examine later, we provide a general overview of the underlying algorithm, as well as how it is implemented in *MOVICS*, if applicable. When we run methods individually, we are trying to follow a robust pipeline for hyperparameter selection and the handling of both continuous and binary types of input data. Whenever we introduce modifications in specific methods (e.g. for the use of binary distances for binary data), this is clearly mentioned in the corresponding section.

### 6 Algorithms

This section explains the methodology behind each algorithm and the corresponding implementation in our analysis. The results for the individual runs of each method are presented in Section 8.

The following methods were not used in the individual method trials, either because they cannot directly handle binary data or because they are uni-modal: Integrative Non-negative Matrix Factorization (IntNMF) [28], moCluster [39], Perturbation clustering for data INtegration and disease Subtyping (PINSPlus) [46–48] and Consensus Clustering [23]. However, they are still described in this section because they are included in the *MOVICS* suite of tools [2] which we use for our baseline analysis.

#### 6.1 Similarity Network methods

Similarity network methods first translate each data view into a sample-by-sample similarity graph, then combine or refine these graphs to produce an integrated, fused affinity matrix whose structure is subsequently used for clustering or other downstream analyses.

**Note:** For all similarity network methods, we pool the pairwise-similarity scores for all affinity matrices that share the same number of neighbours $NN$ and submit them to a one-way ANOVA (with a rank-based Kruskal–Wallis back-up) to test the null hypothesis that varying the regularisation hyperparameter $\sigma$ leaves the distribution unchanged; repeating the procedure with the roles of $NN$ and $\sigma$ swapped indicates whether altering $NN$ matters when $\sigma$ is held constant. A significant $p$-value for a given factor indicates that its different settings generate statistically distinct similarity structures, while the relative spread of the group means (or medians) allows us to judge which factor exerts the stronger practical influence. When $p > 0.05$, we use $\sigma = 0.5$

and $NN = 30$, respectively. Otherwise, we use all similarity matrices with significant differences for spectral clustering and select the optimal clustering based on the silhouette index. In case of contradicting results between ANOVA and Kruskal-Wallis, the Kruskal-Wallis results are used.

##### 6.1.1 Similarity Network Fusion (SNF)

###### Overview

SNF first builds a similarity graph for each data modality by defining edge weights

$$W_{ij} = \exp\Big(-\tfrac{\rho^2(x_i,x_j)}{\mu\,\varepsilon_{ij}}\Big),$$

where $\rho^2$ is the squared Euclidean distance between samples $i$ and $j$, $\mu$ is a scaling hyperparameter (typically in $[0.3, 0.8]$; referred to as `sigma` in code and $\sigma$ in the paper), and $\varepsilon_{ij}$ adapts to local neighbourhood density. Each matrix $W$ is then normalised into a full affinity matrix $P$ and a sparse matrix $S$:

$$P_{ij} = \begin{cases} \dfrac{W_{ij}}{2\sum_{k\neq i} W_{ik}}, & j \neq i, \\ 0.5, & j = i, \end{cases} \qquad S_{ij} = \begin{cases} \dfrac{W_{ij}}{\sum_{k\in N_i} W_{ik}}, & j \in N_i, \\ 0, & \text{otherwise.} \end{cases}$$

An iterative fusion step then exchanges information across modalities:

$$P^{(\nu)} \;\leftarrow\; S^{(\nu)}\Big(\tfrac{1}{m-1}\sum_{u\neq\nu} P^{(u)}\Big)S^{(\nu)T},$$

repeating for $T$ rounds until a consensus network emerges. The final fused affinity matrix is used for spectral clustering.

###### Implementation

We used the *SNFtool* package [86] in R, to perform SNF on the multi-omic data. Before providing the `affinityMatrix()` function of the *SNFtool* package with the distance matrices for each separate modality, we calculated Euclidean distances for the continuous modalities (normalised RNA-seq, normalised CNVs, normalised miRNAs and normalised Methylation) and binary distances for SNPs calculated with `dist(..., method="binary")`.

There are three main tunable hyperparameters in SNF: the number of neighbours ($NN$), the regularisation parameter $\sigma$, and the number of iterations $T$. Here, we used a range of number of neighbours and $\sigma$ values:

$$NN \in \{10, 15, 20, 25, 30, 35, 40, 45, 50\}$$

$$\sigma \in \{0.3, 0.4, 0.5, 0.6, 0.7, 0.8\}$$

and a number of iterations $t = 50$. Results for larger $T$ are usually identical because the algorithm usually converges after roughly 20 iterations [54].

We compared the final fused affinity matrices for a given $NN$ and/or a given $sigma$ (Pearson and Frobenius similarities between matrices). Overall:

- the choice of $\sigma$ does not significantly affect the results for a given $NN$
- the choice of $NN$ significantly affects the results for a given $\sigma$

We therefore set $sigma = 0.5$, which is close to the median $sigma$ value (0.55, but rounded down to 0.5 which was actually included in our range of tested $sigma$ values). The fused matrices for $sigma = 0.5$ and $NN \in \{10, 15, 20, 25, 30, 35, 40, 45, 50\}$, are then used as input in the `estimateNumberOf ClustersGivenGraph()` function of *SNFtool* to determine the optimal number of clusters in the data based on two methods: i) the eigengap method and ii) the discretisation (rotation) method. Table 3 summarizes the results of this process and illustrates how the optimal number of clusters is 2.

| Fusion | K1 | K2 | Eigengap score | Rotation score |
|---|---|---|---|---|
| $NN = 10$ | 2 | 2 | 0.312 | 432.500 |
| $NN = 15$ | 2 | 2 | 0.335 | 414.996 |
| $NN = 20$ | 2 | 2 | 0.358 | 398.877 |
| $NN = 25$ | 2 | 2 | 0.374 | 386.790 |
| $NN = 30$ | 2 | 2 | 0.376 | 380.093 |
| $NN = 35$ | 2 | 2 | 0.375 | 374.583 |
| $NN = 40$ | 2 | 2 | 0.374 | 369.923 |
| $NN = 45$ | 2 | 2 | 0.369 | 366.592 |
| $NN = 50$ | 2 | 2 | 0.365 | 363.174 |

**Table 3. Optimal number of clusters for varying values of nearest neighbours. K1**: eigengap method. **K2**: rotation method. $\sigma = 0.5$

Therefore, we run spectral clustering with $k = 2$ for the fused affinity matrices ($\sigma = 0.5$) and select the $NN = 25, \sigma = 0.5$ - fusion as optimal, because it achieves the best silhouette score (0.792).

##### 6.1.2 annotation signal boosted Similarity Network Fusion (ab-SNF)

###### Overview

association-signal-annotation boosted Similarity Network Fusion (ab-SNF) [15] extends SNF by incorporating feature-level weights derived from external association signals. First, for each omics modality, a weight $w_k$ for feature $k$ is computed, for example, as

$$w_k = \frac{-\log_{10}(p_k)}{\sum_{k'} -\log_{10}(p_{k'})},$$

where $p_k$ is the association $p$-value of feature $k$ with an outcome of interest (e.g. survival or case–control status). This up-weights features with strong signals and down-weights noisy ones.

Next, a boosted similarity network is built for each modality by replacing the unweighted distance in SNF with a weight-adjusted metric. The weighted distance between samples $i$ and $j$ is computed (e.g. weighted Euclidean or Hamming distance using $w_k$) and then transformed via the same exponential kernel as in SNF. The resulting affinity matrix is normalised into a full network $P$ and a sparse local network $S$ following SNF's normalisation rules.

These boosted networks are fused across modalities by the standard SNF diffusion process, iterating for a fixed number of rounds $T$ until a consensus boosted network emerges.

Finally, the fused boosted affinity matrix is used for spectral clustering. By embedding association-signal weights into the similarity construction and leveraging SNF's diffusion, ab-SNF amplifies biologically relevant features and improves subtype discovery and downstream prediction.

###### Implementation

We weight each continuous feature in proportion to its median absolute deviation (weights normalised to sum to one) and give binary SNPs a weight of 0.8 if the locus maps to a COSMIC breast-cancer driver gene and 0.2 otherwise, using the resulting weights to compute weighted squared Euclidean (continuous) or weighted binary distance matrices. As in the case of SNF, the maximum number of iterations is set to $T = 50$ and the same grid of neighbours and $\sigma$ values are tested during tuning.

Identically to SNF, the effect of the number of neighbours for a fixed $\sigma$ is significant, while the effect of $\sigma$ for a fixed value of number of neighbours is not significant. We select $\sigma = 0.5$ and based on the `estimateNumberOfClustersGivenGraph()` function, we conclude $k_{\text{opt}}^{\text{ab-SNF}} = 2$. Spectral clustering on the fused matrices with $\sigma = 0.5$ revealed that the combination $\{NN = 10, \sigma = 0.5\}$ achieves the clustering with the highest silhouette index (0.789).

##### 6.1.3 Affinity Network Fusion (ANF)

###### Overview

Affinity Network Fusion (ANF) [16] streamlines similarity network fusion by replacing SNF's diffusion with one or two closed-form linear updates. For each omics layer, it first computes a $K$-NN affinity matrix $W_j$ using a Gaussian kernel whose adaptive bandwidth is:

$$\sigma_{ij} = \alpha(d_i + d_j) + \beta D_{ij} + \varepsilon$$

where $d_i$ is the average $K$-NN distance of sample $i$ and $D_{ij}$ is the direct distance between samples $i$ and $j$. The scalar $a$ (`alpha` in `ANF::affinity_matrix()`; default `1/6`) weights the *local* term $(d_i + d_j)$, so increasing $\alpha$ makes the kernel width depend more on each sample's immediate neighbourhood and hence emphasizes fine-grained structure. The scalar $\beta$ (`beta` in `ANF::affinity_matrix()`; default `1/6`) multiplies the *global* term $D_{ij}$; larger values smooth the affinities by giving more influence to absolute distances and less to local noise.

In "one-step" mode ANF simply averages the sparsified affinity matrices across omics. When ANF is run in its "two-step" mode, the fusion update for omic $j$ is a weighted sum of eight possible interaction paths between that omic's $K$-NN graph $S_j$, its raw similarity matrix $W_j$, and the cross-omic averages $S$ and $W$:

$$W_j^{\text{new}} = \alpha_1 S_j S + \alpha_2 S S_j + \alpha_3 S_j W + \alpha_4 W S_j + \alpha_5 W_j S + \alpha_6 S W_j + \alpha_7 W_j W + \alpha_8 W W_j$$

The length-8 vector `alpha` in `ANF()` contains these coefficients (default $(1, 1, 0, 0, 0, 0, 0, 0)$). Setting a coefficient high strengthens the corresponding information path, so users can tune how aggressively similarities propagate between omics; leaving most entries at zero reverts to a simpler, faster update. The resulting ANF matrix is used for spectral clustering.

###### Implementation

In ANF, we used the same range of number of neighbours ($\sigma$ not directly applicable here) and fixed values for the ANF affinity parameters `alpha = beta = 1/6` in `affinity_matrix()` and the separate fusion random walk `alpha = (1,1,0,0,0,0,0,0)` vector in `ANF()` (values suggested by the authors) [16].

The fused matrices produced for different values of nearest neighbours are practically identical (mean pairwise Pearson correlation $\sim 0.85$). Similarly to SNF and ab-SNF, the optimal number of clusters is $k_{\text{opt}}^{\text{ANF}} = 2$. The highest silhouette score is achieved for $NN = 30$ (0.786).

##### 6.1.4 Multi-omics Data Integration for Clustering to identify Cancer subtypes (MDICC)

###### Overview

MDICC [36] starts from a user–supplied similarity matrix $R^{(m)}$ for each of $M$ omics and an equal weight vector $\alpha_m = 1/M$. Their average $S^{(0)} = \frac{1}{M}\sum_m R^{(m)}$ is taken as the initial fused network.

In every iteration, the method first projects $S^{(t)}$ onto the $c$ leading eigenvectors of its graph Laplacian, yielding a denoised, low-rank matrix $F^{(t)}F^{(t)\top}$. It then sparsifies local structure by keeping only the $K$ strongest links per sample, producing $A^{(t)}$. Finally it updates each kernel weight with

$$\alpha_m^{(t+1)} \propto \exp\bigl(-\text{mean}\bigl((R^{(m)} - A^{(t)}) \odot S^{(t)}\bigr)\bigr)$$

It then renormalizes the weights so that $\sum_m \alpha_m = 1$ and recomputes the fused matrix $S^{(t+1)} = \sum_m \alpha_m^{(t+1)} R^{(m)}$. Omics whose topology is closer to the current fusion therefore receive larger weights.

After convergence, MDICC performs $k$-means clustering on the rows of the final low-rank representation $F$, returning sample clusters along with the fused similarity matrix $S$ and the learned weight vector $\boldsymbol{\alpha}$, which provides a direct measure of each omic's relative importance.

###### Implementation

Distances for original similarities were calculated as described earlier for SNF. We fix the number of neighbours used for the local affinity matrices to 18, as used by the authors (argument `k` in `testaff()`) [36]. We tuned the number of subspace dimensions (top $c$ eigenvalues: $\{2, 3, \ldots, 10\}$; argument `c` in `MDICC()`) and the number of nearest neighbours used when computing the adaptive entropy-sparsity regularisation parameters ($K \in \{41, 42, 43, 44\}$; range used in the publication; argument `k` in `MDICC()`) [36].

Silhouettes were computed using "$1 - S$" distances in each case. The highest average silhouette width ($3 \cdot 10^{-4}$) was obtained for $\{c = 6, K = 43\}$ and $k_{\text{opt}}^{\text{MDICC}} = 2$. The two clusters are largely imbalanced and therefore, not plausible biologically. No further MDICC results are presented in this supplementary document, but are available on the paper's GitHub repository (MDICC results, GitHub) and through the interactive R Shiny app (https://github.com/sionaris/MO_survey_Shiny).

##### 6.1.5 Multiple Similarity Network Embedding (MSNE)

###### Overview

Multiple Similarity Network Embedding (MSNE) [42] represents each omics layer as an undirected $K$-NN graph, performs short random walks on every graph to capture higher-order sample relationships, and feeds the resulting node sequences into a skip-gram model (Word2Vec) whose shared embedding space (default dimensions: `embed_size=128`) integrates the multiple networks. The learned low-dimensional vectors are then clustered with $k$-means to derive multi-omic subtypes.

The construction of every single-omic graph keeps the $K$ strongest affinities (default $K = 20$). For embedding, the routine `MSNE()` launches `num_walks` random walks per node on each graph (default 10). Each walk has `walk_length` steps (default 80) and produces a sequence that the skip-gram network scans with a context textttwindow_size (default 10) to learn co-occurrence statistics. The parameter `input_type` (`"symmetric relationships"` or `"directed relationships"`) specifies whether edges are treated as undirected.

##### Implementation

We modified the method's source code to also accept precomputed (dis)similarity matrices. We tuned values for $K \in \{10, 20, 30, 4050\}$, `num_walks` $\in \{50, 100, 200\}$, `embed_size` $\in \{50, 100, 200\}$, `window_size` $\in \{5, 10, 15\}$ and `walk_length` $\in \{20, 30, 40\}$. We use `workers=10` for every job. Distances are pre-calculated and provided to `MSNE()`, as described previously.

MSNE itself does not pick the number of clusters $k$. We prespecify $k = 5$ (reasonable option for breast cancer; five intrinsic molecular PAM50 subtypes), to reduce the total number of jobs to be run from 3645 to 405.

We calculate the silhouette scores in the embeddings' space of each run and select the results for the parameter combination:

$$\{K = 50, \texttt{num\_walks=200}, \texttt{embed\_size=50}, \texttt{window\_size=5}, \texttt{walk\_length=40}\}$$

(silhouette score: 0.0379). All these steps are performed in a HPC. The corresponding scripts can be found in the GitHub repository (https://github.com/sionaris/MultiOmicsSurvey/tree/main/Python/MSNE/MSNE), under `Python/MSNE/MSNE`.

##### 6.1.6 Neighbourhood-based Multi-Omics clustering (NEMO)

###### Overview

NEMO processes data by taking matrices from multiple omics types and calculating similarity matrices for each type using a radial basis function kernel with a normalisation factor $\sigma$, similarly to SNF [43]. NEMO then creates a relative similarity matrix for each omic to adjust similarities based on local neighbourhood data, making the comparison between different omics more consistent. The algorithm averages these relative similarity matrices to form an **Average Relative Similarity matrix**. This Average Relative Similarity matrix, which treats similarities as transition probabilities (akin to a random walk on a graph), is used for spectral clustering. The number of clusters is determined using a variant of the eigengap method, optimised to enhance the prognostic value by possibly suggesting a higher number of clusters. This process is implemented in the `nemo.num.clusters()` function, and we test values ranging from 2 to 10.

###### Implementation

We tested the same grid of $\sigma$ and $NN$ values as in other similarity network methods (ab-SNF, SNF). Both parametric and non-parametric tests confirm that the effect of both $\sigma$ and $NN$ on the resulting Average Relative Similarity matrix is significant. Therefore, all Average Relative Similarity matrices are subjected to the `nemo.num.clusters()` to choose the corresponding optimal number of clusters. Spectral clustering follows for each matrix for the corresponding $k$ and the optimal clustering is selected based on the silhouette index ($\{NN = 10, \sigma = 0.5\}, k_{\text{opt}}^{\text{NEMO}} = 2, avg.sil.width = 0.757$).

##### 6.1.7 Random Walk with Restart for multi-dimensional data Fusion (RWR-F)

###### Overview

In Random Walk with Restart for multi-dimensional data Fusion (RWR-F) [52], for every omics layer, a patient-by-patient similarity matrix is first built from the pairwise distances using SNF functionality: each sample keeps its $K$ strongest links and the distances are converted into affinities with a Gaussian kernel whose width is `sigma` ($\sigma$) (`K` and `sigma` in `affinityMatrix()` from *SNFtool* [86]). All layers are then stacked into a multiplex graph by adding inter-layer edges of fixed equal weight. A random walk with restart is run on this multiplex: at each step the walker moves to a neighbour with probability $1-\gamma$ or returns to its starting node with probability ($\gamma = 0.7$) (`gama` argument in `RWR_fusion()`). Larger $\gamma$ forces the walk to stay closer to the seed sample. The process is iterated up to `iteration_max` (default 1000) times or until convergence. The resulting stationary probabilities form a fused similarity matrix that is subsequently clustered.

###### Implementation

RWR-F is run on a HPC and results for each parameter combination are extracted and then processed locally. In our case, we used the same grid of parameter values for $\sigma$ and nearest neighbours $K$ we have been consistently using throughout this work. Maximum iterations and $\gamma$ are fixed to their default values ($\{1000, 0.7\}$). Both parametric and non-parametric tests confirm that the effect of $\sigma$ on similarity matrices is not significant, but the effect of $K$ (nearest neighbours) is significant. We set $\sigma = 0.5$ and use spectral clustering as it is implemented in the *kernlab* package's `specc()` function [93, 94], which is what the RWR-F source code uses. The corresponding eigenspace embeddings are used for distance calculations for silhouette scores. The highest silhouette score is achieved for $\{K = 50, \sigma = 0.5\}, k_{\text{opt}}^{\text{RWR-F}} = 2$ (0.667).

##### 6.1.8 Random Walk with Restart and neighbour information-based multi-dimensional data Fusion (RWR-NF)

###### Overview

Random Walk with Restart and Neighbor information-based multi-dimensional data Fusion (RWR-NF) builds on the methodology of RWR-F [52]. It normalizes each layer before the walk, so that densely connected omics do not dominate. The normalisation uses its own neighbour cut-off (`neighbour_num` argument in `RWR_fusion_neighbour`; default value: 10) and two scaling factors (`alpha` and `beta` in `RWR_fusion_neighbour`; default values 0.9 for both) that balance intra-layer and inter-layer transitions. Apart from this degree-normalisation step, the algorithm, restart probability (`gama`) and iteration limit remain the same as in RWR-F.

##### Implementation

RWR-NF is run on a HPC and results for each parameter combination are extracted and then processed locally. We used the same grid of parameter values for $\sigma$ and nearest neighbours $K$ we have been consistently using throughout this work. Maximum iterations, $\gamma$, `neighbour_num`, `alpha` and `beta` are fixed to their default values ($\{1000, 0.7, 10, 0.9$
$, 0.9\}$; author defaults). Both parametric and non-parametric tests confirm that the effect of $\sigma$ on similarity matrices is not significant, but the effect of $K$ (nearest neighbours) is significant. We set $\sigma = 0.5$ and use spectral clustering as it is implemented in the *kernlab* package's `specc()` function [93, 94], which is what the RWR-F source code uses. The corresponding eigenspace embeddings are used for distance calculations for silhouette scores. The highest silhouette score is achieved for $\{K = 10, \sigma = 0.5\}$, $k_{\text{opt}}^{\text{RWR-NF}} = 2$ (0.720).

##### 6.1.9 Spectrum

###### Overview

Spectrum [55, 95] first converts each omics table into a sample-by-sample similarity matrix: by default an adaptive, density-aware common-nearest-neighbour kernel is used. The view-specific kernels are summed into one affinity matrix, trimmed to its $K$-NN neighbourhood (`KNNs_p` argument in `Spectrum()`) and smoothed for `diffusion_iters` random-walk steps (if `diffusion=TRUE`) to strengthen indirect yet well-supported links. A normalised graph Laplacian is then eigen-decomposed. The optimal number of clusters is selected either by the classic eigengap (`method=1`) or by the "multimodality-gap" routine (`method=2`), which applies a dip test to the first `maxk` eigenvectors and locates the last substantial decrease governed by `frac` (fraction to find the last substantial drop) and `thresh` (how many points ahead to keep searching). The top $c$ eigenvectors are row-normalised and clustered with either $k$-means (`clusteralg='km'`) or a Gaussian mixture (`'GMM'`), yielding integrated multi-omic subtypes. Key tunables therefore include the neighbourhood scales (`NN`, `NN2`, `KNNs_p`), diffusion depth, kernel-tuning flag, and the gap-search parameters, while non-tunable, but crucial settings are the choice of kernel family and whether diffusion is applied.

###### Implementation

We modified Spectrum source functions to accommodate the usage of various distance metrics and parallelisation. We use Manhattan distance for SNPs and euclidean distance for all other modalities. The multimodality–gap selector is chosen as `method = 2`, which activates the eigenvector "dip test" routine. We use the adaptive density-aware kernel and fix `maxk=10`. We use five diffusion iterations and set `NN=3,NN2=7,frac=2,thresh=7` (default values) and tune the $K$-NN value (`KNNs_p`; range $\{10, 15, 20, 25, 30, 35, 40, 45, 50\}$) . The default $k$-means procedure is used for clustering.

There are no significant differences between the fused similarity matrices produced for different values of `KNNs_p` by Spectrum (mean Pearson correlation: 0.94). Silhouettes were then calculated in the eigenspace of each run. The maximum average silhouette width is obtained for `KNNs_p = 20` (0.734) and $k_{\mathrm{opt}}^{\mathrm{Spectrum}} = 2$.

#### 6.2 Multiple-kernel learning methods

Multiple-kernel learning methods fuse several pre-computed similarity kernels, each capturing different aspects or omics views of the data, into a single optimally weighted kernel so that downstream tasks such as clustering or classification can exploit complementary biological signals while automatically down-weighting uninformative sources.

##### 6.2.1 Cancer Integration via Multikernel LeaRning (CIMLR)

###### Overview

CIMLR learns a measure of similarity between each pair of samples in a multi-omic dataset by combining multiple Gaussian kernels per data type, corresponding to different, complementary representations of the data [96]. It enforces a block structure in the resulting similarity matrix, which is then used for dimension reduction and $k$-means clustering. CIMLR is building on the SIMLR approach [97].

The method first constructs multiple kernel matrices per modality, each of which captures local similarities under different neighbourhood settings. All these kernels are normalised and concatenated, so that both local and global sample relationships can be integrated. An iterative optimisation then learns the relative importance (weights) of each kernel while simultaneously refining a low-dimensional representation ($L$) and the final similarity matrix ($S$).

To enhance the clarity of cluster structures, CIMLR applies a diffusion process on this matrix, ensuring that samples connected through neighbours can influence each other's similarity scores. This step is designed to smooth and propagate information across the sample graph. Finally, a dimensionality reduction step (using spectral embedding or $t$-SNE) produces coordinates suitable for visualisation, and $k$-means clustering is applied to assign samples to clusters.

###### Implementation

We use a modified version of the main `CIMLR()` function (and some supportive functions) to allow for the calculation of diverse distance metrics depending on data type. We use the same distance metrics used in SNF and test the same range of nearest neighbours for the kernels. The obtained similarity matrices vary significantly between different values of nearest neighbours (average Pearson correlation $\sim 0.3$).

Therefore, the nine low-dimensional representations produced by the nine different values of nearest neighbours, were used as input for reference-based Monte Carlo consensus clustering using the *M3C* package [87, 98]. We ran the algorithm for every distinct low-dimensional representation, for a range of $k \in \{2, \ldots, 10\}$. We set the number of Monte Carlo iterations to 100, the number of resampling iterations for both reference and real data to 250, the objective function to "entropy" (default), the maximum number of clusters to 10, the reference data generation method to "reverse PCA" (default) and the fraction of points resampled in each iteration to 0.8 (default). This parameter configuration was used whenever the *M3C* package is used in this work. The total of 81 clusterings was examined in terms of stability, as measured by the Relative Cluster Stability Index (RCSI) and statistical significance (clustering Monte-Carlo $p$-value). The best performance is attained for $k_{\text{opt}}^{\text{CIMLR}} = 2$ and 15 nearest neighbours: $p = 0.046$ & RCSI $= 2.03$. This clustering ranked the highest in terms of RCSI, between the clusterings with $p < 0.05$.

##### 6.2.2 Kernel Learning Integrative Clustering (KLIC)

###### Overview

KLIC starts from $P$ pre-computed similarity kernels and seeks a patient-specific weighting of these kernels so that their weighted sum best exposes a block-diagonal (cluster) structure [22]. First, each omics view is clustered separately (kernel $k$-means) to get an initial number of patient groups $c$ and a baseline objective value. With these provisional labels fixed, the method solves a quadratic programme that, for every patient $i$, reallocates its kernel weights $\boldsymbol{\theta}_i = (\theta_{i1}, \dots, \theta_{iP})$ so that kernels whose similarities are most consistent with the current labels gain weight, while discordant kernels are down-weighted; the per-patient weights are constrained to be non-negative and to sum to one. The new weights define an updated composite kernel $K_\Theta = \sum_{m=1}^{P} \boldsymbol{\theta}_{\bullet m} \boldsymbol{\theta}_{\bullet m}^{\mathsf{T}} \odot K^{(m)}$, whose top $c$ eigenvectors provide a low-dimensional embedding on which $k$-means is reapplied to refresh the cluster labels.

The outcome is two-fold: (i) a final clustering of all samples, derived from the last eigenspace/$k$-means round, and (ii) an $n \times P$ weight matrix $\Theta$ whose rows reveal, for each patient, which omics kernels dominated its similarity profile. This local weighting contrasts with CIMLR, where a single global weight is learned for each kernel and thus cannot accommodate patient-specific data quality or biological heterogeneity. Kernels and parameters such as the number of clusters in each view (used only for the initialisation), the global cluster count $c$, and stopping tolerance govern KLIC's behaviour.

###### Implementation

We begin by building a distance matrix for each omics view similar to SNF. For every candidate within-omic cluster count $k_i \in \{2, 3, 4, 5\}$, we run a bootstrap consensus procedure adapted from Cluster Of Cluster Assignments (COCA) [20, 21] with `B=250` resamples and sampling proportion `pItem=0.8`. Each resulting $n \times n$ consensus matrix is made positive-semidefinite with `spectrumShift` and stored.

To choose the global number of clusters, we test all $4^5 = 1024$ combinations of within-omic clusters. Each consensus tensor is used as input for Local Multiple Kernel $k$-means (`lmkkmeans`) with `iteration_count`=100, for every candidate global cluster count $g \in \{2, \dots, 10\}$. The algorithm outputs patient-specific kernel weights $\Theta$; these are used to form the fused kernel $K_\Theta = \sum_{m=1}^{5} \theta_{\bullet m} \theta_{\bullet m}^{\mathsf{T}} \odot \text{CM}^{(m)}$. A silhouette score computed on the dissimilarity $1 - K_\Theta$ evaluates the clustering quality. The best silhouette across $g$ is retained for the current $(k_1, \ldots, k_5)$ combination. We select the clustering produced by the combination for which the silhouette score is the highest (0.023, $k_{\text{opt}}^{\text{KLIC}} = 2$).

##### 6.2.3 weight-boosted Multi-Kernel Learning (wMKL)

###### Overview

wMKL is building on the CIMLR approach [56, 96]. wMKL integrates several pre–computed similarity kernels and learns a non-negative weight for each kernel so that their weighted sum best exposes a chosen number of sample clusters [56]. More specifically, the steps of the algorithm include: (i) build individual kernels from every data view; (ii) use an optimisation that balances two goals—maximizing the between-cluster separation of the fused kernel's graph Laplacian and keeping the weight vector $\mathbf{w}$ small via an $\ell_1$-type penalty; (iii) solve the resulting convex quadratic (tunable penalty $\lambda$, fixed cluster count $k$) to obtain the optimal weights; (iv) perform spectral or kernel $k$-means clustering on the fused kernel $K_{\text{fused}} = \sum_m w_m K_m$.

Because the weights are learned once, not iteratively with the labels, wMKL is fast and the final $\mathbf{w}$ offers immediate interpretability—large values highlight omics that drive the separation, while small or zero weights flag uninformative sources.

###### Implementation

During the installation of the package we encountered a consistent error emerging due to underlying C++ functionality. See Section 11.4 at the end of this document for more information.

Feature weights were computed as in the case of ab-SNF (see the corresponding Section). The `CIMLR_Estimate_Number_of_Clusters_weight()` function was modified to incorporate additional distance metrics and, was used to estimate the optimal number of multi-omic clusters for the five modalities. By default, this process uses a fixed value for the number of nearest neighbours: $\max(\lceil n/20 \rceil, 10)$, where $n$ is the number of samples (in our case: $\max(\lceil 625/20 \rceil, 10) = 32$ neighbours are used). This procedure yields $k_{\text{opt}}^{\text{wMKL}} = 8$.

To obtain the final clustering, we run the `CIMLR.weight()` function (modified to incorporate additional distance metrics) and use the `c=8` (optimal number of clusters) and `k=32` (number of neighbours based on which 8 was the optimal number of clusters). We use the spectral clustering results produced by the function as our final clustering.

#### 6.3 Matrix Factorisation methods

Matrix-factorisation approaches integrate multi-omic datasets by decomposing each data matrix into a small set of shared (and sometimes view-specific) latent factors, thereby capturing the dominant cross-modal biological signals in a common low-dimensional space that can be clustered, visualised, or associated with clinical variables.

##### 6.3.1 Integrative Non-negative Matrix Factorisation (IntNMF)

###### Overview

IntNMF is a Non-negative Matrix Factorization (NMF) approach to multi-omics where one common basis is used for all modalities, but separate sets of coefficients for each modality's features [28]. Through iterative updates, a final set of basis vectors (each of which represents a sample's association with a distinct cluster) is produced, and each sample is allocated a label corresponding to the index of the basis vector with the highest value for that sample. The first $k$ basis vectors are examined, where $k$ is specified by the user or is determined based on the Cluster Prediction Index (CPI) of the examined $k$'s.

###### Implementation

This method is only used in the baseline analysis with the *MOVICS* package, using default settings, because extensive modifications are required to allow for diverse data distribution handling (see Section 7.3).

##### 6.3.2 moCluster

###### Overview

The moCluster algorithm begins by employing a modified Consensus Principal Component Analysis (CPCA) to define a Joint Latent Variable (JLV) from the input matrices of observations and features. It normalizes dataset levels using inverse eigenvalue weighting and iteratively updates the model through regressions of coefficient vectors and block latent variables, combined with a soft feature selection method, until convergence. Residual matrices are then computed differently depending on the specific integrative analysis, ensuring JLV orthogonality. Finally, the JLVs are clustered using hierarchical clustering, with the optimal number of clusters determined by the gap statistic [39].

###### Implementation

This method is only used in the baseline analysis with the *MOVICS* package, using default settings, because extensive modifications are required to allow for diverse data distribution handling (see Section 7.3).

##### 6.3.3 Low Rank Approximation clustering (LRAcluster)

###### Overview

Low Rank Approximation clustering (LRAcluster) assumes that a few major biological factors determine a set of high-dimensional but low-rank systems parameters, and the observed cancer omics data are generated based on these parameters [33]. The probabilistic assumption is that each observed molecular feature of each sample is a random variable conditional on a hidden parameter. Thus, each observed data matrix is conditional on a size-matched parameter matrix and different types of data follow different probabilistic models. The low-rank assumption of the parameter matrix leads to a penalty function corresponding to a structural complexity constraint of the model. Then, the low-rank parameter matrix can be decomposed into a low-dimensional representation of the original data, which will be used to identify candidate molecular subtypes.

###### Implementation

The LRAcluster algorithm is run on a HPC and results are extracted and then further processed locally. For the main LRAcluster function, we specify the `type` argument to be `"binary"` for SNPs and `"Gaussian"` for all other modalities. We restricted the choice of latent dimensions to the range $\{2, 3, \ldots, 10\}$, and selected the optimal number of dimensions ($r_{opt} = 7$) by inspecting the corresponding "explained variance" plot and selecting the number of dimensions where an elbow is present and, slow incremental increase follows in explained variance from then on (Figure 22d). The coordinates of the samples in the $r_{opt} = 7$ dimensions are used as input for reference-based consensus $k$-means with the *M3C* package as described earlier [87, 98]. Through this procedure, we identify $k_{\text{opt}}^{\text{LRAcluster}} = 2, p = 0.046$ (only statistically significant result).

##### 6.3.4 Multiple Factor Analysis (MFA)

###### Overview

MFA extends principal-component ideas to tabular data in which the same set of observations is described by several, conceptually distinct blocks of variables. For every block, it first performs its native ordination (PCA for quantitative variables, Multiple Correspondence Analysis (MCA) for categorical ones), then rescales the resulting factor scores so that each block contributes equally to the total inertia; this is done by dividing every block by the square-root of its first eigenvalue. All rescaled tables are concatenated and a global PCA is run on the merged matrix, yielding a common latent space where observations, variables and blocks can be jointly interpreted [37].

Because the blocks keep their identities throughout the analysis, MFA offers several levels of insight. The global factor map visualizes overall sample patterns, while "partial" maps show how each block projects those same samples, making it easy to spot blocks that drive—or disagree with—the consensus. Supplementary statistics quantify each block's contribution to every dimension.

###### Implementation

The MFA method is run on a HPC and results are extracted and then further processed locally. Due to long running times, MFA was run at the highest proportion of features per modality (33%) that allowed job completion within 12 hours. The subsets were selected based on highest mean absolute deviation (continuous modalities: RNA-seq, CNV, methylation and miRNA), while for SNPs we kept the COSMIC cancer drivers plus the top 33% of other SNPs, ranked by proportion of samples that carry a mutation. The maximum number of components was set to 200. Similarly to LRAcluster, by visually inspecting the "explained variance" plot we selected 11 as the optimal number of dimensions for MFA. We also used the Unit Invariant Knee (UIK) method from the *inflection* package [99, 100] to determine elbow points in a more objective way. UIK identifies 37 as the elbow point in terms of eigenvalues and variance explained, as well as 134 as the elbow point in cumulative variance explained. All three cases of latent dimensions were used as input for reference-based consensus $k$-means with the *M3C* package as described earlier [87, 98]. Through this procedure, we identify $k_{\text{opt}}^{\text{MFA}} = 3, p = 4.5 \cdot 10^{-6}, \text{RCSI} = 1.15$.

##### 6.3.5 Multi-Omics Factor Analysis (MOFA)

###### Overview

MOFA frames multi-omics integration as a group-factor model in which a small set of latent factors captures the principal sources of variation shared across—or specific to—each data modality [40]. Starting from normalised feature matrices that may differ in scale, distribution or even sample coverage, MOFA decomposes every matrix $Y^{(m)}$ into the product of a sample-by-factor score matrix $Z$ and a factor-by-feature loading matrix $W^{(m)}$ plus noise; priors on $W^{(m)}$ enforce element-wise sparsity, so each factor is driven by a concise subset of features. Variational inference learns the posterior distributions of $Z$, $W^{(m)}$ and the noise parameters simultaneously.

Once trained, the model delivers several layers of interpretability. Factor scores position every sample along the latent axes and can be clustered, correlated with phenotypes or visualised directly. Loadings highlight the most influential features per factor. The proportion of variance explained is broken down by factor and by omic, revealing which biological signal is global and which is modality-specific. MOFA can work with Gaussian, Bernoulli and Poisson likelihoods, automatic relevance determination for selecting the number of factors, and a flexible regression layer for covariate adjustment.

###### Implementation

MOFA is run on a HPC on a Graphics Processing Unit (GPU), and results are extracted and then further processed locally. We fit MOFA models with possible number of factors $F \in \{2, \ldots, 10, 15, 20, 25\}$. The models were run with a fixed number of maximum iterations (20,000). The likelihood types were set to `"bernoulli"` for SNPs and `"gaussian"` for the continuous modalities. Automatic Relevance Determination (ARD)-a Bayesian shrinkage prior that automatically suppresses non-informative factors- is enabled on the weights (`ard_weights=True`) so irrelevant features can be shrunk towards zero, while ARD on the factors (`ard_factors=False`) and spike-and-slab sparsity on either weights or factors (`spikeslab_weights=False`, `spikeslab_factors=False`) are disabled. The optimizer is run for in slow convergence mode (`convergence_mode="slow"`). The evidence lower bound is monitored every 5 iterations (`freqELBO=5`) starting from iteration 1 (`startELBO=1`). No factor dropping threshold is applied (`dropR2=-1`, i.e. keep all factors regardless of variance explained). A separate run with number of factors fixed to 25 and ARD on the factors is enabled for comparisons with non-ARD results.

We import MOFA results using functionality already included in the MOFA package in R [40]. The critical point in the cumulative explained variance (Figure 31) and consistently low inter-factor correlations up to $F = 7$ pointed to $F_{\text{opt}} = 7$. The 25-factor run with ARD reproduced the same "variance-explained" profile, confirming this choice for $F_{\text{opt}}$. The corresponding latent embeddings were used as input for reference-based consensus $k$-means with the *M3C* package as described earlier [87, 98]. Through this procedure, we identify $k_{\text{opt}}^{\text{MOFA}} = 3, p = 1.1 \cdot 10^{-6}, \text{RCSI} = 0.86$.

#### 6.4 Graph-based methods

##### 6.4.1 Multi Omic clustering by Non-Exhaustive Types (MONET)

###### Overview

MONET builds one weighted patient-similarity graph for each omics layer, then searches these graphs simultaneously to identify modules (sets of patients that are tightly connected in a subset of the layers) [41]. Starting from every patient as a single-node module, the algorithm repeatedly applies heuristics that grow, merge, split and refine modules in order to maximize a modularity score: edges that belong to the currently selected omic subset are rewarded, while edges outside the subset (or between modules) are penalised. Because each module is allowed to choose its own combination of omics, the final solution can contain, for example, an RNA–methylation module, a pan-omic module and a third module supported only by CNV—capturing heterogeneity that global-weight methods miss.

Key inputs are the per-omic similarity matrices (optionally offset to centre the edge-weight distribution), a minimum module size, and significance thresholds that control when a module is enlarged, merged or discarded. The output comprises (i) the list of patient modules with their chosen omic subsets, (ii) per-module edge weights that can be inspected to see which layers drove each grouping, and (iii) a residual graph of unassigned patients.

###### Implementation

MONET is run on a HPC, and results are extracted and then further processed locally. We computed a $625 \times 625$ Pearson-correlation matrix for each omic (equivalent to $\phi$-correlation for binary data), and, for a sensitivity check, an "offset" version ($-0.2$ shift) as in the original MONET paper [41]. We then ran the main MONET loop twice: once with the original versions and once with the "offset" versions of the correlation matrices. We fixed the maximum number of iterations to 10,000, the number of different seeds to be tried (`num_of_seeds`) to 100, the number of samples in a seed (`num_of_samples_in_seed`) to 10, the minimum size of modules (`min_mod_size`) to 10, the maximum number of patients per action (`max_pats_per_action`) to 10 (upper bound on patients moved per MONET step), the percentile of removed edges (`percentile_remove_edge`) to 80% (drops the lowest 20% of edge weights), and applied no global re-centering of weights.

We import MONET results in RStudio using the *reticulate* package. Weight histograms revealed that the 0.2 offset drove $> 95\%$ of edges negative in every omic, whereas the raw matrices were well-balanced. We, therefore, kept the "no-offset" solution, which yields $k_{\text{opt}}^{\text{MONET}} = 3$.

#### 6.5 Bayesian methods

##### 6.5.1 iClusterBayes

###### Overview

iClusterBayes uses a few latent variables to capture the inherent structure of multiple omics datasets to achieve joint dimension reduction [25]. As a result, the tumour samples can be clustered in the latent variable space and relevant omics features that drive the sample clustering are identified through Bayesian variable selection. In essence, iClusterBayes posits that all measured omic profiles can be summarised by a small number of hidden (latent) factors. Each sample is thus associated with a continuous latent vector, assumed to follow a standard multivariate normal distribution. This vector captures the primary patterns of variability across all data types, enabling iClusterBayes to model multiple high-dimensional datasets simultaneously.

For each feature, an indicator parameter $\gamma_{jt}$ determines whether it actively contributes to the latent space (and therefore to sample clustering) or is essentially "switched off". If $\gamma_{jt} = 1$, then feature $j$ from omic dataset $t$ is actively shaping the latent variables (and hence the clusters), whereas if $\gamma_{jt} = 0$, then feature $j$ from omic dataset $t$ the feature "switched off". This indicator is given a Bernoulli prior in the Bayesian model, which leads to a posterior distribution (i.e., an updated belief) about whether it should be 0 or 1 after seeing the data. This allows the method to focus on a subset of biologically meaningful features without discarding the rest of the data prematurely.

Different types of omics data are handled by linking the latent variables to appropriate statistical models—linear (Gaussian) for continuous measurements, logistic for binary measurements, and Poisson for count data. Throughout the process, the posterior distribution of all unknown parameters is explored, including the latent variables and the feature-selection indicators. By repeatedly drawing samples from these posterior distributions and updating model parameters, iClusterBayes builds a robust probabilistic picture of how samples cluster in the low-dimensional space and which features underpin those clusters.

Once the procedure converges, each sample's latent factor estimates can be used with classical clustering algorithms (e.g., $k$-means) to assign samples into tumour subtypes. Simultaneously, features with high posterior probability of being "selected" are deemed strong candidates for driving the observed clustering structure. As a result, iClusterBayes not only groups tumours in a biologically informed manner, but also pinpoints the key features that give rise to these clusters.

###### Implementation

iClusterBayes is run on a HPC, and results are extracted and then further processed locally. For iClusterBayes, we performed a grid search over $s_{\text{dev}} \in \{0.005, 0.01, 0.015, 0.02, 0.025, 0.03, 0.05\}$ and $\beta_{\text{var}} \in \{0.1, 0.2, 0.3, 0.4, 0.5, 0.8, 1, 1.25, 1.5, 2, 2.5, 3\}$, fixing $n_{\text{burnin}} = 1200$, $n_{\text{draw}} = 1800$, `thin`=3, $\text{pp}_{\text{cutoff}} = 0.5$ and $\boldsymbol{\gamma} = (0.5, 0.5, 0.5, 0.5, 0.5)$ as in the original specification (which also allows for running times below 12 hours). Each of the $7 \times 12 = 84$ pairs was run for $K = 2{:}10$ (`K=1:9` in code) on a HPC. For every fit we recorded the median acceptance rates $\tilde{Z}_{\text{ar}}, \tilde{\beta}_{\text{ar}}, \tilde{\gamma}_{\text{ar}}$ and defined

$$S = \sum_{K=1}^{9} \left( |\tilde{Z}_{\text{ar}} - 0.234| + |\tilde{\beta}_{\text{ar}} - 0.234| + |\tilde{\gamma}_{\text{ar}} - 0.234| \right),$$

using 0.234 as the optimal random-walk acceptance target [101, 102]. A penalty of +0.05 was added whenever a median laid outside [0.1, 0.8]. The lowest penalised score was obtained for $s_{\text{dev}} = 0.015$, $\beta_{\text{var}} = 0.5$.

Within the selected hyperparameter setting, the minimum Bayesian Information Criterion (BIC), a deviance-ratio elbow and a well-structured heatmap all agreed at $K = 5$; we therefore selected $K_{\text{opt}} = 5$ and the corresponding cluster labels for downstream analyses.

#### 6.6 Ensemble clustering methods

##### 6.6.1 Perturbation clustering for data INtegration and disease Subtyping (PINSPlus)

###### Overview

PINSPlus is an unsupervised approach for subtype discovery without using any a priori knowledge (such as clinical variables or known subtypes) [47]. The method is based on the observation that small changes in quantitative assays will be inherently present between individuals, even in a truly homogeneous population. If distinct molecular subtypes do exist, they must be stable with respect to small changes in quantitative assays. In order to discover reliable subtypes, PINSPlus estimates how often each pair of patients is grouped together in the following scenarios: (i) when data are perturbed, (ii) when using different data types and (iii) when using different clustering techniques. PINSPlus then partitions patients into subgroups that are strongly connected in all scenarios.

###### Implementation

This method is only used in the baseline analysis with the *MOVICS* package, using default settings, because extensive modifications are required to allow for diverse data distribution handling (see Section 7.3).

##### 6.6.2 Cluster Of Cluster Assignments (COCA)

###### Overview

COCA is a two-stage consensus strategy that first derives clusters within each omics layer and then integrates those partitionings into a single, cross-omic subtype solution [20, 21]. For every data type, COCA performs clustering e.g., with hierarchical clustering. Each run yields a connectivity matrix that records whether two samples co-occur in the same single-omic cluster.

The connectivity matrices from all resamples and all omics are averaged into one consensus matrix whose entries represent the probability that a pair of samples was grouped together anywhere in the bootstrap procedure. This consensus is then re-clustered (e.g., by hierarchical clustering) to produce the final multi-omic subtypes.

###### Implementation

We first built a $625 \times 625$ Multi-Omic Consensus (MOC) matrix with `buildMOC()` using all five omics ($M = 5$) and evaluating $K \in \{2, \ldots, 10\}$; single-omic partitions were produced with Ward hierarchical clustering on Jaccard distance for SNPs and Euclidean distance for the four continuous omics. The resulting MOC was converted to a Jaccard dissimilarity matrix and reclustered with Ward's method, yielding a consensus dendrogram. Based on average silhouette and the gap statistic, a number of clusters of 5 or 6 would seem optimal. However, both of these solutions included 3 or 4 clusters with no more than 3 samples in each, respectively. We therefore proceeded with $k_{\text{opt}}^{\text{COCA}} = 2$, which produces a 557/68 sample partition.

##### 6.6.3 Consensus Clustering

###### Overview

Consensus clustering is running multiple clusterings on subsamples of the input data, records the sample allocations in each clustering and eventually generates a consensus matrix which essentially describes sample similarities based on the proportion of times these samples clustered together [23]. The final matrix is used to cluster the full input. The functionality from the *ConsensusClusterPlus* package is used [24].

###### Implementation

This method is only used in the baseline analysis with the *MOVICS* package, using default settings, because it works with the concatenation of the omic matrices (see Section 7.3).

#### 7 Baseline analysis

The analysis described in this section has been used as a baseline, to which, results from all other individual methods and the final consensus pipeline were compared.

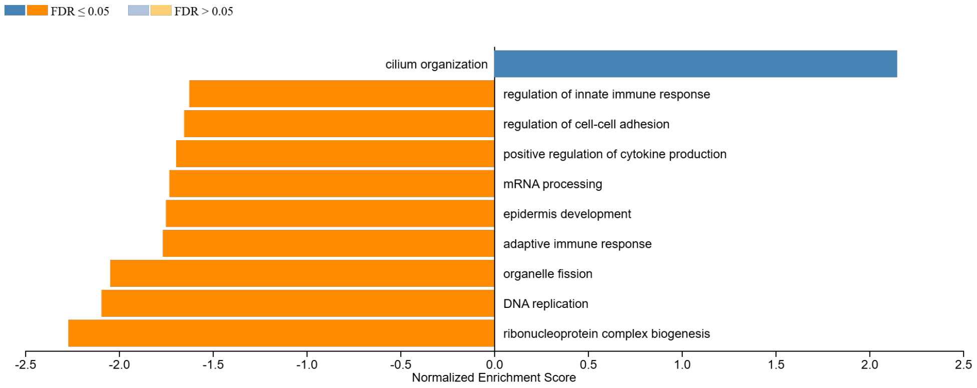

**Figure 2. Enriched pathways in the ER+ cluster.** The opposite direction of enrichment holds for the ER- cluster. Only $FDR < 0.05$ results shown here.

#### 7.1 Overview

We carry out an analysis on pure ER status-based clusters (biological pattenrs of the clusters can be seen on Figure 2), plus, *MOVICS* is used to perform multi-omics integrative clustering and visualisation for cancer subtyping research. *MOVICS* provides a unified interface for 10 state-of-the-art multi-omics clustering algorithms: CIMLR [96], iClusterBayes [25], moCluster [39], COCA [20], Consensus Clustering [23], IntNMF [28], LRAcluster [33], NEMO [43], PINSPlus [47] and SNF [54], where the former three methods can also perform the process of feature selection. *MOVICS* suggests an estimate of the optimal number of multi-omic clusters $k$ and then proceeds by using each clustering algorithm to cluster the multi-omic data in $k$ clusters. Functionality is also provided to evaluate results on an external cohort; here we choose the transNEO cohort for this purpose [73].

#### 7.2 Optimal number of clusters

The first part of the *MOVICS* pipeline is to identify the optimal number of multi-omic clusters in our data. For this, we use the `getClustNum()` function. This particular step involves setting negative values to 0 (by adjusting them based on the minimum value), and scaling all values to a maximum of 1. The function uses the `nmf.opt.k()` function from the *IntNMF* package [29] to compute the CPI for a range of cluster numbers, specified by the argument `try.N.clust`, which we set to the range $\{2, 3, 4, 5, 6, 7, 8, 9, 10\}$. This function performs NMF, which is used to reduce dimensionality and detect patterns in multi-omics data, and estimates the quality of clustering for each possible number of clusters by assessing the stability of clusters across multiple runs. This way, the CPI is calculated. The optimal $k$ is determined based on the CPI using a resampling-based cross-validation technique.

getClustNum() also computes gap statistics using the moGap() function of the *mogsa* package [103], after performing Multiple Omics data integrative Cluster Analysis (MOCA) with mbpca(). Gap statistics compare the total within-cluster variation for different numbers of clusters against the expected variation under a reference null distribution that is generated using Monte Carlo simulations. This helps in identifying a clustering structure that has a significantly lower within-cluster variation compared to the null model.

In the final step, getClustNum() identifies the optimal number of clusters **by combining the results from the CPIs and the gap statistics**. It computes the sum of the normalised values from both metrics for each cluster number and identifies the number where this sum is maximised. In our work, this was $k = 2$ as shown in the main text and Figure 3a.

As shown in Figure 3e, both COSMIC criteria-based and simple criteria-based FGA patterns show significant differences between subtypes, but with opposite directionality. Unique biological processes enriched in each subtype were derived through the computation of GSVA scores after carrying out GSEA based on the unique expression markers of each subtype (Figure 3f,i). No distinguishable patterns were observed with respect to immune biological processes between the two subtypes (Figure 3f). CS1 is predicted to be significantly less sensitive to sorafenib ($p = 2.2 \cdot 10^{-7}$) compared to CS2 (Figure 3h).

Consensus subtype labels were obtained for the external cohort samples using NTP and PAM, with the external standardised expression profiles as input. Label mapping with NTP and PAM demonstrates a high level of agreement on both the transNEO ($\kappa = 0.974, p < 0.001$, 151/153 external samples labelled identically) and the TCGA holdout set ($\kappa = 0.893, p < 0.001$, 441/466 external samples labelled identically). In the TCGA holdout cohort, the consensus subtypes are associated with ethnicity ($p = 0.023$), race ($p < 0.001$), age ($p = 0.031$), histological type ($p < 0.001$), ER status ($p < 0.001$), progesterone receptor (PR) status ($p < 0.001$), HER2 status ($p < 0.001$) and stage ($p = 0.015$). The two subtypes in transNEO are associated with pCR ($p = 0.049$), lymph node status at diagnosis ($p = 0.019$), ER status ($p = 0.01$), but not HER2 status ($p = 0.392$), Prediction Analysis of Microarray 50 (PAM50) subtype ($p < 0.001$), iC10 subtype ($p < 0.001$), tumour grade ($p < 0.001$), residual cancer burden (RCB) score ($p < 0.001$), STAT1.gsva score ($p < 0.001$), GGI.gsva score ($p < 0.001$), ESC.gsva score ($p < 0.001$), HRD sum ($p < 0.001$). With regard to ER status (only common clinical variable between cohorts with significant association with the subtypes), the same direction of association is observed: CS2 demonstrates a higher proportion of ER+ samples.

#### 7.3 Parametrisation of methods in MOVICS

This section describes the default parameters used by *MOVICS* when multiple methods are run on the same input with one function call. We use this approach in our baseline analysis.

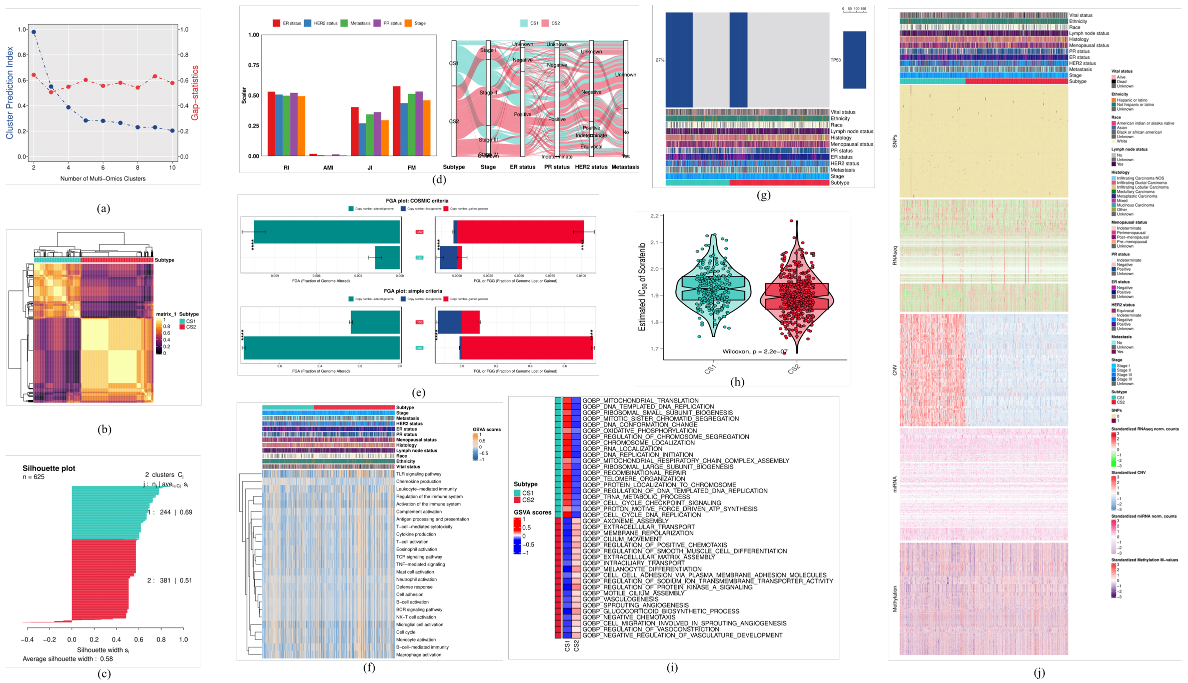

**Figure 3. Baseline MOVICS analysis. (a)** Optimal number of clusters determined by *MOVICS*. **(b)** Consensus similarity heatmap showing two subtypes (CS1 and CS2). **(c)** Silhouette plot for the consensus clusters. **(d)** Subtype agreement metrics and alluvial plot. **(e)** Fraction of Genome Altered plots: COSMIC criteria and simple criteria. **(f)** Immune gene sets GSVA scores across clusters. **(g)** Oncoprint plot. **(h)** Drug sensitivity (IC50) violin plot for sorafenib. **(i)** Top 20 representative up-regulated pathways per subtype. **(j)** Comprehensive heatmap of consensus subtypes and omic modalities, showing clinical annotations at the top.

#### SNF

In *MOVICS*, the SNF algorithm is run for $K = 30$ nearest neighbours, $\sigma = 0.5$ and number of iterations $t = 20$. A grid of parameters can be used if the single algorithm option of `getMOIC()` is used. Additionally, Euclidean distance is used regardless of the type of data (continuous vs. binary).

#### NEMO

*MOVICS* uses default parameters with NEMO. The number of neighbours is defined as $k = \frac{\#samples(l)}{6}$ for modality $l$, and $\sigma$ is set to 0.5.

#### CIMLR

The dimensionality of $L$ ($C$) is defined by the selected number of clusters (e.g. 2). The weights for each kernel are initialised uniformly ($w_{l_0} = 1/55$). The similarity matrix $S$ is iteratively adjusted through a combination of an adjacency matrix $A$, computed based on nearest neighbours and distances, and diffused and adjusted by the $\delta$ parameter, mixing the prior and current states of $S$. The number of nearest neighbours for matrix $S$ is set to $nn' = 10$, by default. The loop runs `NITER = 30` times (internal parameter). After the final similarity matrix $S$ is obtained, $t$-SNE is used to project to low dimensions and perform $k$-means clustering there for the preset value of $k$. The only parameter that *MOVICS* sets when implementing CIMLR is `Nclust` (selected number of clusters) which also controls the dimensionality of $L$ ($C$), unless specified otherwise.

#### IntNMF

The number of clusters $k$ is set to 2. The Non-Negative Double Singular Value Decomposition (NNDSVD) method is used for the initialisations (`30` initialisations) [104]. The maximum number of iterations is set to `200` and the stability count for convergence is set to `20`. **No weights** are used for different datasets. The algorithm is run in a nested loop, for each initialisation ($1 \ldots 30$) and for each iteration ($1 \ldots 200$). The best solution across initialisations is the $W$ with the minimum $Q_W$ and the corresponding $H_i$'s. Using the final $W$, each sample is allocated to the cluster (column index) that has the highest value for that sample.

#### LRAcluster

The *MOVICS* implementation of LRAcluster lets the user define the number of components in the low-dimensional space (in our case `N.clust = 2`) and uses hierarchical clustering in the output (Ward method), instead of the default $k$-means.

#### moCluster

In *MOVICS*, the CPCA focuses on maximizing the variance explained across all datasets collectively (`method = "globalScore"` - concatenated input matrices). Euclidean distances are calculated in the 2-dimensional PCA space and then used for hierarchical clustering and pruning at two groups using the Ward method.

##### iClusterBayes

The $\gamma$ prior is set to 0.5 for all features. 18000 burn-in iterations are used for the Markov Chain Monte Carlo (MCMC) sampling. 12000 draws (samples) from the posterior after burn-in are used to estimate the posterior distribution of parameters. Burn-in iterations help the MCMC mechanism settle into its stationary distribution. There is a thinning parameter for the MCMC which is set to 3, which means only every third sample is kept to reduce auto-correlation between consecutive samples. **We set the number of burn-in iterations to 1800 and the number of draws to 1200 to reduce running times**.

##### COCA

*MOVICS* uses default settings for COCA, i.e. Euclidean distance, hierarchical clustering during and after COCA, complete linkage during COCA, Ward after COCA. When pre-specifying a number of clusters - as in our case (`Nclust = 2`) - all modalities are hierarchically clustered and the tree for each modality is cut at 2 groups. Subsequently, the final hierarchical clustering is performed and cut at 2 clusters as well.

##### PINSPlus

*MOVICS* is using the default settings of PINSPlus. It uses iterations $n$ by specifying $n_{min} = 50$ and $n_{max} = 500$. It uses the introduction of Gaussian noise as perturbation method and $k$-means as the clustering algorithm. Our dataset has more than 200 samples, so PCA results in $N - 1$ dimensions. However, *MOVICS* is **concatenating the different omics datasets into one dataset** of dimensions $N \times \sum_{i=1}^{T} d_i$ and performs perturbation clustering on the concatenated dataset (according to the source code; `View(MOVICS::getPINSPlus)`). Essentially, $k$-means clustering is performed on the concatenated matrices.

##### Consensus Clustering

In *MOVICS*, the data from multiple modalities are concatenated based on common samples. The proportion of features and samples selected at each subsampling is 0.8. The number of iterations is $T = 500$. Pearson correlation is used for the distances. The clustering algorithm used for the initial clusterings and the final clustering is hierarchical clustering using Ward linkage. The dendrograms are pruned to create $k$ groups.

#### 7.4 Consensus approach

We selected all ten algorithms for this consensus pipeline. *MOVICS* runs them all with default parameters (see Section 7.3 for details on these parameters) and then uses the predicted cluster labels to get the consensus labels, using the classical consensus clustering approach [23].

#### 7.5 MOVICS main results

The main MOVICS baseline results are presented in the paper and here in Figure 3. Figure 3b shows the hierarchical clustering of the two consensus subtypes and is coloured by the pairwise consensus similarity of samples. The corresponding silhouette plot is shown in Figure 3c. The vast majority of samples have positive silhouette values. Average silhouette width is 0.58, which indicates that samples within each cluster are sufficiently similar to one another and sufficiently dissimilar to samples from the other cluster. A heatmap of the five input matrices, accompanied by additional clinical annotation is shown in Figure 3j. Continuous data (RNA-seq, CNV, miRNA and Methylation) have had their extreme values truncated at $-3$ and 3 for better colouring. CNV is the modality with the clearest separation across subtypes.

With respect to clinical variables, noteworthy statistically significant associations were found between the baseline consensus subtypes and histological type ($p < 0.001$), ER status ($p < 0.001$), HER2 status, and ER cell level percentages and stage (all sub-stages included. The FGA plots for the consensus subtypes show significant genome alterations across clusters. However, according to the simple criteria FGA plot (Figure 3e) there is significantly increased genome gain in CS1 and increased loss in CS2, while opposite effects are observed in the plot drawn using COSMIC criteria. A heatmap of the top 20 represnetative (hierarchically clustered) deregulated pathways is produced for the baseline consensus subtypes, that is shown in Figure 3i. The GSVA plot for immune pathways is shown in Figure 3f. CS1 is significantly enriched in *TP53* mutations (Figure 3g; 42.2% of samples have *TP53* mutations, compared to CS2 where this is reduced to 17.8%, $p_{adj} = 4.31 \cdot 10^{-10}$). Comparisons of drug sensitivity (Figure 3h) across the baseline consensus subtypes showed that CS1 has increased predicted sensitivity to sorafenib.

**Overall**:

1. There are 2 predicted clusters (consensus subtypes) in the TCGA cohort.
2. There are 244 CS1 samples and 381 CS2 samples.
3. Normalised CNV data seem to drive the separation between CS1 and CS2, based on the multi-omic heatmap.
4. Special findings about the consensus subtypes:

- CS1 has 77.8% Invasive Ductal Carcinoma (IDC) samples, while CS2 has 64.8%. CS1 has 11.5% Invasive Lobular Carcinoma (ILC), while CS2 has 23.6%.
- 69.5% of CS1 samples are ER+, while 83% of CS2 samples are ER+.
- 59.6% of CS1 samples are PR+, while 73.6% of CS2 samples are PR+.
- There are significant differences in the FGA bars of the two subtypes.
- CS1 is associated with decreased predicted sensitivity to sorafenib.
- Oncoprint showed association of CS1 with TP53 mutations.

##### 7.5.1 Agreement between algorithms

The similarity of clustering allocations and the clusters' biological profiles produced by different algorithms employed by *MOVICS*, is shown in Figure 4. A subtype's biological profile was determined based on the identified subtype-specific significant pathways. Clustering allocations are more similar between iClusterBayes, IntNMF, Consensus Clustering, LRAcluster and PINSPlus. The clusters produced by NEMO and SNF are also similar to each other, which may be a result of the similarities in the underlying methodology. SNF and NEMO clusters also demonstrate moderate similarity to the results of moCluster.

In terms of biological profiles of the identified clusters, most pairs exhibit overlap coefficients higher than 0.5, indicating relatively similar biological profiles. The highest overlaps are noted in the pairs with strong clustering allocation similarities, i.e. the pairs with purple and dark purple colours in the corresponding cells on the heatmap.

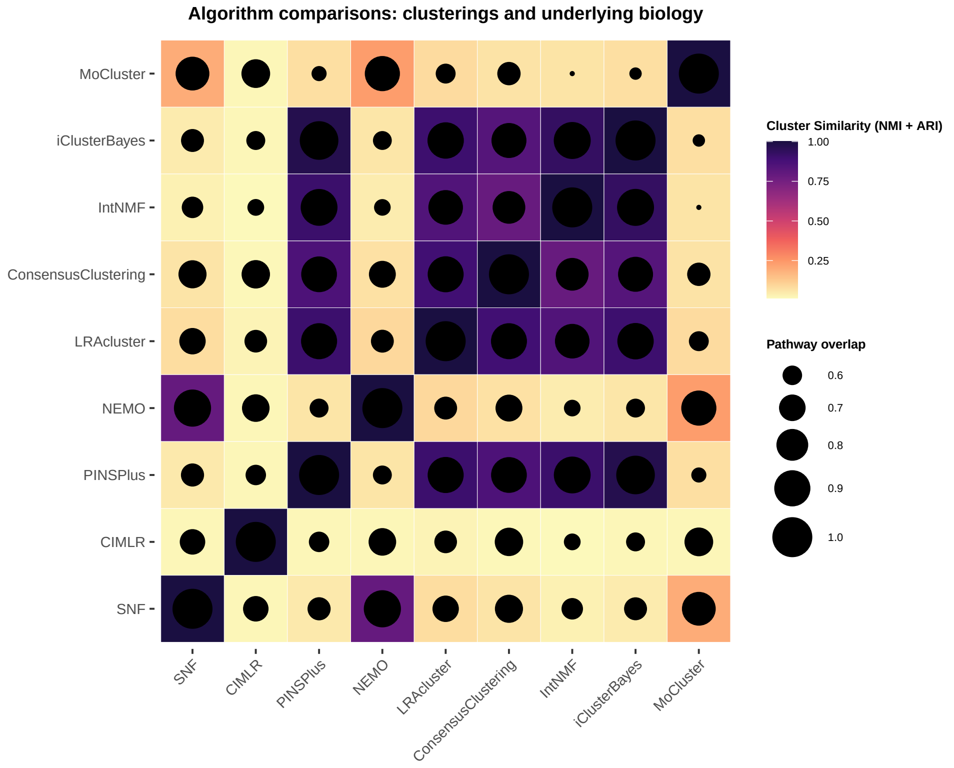

**Figure 4. Similarities between algorithm clustering outputs.** The clustering allocation similarity between algorithms is measured by the sum of the Adjusted Rand Index (ARI) and the NMI for each pair and coloured using the magma palette. The pair-wise similarities in terms of significant pathways is measured by the overlap coefficient $(A \cap B \div \min(A, B))$ and determines the size of the black circles in each heatmap cell.

### 8 Results from individual method runs

#### 8.1 Similarity Network methods

##### 8.1.1 ab-SNF, ANF, NEMO, RWR-NF, SNF and Spectrum

Results from ab-SNF, ANF, NEMO, RWR-NF, SNF and Spectrum are highly similar and are presented together below. Silhouette scores, ER status bar charts, oncoprint plots, network graphs, FGA plots with COSMIC criteria, feature rankings (not for Spectrum) and methylation data dimensionality reduction plots are presented in Figures 5, 6, 7, 8, 9, 10 and 11. Pathways which are repsentative and up-regulated in each cluster are illustrated in Figure 12.

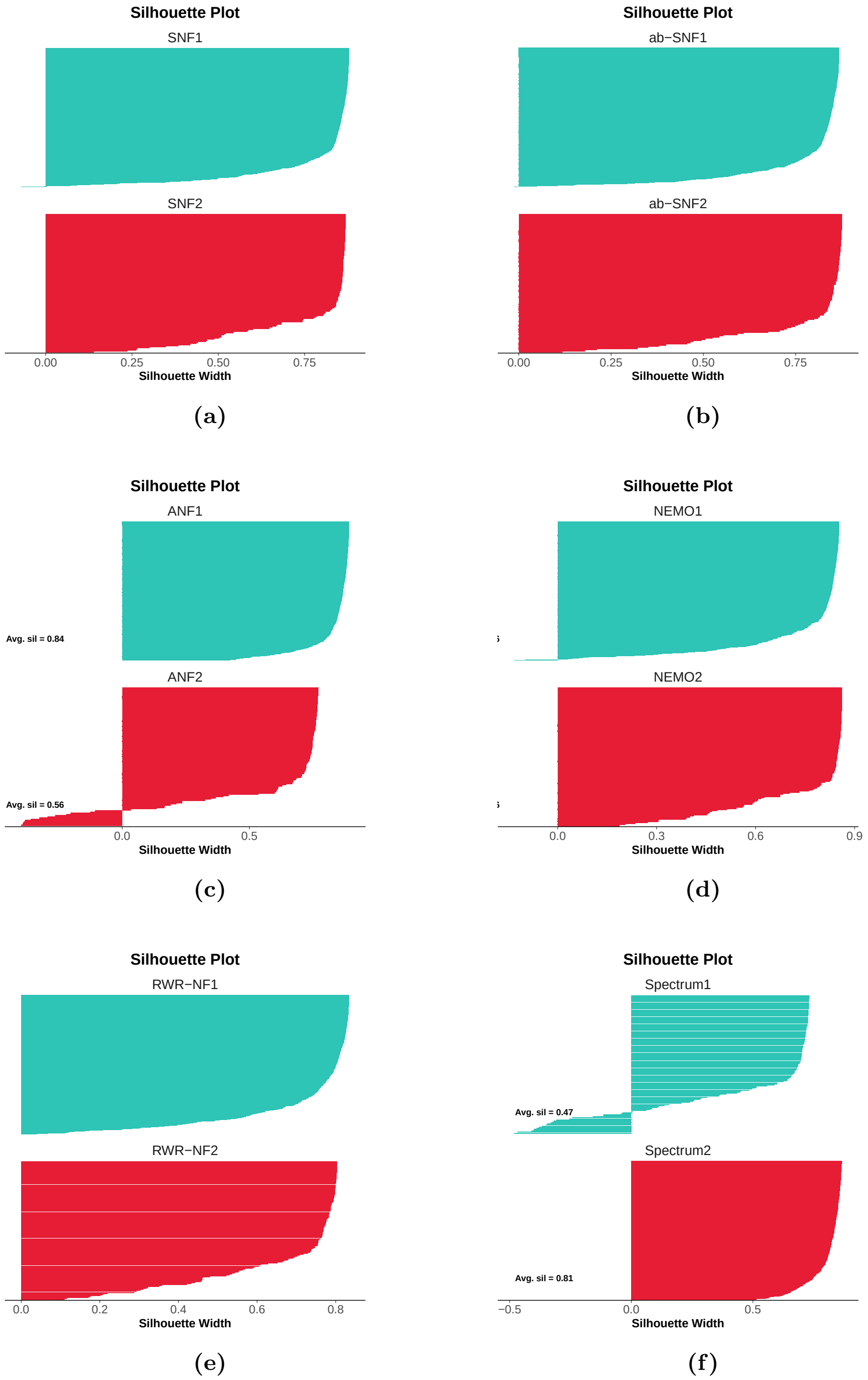

**Figure 5. Silhouette score comparisons for six similarity network algorithms:** (a) SNF, (b) ab-SNF, (c) ANF, (d) NEMO, (e) RWR-NF and (f) Spectrum.

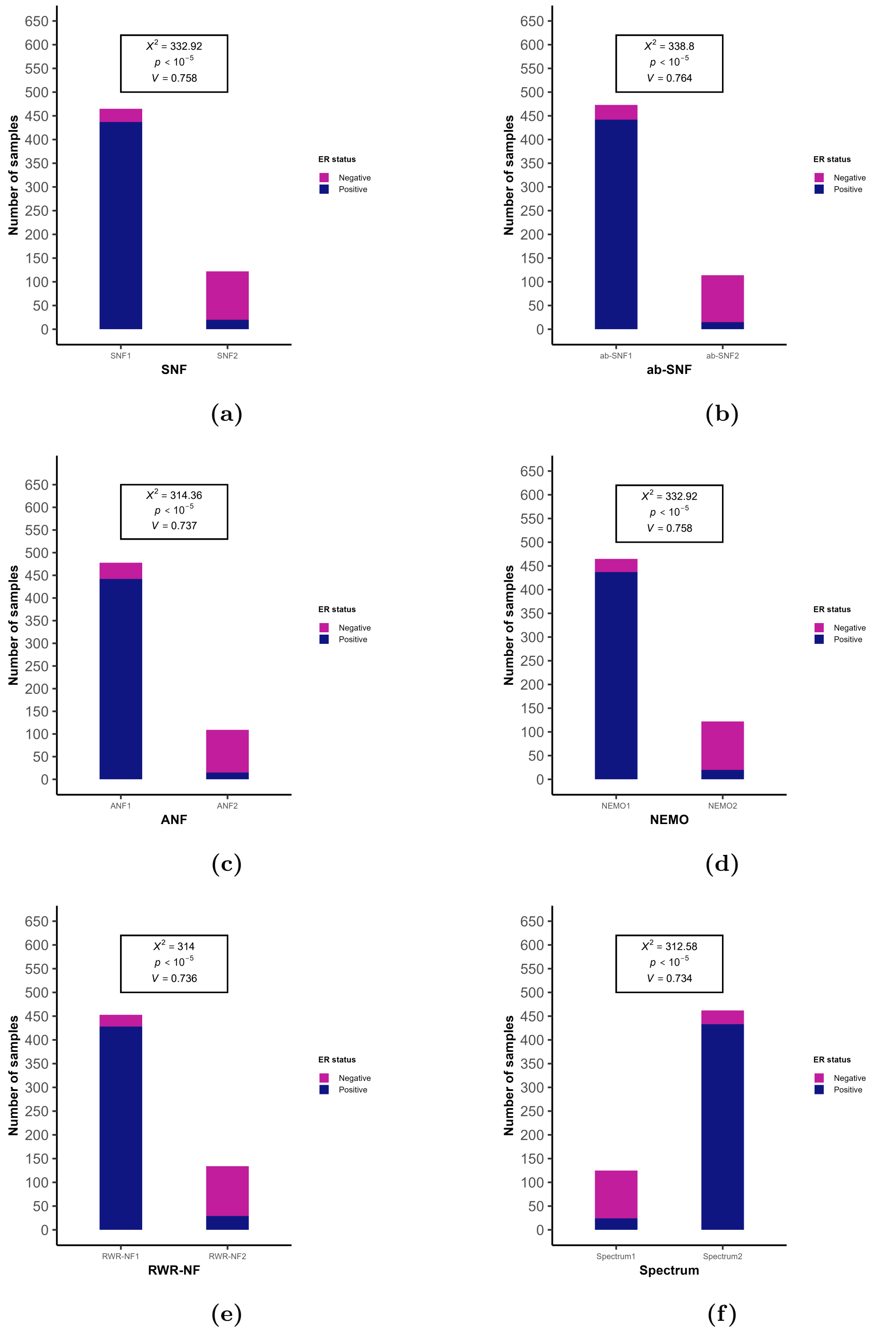

**Figure 6. ER status - Cluster bar charts with Chi-square test and Cramér's V:** (a) SNF, (b) ab-SNF, (c) ANF, (d) NEMO, (e) RWR-NF and (f) Spectrum.

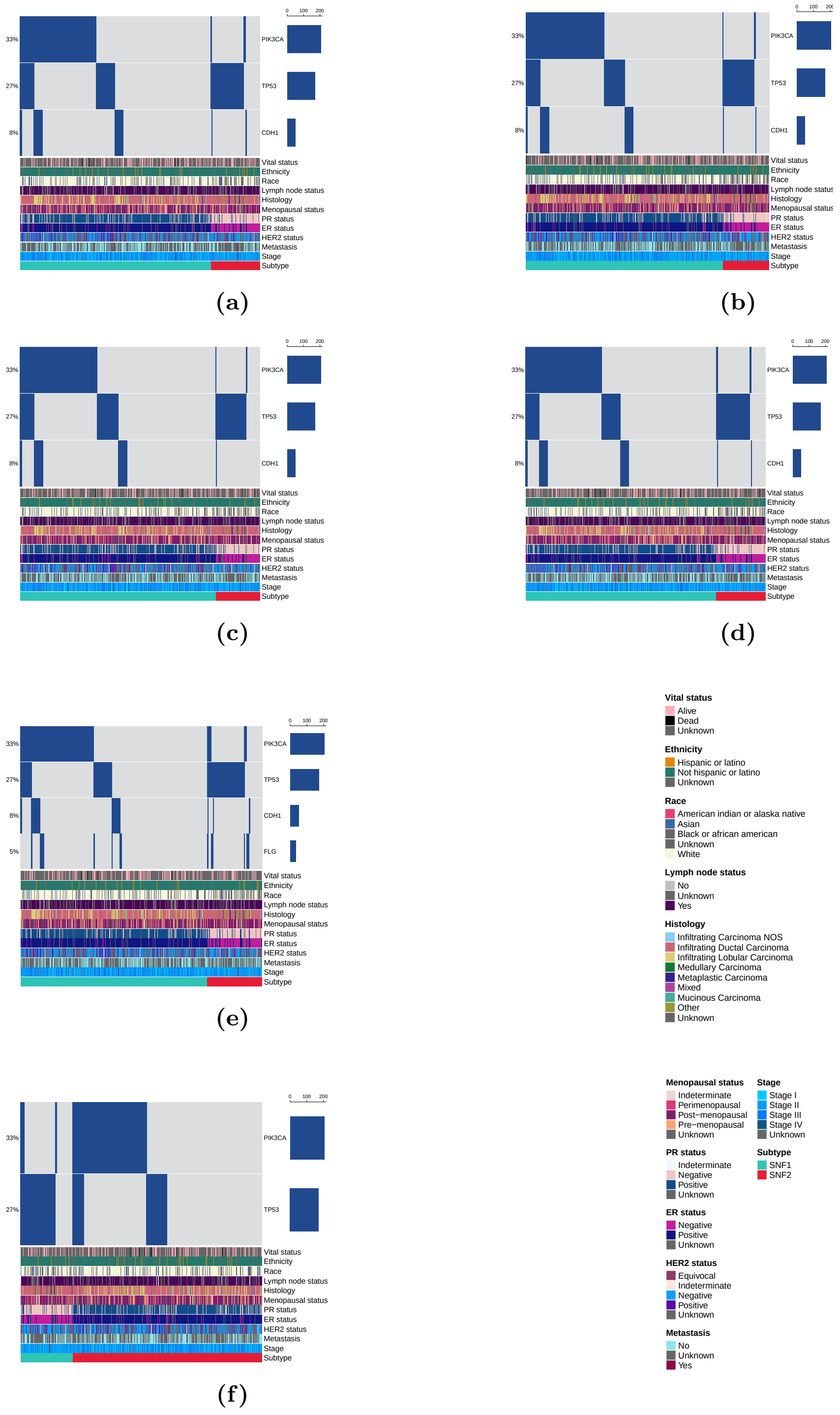

**Figure 7. Oncoprint plots for six similarity network algorithms:** (a) SNF, (b) ab-SNF, (c) ANF, (d) NEMO, (e) RWR-NF and (f) Spectrum.

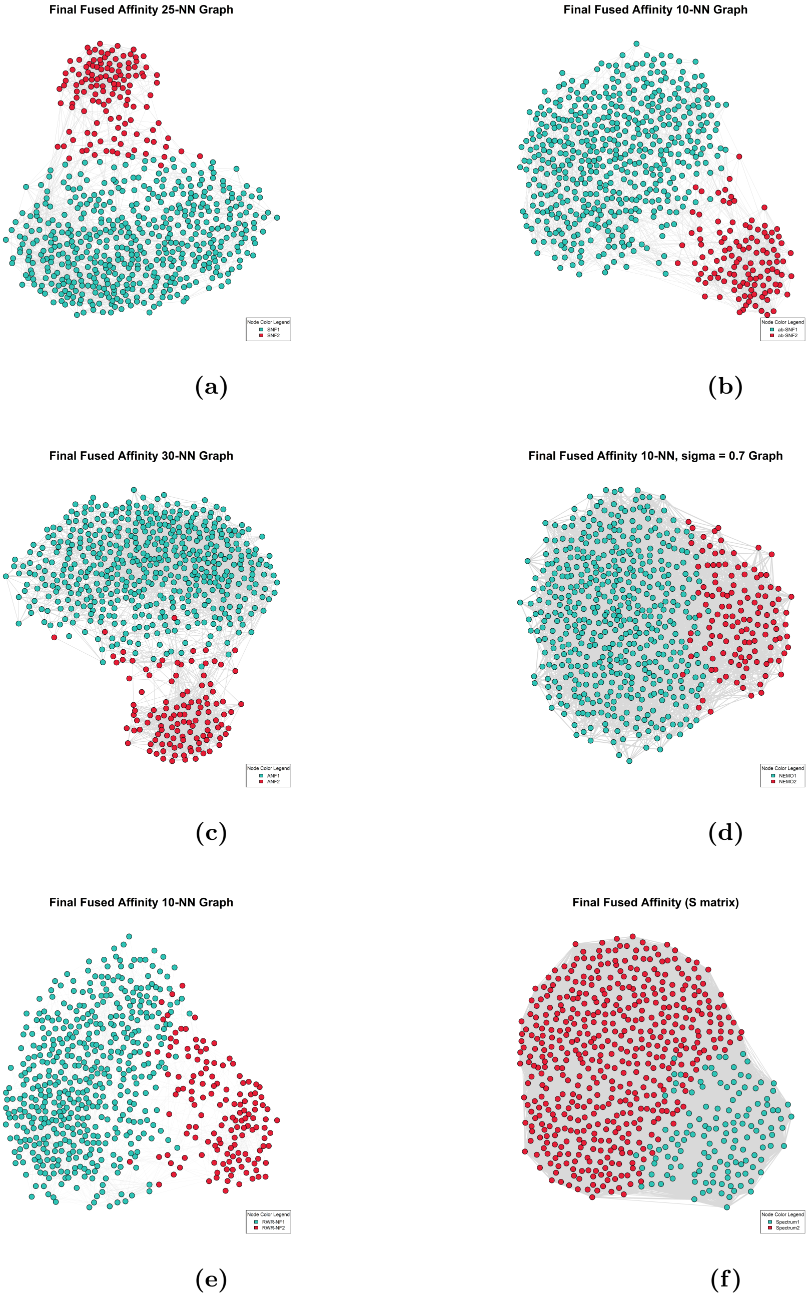

**Figure 8. Network graphs for six similarity network algorithms:** (a) SNF (25 nearest neighbours), (b) ab-SNF (10 nearest neighbours), (c) ANF (30 nearest neighbours), (d) NEMO (10 nearest neighbours), (e) RWR-NF (10 nearest neighbours) and (f) Spectrum (20 nearest neighbours).

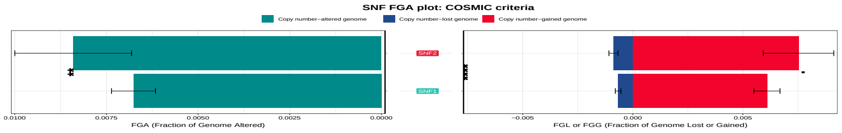

**(a)**

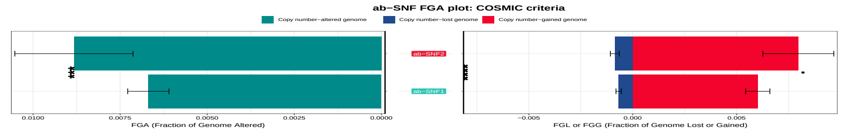

**(b)**

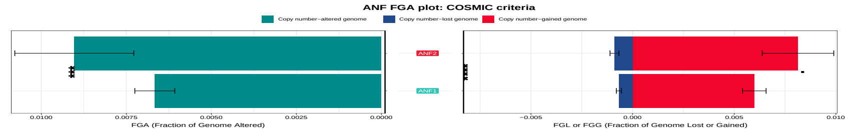

**(c)**

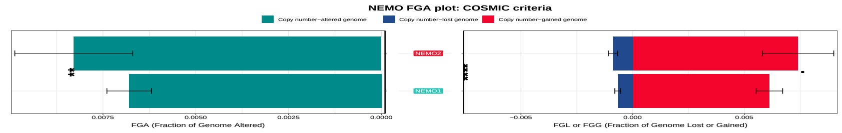

**(d)**

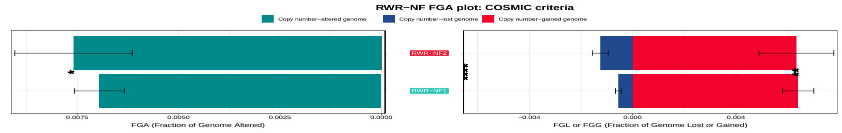

**(e)**

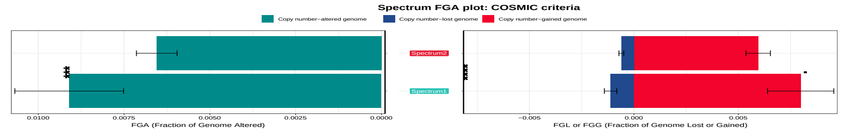

**(f)**

**Figure 9. Fraction of Genome Altered (FGA) plots for six similarity network algorithms (COSMIC criteria):** (a) SNF, (b) ab-SNF, (c) ANF, (d) NEMO, (e) RWR-NF and (f) Spectrum.

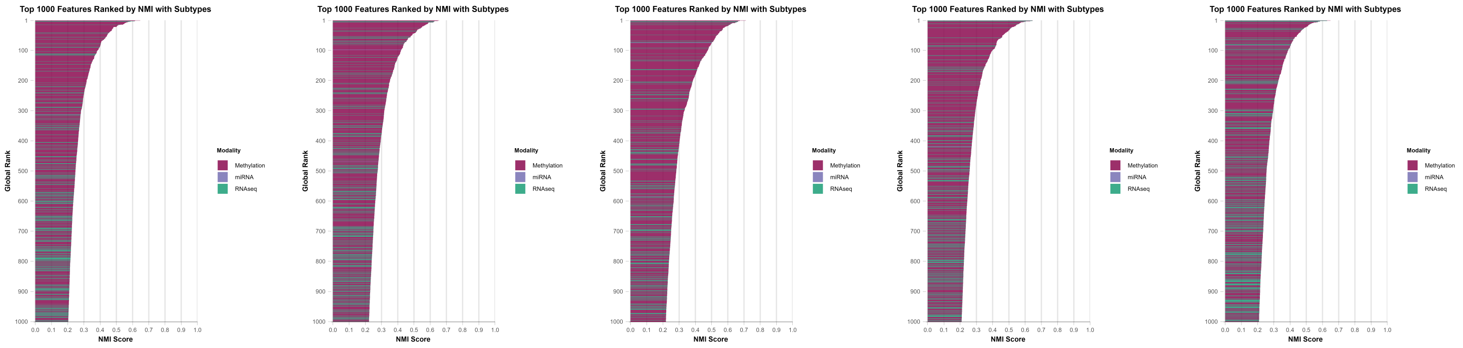

(a) (b) (c) (d) (e)

**Figure 10. Feature-rank comparisons for five similarity networks algorithms (top 1000 features):** (a) SNF, (b) ab-SNF, (c) ANF, (d) NEMO, and (e) RWR-NF.

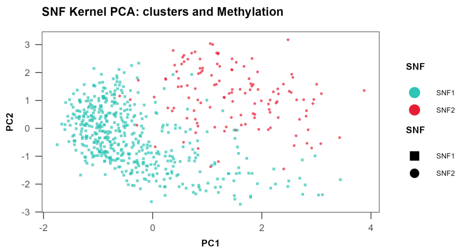

**(a)**

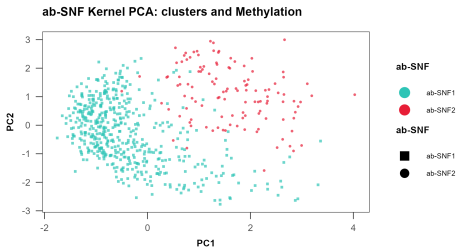

**(b)**

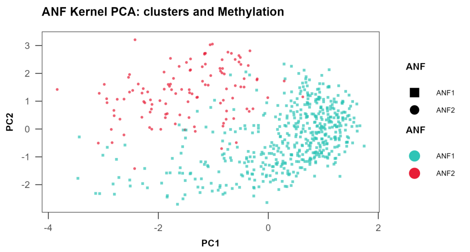

**(c)**

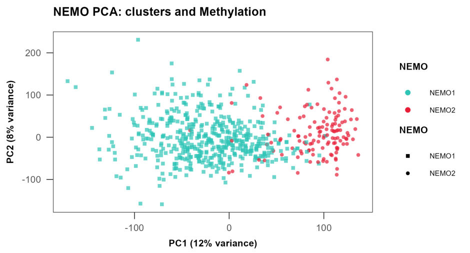

**(d)**

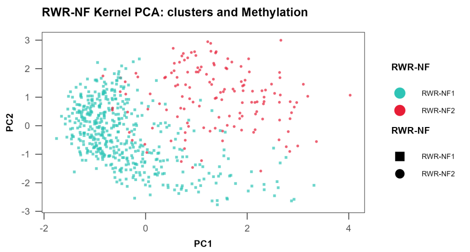

**(e)**

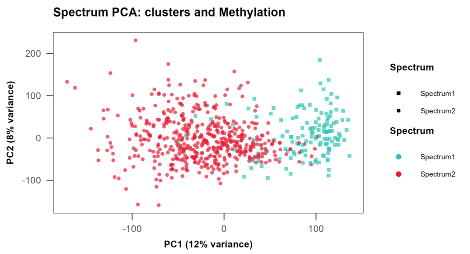

**(f)**

**Figure 11. Dimensionality reduction plots for methylation data and clusters:** (a) SNF (kernel PCA), (b) ab-SNF (kernel PCA), (c) ANF (kernel PCA), (d) NEMO (original matrix PCA), (e) RWR-NF (kernel PCA) and (f) Spectrum (original PCA).

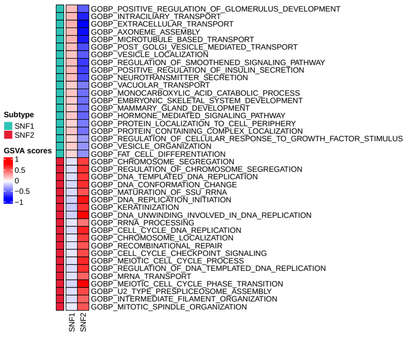

**(a)**

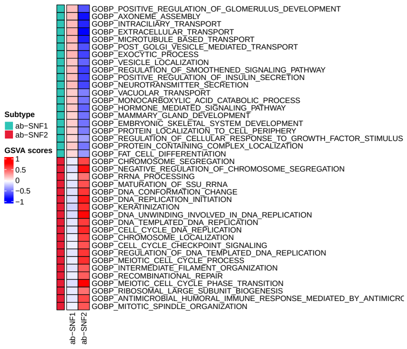

**(b)**

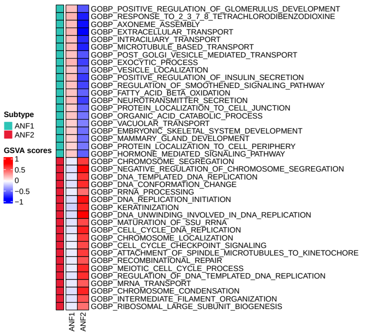

**(c)**

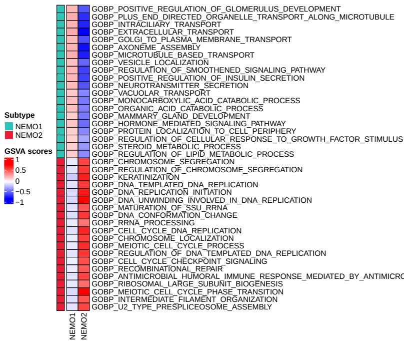

**(d)**

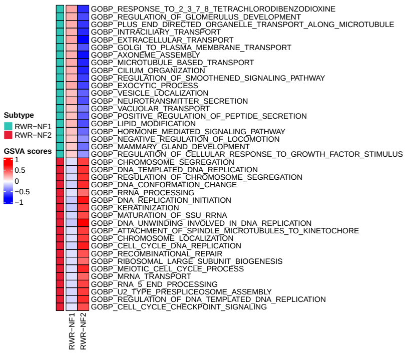

**(e)**

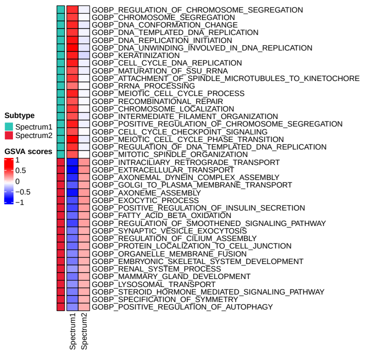

**(f)**

**Figure 12. Up-regulated pathways in each cluster for six similarity network methods:** (a) SNF, (b) ab-SNF, (c) ANF, (d) NEMO, (e) RWR-NF and (f) Spectrum.

##### 8.1.2 MSNE

MSNE low-dimensional projection plots, oncoprint, ER status bar chart and FGA plots are shown in Figure 13.

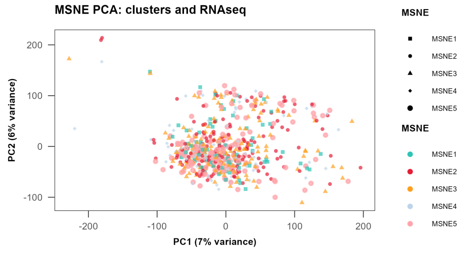

(a)

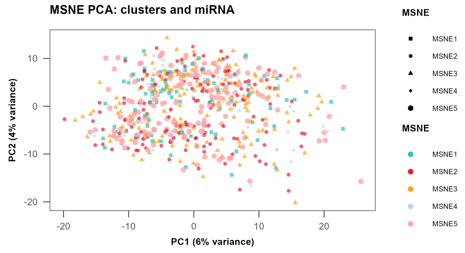

(b)

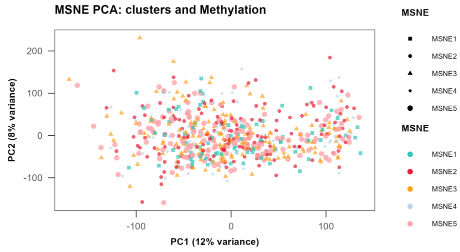

(c)

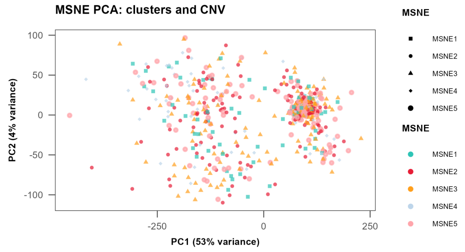

(d)

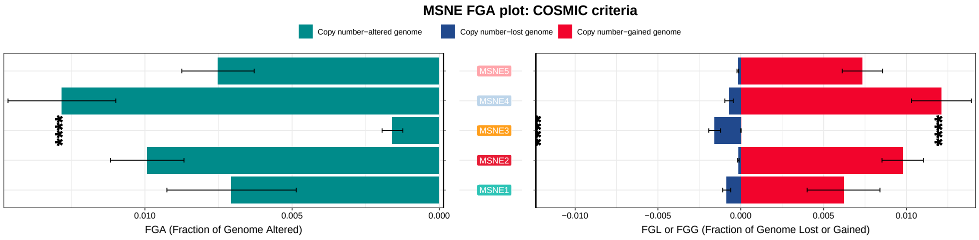

(e)

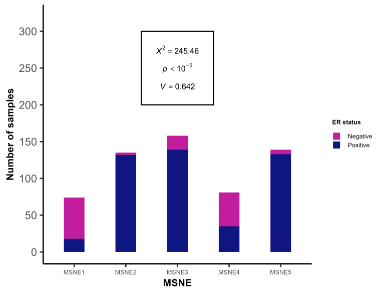

(f)

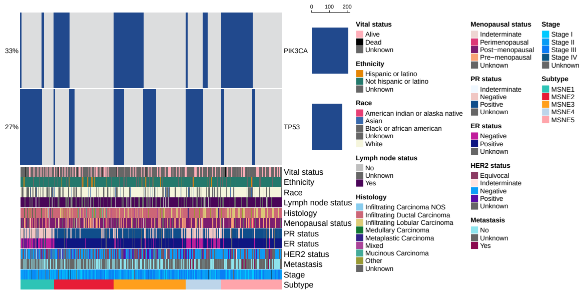

(g)

**Figure 13. MSNE results:** (a) RNA-seq data PCA plot, (b) miRNA data PCA plot, (c) Methylation data PCA plot, (d) CNV data PCA plot, (e) FGA plot, (f) ER status bar chart and (f) oncoprint.

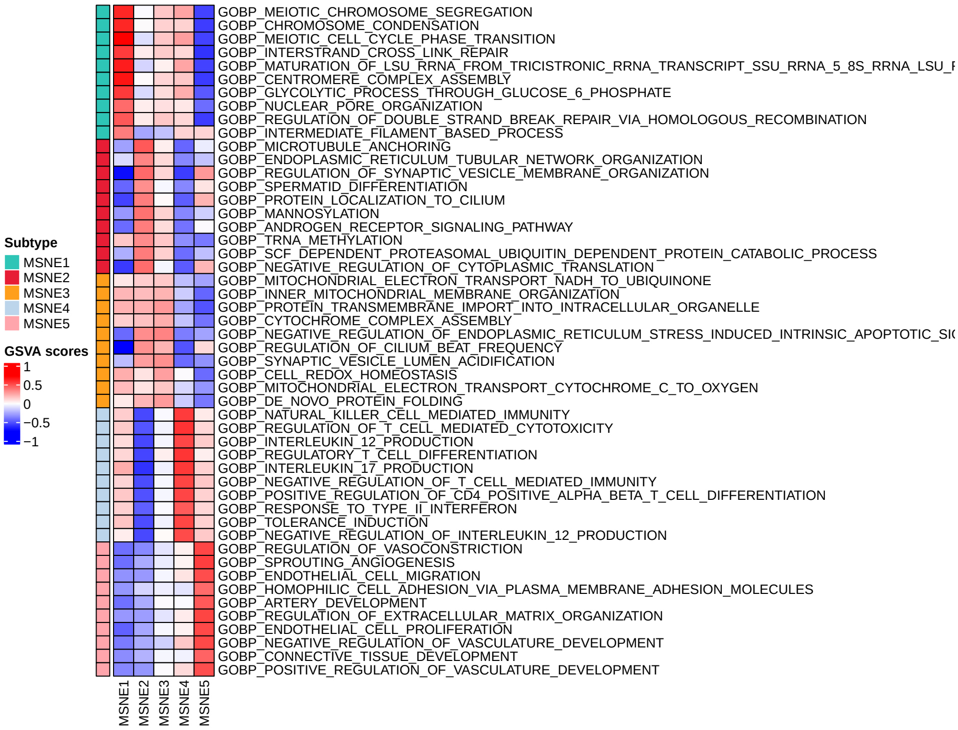

**Figure 14. Up-regulated pathways in MSNE clusters.**

##### 8.1.3 RWR-F

RWR-F results are presented in Figure 15. RWR-F distinguishes two subtypes which are not as pure in ER status composition as results from other similarity network methods Figure 15a. Results are mainly driven by CNV data as shown in the corresponding PCA plot and the FGA plot. Finally, up-regulated pathways are presented in Figure 15d and the ranking of the top 1000 features is shown in Figure 15e. CNV, SNP, RNA-seq and methylation features are amongst the highest ranking features.

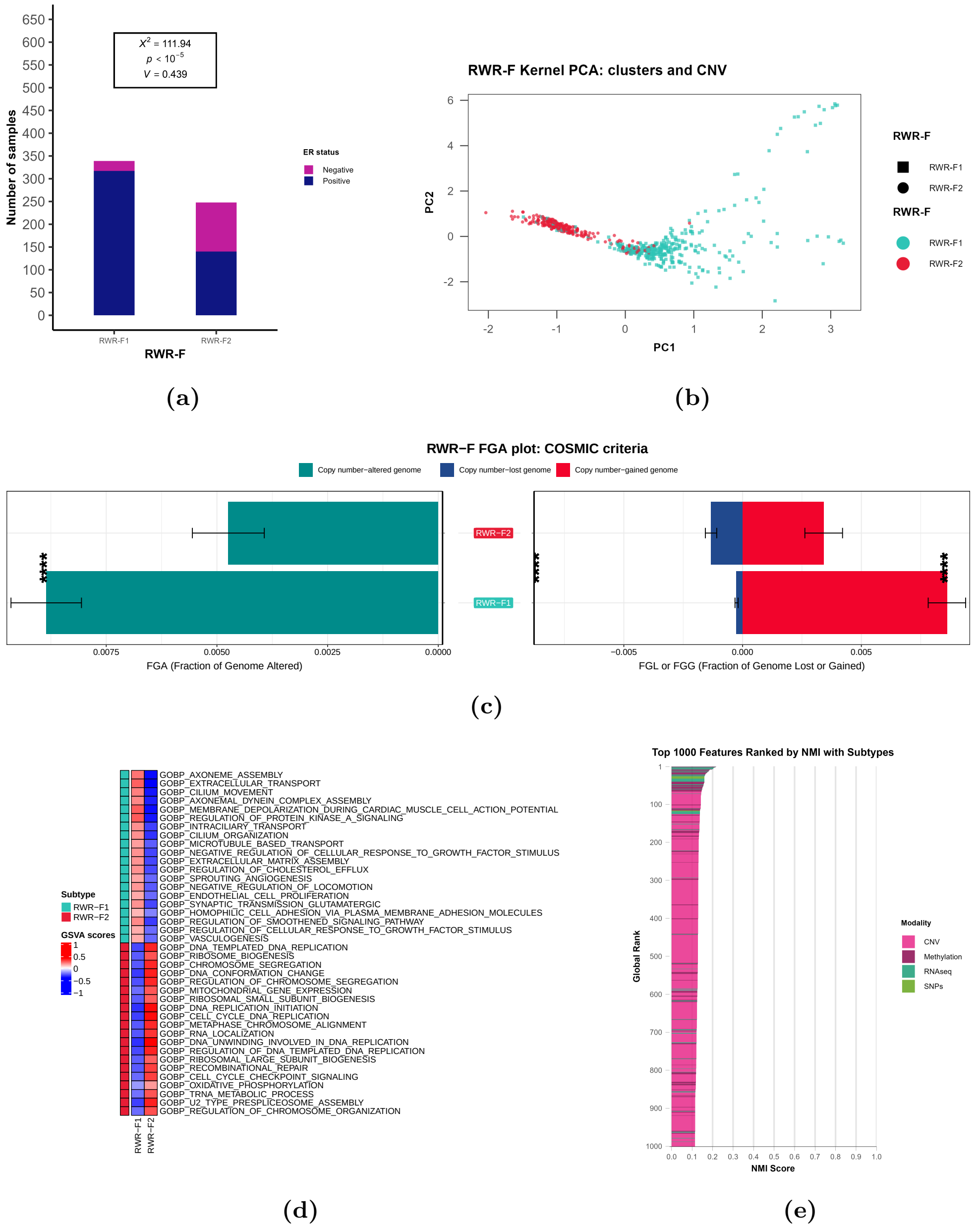

**Figure 15. RWR-F results:** (a) ER status bar chart, (b) CNV data PCA, (c) FGA plot, (d) up-regulated pathways and (e) top 1000 features ranking.

#### 8.2 Multiple-kernel learning methods

##### 8.2.1 CIMLR

Figure 16 presents results for CIMLR clusters: ER status bar chart (Figure 16a), pathways (Figure 16b), FGA (Figure 16c) and oncoprint (Figure 16d).

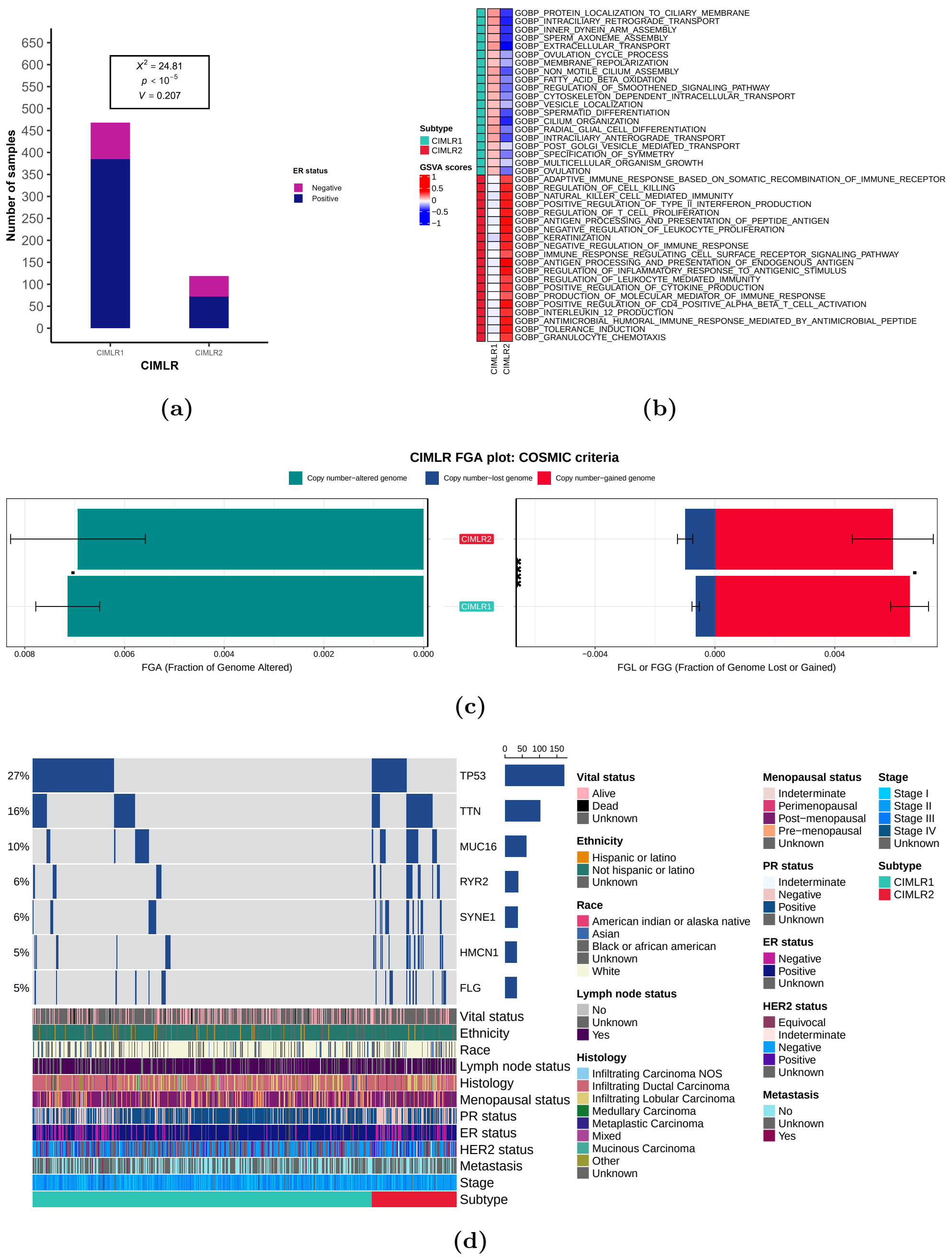

**Figure 16. CIMLR results:** (a) ER status bar chart, (b) up-regulated pathways, (c) FGA plot and (d) oncoprint.

##### 8.2.2 KLIC

The KLIC multi-omic heatmap (Figure 17), ER status bar chart (Figure 18a) and silhouette scores (Figure 18b), demonstrate how the optimal partitions for KLIC are imbalanced and show little plausibility.

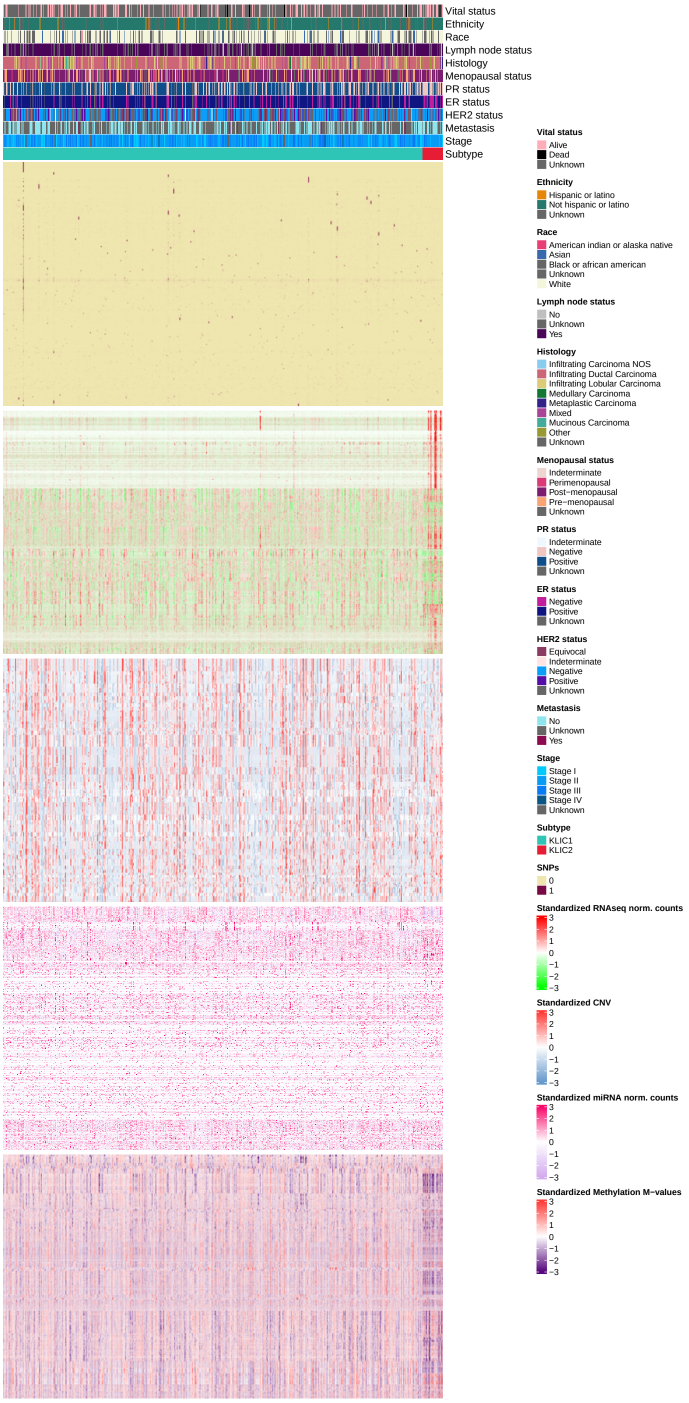

**Figure 17. KLIC multi-omic heatmap.** Distinct profiles can be seen in the RNA-seq and methylation sub-heatmaps.

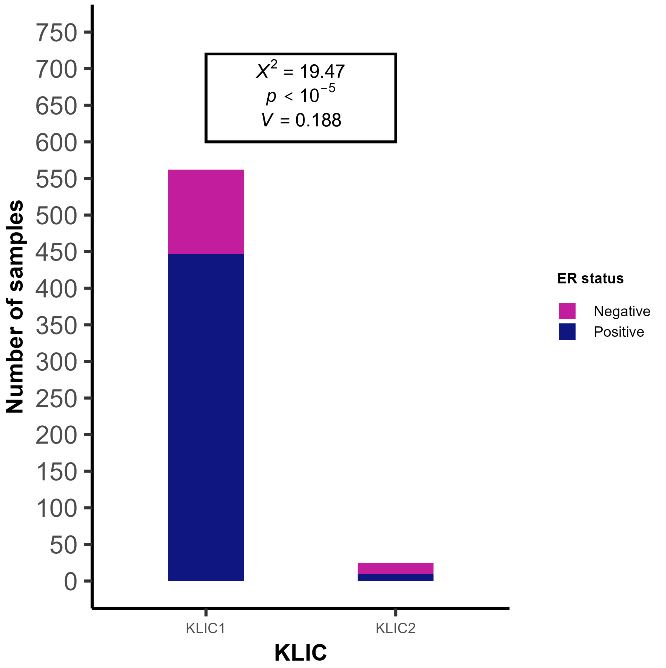

**(a)**

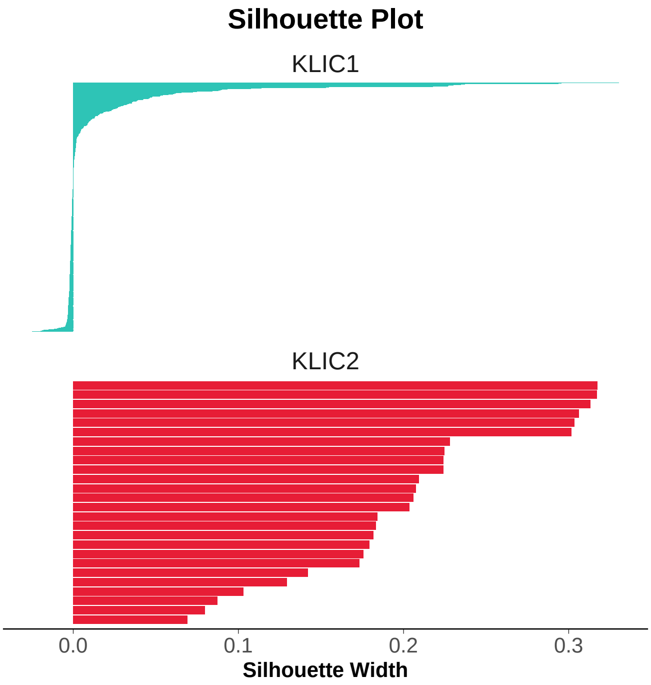

**(b)**

**Figure 18. KLIC results:** (a) ER status bar chart and (b) silhouette scores.

##### 8.2.3 wMKL

wMKL results are presented in three Figures: i) multi-omic heatmap (Figure 20), ii) a composite plot showing the ER (Figure 20a) and HER2 (Figure 20b) status bar charts, the FGA patterns across clusters (Figure 20c), the oncoprint (Figure 20d), the final similarity kernel PCA (Figure 20e) and the CNV PCA plot (Figure 20f), and iii) a plot of the representative up-regulated pathways in each cluster (Figure 21).

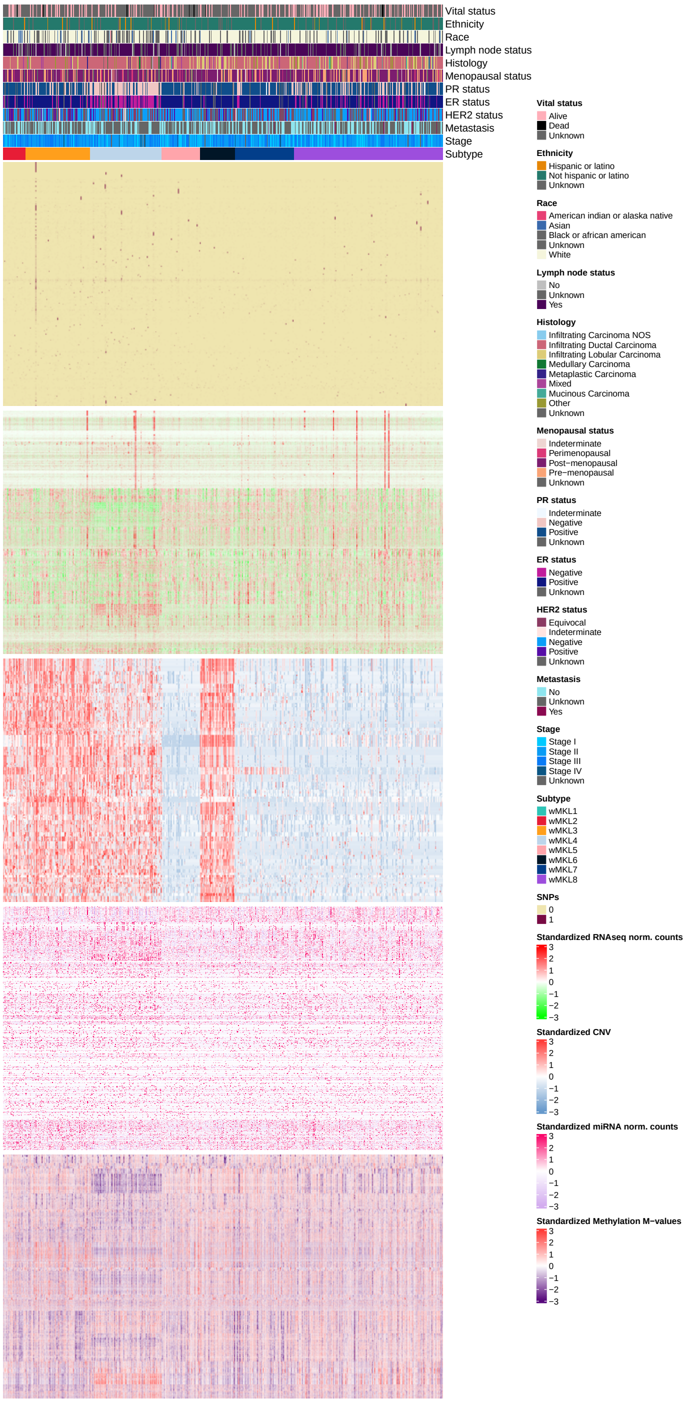

**Figure 19. wMKL multi-omic heatmap.**

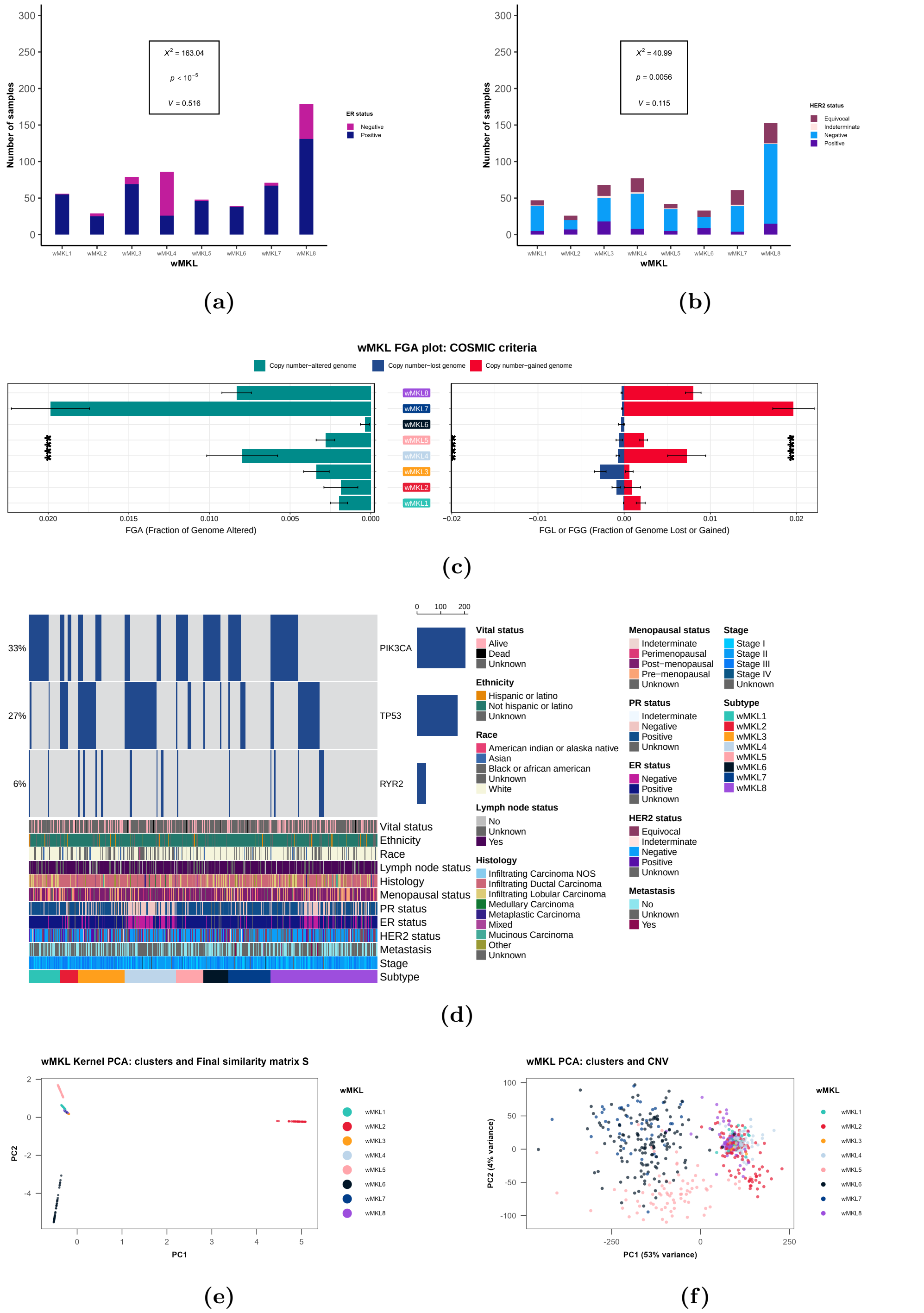

**Figure 20. wMKL results:** (a) ER status bar chart, (b) HER2 status bar chart, (c) FGA plot, (d) oncoprint, (e) final similarity kernel PCA and (f) CNV data PCA.

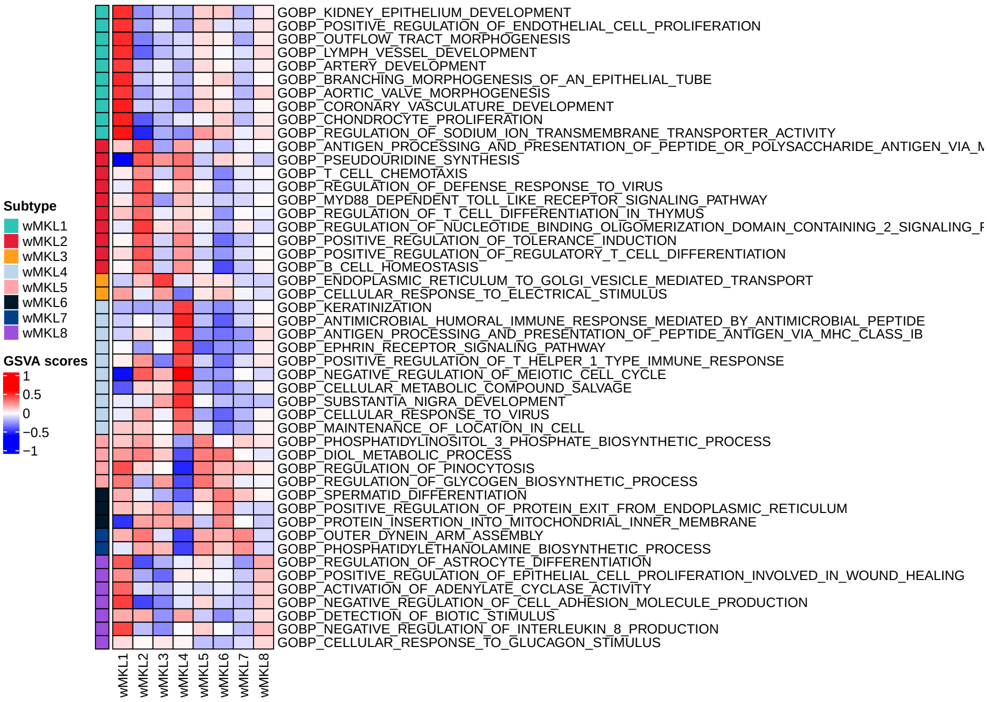

**Figure 21. Up-regulated pathways in wMKL clusters.**

#### 8.3 Matrix Factorisation methods

##### 8.3.1 LRAcluster

LRAcluster results are summarised in Figures 22, 23. Figure 22 is a composite plot showing the ER (Figure 22a) and HER2 (Figure 22b) status bar charts, the FGA patterns across clusters (Figure 22c), the incremental increase in explained variance (Figure 22d) and the oncoprint (Figure 22e), whereas the representative up-regulated pathways are shown in Figure 23.

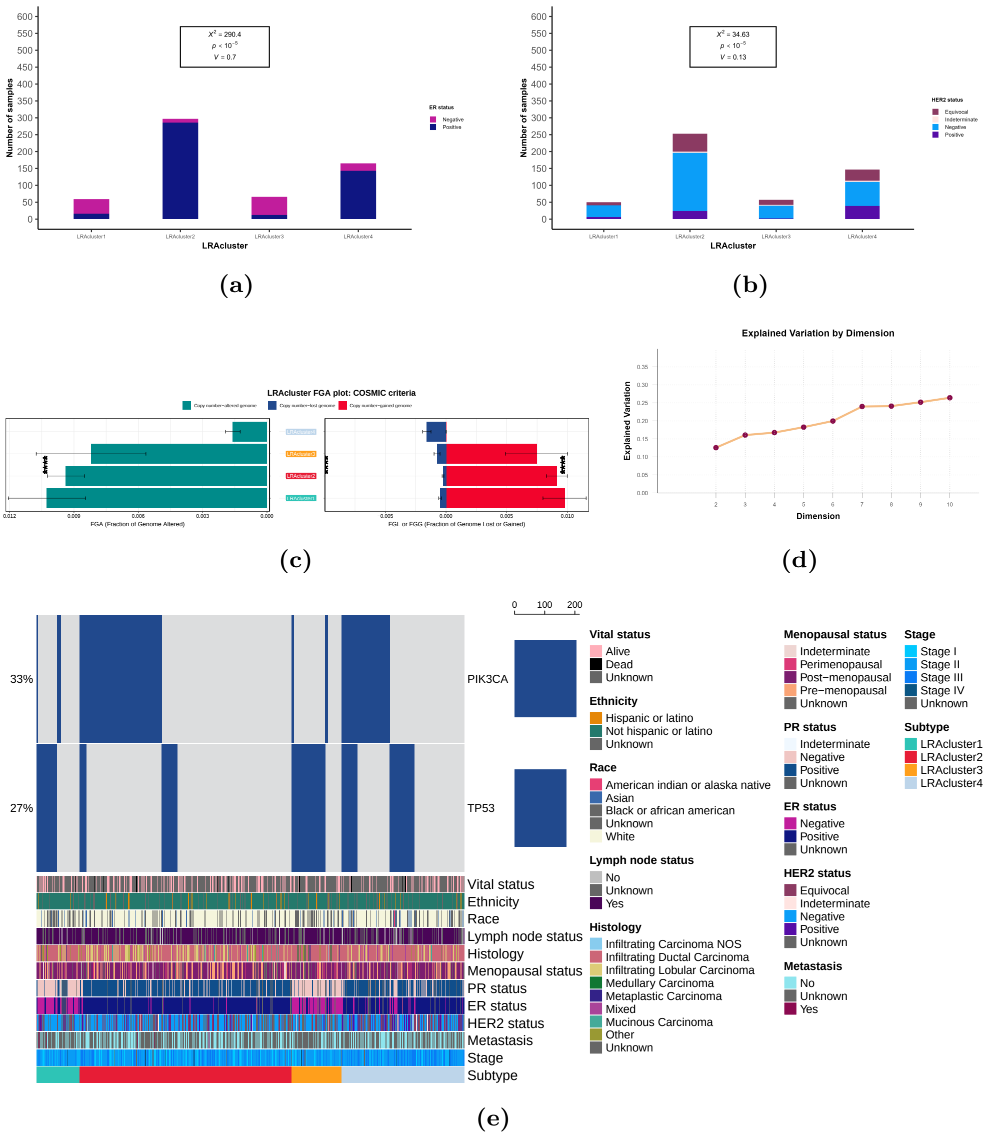

**Figure 22. LRAcluster results:** (a) ER status bar chart, (b) HER2 status bar chart, (c) FGA plot and (d) oncoprint

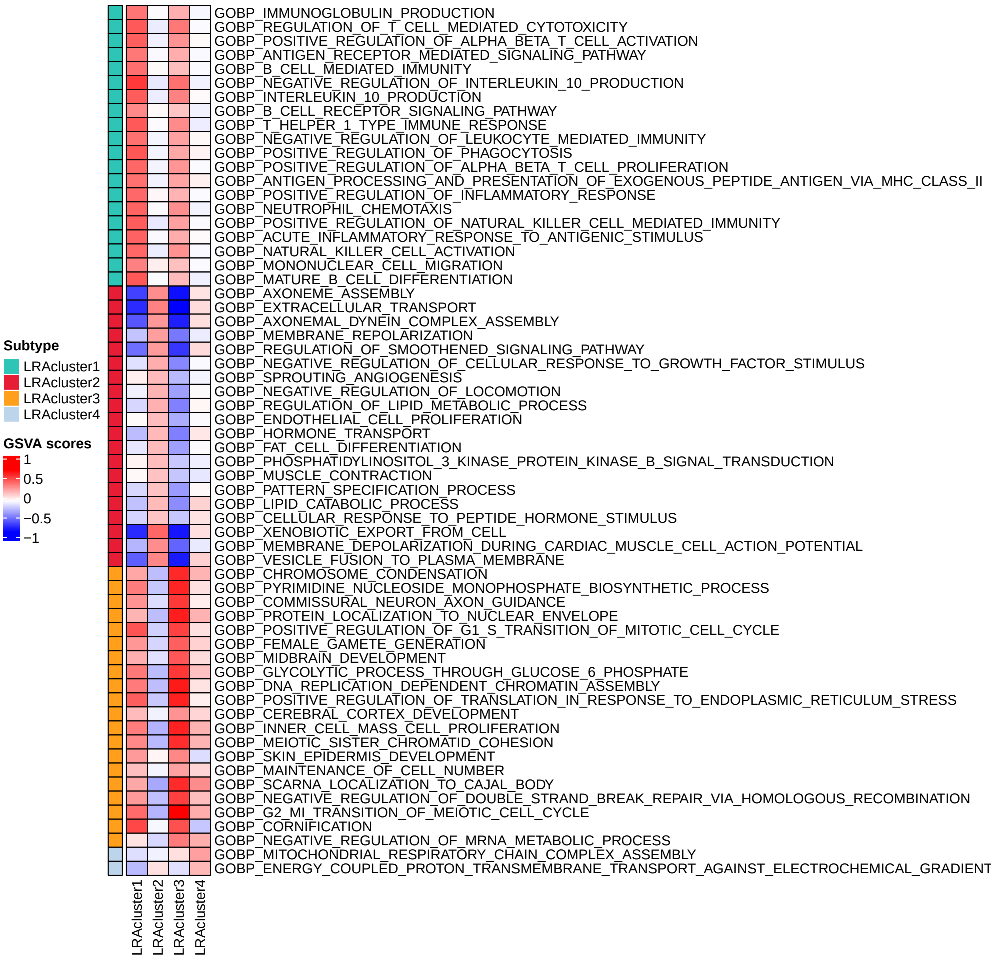

**Figure 23. Up-regulated pathways in LRAcluster clusters.**

##### 8.3.2 MFA

MFA results are presented below. Figure 24 presents the ER status bar chart, the silhouette scores, the FGA distributions across clusters and the oncoprint. Figure 26 shows the up-regulated pathways in the three MFA clusters. Figure 25 shows the low-dimensional projection of the five modalities and the distribution of the three clusters within them, in addition to a distance matrix obtained from the embeddings of the samples in the $r_{\text{opt}} = 11$ factors. Feature contribution plots (Figure 27), modality contribution plots (Figure 28), ER (Figure 29) and HER2 (Figure 30) distribution plots follow.

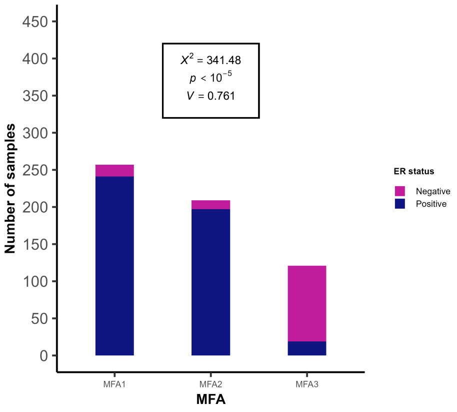

(a)

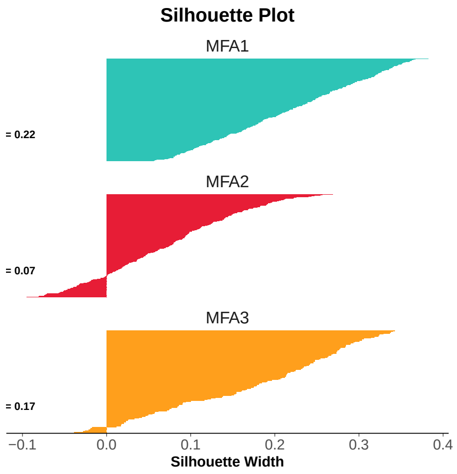

(b)

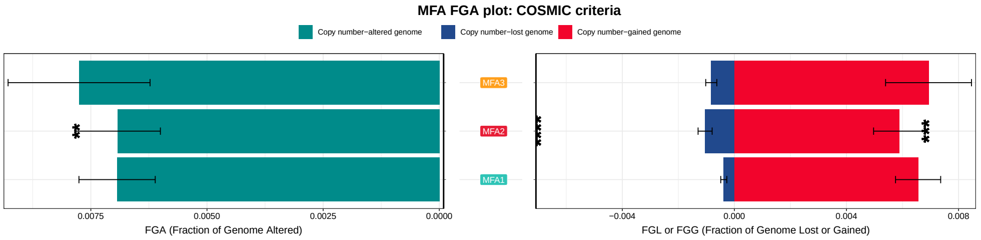

(c)

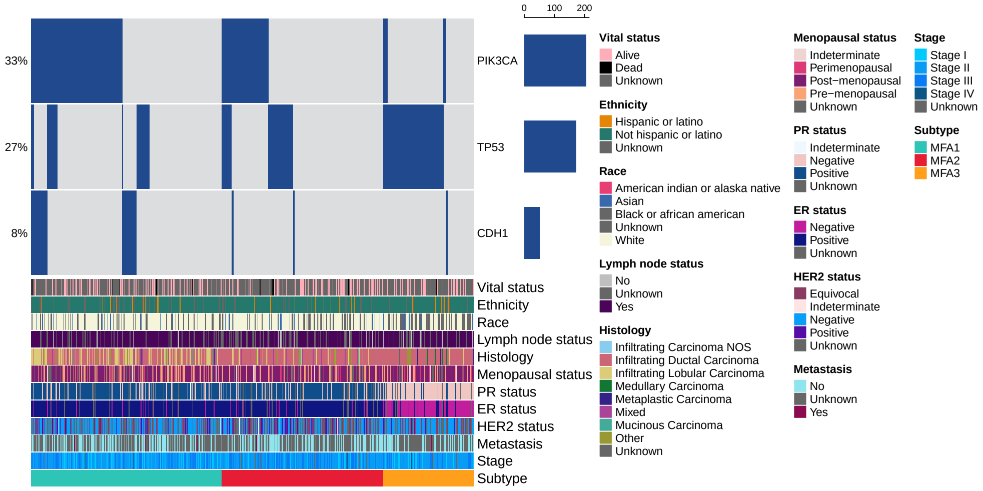

(d)

**Figure 24. MFA results:** (a) ER status bar chart, (b) HER2 status bar chart, (c) FGA plot and (d) oncoprint

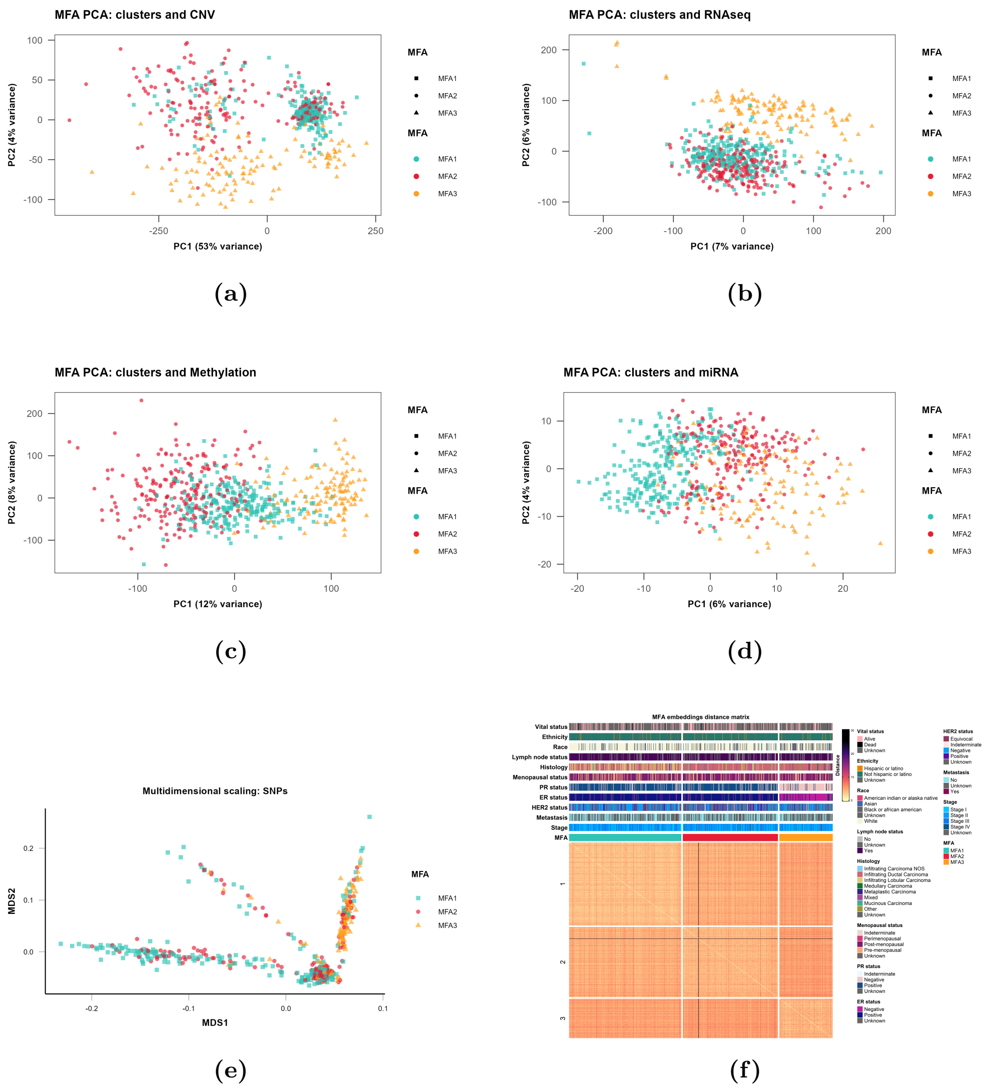

**Figure 25. MFA dimensionality reduction plots and embeddings distance matrix.** (a) CNV data PCA, (b) RNA-seq data PCA, (c) methylation data PCA, (d) miRNA data PCA, (e) SNP MDS plot and (f) embeddings distance matrix.

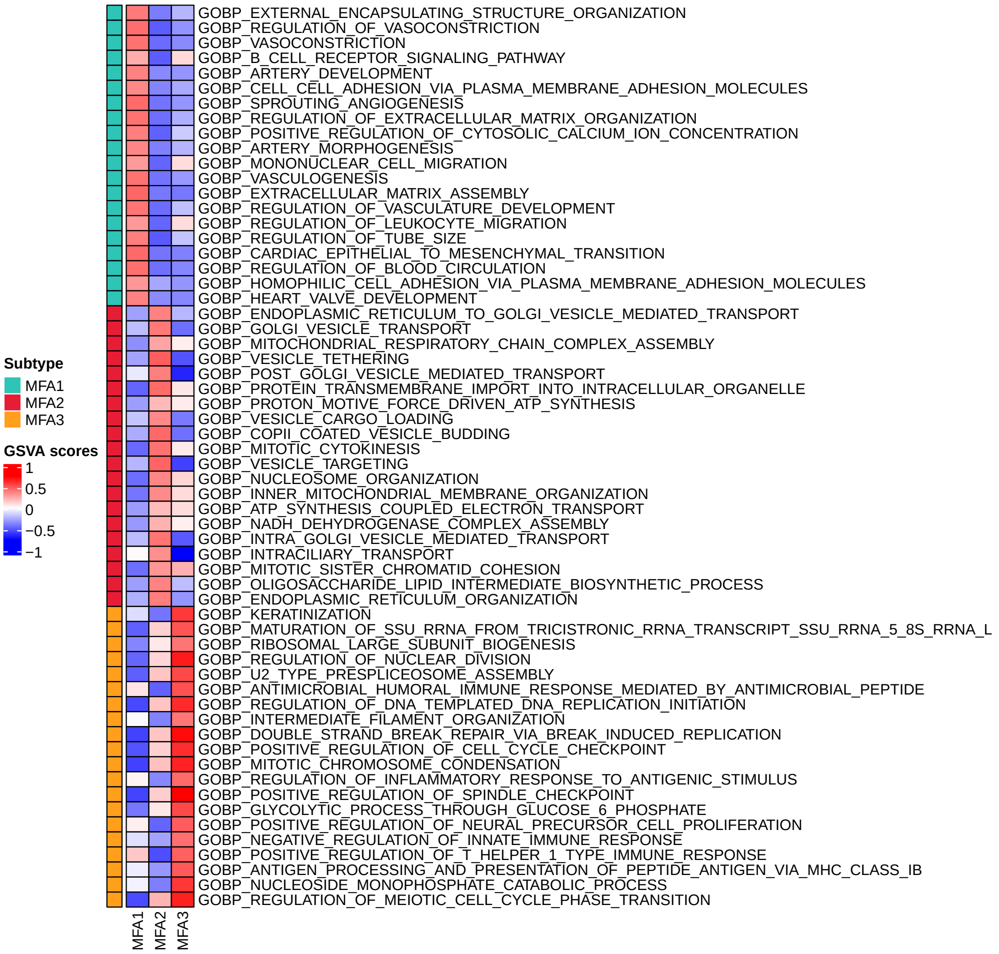

**Figure 26. Up-regulated pathways in MFA clusters.**

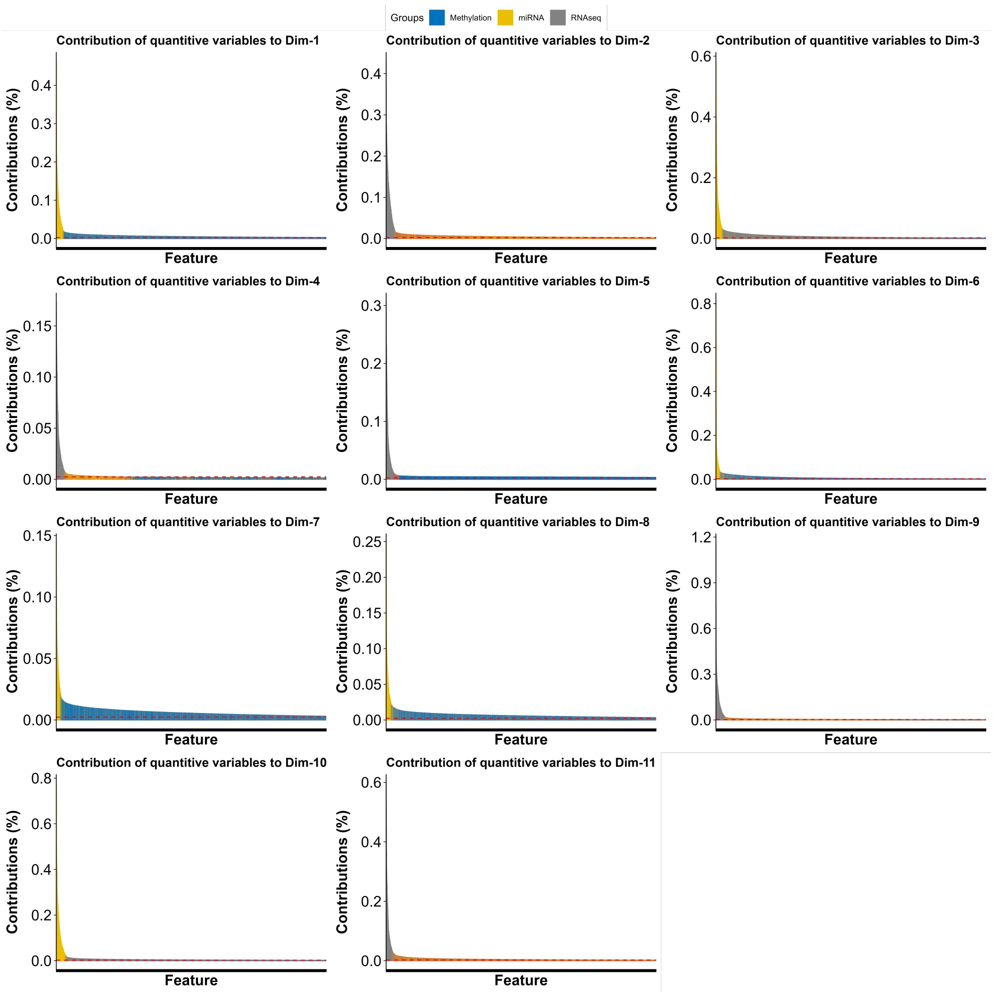

**Figure 27. Feature contributions (top 10000 features) across all 11 latent factors.**

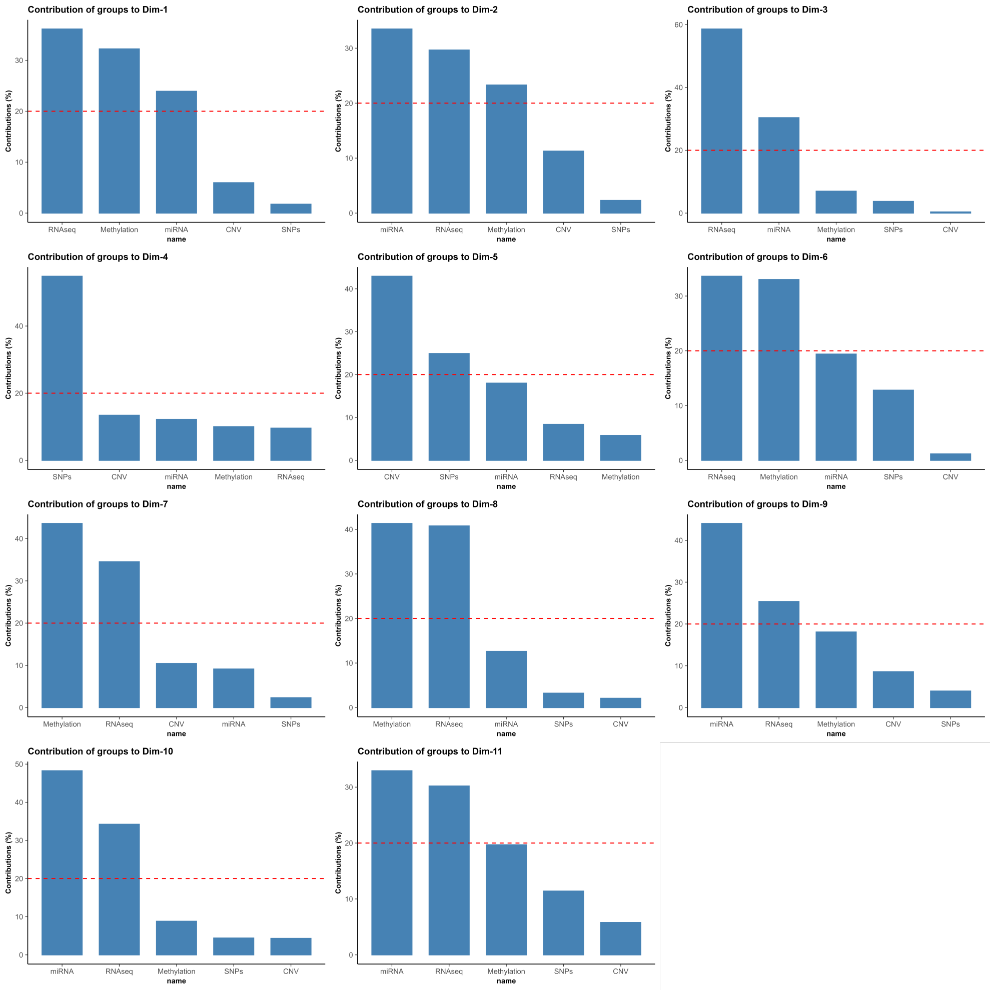

**Figure 28. Modality contributions across all 11 latent factors.**

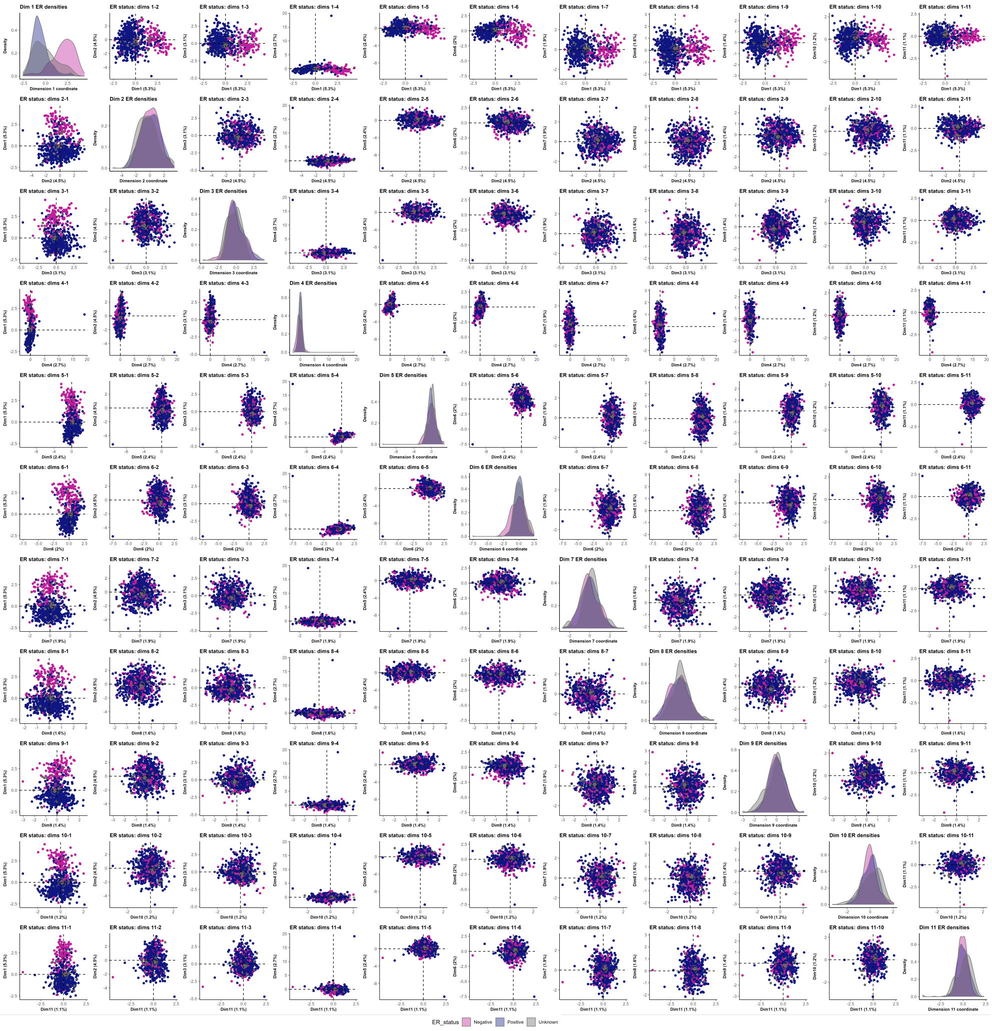

**Figure 29. ER status distribution across all 11 latent factors.**

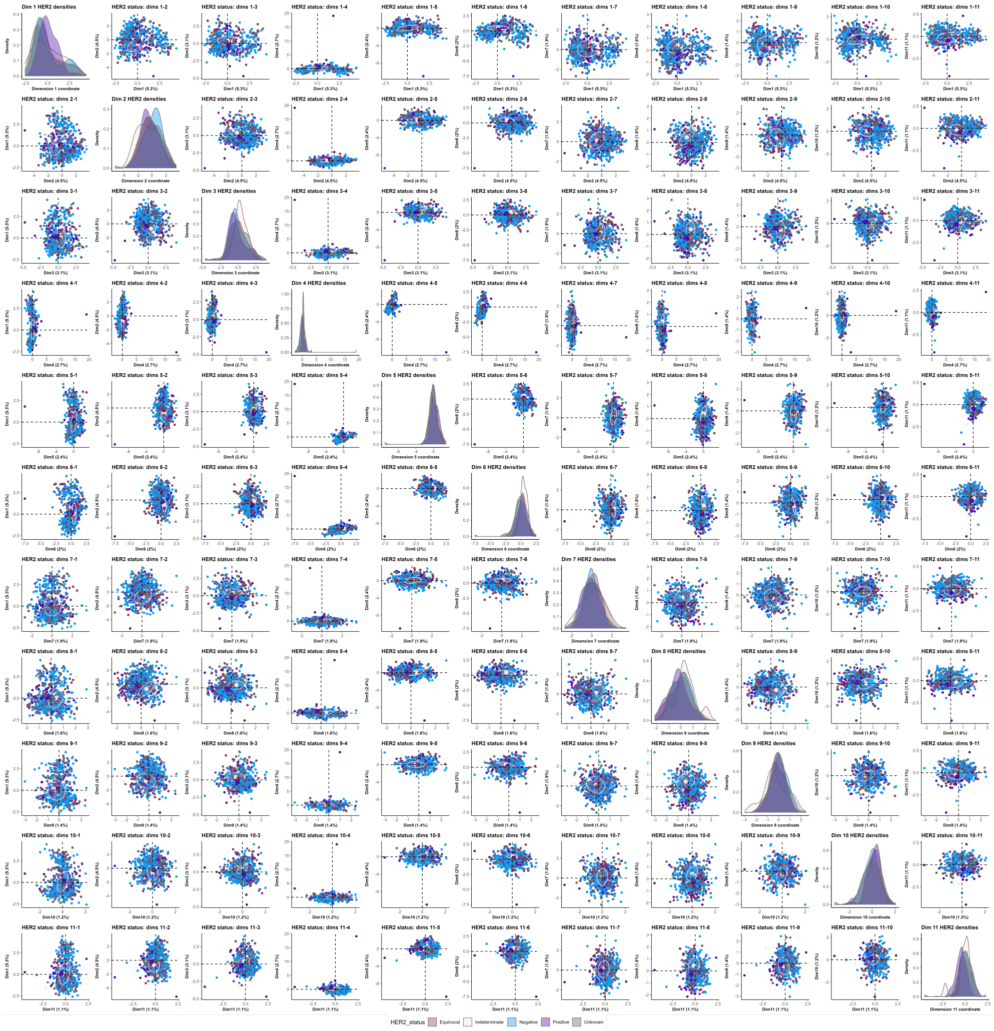

**Figure 30. HER2 status distribution across all 11 latent factors.**

##### 8.3.3 MOFA

MOFA results are presented below. Figure 31 demonstrates the increments in cumulative explained variance for the tests numbers of candidate factors. Figure 32 presents the ER status bar chart, the low-dimensional CNV space, the FGA distributions across clusters and the oncoprint. Figure 33 shows the representative up-regulated pathways in the three MOFA clusters. Feature contribution plots (Figure 34), ER (Figure 35) and HER2 (Figure 36) distribution plots follow.

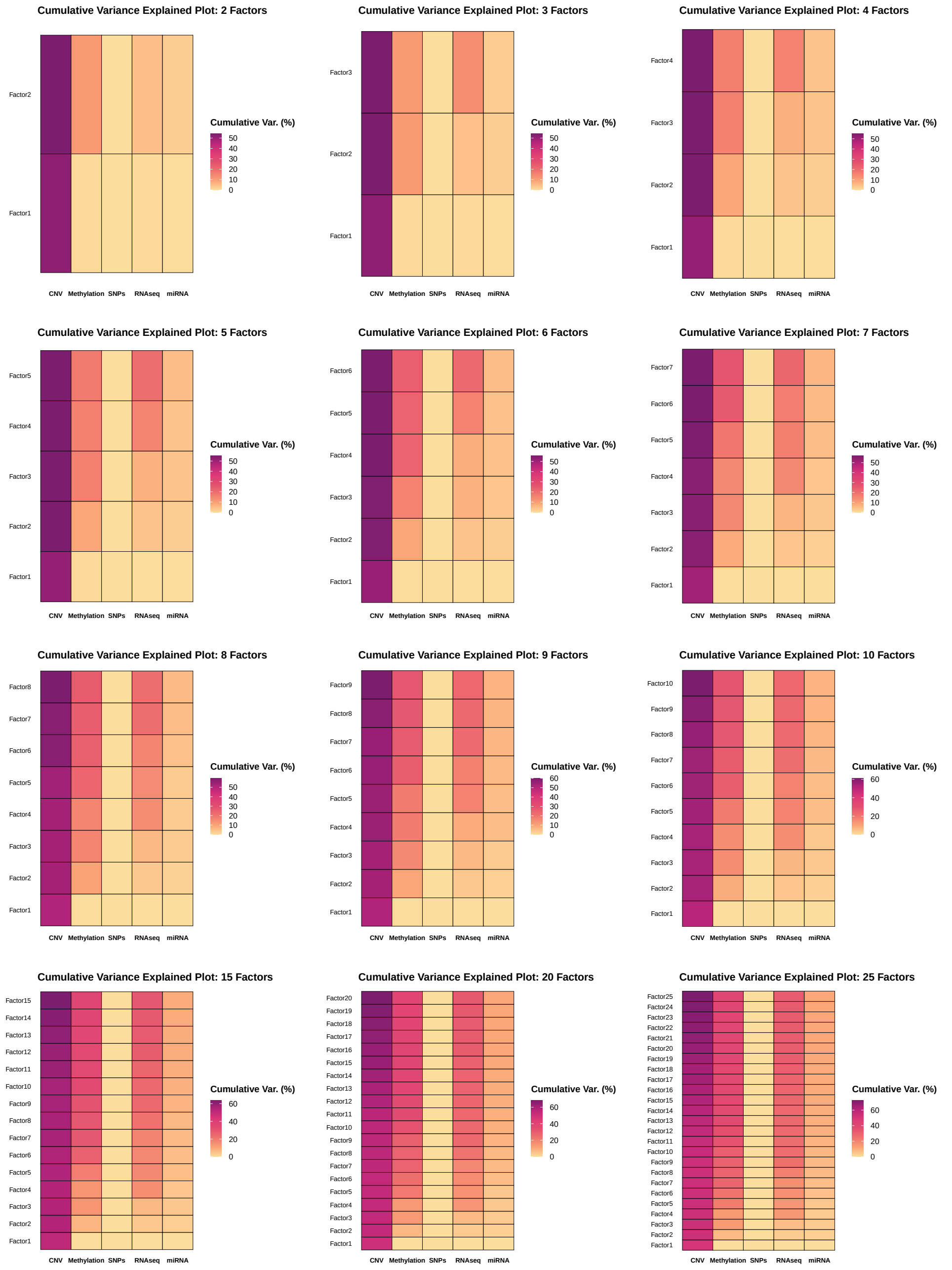

**Figure 31. Cumulative explained variance across all candidate factors.**

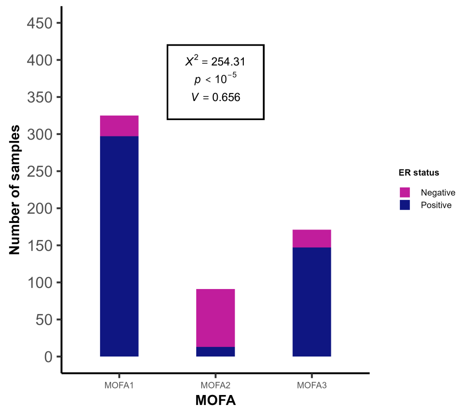

**(a)**

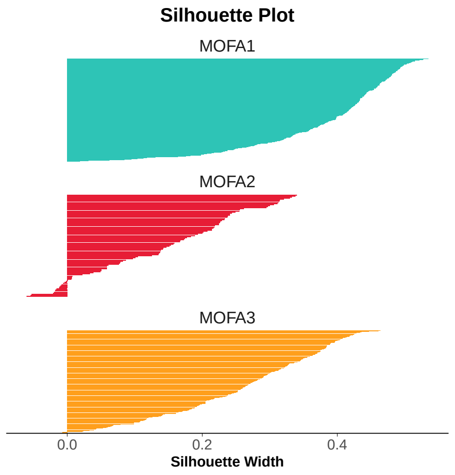

**(b)**

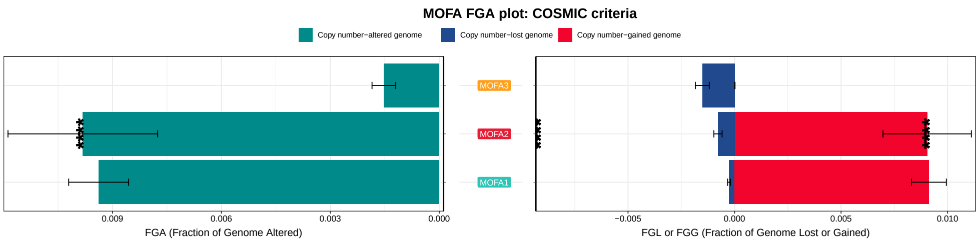

**(c)**

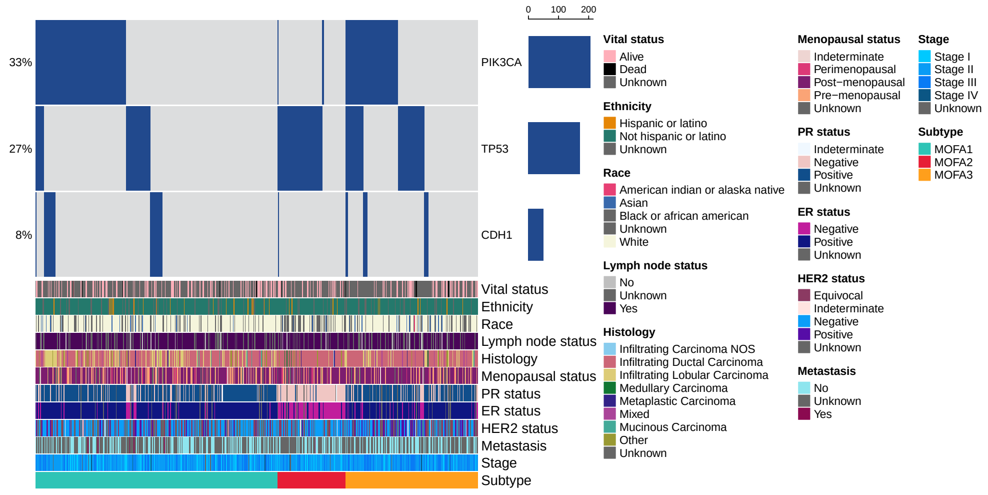

**(d)**

**Figure 32. MOFA results:** (a) ER status bar chart, (b) HER2 status bar chart, (c) FGA plot and (d) oncoprint

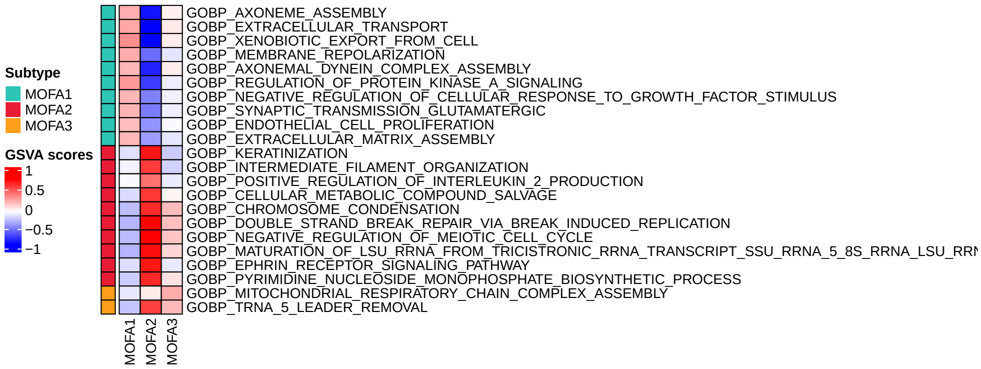

**Figure 33. Up-regulated pathways in MOFA clusters.**

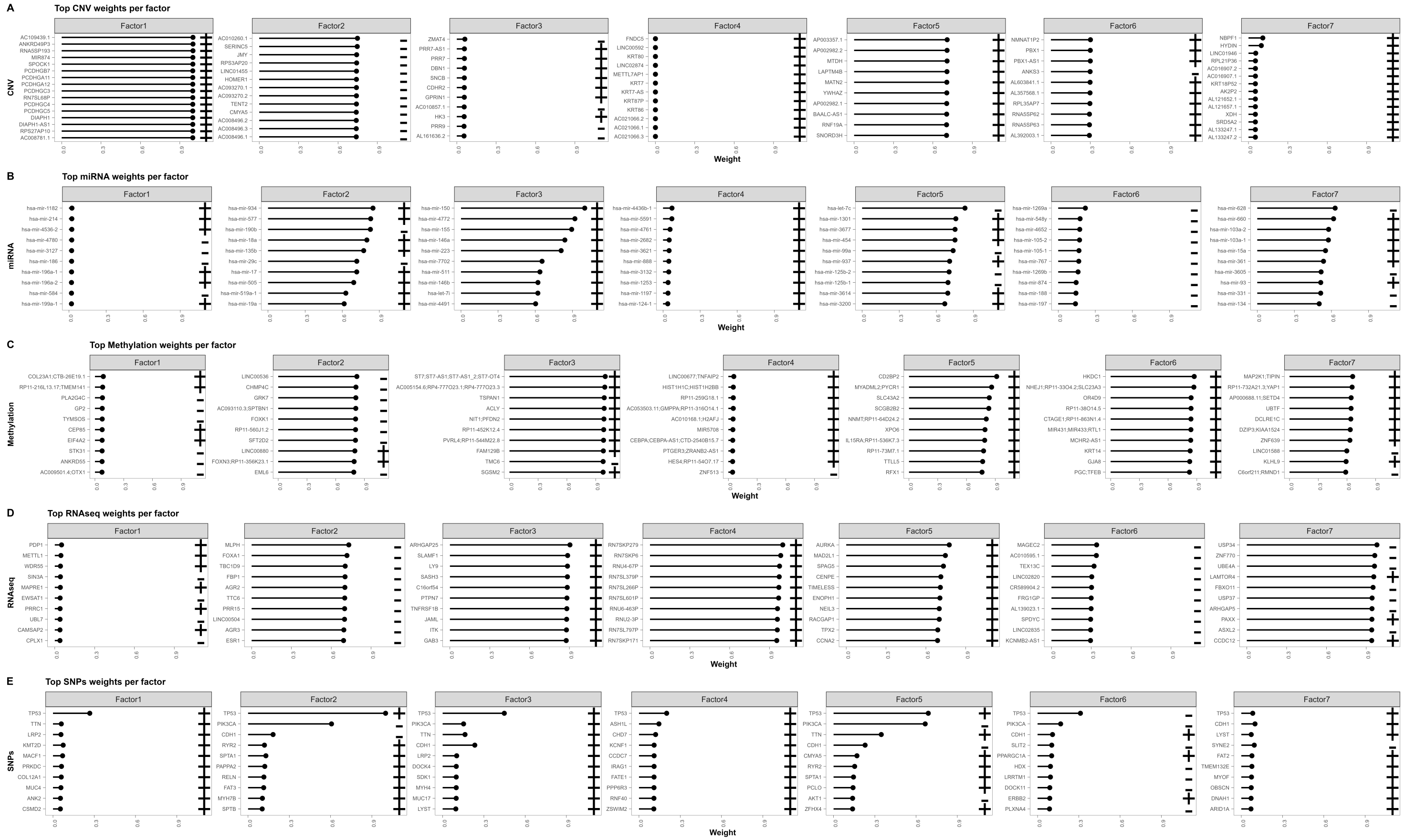

**Figure 34. Feature contributions (top 10 features) across all 7 latent factors.**

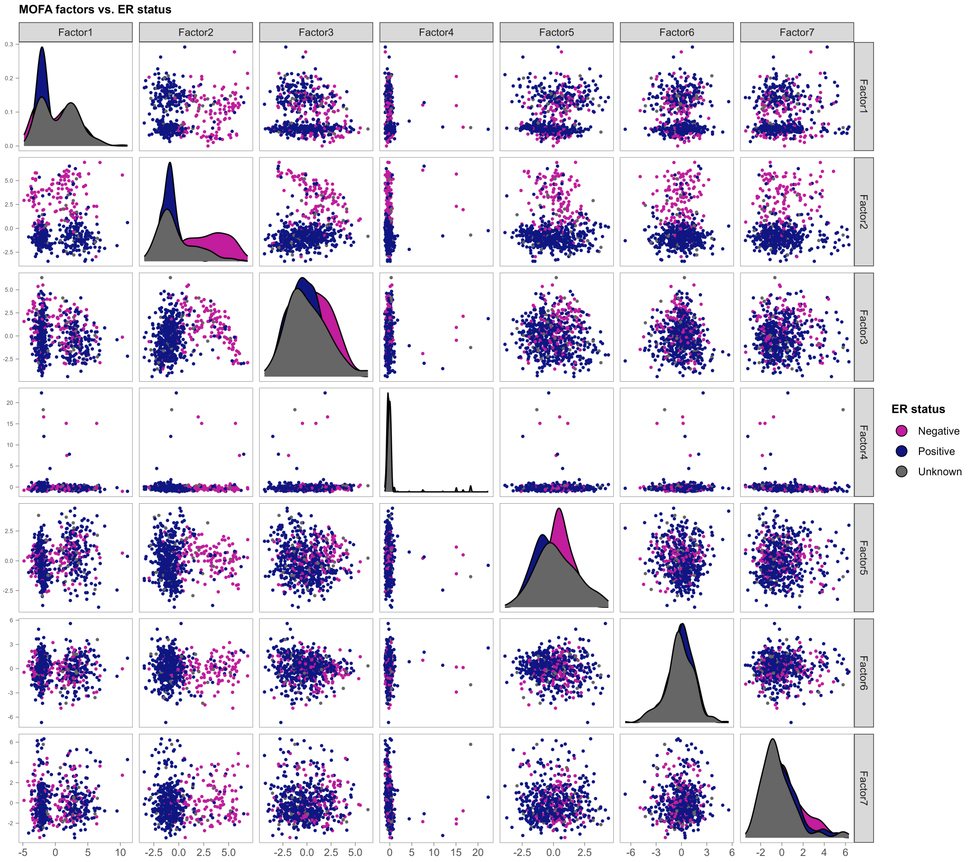

**Figure 35. ER status distribution across all 7 latent factors.**

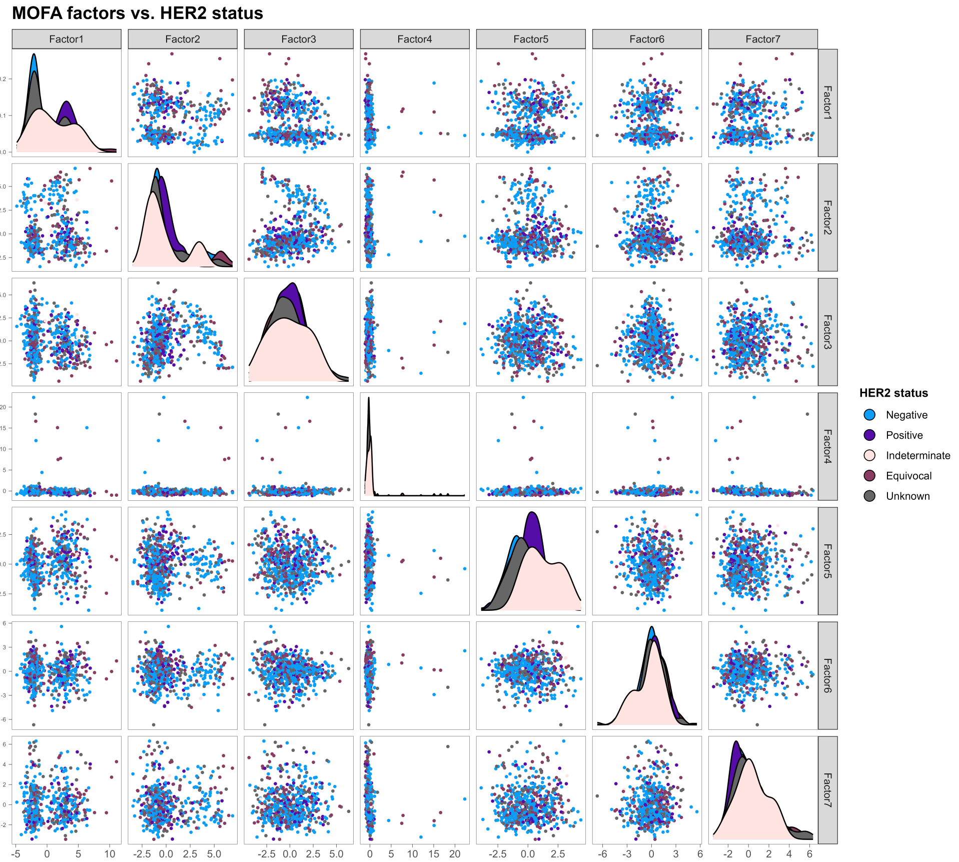

**Figure 36. HER2 status distribution across all 7 latent factors.**

#### 8.4 Bayesian methods

##### 8.4.1 iClusterBayes

iClusterBayes results are presented below. Figure 37 presents the ER status bar chart, the low-dimensional CNV space, the FGA distributions across clusters and the oncoprint. Figure 37c shows the representative up-regulated pathways in the three iClusterBayes clusters.

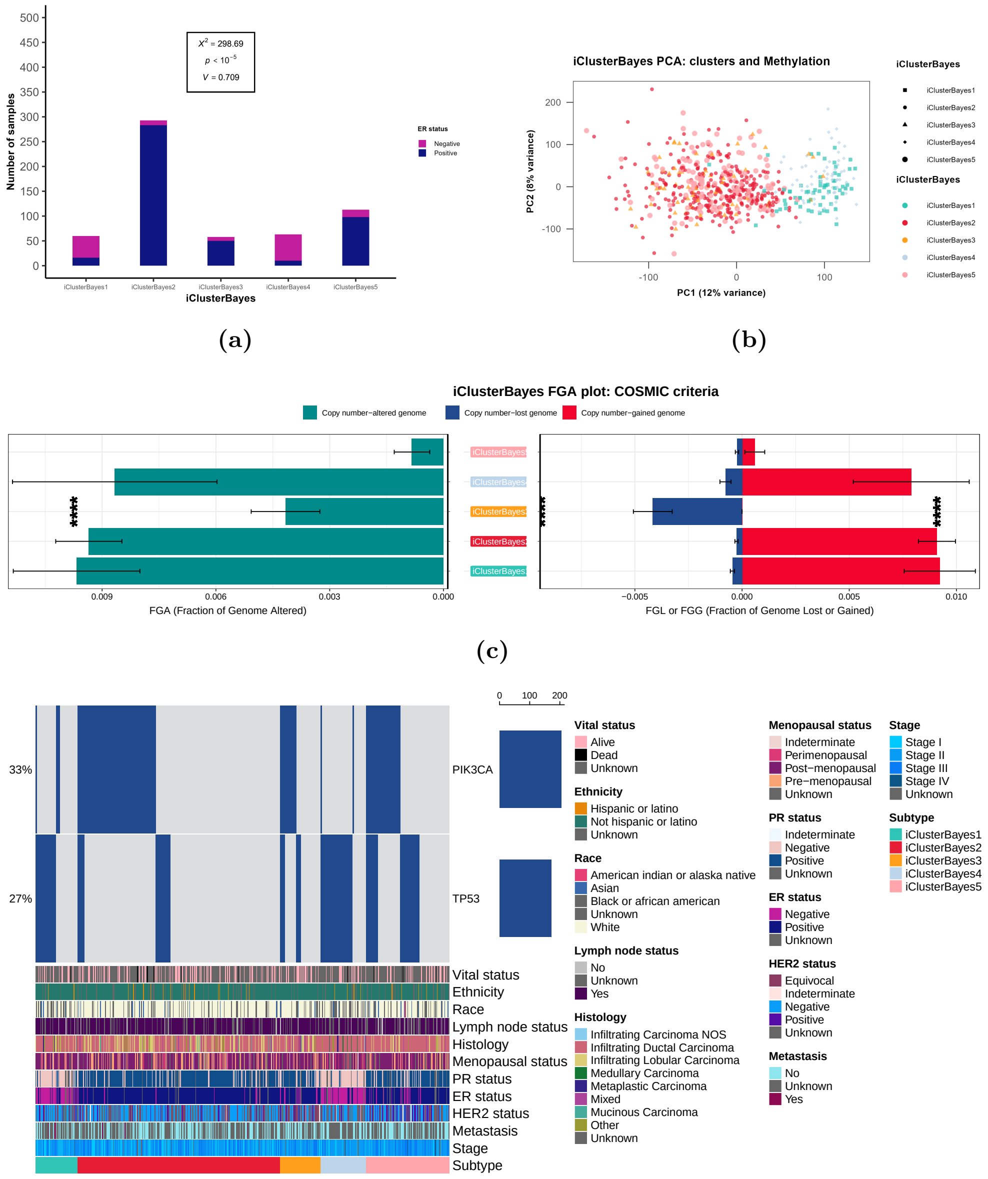

(d)

**Figure 37. iClusterBayes results:** (a) ER status bar chart, (b) Methylation data PCA, (c) FGA plot and (d) oncoprint

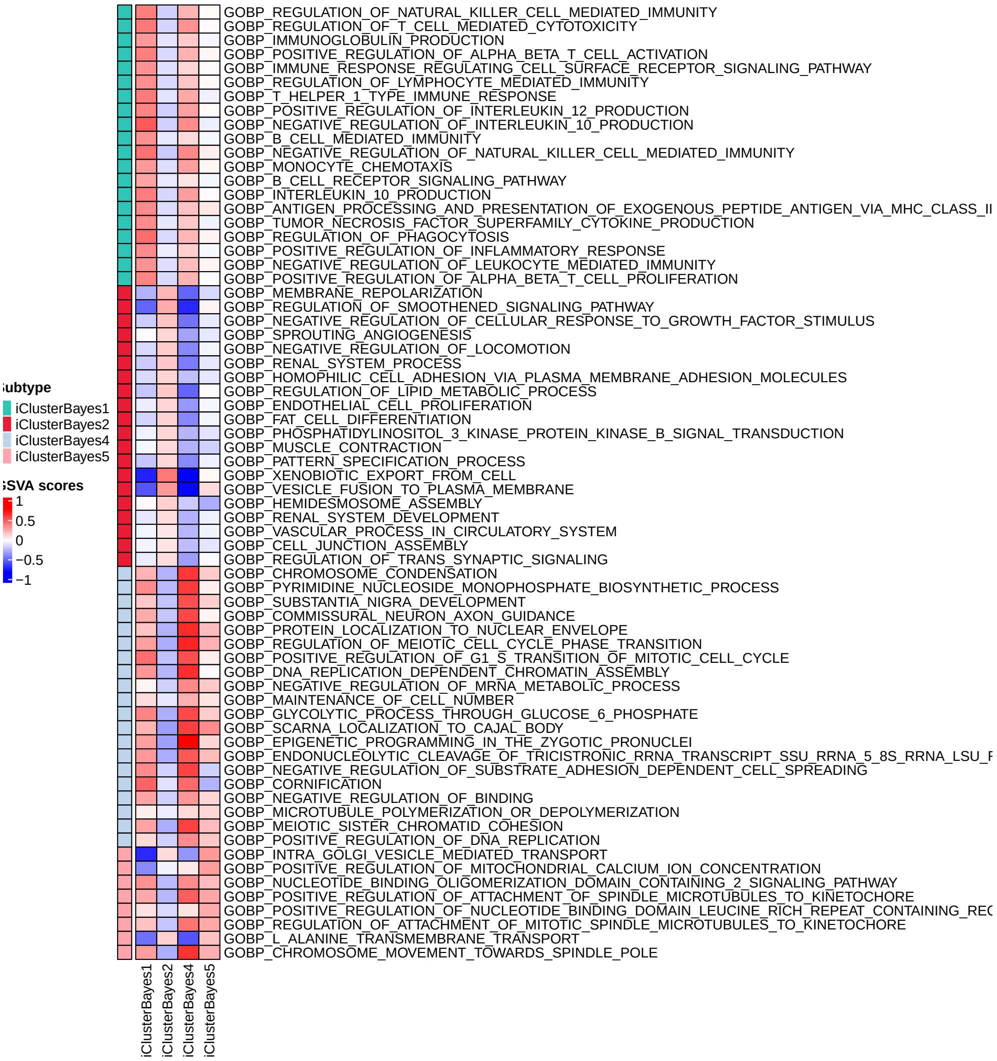

**Figure 38. Up-regulated pathways in iClusterBayes clusters.**

#### 8.5 Graph-based methods

##### 8.5.1 MONET

MONET results are presented below. Figure 39 shows the bar charts of MONET clusters and ER status, HER2 status, lymph node status and tumour stage. Figure 40 presents the FGA distributions across clusters and the oncoprint. The multi-omic heatmap of MONET is shown in Figure 41, whereas the kernel PCA plots for the adjacencies within individual modalities and the average multi-omic adjacency are shown in Figure 42. Figure 43 shows the representative up-regulated pathways in the three MONET clusters. Figure 44 illustrates the composition of the identified MONET modules with respect to ER and HER2 status.

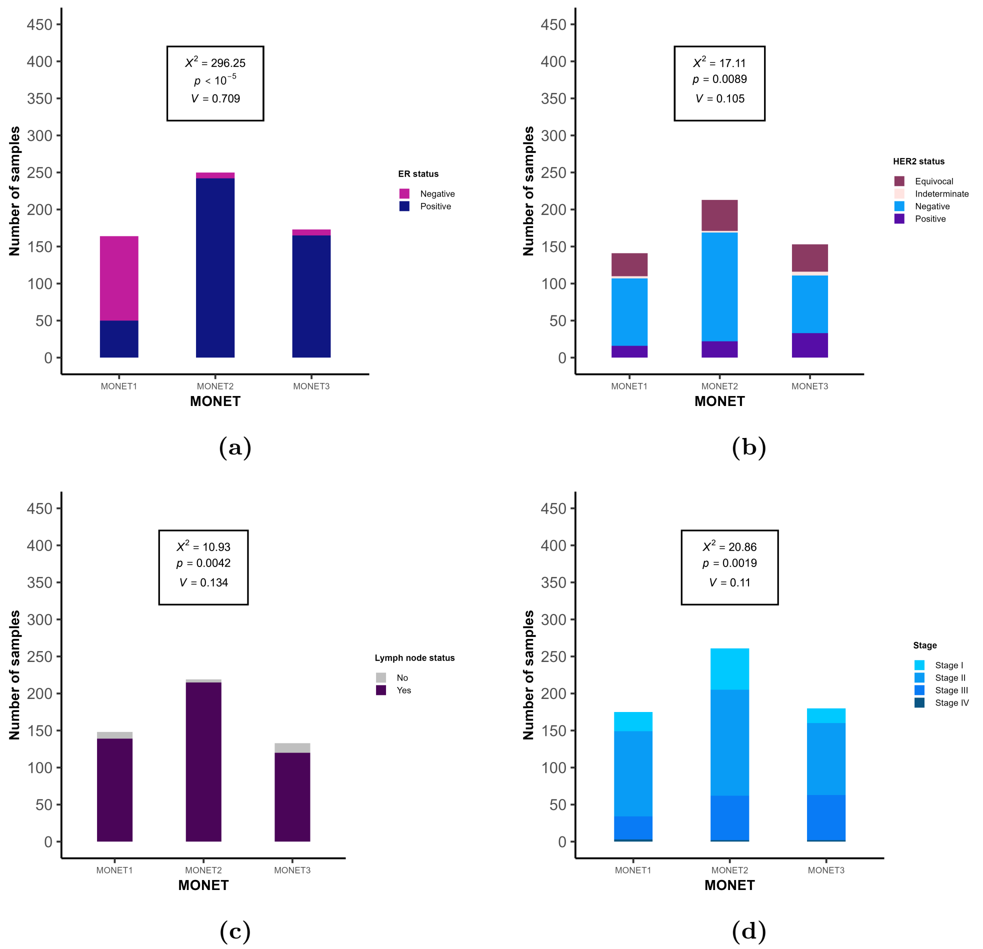

**Figure 39. MONET bar charts:** (a) ER status, (b) HER2 status, (c) Lymph node status and (d) tumour stage.

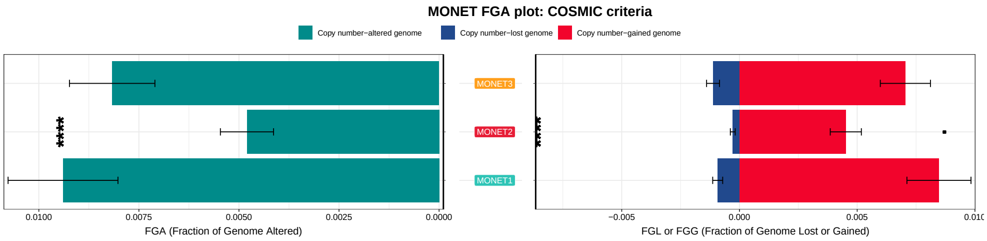

(a)

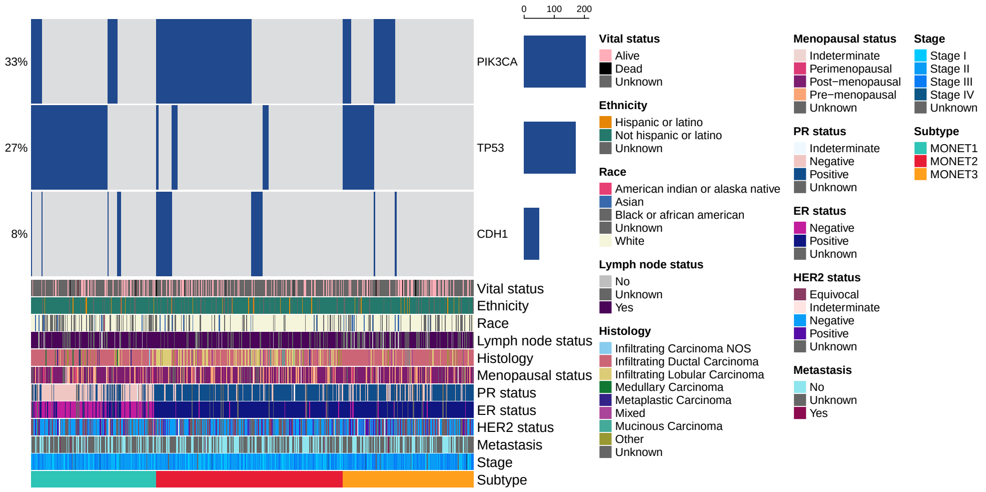

(b)

**Figure 40. MONET results:** (a) FGA plot and (b) oncoprint.

#### 8.6 Ensemble clustering methods

##### 8.6.1 COCA

Although COCA resolves in a two-cluster solution, the partition achieved is significantly associated only with HER2 status ($p = 0.01$). The separation of samples is mainly driven by CNV data. There is significant FGL in COCA2 and significant FGG in COCA1 (Figure 45). DGEA between the two clusters only revealed five genes as significantly differentially expressed, which is a set of small size for any meaningful pathway analysis.

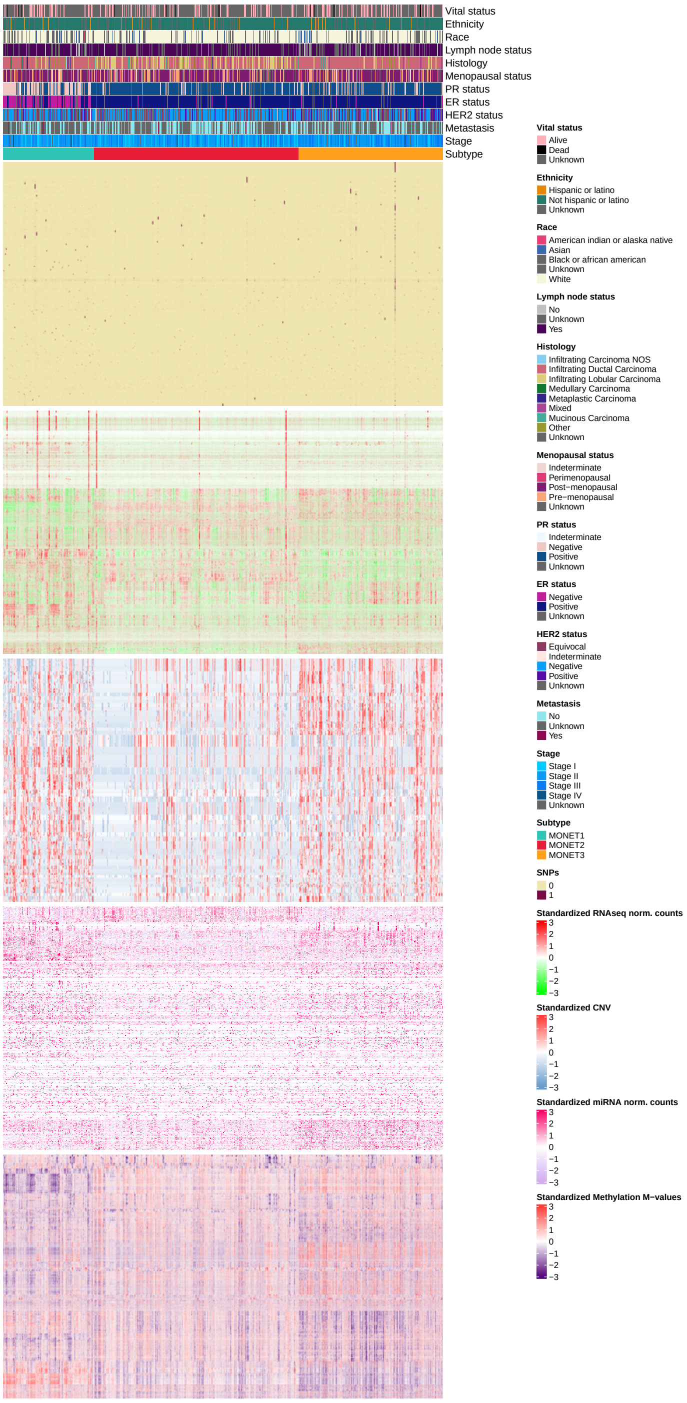

**Figure 41. MONET multi-omic heatmap.**

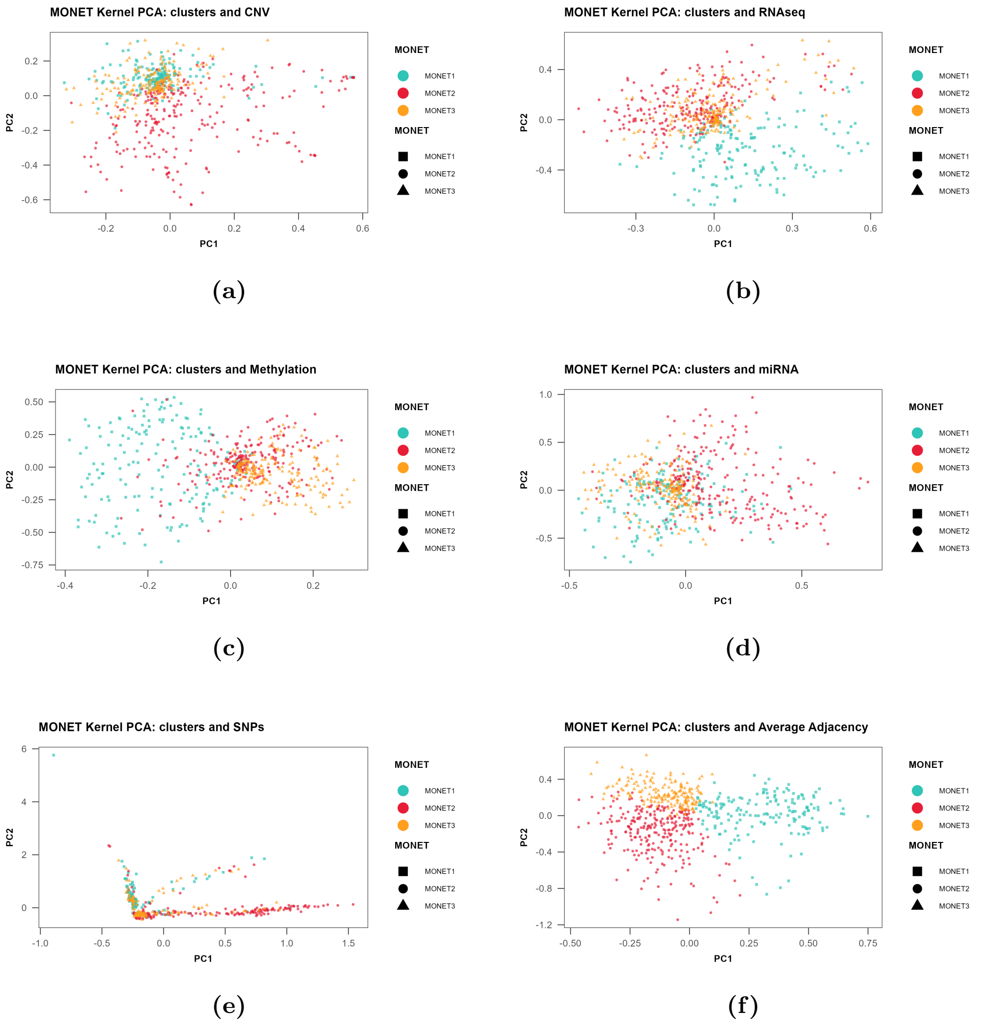

**Figure 42. MONET dimensionality reduction plots and embeddings distance matrix.** (a) CNV adjacency kernel PCA, (b) RNA-seq adjacency kernel PCA, (c) methylation adjacency kernel PCA, (d) miRNA adjacency kernel PCA, (e) SNP adjacency kernel PCA and (f) average adjacency kernel PCA.

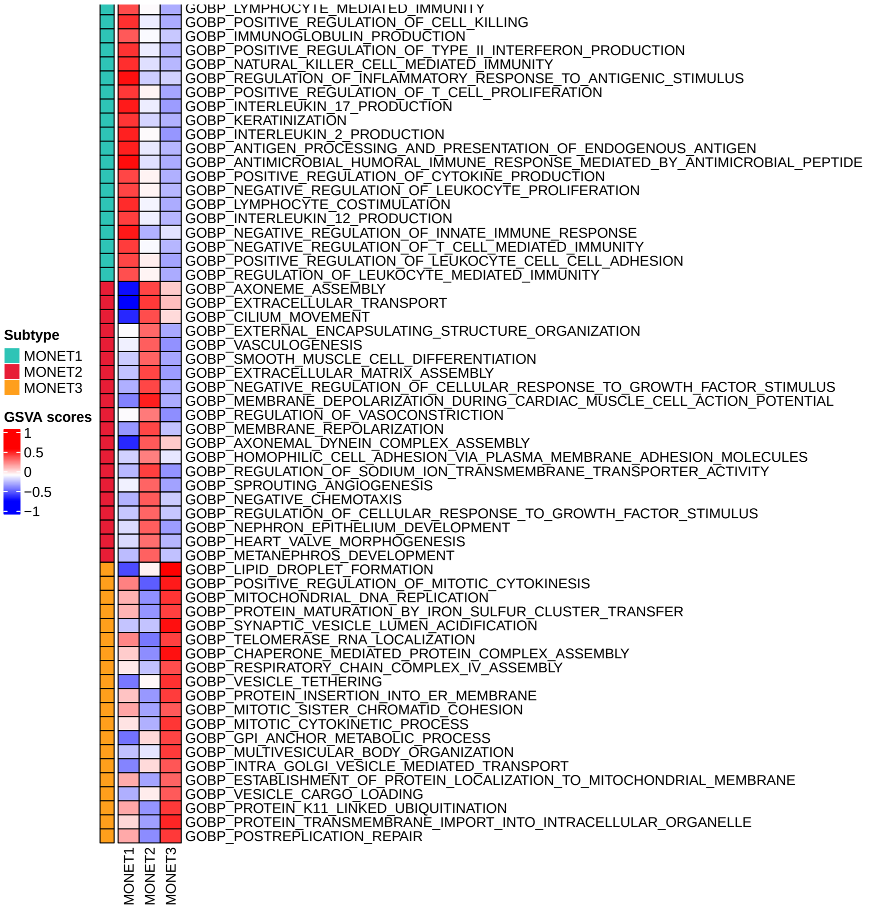

**Figure 43. Up-regulated pathways in MONET clusters.**

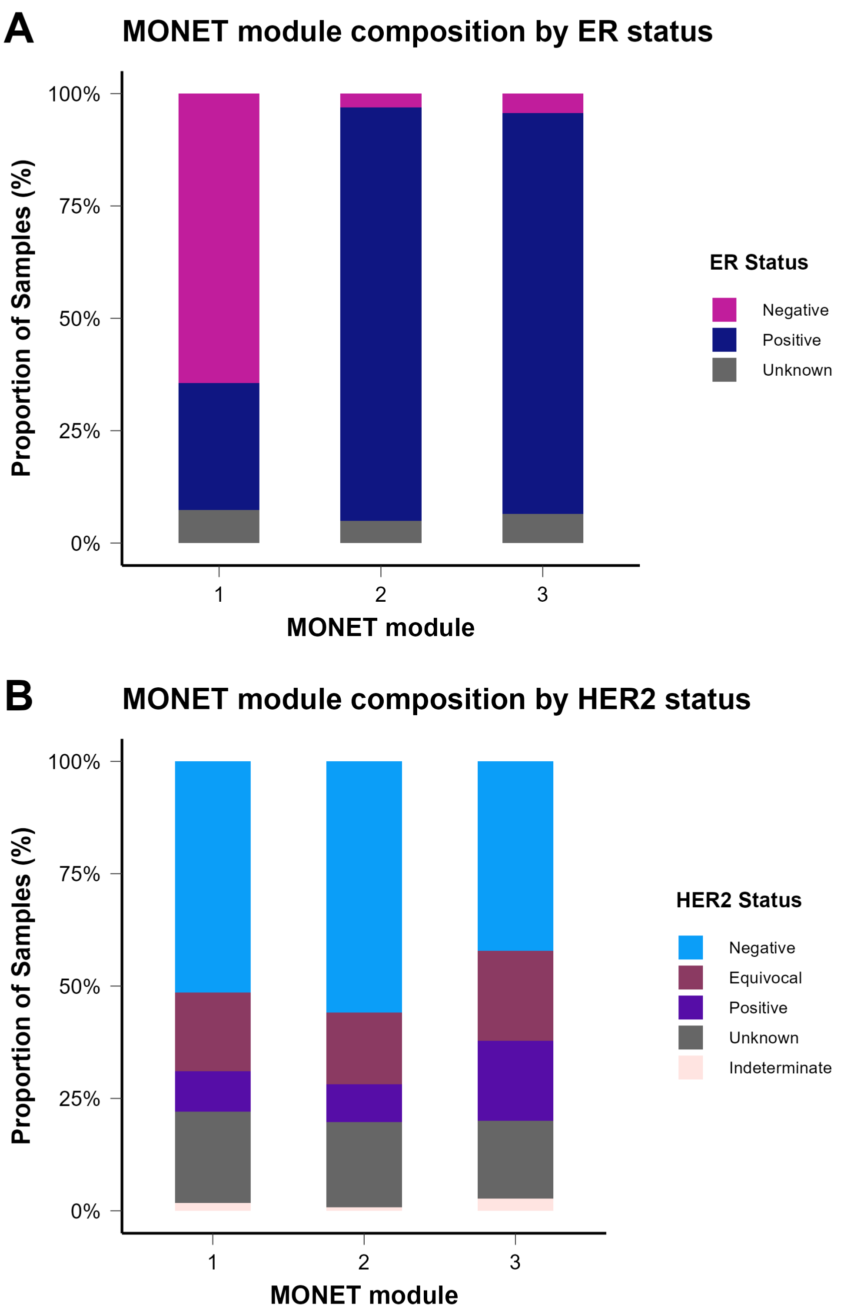

Figure 44. MONET module composition.

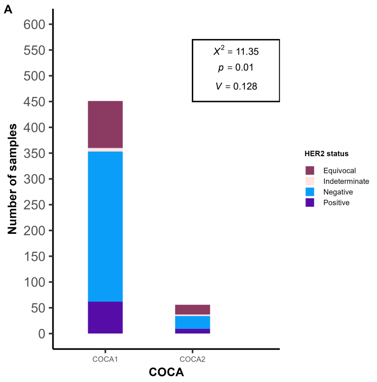

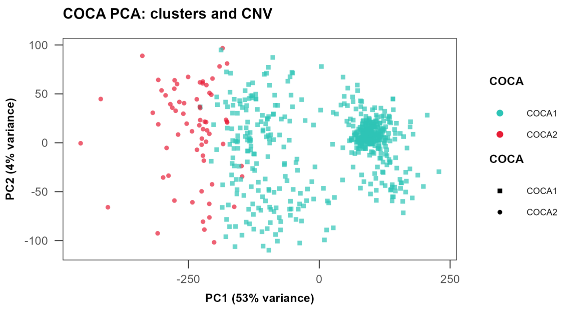

**(a)** **(b)**

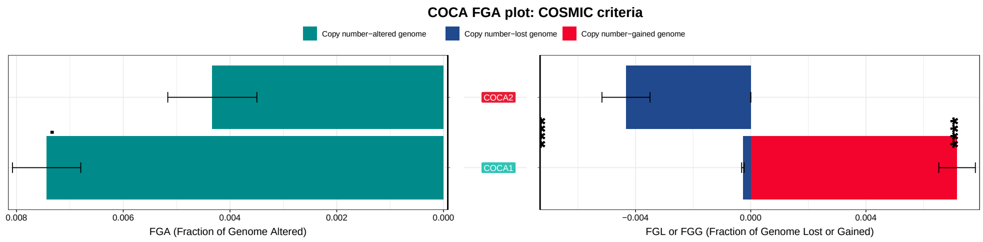

**(c)**

**Figure 45. COCA results:** (a) HER2 status bar chart, (b) CNV PCA plot and (c) FGA plot.

### 9 Final consensus pipeline

#### 9.1 Consensus for all methods

In this consensus approach, results from all methods are combined. Figure 46 presents the ER status bar chart, the silhouette scores, the FGA patterns and the oncoprint of the consensus subtypes Consensus Cluster (CC)1 and CC2. Figure 47 presents the up-regulated pathways in the two clusters.

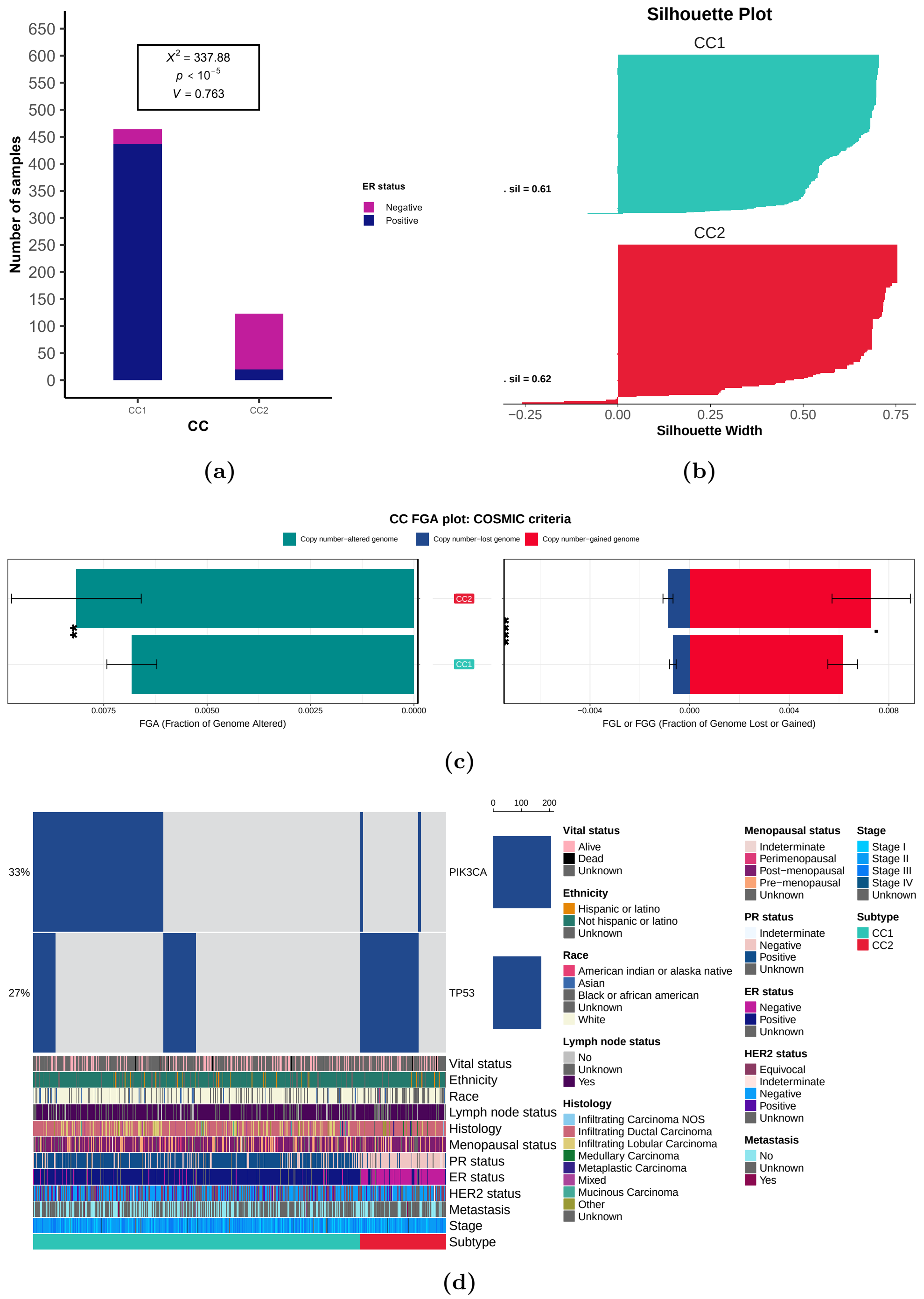

**Figure 46. CC results:** (a) ER status bar chart, (b) silhouette scores, (c) FGA plot and (d) oncoprint

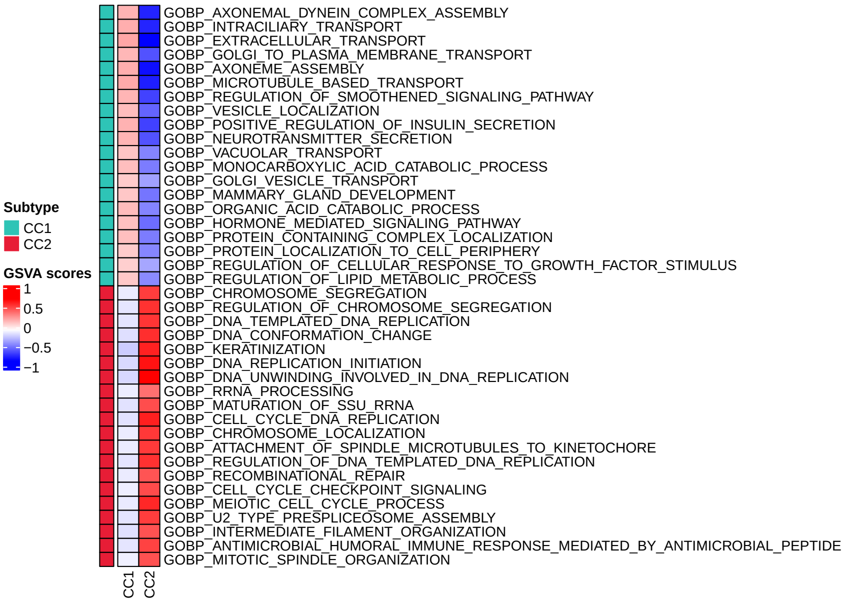

**Figure 47. Up-regulated pathways in CC clusters.**

#### 9.2 Consensus for methods with $k > 2$

In this consensus approach, results from methods which yielded more than 2 clusters are combined. Figure 48 presents the ER status bar chart, the HER2 status bar chart, the FGA patterns and the oncoprint of the consensus subtypes CC1 and CC2. Figure 49 presents the up-regulated pathways in the two clusters. Figure 50 is showing the dimensionality reduction plots for RNA-seq, CNV and methylation data, where cluster separation is more prominent.

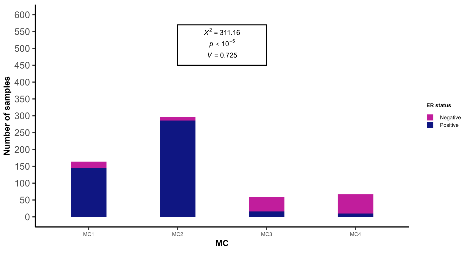

(a)

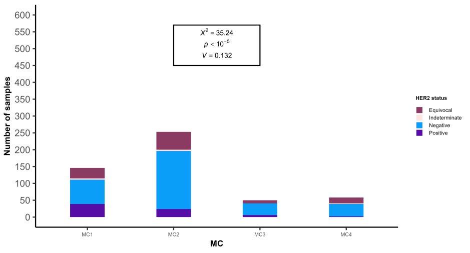

(b)

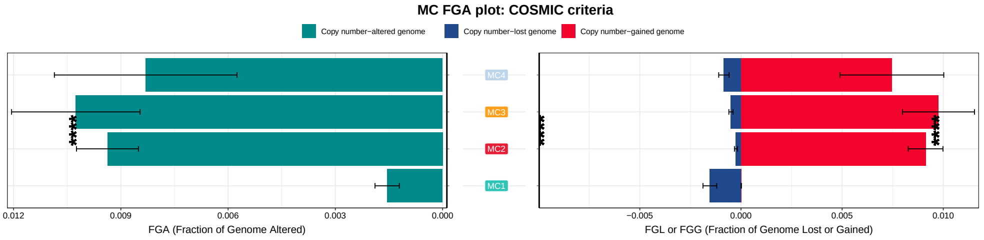

(c)

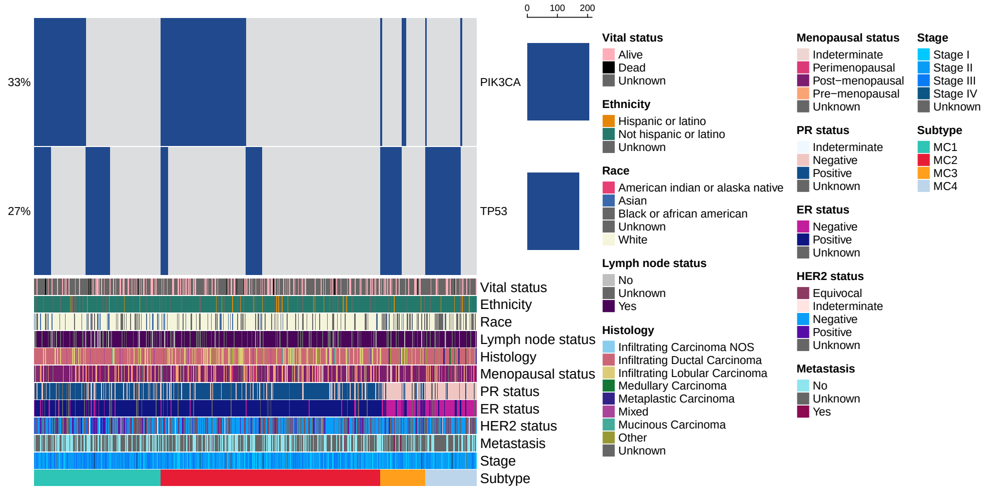

(d)

**Figure 48. MC results:** (a) ER status bar chart, (b) silhouette scores, (c) FGA plot and (d) oncoprint

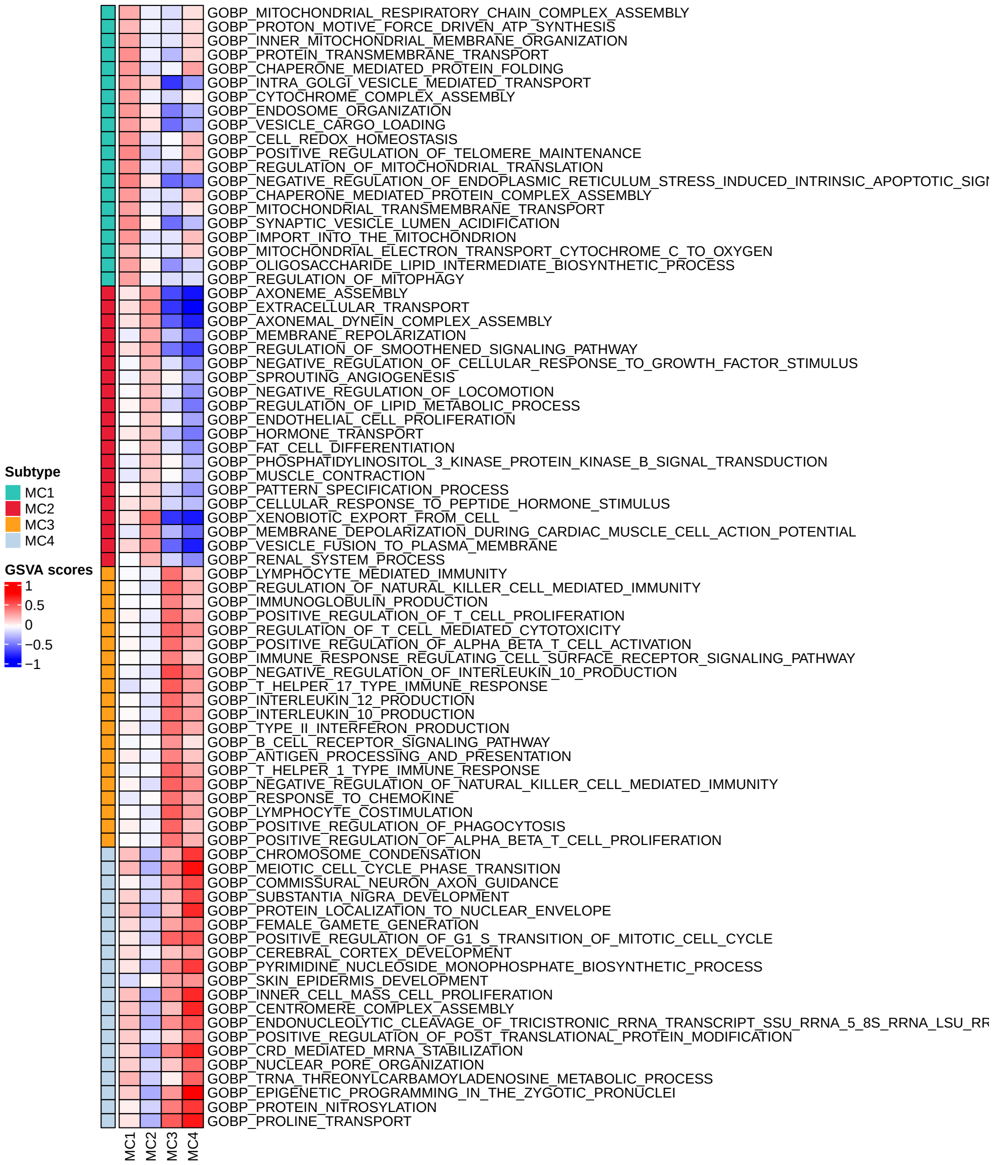

**Figure 49. Up-regulated pathways in MC clusters.**

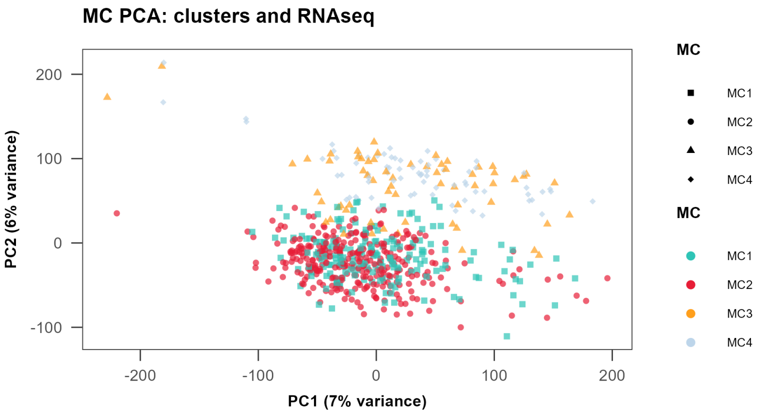

**(a)**

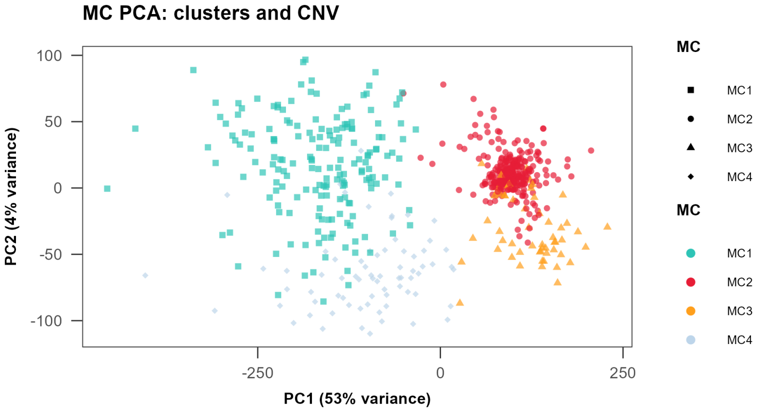

**(b)**

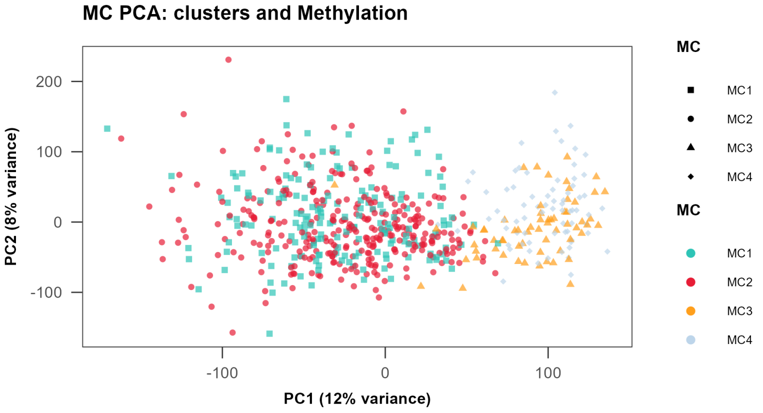

**(c)**

**Figure 50. MC dimensionality reduction plots:** (a) RNA-seq data PCA, (b) CNV data PCA and (c) methylation data PCA

### 10 Benchmarks

MFA, RWR-NF and MOFA are excluded from the benchmarks runs. MFA used a subset of the input in the poriginal runs and therefore cannot be directly compared to other methods, because both larger and samller subsets are needed and that would require extensive memory and in terms of runtime, it would extend beyond the 12h limit. RWR-NF benchmark runs failed to stay within the 12h limit, so the corresponding results were excluded as they were not complete for all subsets. This essentially ranks the method as the second slowest (after MFA). MOFA runs on a GPU, which is a major advantage in terms of scalability and comparisons with Central Processing Unit (CPU)-based methods are not straightforward.

For all runs, we constrained execution to a single CPU per task (`--cpus-per-task=1`) and disabled multithreading to minimise variability due to parallelism (e.g., setting BLAS/OpenMP thread environment variables to 1 and using `srun --hint=nomultithread --cpu-bind=cores`). Runtime and peak memory were measured in a consistent way across methods by wrapping the *core* computation of each method (the primary fitting/training routine that produces the latent representation and/or clustering) with system-level profiling (e.g., `/usr/bin/time -v` for Max RSS and elapsed time, or an equivalent method-level timing where appropriate). Downstream post-processing steps (e.g., exporting results, harmonising sample IDs, and clustering of learned embeddings) were kept separate from the measured region when they were **not** part of the method's core optimisation, enabling fair comparisons of how computational cost changes as the number of samples ($n$) and/or features ($p$) varies.

#### 10.1 Robustness

To benchmark multi-omic clustering methods under controlled and comparable conditions with respect to robustness, we ran two families of perturbation experiments while keeping each method's hyperparameters fixed to the optimal hyperparameter values we identified by tuning each method earlier. ARI-measured agremeent with the results of the full dataset runs was used to estimate robustness (method results agreement under feature/sample perturbations).

In the *feature-perturbation* experiments, we reduced the number of features per modality by selecting the top-ranked features according to a precomputed, method-agnostic feature ranking and retaining fixed centiles of that ranking (10%, 20%, 50%, 75%, 90%); the same centile was applied independently within each modality so that all methods saw a consistent proportional reduction in $p$ across views.

In the *sample-perturbation* experiments, we reduced the number of samples by using a shared set of predefined sample subsets (fractions with replicates) so that every method was evaluated on identical sample selections for a given fraction/replicate pair. For each target sample fraction (e.g., 10%, 20%, 50%, 70%, 90%), we pre-generated multiple *replicate* subsets to capture sampling variability while keeping the sampling scheme identical across methods. We constructed a table `sample_subsets.tsv.gz` containing rows of the form `(fraction, replicate, sample_id)`. For each fraction we created a fixed number of replicates (10 replicates per fraction, which retained the original proportion of ER+ and ER- samples), where each replicate corresponds to one random draw (without replacement) of the required number of samples from the valid sample universe. This yields a grid of (fraction × replicate) subset definitions, and each benchmark task corresponds to exactly one such pair; thus, the total number of runs per method is $n_{\text{fractions}} \times n_{\text{replicates}}$ ($5 \times 10 = 50$), and every method is run on the *same* (fraction, replicate) subsets.

For feature experiments, ARI between produced clusterings for each centile subset were compared to the full dataset. For sample experiments, ARI values for each centile were calculated by taking the *median* ARI of all replicates for that centile.

#### 10.2 Resampling stability

We quantified resampling stability under *sample perturbations* at the 90% subset level by comparing replicate clusterings produced from independently resampled patient subsets. For each method, we loaded the cluster assignment files for all 90% centile replicates and computed the ARI for every replicate pair using only the intersection of sample identifiers shared by the two replicates. The resulting set of pairwise ARIs per method provides an empirical stability distribution, which we summarised using the median and interquartile range and visualised with violin and box plots across methods.

#### 10.3 Scalability

To characterise empirical scaling of runtime, we fit a separate power-law model for each method and perturbation type. For feature perturbations, we model runtime as a function of the effective number of features $p$; for sample perturbations, as a function of the effective number of samples $n$. Specifically, for each method we fit an ordinary least squares regression on the log–log scale,

$$\log(\text{Time}) = a + \alpha \log(p) \quad \text{(feature perturbations)}$$

$$\log(\text{Time}) = b + \beta \log(n) \quad \text{(sample perturbations)}$$

In Tables 4, 5, we report the fitted scaling exponent ($\alpha$ or $\beta$), its confidence interval from the linear model, and summarised complexity as $O(p^\alpha)$ or $O(n^\beta)$. For readability, if an exponent was close to a small integer (within a tolerance), we rounded and reported $O(1)$, $O(n)$, $O(n^2)$, etc.; otherwise we reported $O(n^\beta)$ or $O(p^\alpha)$ with the estimated exponent. We also report the $R^2$ of each model as a goodness-of-fit measure for each result.

**Table 4.** Power-law models for feature dimensionality.

| **Algorithm** | Intercept ($a$) | Coefficient ($\alpha$) | **R2** | $\mathcal{O}$-notation |
|---|---|---|---|---|
| ab-SNF | 1.541 | 0.004 | 0.252 | $\mathcal{O}(1)$ |
| ANF | -0.491 | -0.019 | 0.882 | $\mathcal{O}(1)$ |
| CIMLR | 4.345 | 0.217 | 0.983 | $\mathcal{O}(p^{0.22})$ |
| COCA | -11.051 | 1.605 | 0.996 | $\mathcal{O}(p^{1.61})$ |
| iClusterBayes | -2.901 | 1.06 | 1 | $\mathcal{O}(p)$ |
| KLIC | 4.932 | -0.013 | 0.012 | $\mathcal{O}(1)$ |
| LRAcluster | -7.993 | 1.231 | 0.999 | $\mathcal{O}(p^{1.23})$ |
| MDICC | -0.556 | 0.323 | 0.868 | $\mathcal{O}(p^{0.32})$ |
| MONET | 4.677 | 0.052 | 0.482 | $\mathcal{O}(1)$ |
| MSNE | 5.583 | -0.022 | 0.166 | $\mathcal{O}(1)$ |
| NEMO | -11.195 | 1.275 | 0.982 | $\mathcal{O}(p^{1.27})$ |
| RWR-F | 9.423 | 0.024 | 0.31 | $\mathcal{O}(1)$ |
| SNF | 1.733 | -0.004 | 0.166 | $\mathcal{O}(1)$ |
| Spectrum | 1.367 | 0.364 | 0.965 | $\mathcal{O}(p^{0.36})$ |
| wMKL | 6.608 | -0.111 | 0.063 | $\mathcal{O}(1)$ |

**Table 5.** Power-law models for sample size.

| **Algorithm** | Intercept ($b$) | Coefficient ($\beta$) | **R2** | $\mathcal{O}$-notation |
|---|---|---|---|---|
| ab-SNF | -11.851 | 2.079 | 0.998 | $\mathcal{O}(n^2)$ |
| ANF | -12.588 | 1.812 | 0.994 | $\mathcal{O}(n^{1.81})$ |
| CIMLR | 1.148 | 0.891 | 0.995 | $\mathcal{O}(n)$ |
| COCA | -4.692 | 1.949 | 0.999 | $\mathcal{O}(n^2)$ |
| iClusterBayes | 3.688 | 0.935 | 1 | $\mathcal{O}(n)$ |
| KLIC | -5.102 | 1.517 | 0.984 | $\mathcal{O}(n^{1.52})$ |
| LRAcluster | -3.036 | 1.493 | 0.999 | $\mathcal{O}(n^{1.49})$ |
| MDICC | -7.02 | 1.583 | 0.987 | $\mathcal{O}(n^{1.58})$ |
| MONET | -8.83 | 2.157 | 0.991 | $\mathcal{O}(n^{2.16})$ |
| MSNE | -3.157 | 1.306 | 0.992 | $\mathcal{O}(n^{1.31})$ |
| NEMO | -7.589 | 1.603 | 0.997 | $\mathcal{O}(n^{1.60})$ |
| RWR-F | -14.314 | 3.686 | 0.996 | $\mathcal{O}(n^{3.69})$ |
| SNF | -10.911 | 1.933 | 0.998 | $\mathcal{O}(n^2)$ |
| Spectrum | -5.044 | 1.668 | 0.998 | $\mathcal{O}(n^{1.67})$ |
| wMKL | -3.701 | 1.458 | 0.959 | $\mathcal{O}(n^{1.46})$ |

### 11 Appendix

#### 11.1 Running MONET

Modifications were made to the MONET source code to improve compatibility and performance. Specifically, the graph-building function `build_a_graph_from_similarity()`'s source code was altered by changing a `from_numpy_matrix` method call from the *networkx* library to `from_numpy_array`, thus avoiding errors. Additional modifications were introduced, for example, to avoid runtime errors associated with modifying dictionaries in-place instead of iterating through copies of them.

#### 11.2 Running MDICC

We follow the installation notes in the MDICC repository. After downloading and un-zipping, we place the folder under the first entry of `.libPaths()`. Then we set:

```
MDICC_dir <- paste0(.libPaths()[1], "/MDICC")
```

**Suppress verbose warnings produced when calling `MDICClabel()`.**

- Limit the number of physical cores detected by `joblib`:
  `Sys.setenv(LOKY_MAX_CPU_COUNT = 8)`
- Prevent the $k$-means / multiple-kernel learning memory-leak on Windows:
  `Sys.setenv(OMP_NUM_THREADS = 3)`

**Load the MDICC R sources and the companion `.dll`.**

```
library(Rcpp)
library(parallel)
library(Matrix)
source(file.path(MDICC_dir, "NetworkFusion.R"))
dyn.load(file.path(MDICC_dir, "projsplx_R.dll"))
```

**Configure the Python back-end through `reticulate`. For example:**

```
library(reticulate)
Sys.setenv(RETICULATE_PYTHON = "C:/ProgramData/anaconda3/python.exe")
use_python("C:/ProgramData/anaconda3/python.exe")
```

**Quick checks:** `py_config()` and `py_available()`

**Import the MDICC helper modules written in Python.**

```
source_python(file.path(MDICC_dir, "LocalAffinityMatrix.py"))
source_python(file.path(MDICC_dir, "score.py"))
source_python(file.path(MDICC_dir, "label.py"))
```

Then you can proceed with running MDICC.

#### 11.3 Running KLIC

##### Installing the `klic` dependency `Rmosek` on Windows

1. Install `rtools` if it is not already available.
2. Download the current MOSEK bundle for Windows from here and unzip it so that you obtain the folder `C:/Users/`*`username`*`/mosek`.
3. Extend the system `PATH` every time R starts by adding the lines

```
##  .Rprofile  -----
mosek_path <- "C:/Users/username/mosek/10.2/tools/platform/win64x86/bin"
paths <- strsplit(Sys.getenv("PATH"), .Platform$path.sep)[[1]]
if (!mosek_path %in% paths) {
  Sys.setenv(PATH = paste(c(mosek_path, paths),
                         collapse = .Platform$path.sep))
}
```

4. Restart R or RStudio and execute

```
source("C:/Users/username/mosek/10.2/tools/platform/win64x86/rmosek/builder.R")
attachbuilder()
install.rmosek()
```

5. If MOSEK asks for a licence, request an academic key here and place the licence file inside `C:/Users/username/mosek`.

#### 11.4 wMKL installation error

The issue stems from the tsne.cpp file, line 883 (see here). More specifically, there is an argument mismatch: the dgemm_ call in tsne.cpp at line 883 is missing arguments compared to the definition in the computer's include/R_ext/BLAS.h file (lines 217-227).

The problem arises because on Windows (and certain other platforms), the BLAS routine dgemm called from C/C++ requires two extra hidden parameters for the Fortran string lengths. In other words, the function signature in R_ext/BLAS.h expects 15 arguments (13 explicit plus 2 string-length integers), but the initial call in tsne.cpp was only providing 13. To fix this, we explicitly supply FCONE for each string argument when calling F77_CALL(dgemm).

We also include at the top of tsne.cpp:

```
extern "C" {
  #include <R_ext/BLAS.h>
}
```

early in the file, ensuring the correct Fortran prototypes were in scope. Thus, changing the preamble and calling

```
F77_CALL(dgemm)(
   &transT, &transN,
   &N, &N, &D,
   &a1, X, &D,
        X, &D,
   &a2, DD, &N
   FCONE FCONE
);
```

eliminated the argument mismatch error and allowed the package to install successfully.

The summary of the steps we take to install the library are:

1. git clone to a directory (other than the R libraries directory)
2. Fix the tsne.cpp code as discussed above
3. Install the package from the local source, by navigating to the directory in R and running
   install.packages("wMKL_source", repos = NULL, type = "source")

[20] Koboldt, D.C., Fulton, R.S., McLellan, M.D., Schmidt, H., Kalicki-Veizer, J., McMichael, J.F., Fulton, L.L., Dooling, D.J., Ding, L., Mardis, E.R., Wilson, R.K., Ally, A., Balasundaram, M., Butterfield, Y.S.N., Carlsen, R., Carter, C., Chu, A., Chuah, E., Chun, H.-J.E., Coope, R.J.N., Dhalla, N., Guin, R., Hirst, C., Hirst, M., Holt, R.A., Lee, D., Li, H.I., Mayo, M., Moore, R.A., Mungall, A.J., Pleasance, E., Gordon Robertson, A., Schein, J.E., Shafiei, A., Sipahimalani, P., Slobodan, J.R., Stoll, D., Tam, A., Thiessen, N., Varhol, R.J., Wye, N., Zeng, T., Zhao, Y., Birol, I., Jones, S.J.M., Marra, M.A., Cherniack, A.D., Saksena, G.,

Onofrio, R.C., Pho, N.H., Carter, S.L., Schumacher, S.E., Tabak, B., Hernandez, B., Gentry, J., Nguyen, H., Crenshaw, A., Ardlie, K., Beroukhim, R., Winckler, W., Getz, G., Gabriel, S.B., Meyerson, M., Chin, L., Park, P.J., Kucherlapati, R., Hoadley, K.A., Todd Auman, J., Fan, C., Turman, Y.J., Shi, Y., Li, L., Topal, M.D., He, X., Chao, H.-H., Prat, A., Silva, G.O., Iglesia, M.D., Zhao, W., Usary, J., Berg, J.S., Adams, M., Booker, J., Wu, J., Gulabani, A., Bodenheimer, T., Hoyle, A.P., Simons, J.V., Soloway, M.G., Mose, L.E., Jefferys, S.R., Balu, S., Parker, J.S., Neil Hayes, D., Perou, C.M., Malik, S., Mahurkar, S., Shen, H., Weisenberger, D.J., Triche Jr, T., Lai, P.H., Bootwalla, M.S., Maglinte, D.T., Berman, B.P., Van Den Berg, D.J., Baylin, S.B., Laird, P.W., Creighton, C.J., Donehower, L.A., Noble, M., Voet, D., Gehlenborg, N., DiCara, D., Zhang, J., Zhang, H., Wu, C.-J., Yingchun Liu, S., Lawrence, M.S., Zou, L., Sivachenko, A., Lin, P., Stojanov, P., Jing, R., Cho, J., Sinha, R., Park, R.W., Nazaire, M.-D., Robinson, J., Thorvaldsdottir, H., Mesirov, J., Reynolds, S., Kreisberg, R.B., Bernard, B., Bressler, R., Erkkila, T., Lin, J., Thorsson, V., Zhang, W., Shmulevich, I., Ciriello, G., Weinhold, N., Schultz, N., Gao, J., Cerami, E., Gross, B., Jacobsen, A., Sinha, R., Arman Aksoy, B., Antipin, Y., Reva, B., Shen, R., Taylor, B.S., Ladanyi, M., Sander, C., Anur, P., Spellman, P.T., Lu, Y., Liu, W., Verhaak, R.R.G., Mills, G.B., Akbani, R., Zhang, N., Broom, B.M., Casasent, T.D., Wakefield, C., Unruh, A.K., Baggerly, K., Coombes, K., Weinstein, J.N., Haussler, D., Benz, C.C., Stuart, J.M., Benz, S.C., Zhu, J., Szeto, C.C., Scott, G.K., Yau, C., Paull, E.O., Carlin, D., Wong, C., Sokolov, A., Thusberg, J., Mooney, S., Ng, S., Goldstein, T.C., Ellrott, K., Grifford, M., Wilks, C., Ma, S., Craft, B., Yan, C., Hu, Y., Meerzaman, D., Gastier-Foster, J.M., Bowen, J., Ramirez, N.C., Black, A.D., Pyatt, R.E., White, P., Zmuda, E.J., Frick, J., Lichtenberg, T.M., Brookens, R., George, M.M., Gerken, M.A., Harper, H.A., Leraas, K.M., Wise, L.J., Tabler, T.R., McAllister, C., Barr, T., Hart-Kothari, M., Tarvin, K., Saller, C., Sandusky, G., Mitchell, C., Iacocca, M.V., Brown, J., Rabeno, B., Czerwinski, C., Petrelli, N., Dolzhansky, O., Abramov, M., Voronina, O., Potapova, O., Marks, J.R., Suchorska, W.M., Murawa, D., Kycler, W., Ibbs, M., Korski, K., Spychała, A., Murawa, P., Brzeziński, J.J., Perz, H., Łaźniak, R., Teresiak, M., Tatka, H., Leporowska, E., Bogusz-Czerniewicz, M., Malicki, J., Mackiewicz, A., Wiznerowicz, M., Van Le, X., Kohl, B., Viet Tien, N., Thorp, R., Van Bang, N., Sussman, H., Duc Phu, B., Hajek, R., Phi Hung, N., Viet The Phuong, T., Quyet Thang, H., Zaki Khan, K., Penny, R., Mallery, D., Curley, E., Shelton, C., Yena, P., Ingle, J.N., Couch, F.J., Lingle, W.L., King, T.A., Maria Gonzalez-Angulo, A., Dyer, M.D., Liu, S., Meng, X., Patangan, M., Network, T.C.G.A., St Louis, G.s.c.W.U., BC Cancer Agency, G., Institute, B., & Harvard Medical School, B.&.W.H., North Carolina, C.H., Southern California/Johns Hopkins, U., Medicine, G.d.a.B.C., Systems Biology, I., Center, M.S.-K.C., & Science University, O.H., Texas MD Anderson Cancer Center, T.U., California, S.C.I., NCI, Nationwide Children's Hospital Biospecimen Core Resource, B., ABS-IUPUI, T., Christiana, Cureline, Center, D.U.M., Centre, T.G.P.C., ILSBio, Consortium, I.G., Clinic, M., MSKCC, Center, M.A.C.: Comprehensive molecular portraits of human breast tumours. Nature **490**(7418), 61–70 (2012) https://doi.org/10.1038/nature11412

[26] Mo, Q., Shen, R.: iClusterPlus: Integrative Clustering of Multi-type Genomic Data. (2024). R package version 1.40.0

[46] Nguyen, T., Tagett, R., Diaz, D., Draghici, S.: A novel approach for data integration and disease subtyping. Genome Research (2017)

[47] Nguyen, H., Shrestha, S., Draghici, S., Nguyen, T.: Pinsplus: a tool for tumor subtype discovery in integrated genomic data. Bioinformatics **35**(16), 2843–2846 (2018) https://doi.org/10.1093/bioinformatics/bty1049

[48] Nguyen, H., Tran, D., Tran, B., Roy, M., Cassell, A., Dascalu, S., Draghici, S., Nguyen, T.: Smrt: Randomized data transformation for cancer subtyping and big data analysis. Frontiers in oncology (2021)

[61] Taylor, A.M., Shih, J., Ha, G., Gao, G.F., Zhang, X., Berger, A.C., Schumacher, S.E., Wang, C., Hu, H., Liu, J., Lazar, A.J., Cherniack, A.D., Beroukhim, R., Meyerson, M., Caesar-Johnson, S.J., Demchok, J.A., Felau, I., Kasapi, M., Ferguson, M.L., Hutter, C.M., Sofia, H.J., Tarnuzzer, R., Wang, Z., Yang, L., Zenklusen, J.C., Zhang, J.J., Chudamani, S., Liu, J., Lolla, L., Naresh, R., Pihl, T., Sun, Q., Wan, Y., Wu, Y., Cho, J., DeFreitas, T., Frazer, S., Gehlenborg, N., Getz, G., Heiman, D.I., Kim, J., Lawrence, M.S., Lin, P., Meier, S., Noble, M.S., Saksena, G., Voet, D., Zhang, H., Bernard, B., Chambwe, N., Dhankani,

V., Knijnenburg, T., Kramer, R., Leinonen, K., Liu, Y., Miller, M., Reynolds, S., Shmulevich, I., Thorsson, V., Zhang, W., Akbani, R., Broom, B.M., Hegde, A.M., Ju, Z., Kanchi, R.S., Korkut, A., Li, J., Liang, H., Ling, S., Liu, W., Lu, Y., Mills, G.B., Ng, K.-S., Rao, A., Ryan, M., Wang, J., Weinstein, J.N., Zhang, J., Abeshouse, A., Armenia, J., Chakravarty, D., Chatila, W.K., Bruijn, I., Gao, J., Gross, B.E., Heins, Z.J., Kundra, R., La, K., Ladanyi, M., Luna, A., Nissan, M.G., Ochoa, A., Phillips, S.M., Reznik, E., Sanchez-Vega, F., Sander, C., Schultz, N., Sheridan, R., Sumer, S.O., Sun, Y., Taylor, B.S., Wang, J., Zhang, H., Anur, P., Peto, M., Spellman, P., Benz, C., Stuart, J.M., Wong, C.K., Yau, C., Hayes, D.N., Parker, J.S., Wilkerson, M.D., Ally, A., Balasundaram, M., Bowlby, R., Brooks, D., Carlsen, R., Chuah, E., Dhalla, N., Holt, R., Jones, S.J.M., Kasaian, K., Lee, D., Ma, Y., Marra, M.A., Mayo, M., Moore, R.A., Mungall, A.J., Mungall, K., Robertson, A.G., Sadeghi, S., Schein, J.E., Sipahimalani, P., Tam, A., Thiessen, N., Tse, K., Wong, T., Berger, A.C., Beroukhim, R., Cherniack, A.D., Cibulskis, C., Gabriel, S.B., Gao, G.F., Ha, G., Meyerson, M., Schumacher, S.E., Shih, J., Kucherlapati, M.H., Kucherlapati, R.S., Baylin, S., Cope, L., Danilova, L., Bootwalla, M.S., Lai, P.H., Maglinte, D.T., Van Den Berg, D.J., Weisenberger, D.J., Auman, J.T., Balu, S., Bodenheimer, T., Fan, C., Hoadley, K.A., Hoyle, A.P., Jefferys, S.R., Jones, C.D., Meng, S., Mieczkowski, P.A., Mose, L.E., Perou, A.H., Perou, C.M., Roach, J., Shi, Y., Simons, J.V., Skelly, T., Soloway, M.G., Tan, D., Veluvolu, U., Fan, H., Hinoue, T., Laird, P.W., Shen, H., Zhou, W., Bellair, M., Chang, K., Covington, K., Creighton, C.J., Dinh, H., Doddapaneni, H., Donehower, L.A., Drummond, J., Gibbs, R.A., Glenn, R., Hale, W., Han, Y., Hu, J., Korchina, V., Lee, S., Lewis, L., Li, W., Liu, X., Morgan, M., Morton, D., Muzny, D., Santibanez, J., Sheth, M., Shinbrot, E., Wang, L., Wang, M., Wheeler, D.A., Xi, L., Zhao, F., Hess, J., Appelbaum, E.L., Bailey, M., Cordes, M.G., Ding, L., Fronick, C.C., Fulton, L.A., Fulton, R.S., Kandoth, C., Mardis, E.R., McLellan, M.D., Miller, C.A., Schmidt, H.K., Wilson, R.K., Crain, D., Curley, E., Gardner, J., Lau, K., Mallery, D., Morris, S., Paulauskis, J., Penny, R., Shelton, C., Shelton, T., Sherman, M., Thompson, E., Yena, P., Bowen, J., Gastier-Foster, J.M., Gerken, M., Leraas, K.M., Lichtenberg, T.M., Ramirez, N.C., Wise, L., Zmuda, E., Corcoran, N., Costello, T., Hovens, C., Carvalho, A.L., Carvalho, A.C., Fregnani, J.H., Longatto-Filho, A., Reis, R.M., Scapulatempo-Neto, C., Silveira, H.C.S., Vidal, D.O., Burnette, A., Eschbacher, J., Hermes, B., Noss, A., Singh, R., Anderson, M.L., Castro, P.D., Ittmann, M., Huntsman, D., Kohl, B., Le, X., Thorp, R., Andry, C., Duffy, E.R., Lyadov, V., Paklina, O., Setdikova, G., Shabunin, A., Tavobilov, M., McPherson, C., Warnick, R., Berkowitz, R., Cramer, D., Feltmate, C., Horowitz, N., Kibel, A., Muto, M., Raut, C.P., Malykh, A., Barnholtz-Sloan, J.S., Barrett, W., Devine, K., Fulop, J., Ostrom, Q.T., Shimmel, K., Wolinsky, Y., Sloan, A.E., De Rose, A., Giuliante, F., Goodman, M., Karlan, B.Y., Hagedorn, C.H., Eckman, J., Harr, J., Myers, J., Tucker, K., Zach, L.A., Deyarmin, B., Hu, H., Kvecher, L., Larson, C., Mural, R.J., Somiari, S., Vicha, A., Zelinka, T., Bennett, J., Iacocca, M., Rabeno, B., Swanson, P., Latour, M., Lacombe, L., Têtu, B., Bergeron, A., McGraw, M., Staugaitis, S.M., Chabot, J., Hibshoosh, H., Sepulveda, A., Su, T., Wang, T., Potapova, O., Voronina, O.,

Desjardins, L., Mariani, O., Roman-Roman, S., Sastre, X., Stern, M.-H., Cheng, F., Signoretti, S., Berchuck, A., Bigner, D., Lipp, E., Marks, J., McCall, S., McLendon, R., Secord, A., Sharp, A., Behera, M., Brat, D.J., Chen, A., Delman, K., Force, S., Khuri, F., Magliocca, K., Maithel, S., Olson, J.J., Owonikoko, T., Pickens, A., Ramalingam, S., Shin, D.M., Sica, G., Van Meir, E.G., Zhang, H., Eijckenboom, W., Gillis, A., Korpershoek, E., Looijenga, L., Oosterhuis, W., Stoop, H., Kessel, K.E., Zwarthoff, E.C., Calatozzolo, C., Cuppini, L., Cuzzubbo, S., DiMeco, F., Finocchiaro, G., Mattei, L., Perin, A., Pollo, B., Chen, C., Houck, J., Lohavanichbutr, P., Hartmann, A., Stoehr, C., Stoehr, R., Taubert, H., Wach, S., Wullich, B., Kycler, W., Murawa, D., Wiznerowicz, M., Chung, K., Edenfield, W.J., Martin, J., Baudin, E., Bubley, G., Bueno, R., De Rienzo, A., Richards, W.G., Kalkanis, S., Mikkelsen, T., Noushmehr, H., Scarpace, L., Girard, N., Aymerich, M., Campo, E., Giné, E., Guillermo, A.L., Van Bang, N., Hanh, P.T., Phu, B.D., Tang, Y., Colman, H., Evason, K., Dottino, P.R., Martignetti, J.A., Gabra, H., Juhl, H., Akeredolu, T., Stepa, S., Hoon, D., Ahn, K., Kang, K.J., Beuschlein, F., Breggia, A., Birrer, M., Bell, D., Borad, M., Bryce, A.H., Castle, E., Chandan, V., Cheville, J., Copland, J.A., Farnell, M., Flotte, T., Giama, N., Ho, T., Kendrick, M., Kocher, J.-P., Kopp, K., Moser, C., Nagorney, D., O'Brien, D., O'Neill, B.P., Patel, T., Petersen, G., Que, F., Rivera, M., Roberts, L., Smallridge, R., Smyrk, T., Stanton, M., Thompson, R.H., Torbenson, M., Yang, J.D., Zhang, L., Brimo, F., Ajani, J.A., Angulo Gonzalez, A.M., Behrens, C., Bondaruk, J., Broaddus, R., Czerniak, B., Esmaeli, B., Fujimoto, J., Gershenwald, J., Guo, C., Lazar, A.J., Logothetis, C., Meric-Bernstam, F., Moran, C., Ramondetta, L., Rice, D., Sood, A., Tamboli, P., Thompson, T., Troncoso, P., Tsao, A., Wistuba, I., Carter, C., Haydu, L., Hersey, P., Jakrot, V., Kakavand, H., Kefford, R., Lee, K., Long, G., Mann, G., Quinn, M., Saw, R., Scolyer, R., Shannon, K., Spillane, A., Stretch, J., Synott, M., Thompson, J., Wilmott, J., Al-Ahmadie, H., Chan, T.A., Ghossein, R., Gopalan, A., Levine, D.A., Reuter, V., Singer, S., Singh, B., Tien, N.V., Broudy, T., Mirsaidi, C., Nair, P., Drwiega, P., Miller, J., Smith, J., Zaren, H., Park, J.-W., Hung, N.P., Kebebew, E., Linehan, W.M., Metwalli, A.R., Pacak, K., Pinto, P.A., Schiffman, M., Schmidt, L.S., Vocke, C.D., Wentzensen, N., Worrell, R., Yang, H., Moncrieff, M., Goparaju, C., Melamed, J., Pass, H., Botnariuc, N., Caraman, I., Cernat, M., Chemencedji, I., Clipca, A., Doruc, S., Gorincioi, G., Mura, S., Pirtac, M., Stancul, I., Tcaciuc, D., Albert, M., Alexopoulou, I., Arnaout, A., Bartlett, J., Engel, J., Gilbert, S., Parfitt, J., Sekhon, H., Thomas, G., Rassl, D.M., Rintoul, R.C., Bifulco, C., Tamakawa, R., Urba, W., Hayward, N., Timmers, H., Antenucci, A., Facciolo, F., Grazi, G., Marino, M., Merola, R., Krijger, R., Gimenez-Roqueplo, A.-P., Piché, A., Chevalier, S., McKercher, G., Birsoy, K., Barnett, G., Brewer, C., Farver, C., Naska, T., Pennell, N.A., Raymond, D., Schilero, C., Smolenski, K., Williams, F., Morrison, C., Borgia, J.A., Liptay, M.J., Pool, M., Seder, C.W., Junker, K., Omberg, L., Dinkin, M., Manikhas, G., Alvaro, D., Bragazzi, M.C., Cardinale, V., Carpino, G., Gaudio, E., Chesla, D., Cottingham, S., Dubina, M., Moiseenko, F., Dhanasekaran, R., Becker, K.-F., Janssen, K.-P., Slotta-Huspenina, J., Abdel-Rahman, M.H., Aziz, D., Bell, S., Cebulla, C.M., Davis, A., Duell, R., Elder, J.B.,

Hilty, J., Kumar, B., Lang, J., Lehman, N.L., Mandt, R., Nguyen, P., Pilarski, R., Rai, K., Schoenfield, L., Senecal, K., Wakely, P., Hansen, P., Lechan, R., Powers, J., Tischler, A., Grizzle, W.E., Sexton, K.C., Kastl, A., Henderson, J., Porten, S., Waldmann, J., Fassnacht, M., Asa, S.L., Schadendorf, D., Couce, M., Graefen, M., Huland, H., Sauter, G., Schlomm, T., Simon, R., Tennstedt, P., Olabode, O., Nelson, M., Bathe, O., Carroll, P.R., Chan, J.M., Disaia, P., Glenn, P., Kelley, R.K., Landen, C.N., Phillips, J., Prados, M., Simko, J., Smith-McCune, K., VandenBerg, S., Roggin, K., Fehrenbach, A., Kendler, A., Sifri, S., Steele, R., Jimeno, A., Carey, F., Forgie, I., Mannelli, M., Carney, M., Hernandez, B., Campos, B., Herold-Mende, C., Jungk, C., Unterberg, A., Deimling, A., Bossler, A., Galbraith, J., Jacobus, L., Knudson, M., Knutson, T., Ma, D., Milhem, M., Sigmund, R., Godwin, A.K., Madan, R., Rosenthal, H.G., Adebamowo, C., Adebamowo, S.N., Boussioutas, A., Beer, D., Giordano, T., Mes-Masson, A.-M., Saad, F., Bocklage, T., Landrum, L., Mannel, R., Moore, K., Moxley, K., Postier, R., Walker, J., Zuna, R., Feldman, M., Valdivieso, F., Dhir, R., Luketich, J., Mora Pinero, E.M., Quintero-Aguilo, M., Carlotti, C.G., Dos Santos, J.S., Kemp, R., Sankarankuty, A., Tirapelli, D., Catto, J., Agnew, K., Swisher, E., Creaney, J., Robinson, B., Shelley, C.S., Godwin, E.M., Kendall, S., Shipman, C., Bradford, C., Carey, T., Haddad, A., Moyer, J., Peterson, L., Prince, M., Rozek, L., Wolf, G., Bowman, R., Fong, K.M., Yang, I., Korst, R., Rathmell, W.K., Fantacone-Campbell, J.L., Hooke, J.A., Kovatich, A.J., Shriver, C.D., DiPersio, J., Drake, B., Govindan, R., Heath, S., Ley, T., Van Tine, B., Westervelt, P., Rubin, M.A., Lee, J.I., Aredes, N.D., Mariamidze, A.: Genomic and functional approaches to understanding cancer aneuploidy. Cancer Cell **33**(4), 676–6893 (2018) https://doi.org/10.1016/j.ccell.2018.03.007

[68] Hastie, T., Tibshirani, R., Narasimhan, B., Chu, G.: Impute: Imputation for Microarray Data. (2024). R package version 1.78.0

[89] Garnett, M.J., Edelman, E.J., Heidorn, S.J., Greenman, C.D., Dastur, A., Lau, K.W., Greninger, P., Thompson, I.R., Luo, X., Soares, J., Liu, Q., Iorio, F., Surdez, D., Chen, L., Milano, R.J., Bignell, G.R., Tam, A.T., Davies, H., Stevenson, J.A., Barthorpe, S., Lutz, S.R., Kogera, F., Lawrence, K., McLaren-Douglas, A., Mitropoulos, X., Mironenko, T., Thi, H., Richardson, L., Zhou, W., Jewitt, F., Zhang, T., O'Brien, P., Boisvert, J.L., Price, S., Hur, W., Yang, W., Deng, X., Butler, A., Choi, H.G., Chang, J.W., Baselga, J., Stamenkovic, I., Engelman, J.A., Sharma, S.V., Delattre, O., Saez-Rodriguez, J., Gray, N.S., Settleman, J., Futreal, P.A., Haber, D.A., Stratton, M.R., Ramaswamy, S., McDermott, U., Benes, C.H.: Systematic identification of genomic markers of drug sensitivity in cancer cells. Nature **483**(7391), 570–575 (2012) https://doi.org/10.1038/nature11005

[103] Meng, C.: Mogsa: Multiple Omics Data Integrative Clustering and Gene Set Analysis. (2024). R package version 1.38.0

[104] Boutsidis, C., Gallopoulos, E.: Svd based initialization: A head start for non-negative matrix factorization. Pattern Recognition **41**(4), 1350–1362 (2008) https://doi.org/10.1016/j.patcog.2007.09.010
